# Genome-Wide Association Insights into the Genomic Regions Controlling Oil Production Traits in *Acrocomia aculeata* (neotropical native palm)

**DOI:** 10.1101/2024.01.17.576138

**Authors:** Evellyn Giselly de Oliveira Couto, Jonathan Morales-Marroquín, Alessandro Alves-Pereira, Samuel B. Fernandes, Carlos Augusto Colombo, Joaquim Adelino de Azevedo Filho, Cassia Regina Limonta Carvalho, Maria Imaculada Zucchi

## Abstract

Macauba (*Acrocomia aculeata*) is a non-domesticated neotropical palm that has been attracting attention for economical use due to its great potential for oil production comparable to the commercially used oil palm (*Elaeis guineenses*). The discovery of associations between quantitative trait loci and economically important traits represents an advance toward macauba domestication. Pursuing this advance, this study performs single-trait and multi-trait GWAS models to identify candidate genes related to oil production traits in macauba. We randomly selected 201 palms from a natural population and analysed 13 traits related to fruit production, processing, and oil content. Genotyping was performed following the genotyping-by-sequencing protocol. SNP calling was performed using three strategies since macauba doesn’t have a reference genome: using i) de novo pipeline, ii) *Elaeis guineenses* Jacq. reference genome, and iii) transcriptome of *Acrocomia aculeata*. Single-trait analysis was fitted using five models from GAPIT, while multi-trait analysis was fitted using a multivariate stepwise method implemented in the software TASSEL. Multi-trait analyses were conducted in all pairwise trait combinations. Results showed statistically significant differences in all phenotypic traits studied, and heritability values ranged from 0.63 to 0.95. Gene annotation detected 15 candidate genes in seven traits in the single-trait GWAS and four candidate genes in 10 trait combinations in the multi-trait GWAS. We provide new insights on genomic regions that mapped candidate genes involved in macauba oil production phenotypes. Associated markers to the traits of interest may be valuable resources for the development of marker-assisted selection in macauba for both domestication and pre-breeding purposes.

## 1 INTRODUCTION

Macauba (*Acrocomia aculeata*) is a neotropical palm distributed from Mexico to the Northern region of Argentina. In Brazil, macauba occurs in the states of Ceará, Mato Grosso, Mato Grosso do Sul, Minas Gerais, Tocantins, Roraima, Amazonas, and São Paulo (Scariot et al., 1995; Lima et al., 2018; Lorenzi et al., 2010). The genus *Acrocomia* has great potential to become commercially exploited. Within the genus, macauba is the most targeted species due to the high oleic concentration in its fruits. As a perennial plant, this palm has a long-term, regular production. Furthermore, the species is adapted to growing on dry biomes, bare ground, rangeland, or anthropogenic landscapes. These features characterize macauba as a sustainable source of vegetable oil. Nowadays, vegetable oil can be extracted from many species, such as soybeans, sugar cane, and oil palm, among others, of which the palm macauba (*Acrocomia aculeata* (Jacq.) Lood. ex Mart) has stood out (Colombo et al., 2018). Macauba starts producing fruit around the fifth year on the field and guarantees production for up to 50 years, avoiding excessive soil management (Teixeira, 2005), as it would occur in annual species like soybean and sunflower. Macauba can be used in integrated Crop-Livestock-Forest systems or in Regenerative Land Management, optimizing its use and production gains.

In industry, all macauba fruit parts (epicarp, mesocarp, endocarp, and endosperm) can be used for raw material production. Oil is extracted from the mesocarp and endosperm. The mesocarp has up to 70% of an oleic acid-rich oil content (Ciconini et al., 2013) targeted in the biofuel industry. The endosperm has up to 50% of an oil content rich in lauric acid, which is used in cosmetics industries and other saponification products (Coimbra & Jorge, 2012). Furthermore, the fruits are abundant in amino acids, carbohydrates, and minerals (Hiane et al., 2005), and can be used in the animal and human food industry. Finally, the endocarp has high lignin content and could be used in coal production (Silva et al., 1986). Despite the commercial potential, this palm has not been domesticated (Clement, 1999; Clement *et al*., 2021), and most of the oil consumed is obtained from extractivism (Abreu et al., 2012).

In plant breeding, genetic diversity studies can guide the selection of contrasting parents in artificial crosses, maximizing the genetic gains and the efficiency in the development of new cultivars (Cruz et al. 2011; Farias Neto et al. 2013). Diaz et al. (2021) studied genetic diversity in *Acrocomia ssp*. distributed in Brazil and found high genetic variability. That study indicates that productive native palms can be selected to start a pre-breeding cycle. The selection of parental plants with potential for productivity in a natural population is based on their phenotypic traits, and the correlation estimates between its traits enable the selection and understanding of their genetic basis. Previous studies in macauba have observed high genotypic correlations between plant height and number of green, plant height and dry leaves, plant height and stipe thickness; as also between leaf length and stem length, and leaf length and stipe thickness (Berton, 2013). High genetic correlations between traits of interest can facilitate the breeding process once breeders can perform indirect selection, focusing on a few easily measured traits. This is possible because the existence of a high correlation between traits indicates pleiotropic and/or linked genes control them (Falconer & Mackay, 1996).

Agronomic traits that are controlled by multiple genes are qualified as quantitative traits. Quantitative Trait Loci (QTL) are widely used to name chromosomal regions that have genes controlling the expression of quantitative traits (Falconer & Mackay, 1996; Mackay, 2001). Molecular markers are used for mapping QTL for several traits, allowing a better understanding of the genetic expression of these traits (Falconer & Mackay, 1996; Malosetti et al., 2008). This can be directly applied to genetic improvement programs. QTL mapping studies can be based on genome-wide association studies (GWAS) (Oraguzie et al., 2007). GWAS attempts to evaluate causal relationships between genetic variants and the phenotype by surveying a genome-wide set of genetic polymorphisms in a large number of individuals (François & Caye, 2018). The association detected is attributed to linkage disequilibrium between markers and functional polymorphisms in a set of evaluated genotypes (Yu et al., 2006).

Single and multi-trait GWAS approaches are currently employed to detect cross-phenotype associations (Zhou and Stephens, 2014; Cichonska et al., 2016; Joo et al., 2016). Some of the first GWAS models, such as general linear models and mixed-linear models (MLMs), were single-locus and single-trait, created to implement covariates along with kinship matrices (Yu et al., 2006). These simple models resulted in false negatives caused by weakened associations due to population structure. To evaluate big datasets while also reducing false positives and negatives, multi-locus GWAS, such as FarmCPU (Liu et al., 2016) and BLINK (Huang et al., 2019), were developed. Unlike the single-trait models, multi-trait models enable the quantification of simultaneous loci contributions for multiple traits in GWAS studies (Zhou & Stephens, 2014). Many software packages have implemented multi-trait GWAS models (Furlotte & Eskin, 2015; Pritikin et al., 2021; Zhou & Stephens, 2014; Bradbury et al., 2007). Especially when analyzing low heritability traits, multi-trait models can outperform univariate multi-locus models (Fernandes et al., 2022).

To the best of our knowledge, there are no studies applying GWAS on macauba. Results related to the genomic association of traits responsible for oil production are of interest for understanding their genetic architecture, enabling the facilitation of the specie’s domestication and breeding. In addition, markers detected in the multi-trait GWAS can guide studies about the detection of linked or pleiotropic genes, which would enrich the genetic information about this palm. This study presents the first GWAS targeting genomic regions controlling oil production traits in macauba.

## 2 MATERIAL AND METHODS

### 2.1 Plant material and phenotyping

We studied a panel of 201 macauba palms from a natural population located in Dourado, São Paulo state, Brazil (geographical coordinates 22° 6’ 13’’ S, 48° 18’ 50’’ W). Palms were randomly selected in four subpopulations located in four rural áreas, approximately 500 meters apart (**Figure 1**). Data was collected in two different seasons in the years of 2019/2020 and 2020/2021. Subpopulation 1 is formed by individuals numbered 1 to 100, subpopulation 2 by individuals numbered 101 to 130, subpopulation by individuals numbered 131 to 184, and subpopulation 4 by individuals numbered 185 to 201.

**Figure 1.**
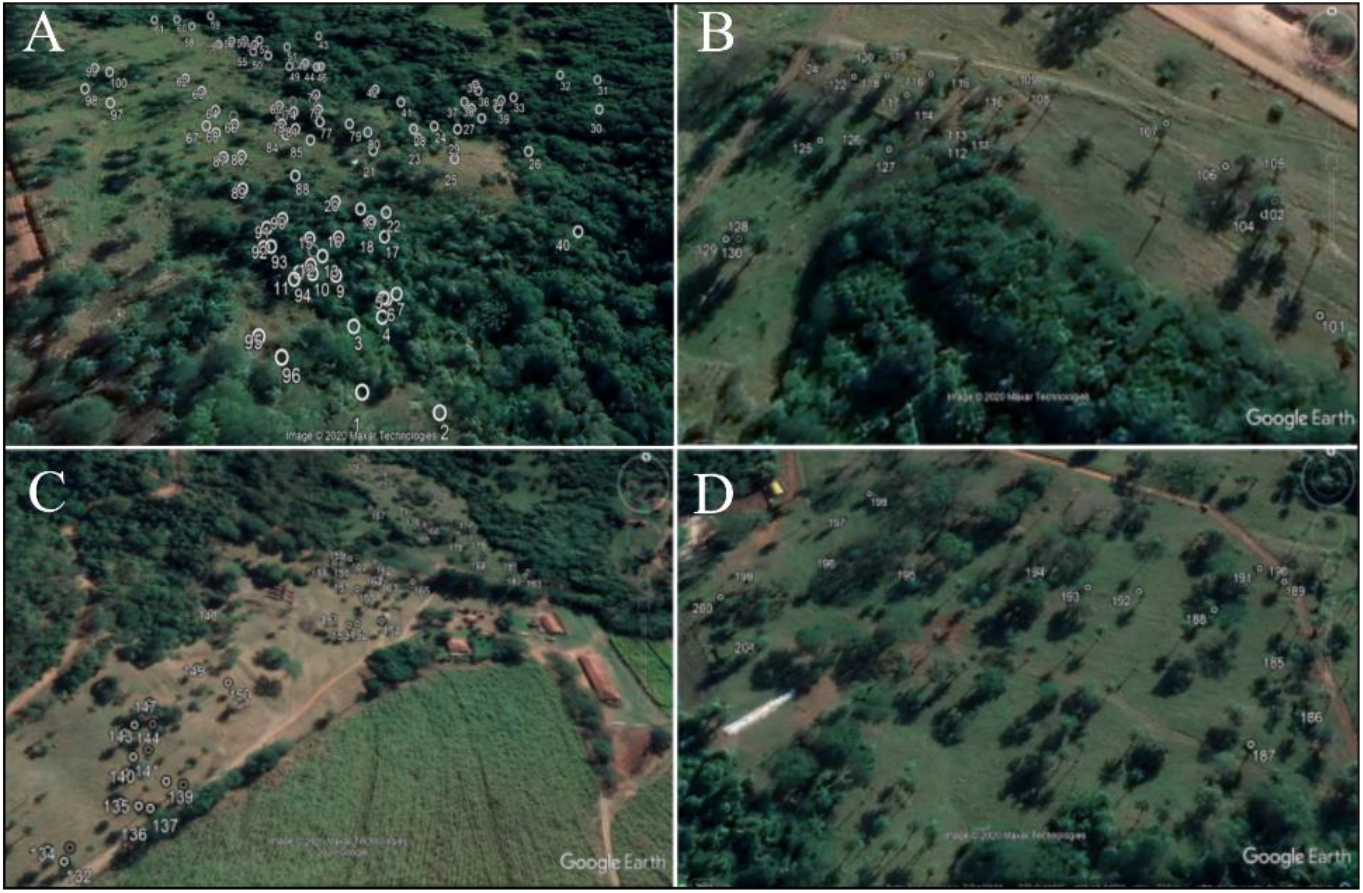
Rural areas where macauba palms were collected. A) Subpopulation 1. B) Subpopulation 2. C) Subpopulation 3. D) Subpopulation 4.

The measured phenotypes were: height (H - in meters), stipe (trunk) diameter at breast height (DHB - in centimeters), number of leaves (LN), leaves length (LL - in centimeters), number of leaf needles (NN), leaf needles length (NL - in centimeters), leaf needles width (NW - in centimeters) and total fruit mass (FM – in grams). The fruit characterization was based on six fruits from each palm (**Figure 2A**) in three repetitions (two fruits per repetition). After obtaining the total weight of the fruits, they were manually separated into four fractions: (1) epicarp (husk), (2) mesocarp (pulp), (3) endocarp (shell), and (4) endosperm (kernel) for fresh and dry mass (**Figure 2B**). After measuring the fresh masses, the fruit fractions were dried in a ventilated oven at 36°C for 36 hours. We identified these traits as: husk fresh mass (HFM), pulp fresh mass (PFM), endocarp fresh mass (EFM), kernel fresh mass (KFM), husk dry mass (HDM), pulp dry mass (PDM), endocarp dry mass (EDM) and kernel dry mass (KDM). The weights of the four dry fractions were added to obtain the value of the total mass of the dry fruit (FDM).

**Figure 2.**
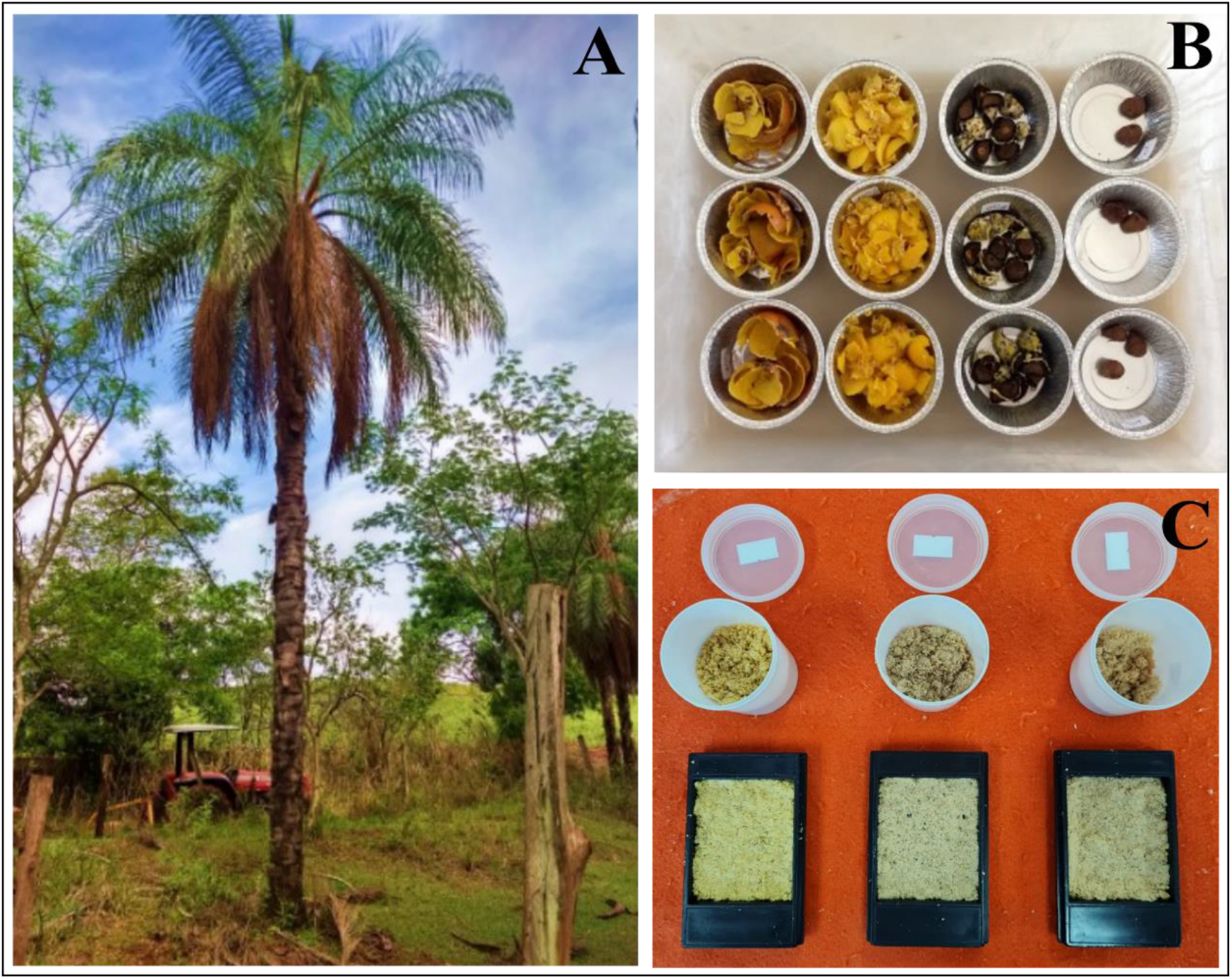
Plant material and phenotyping. A) Macauba in the field. B) Fruit fractions in three repetitions; considering from left to right: epicarp (husk), mesocarp (pulp), endocarp, and kernel. C) Different shades of dry mesocarp flours used in oil content quantification.

### 2.2 Oil content quantification by NIRS

Oil content (OC, % or g/100g) was quantified from the dry mesocarp mass (**Figure 2C**) using Near-infrared Spectroscopy (NIRS). The dry mesocarps of the three replicates were crushed in the analytical mil IKA–A11. Diffuse reflectance of the mesocarp samples was measured in rectangular cells (48 x 58 mm) using a FOSS NIRSystems 6500 spectrophotometer. The spectra measurements of the dry mesocarp mass were performed in triplicates and recorded using the ISIscanTM software, VERSION 3.10 (Infrasoft International, 2007). A multivariate model was built using the software Pirouette 4.5 (Infometrix, 1990-2011) to predict the oil content using the NIRs spectros graphs.

### 2.3 Genotypic dataset

Genotyping was performed with markers based on single-nucleotide polymorphism (SNPs), following the protocol of genotyping-by-sequencing using two restriction enzymes (ddGBS) (Poland et al., 2012). The genomic DNA was isolated from leaf material using the Doyle & Doyle (1990) protocol. We evaluated the extracted DNA quality and quantity by agarose gel electrophoresis (1% w/v) stained with SYBR Safe DNA Gel Stain (Invitrogen) and by visual comparison with lambda DNA (Invitrogen). The quantification and normalization of genomic DNA was performed through fluorescence using the Qubit dsDNA BR Assay (Qubit—Life Technologies). Based on the obtained reading, we standardized the DNA to a concentration of 30ng.μl- 1.

### 2.4 GBS library preparation and high-throughput sequencing

To obtain the SNPs, we prepared three 96-plex genomic libraries using the ddGBS technique according to the protocol described by Poland et al. (2012) and the modifications used by Díaz et al. (2021). We digested the genomic DNA with the combination of enzymes *NsiI* and *Mse1* (New England Biolabs). The ddGBS libraries were quantified through RT-PCR on the CFX 384 Touch Real-Time PCR (BioRad) equipment using a KAPA Library Quantification kit (KAPA Biosystems, cat. KK4824), and the fragments’ profiles were inspected using the Agilent DNA 1000 Kit on a 2100 Bioanalyzer (Agilent Technologies). The 201 sample libraries prepared were sequenced on a single run in an Illumina HiSeq3000 with single-end and 101bp configurations. The overall quality of the sequencing of GBS libraries was evaluated with the FastQC program (Andrews, 2010). Quality control and *demultiplex* were performed with the *process_radtags* module of the Stacks 1.42 program (Catchen et al., 2011), where low-quality reads were removed.

### 2.5 SNP calling

The SNP calling was performed using three different strategies because no reference genome is available for macauba. First, (i) using the de novo pipeline (Stacks v.1.42) (Catchen et al., 2013) based on the alignment of the reads obtained in the genotyping, (ii) using the genome of *Elaeis guineensis* var *tenera* (oil palm) as reference and (iii) using the transcriptome of *Acrocomia aculeata* as reference (Bazzo et al., 2018).

#### SNP calling using de novo pipeline

The identification of SNP was carried out using the de novo pipeline in the software Stacks *v*. 1.42 (Catchen et al., 2011). After the quality control, samples were demultiplexed using process-rad-tags in Stacks. Illumina reads were trimmed to 90 bp; then the *ustacks* module was used to identify groups of putatively homologous reads (putative loci) for each sample separately following the parameters: minimum sequencing depth (m = 3) and maximum mismatches (M = 2). Subsequently, a locus catalog was built using the *cstacks* module, allowing a maximum of two differences between stacks (Boutet et al., 2016) from different individuals. Loci with lower values of probability (lnl_lim -10) were eliminated by the *rxstacks* correction module. The SNPs were filtered using the *populations* module, retaining all SNP per sequence.

#### SNP calling using reference sequences

The *bwa-mem* algorithm of the *bwa* 0.7.17 program (Li, 2013) was used to align the sequences of each sample to the genome of *Elaeis guineensis* var *tenera* (oil palm) EG5 (NCBI GCA_000442705.1) and to the transcriptome sequencing of *Acrocomia aculeata* (Bazzo et al., 2018). Alignment files were processed with SAMtools (Li et al., 2009) and Picard programs (http://broadinstitute.github.io/picard). SNP identification was performed using the program *freebayes* 1.3.4 (Garrison & Marth, 2012) with the configuration --standard_filters. VCFtools 0.1.17 (Danecek et al., 2011) and bcftools 0.1.12 (Li et al., 2009) programs were used to filter SNP markers, retaining all SNP per sequence.

### 2.6 Quality control, imputation, and organization of genotypic data

To filter SNPs in the three SNP calling strategies mentioned above, we used the following criteria: maximum number of alleles = 2, minor allele frequency ≥ 0.01, sequencing depth ≥ 3X, mapping quality ≥ 20, maximum percentage of 30% of missing data per locus and of 45% of missing data per individuals. After filtering, we identified a total of 27410 SNPs in 153 individuals for the de novo genotypic dataset, 10444 SNPs in 158 individuals using the oil palm genotypic dataset, and 4329 SNPs in 167 individuals using the transcriptome genotypic dataset. Missing data were input using the Beagle 5.3 software (Browning et al., 2018, 2021).

Chromosome information was added to the three genotypic datasets to perform GWAS analyses. For de novo and transcriptome genotypic datasets, which do not have a reference genome, we considered that all the SNP markers belonged to the same chromosome. For the oil palm genotypic dataset, chromosome information was obtained from the NCBI GCA_000442705.1. The oil palm reference genome holds information from 16 chromosomes and from the complete chloroplast genome (reference sequence named NC_017602.1). The latter was excluded from our analysis. We also obtained sequences that have not yet been allocated in the oil palm genome, named “SNW”.

### 2.6 Phenotypic data analysis

Statistical analysis of the phenotypic data was performed using a mixed linear model in the lme4 package (Bates et al., 2015) in the R software, version 4.0.5 (R Core Team, 2021). The year information was used as a repetition. Statistical analysis of fruit characteristics was performed considering the mean of the three replications. The statistical model used to obtain the adjusted means:

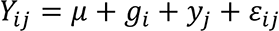

where *Y*_*ij*_ is the phenotype of the *ith* tree in the *jth* year, *μ* is the general mean, *g*_*i*_ is the random effect of the germplasm source *i*, *Y*_*j*_ is the fixed effect of years *j,* and ε_*ij*_ is the residual.

The significance of the random effect was estimated by the Likelihood Ratio Test at 5% probability, and the variance components were used to estimate the narrow sense heritability (H²):

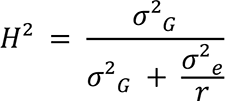

where σ^2^_*G*_ is the genotypic variance, σ^2^_*e*_ is the residual variance, and *r* is the repetition.

Pearson’s phenotypic correlation coefficients were calculated in all studied traits, while the genotypic correlation was calculated with multivariate linear mixed model by Likelihood methods (REML) using the sommer R package, version 4.1.5. (Covarrubias-Pazaran, 2016).

### 2.7 Genetic diversity and population structure

Genetic diversity and population structure were analyzed individually for each genotypic dataset in two different R packages: hierfstat (Goudet et al., 2005) and adegenet (Jombart, 2008). In this analysis, the four macauba subpopulations were considered. Genetic diversity was investigated according to the value of the total number of alleles, the observed and expected heterozygosity, and the inbreeding coefficient, while population structure was inferred by principal component analysis (PCA).

### 2.8 Single-trait and Multi-trait GWAS

To conduct single-trait GWAS, five different statistical models were used and compared. The five models fitted using the GAPIT package (version 3) (Wang et al., 2021) were: (i) general linear model (GLM) (Price et al., 2006), (ii) MLM (Yu et al., 2006), (iii) multiple loci MLM (MLMM) (Segura et al., 2012), (iv) fixed and random model circulating probability unification (FarmCPU) (Liu et al., 2016), and (v) bayesian-information and linkage-disequilibrium iteratively nested keyway (BLINK) (Huang et al., 2019). To account for population structure, we utilize the “Model.selection = TRUE” and “PCA.total=5”, which selects the best number of principal components from 0 to 5 in the GLM and MLM models. The relationship matrix was also calculated by GAPIT utilizing its default parameters.

The multi-trait analysis was fitted using a multivariate stepwise method (MSTEP) implemented in the software TASSEL, being conducted in all pairwise trait combinations (Fernandes et al., 2022). Prior to running MSTEP, each trait was normal quantile transformed using the orderNorm function from the R package bestNormalize (Peterson, 2020). Next, multivariate outliers were removed based based on the aq.plot function from the R package mvoutlier (Filzmoser and Gschwandtner, 2021). In all cases, MSTEP was fitted with and without the first five principal components as covariates obtained from the respective marker data set. SNPs that were significant in both instances were selected as high-confidence multivariate associations.

### 2.9 Identification of candidate genes

Tag sequences containing the significant SNPs from the de novo genotypic dataset were manually recovered from the catalog generated by the *cstacks* module in the Stacks program. The sequences of the significant SNPs from *Acrocomia aculeata* transcripts were obtained by retrieving fasta sequences from the transcriptome assembly with the perl fasta_FetchSeqs.pl script (https://github.com/4ureliek/Fasta). The occurrence of significant SNPs in predicted gene regions for *Elaeis guineenses* was verified with the *intersect* function from BEDTools v.2.30 program (Quinlan & Hall 2010), while the respective predicted protein sequences were recovered with the perl fasta_FetchSeqs.pl script. Blast2GO software (Conesa et al., 2005) was used to search for similarities between the sequences of significant SNPs and candidate genes. The *blastx* function was used for the de novo genotypic dataset, and the *blastp* for the transcriptome and oil palm genotypic dataset. Venn diagrams were obtained from the site InteractiVenn (Heberle et al., 2015).

## 3. RESULTS

### 3.1 Phenotypic analysis

Mean phenotypic values and their range, estimates of genetic, phenotypic, and residual variances, coefficient of variation (CV), heritability (ℎ²), and likelihood ratio test (LRT) are shown in Table 1. The likelihood ratio tests detected significant differences among genotypes for all traits (*p* < 0.001) (**Table 1**). Mean values within each trait studied showed a large interval. FM had the higher genetic, phenotypic, and residual variance values (176.20g, 248.20g, and 72.00g, respectively), while LL had the lower values (0.05cm, 0.07cm, and 0.02cm, respectively). The percentage of OC ranged from 17.47% to 60.80%. The value of the CV ranged from 22.38% in LN to 5.75% in LL. H, LL, OC, FM, PFM, EFM, HDM, EDM, and KDM traits had heritability values higher than 80%.

**Table 1.**
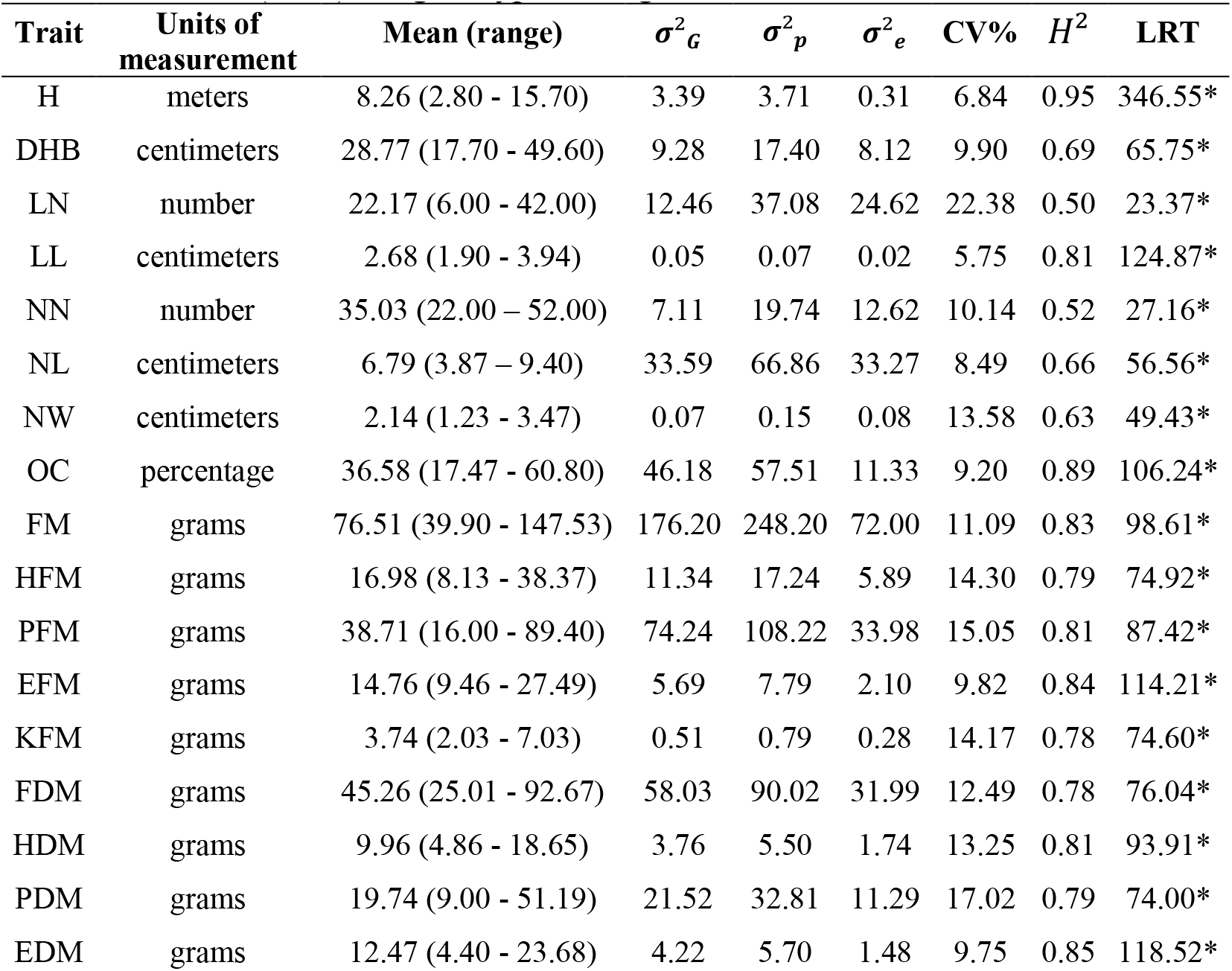
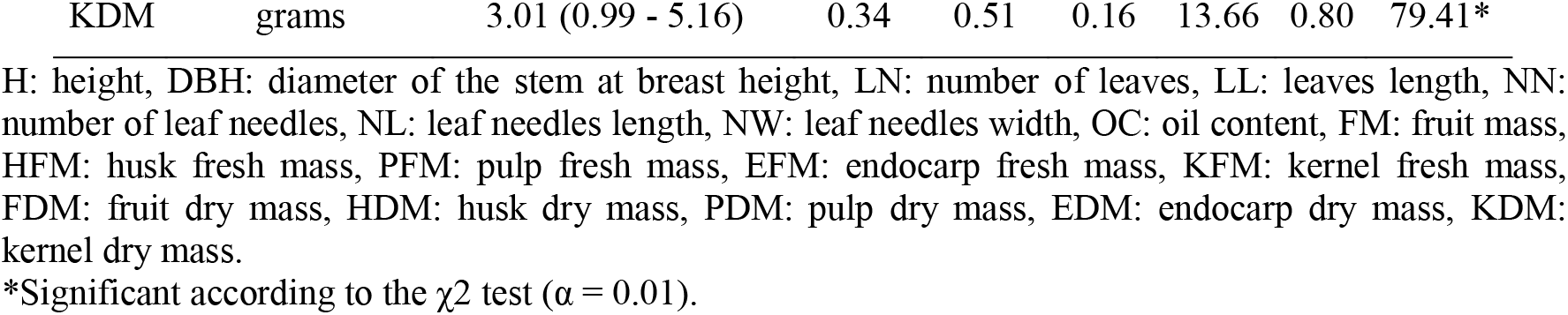
Means (range), estimates of genetic, phenotypic, and residual variances (σ^2^_*G*_, σ^2^_*p*_, and σ^2^_*e*_, respectively), coefficient of variation (CV%), heritability (*h*^2^) and likelihood ratio test (LRT) for genotypes in eighteen evaluated traits.

Pairwise genetic correlation between the eighteen traits studied are shown in **Figure 3**. Genetic correlation values ranged from -1 to 1, indicating a correlation between fruit traits. The traits FM and FDM (0.94), HFM and HDM (0.90), PFM and PDM (0.91), EFM and EDM (1.00), KFM and KDM (0.99) had positive genetic correlation because they are the same traits measured in different ways. Nevertheless, the difference between them is the presence of humidity. Traits that presented the highest correlations were FM and HFM (0.83), FM and PFM (0.95), FDM and PFM (0.88), FM and PDM (0.86), FDM and HDM (0.81), FDM and PDM (0.92). For OC and NN, null and negative values were observed for all the pairwise traits tested. For example, NN and LL showed -0.39, while for NL and NN, the correlation was -0.47. OC had a correlation varying from 0.00 to -0.42 between the pairwise evaluated traits (**Figure 3**).

**Figure 3.**
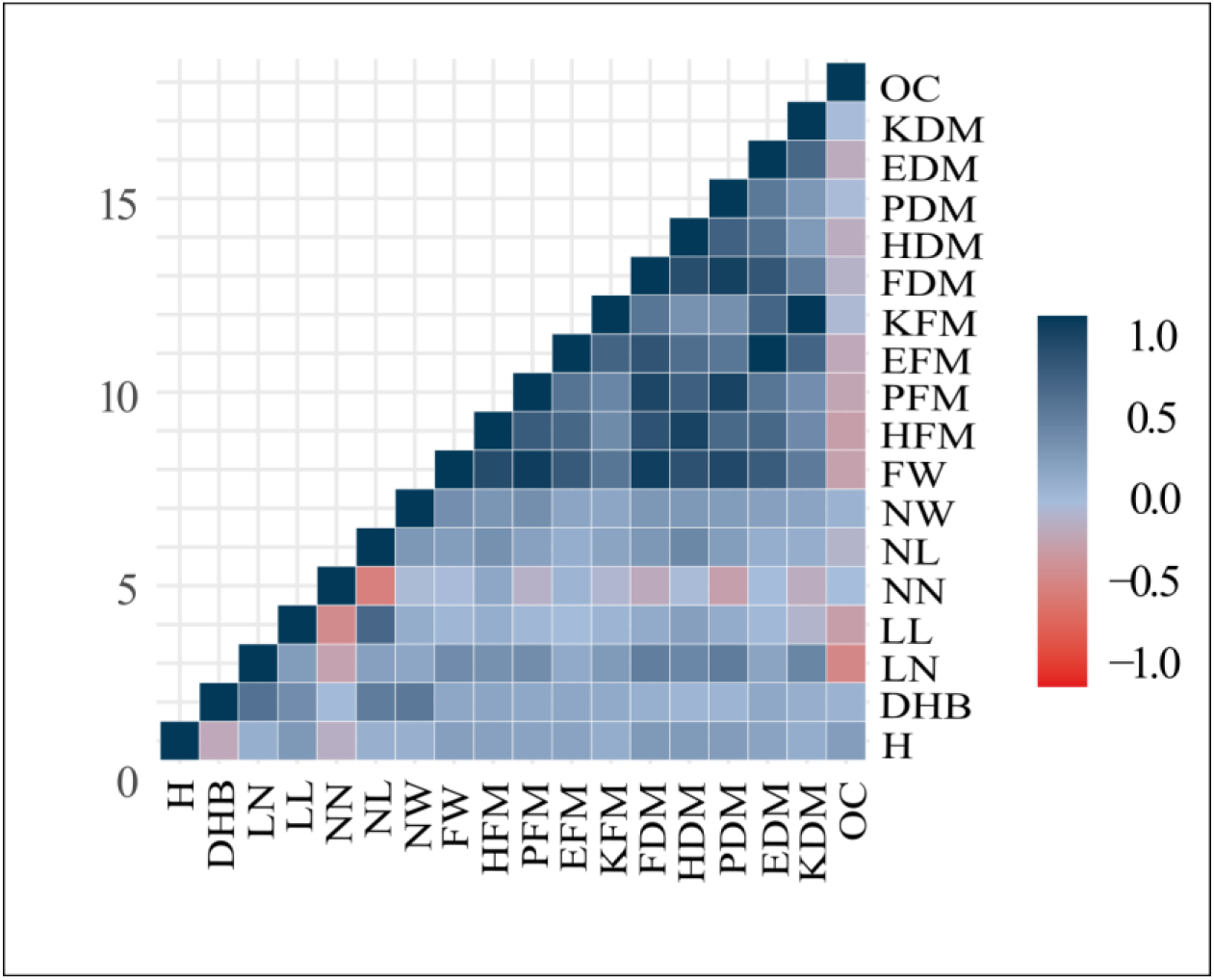
Genetic correlation among macauba traits. H: height, DBH: diameter of the stem at breast height, LN: number of leaves, LL: leaves length, NN: number of leaf needles, NL: leaf needles length, NW: leaf needles width, OC: oil content, FM: fruit mass, HFM: husk fresh mass, PFM: pulp fresh mass, EFM: endocarp fresh mass, KFM: kernel fresh mass, FDM: fruit dry mass, HDM: husk dry mass, PDM: pulp dry mass, EDM: endocarp dry mass, KDM: kernel dry mass.

### 3.2 Genetic diversity and population structure

Genetic parameters from the macauba population studied were different for each SNP calling strategy (**Table 2**). The three genotypic datasets used in this study showed high values of heterozygosity and low values of inbreeding coefficient.

**Table 2.** Population genetic parameters in macauba individuals: observed heterozygosity (*H*_*o*_), expected heterozygosity (*H*_*s*_), overall gene diversity (*H*_*t*_), Wright’s inbreeding coefficient (*Fis*)

| Genotypic dataset | Number of individuals | Number of SNP | $H_o$ | $H_s$ | $H_t$ | $F_{is}$ |
| --- | --- | --- | --- | --- | --- | --- |
| De novo | 153 | 27410 | 0.15 | 0.17 | 0.18 | 0.15 |
| Oil palm | 158 | 10444 | 0.41 | 0.30 | 0.31 | -0.35 |
| Transcriptome | 167 | 4329 | 0.53 | 0.34 | 0.34 | -0.53 |

Principal component analysis (PCA) in the three genotypic datasets revealed that individuals from subgroup 1 are slightly separated from individuals from subgroups 2, 3, and 4 (**Figure 4**). The four subgroups in the de novo genotypic dataset are evenly distributed in the graph, with some individuals of subgroup 1 separated from the other subgroups. The same situation was observed in the transcriptome genotypic dataset for the individuals from subgroup 1 and subgroups 2, 3, and 4. On the contrary, in the oil palm genotypic dataset, individuals from subgroups 2, 3, and 4 are tightly clustered, while the individuals from subgroup 1 are distant from the individuals of the other subgroups, with little overlap between them.

**Figure 4.**
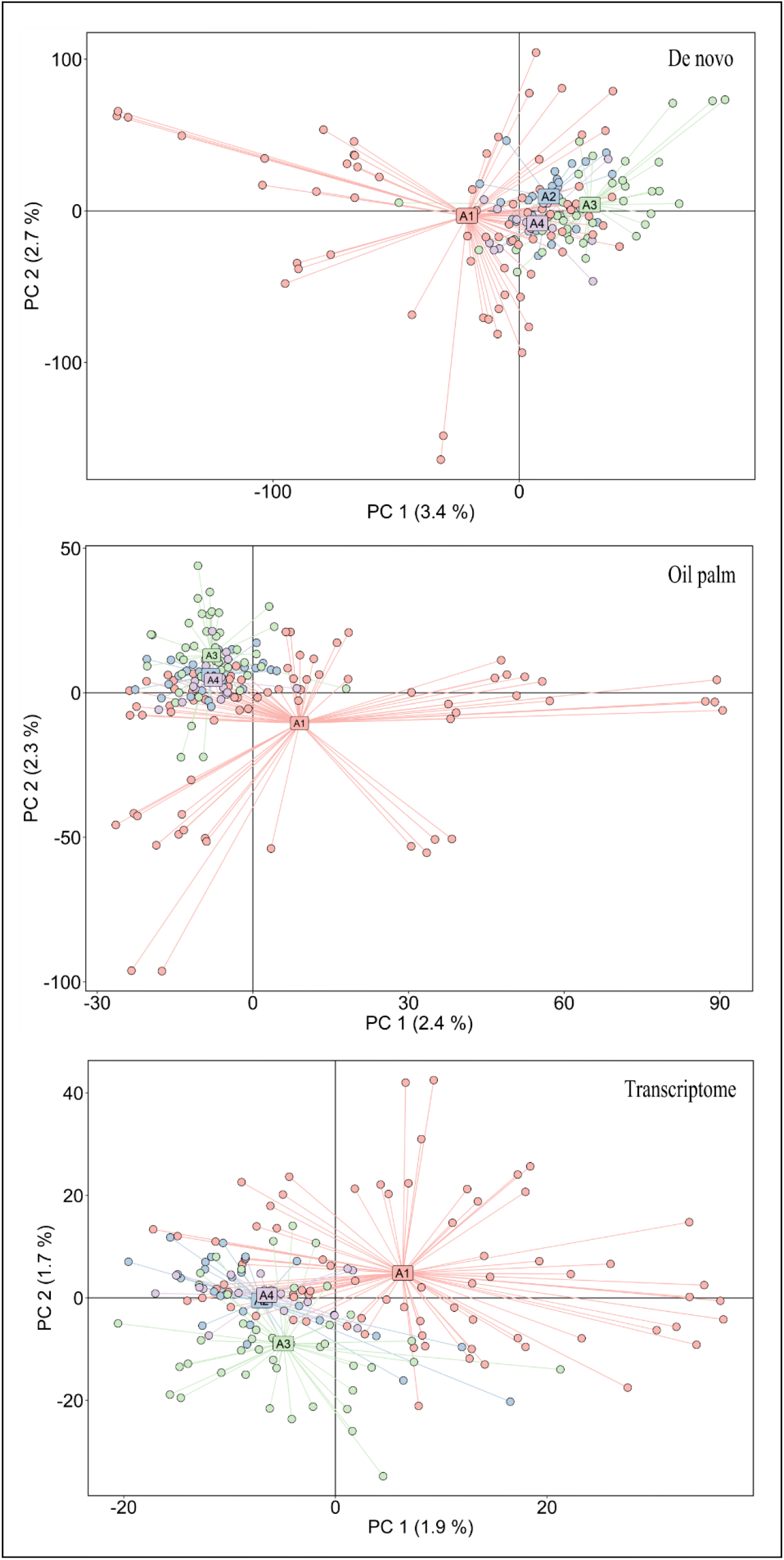
Principal component in the genotypic datasets de novo, oil palm and transcriptome. PCA shows the first two principal components in parentheses in the axis titles. Colors dots represent the four macauba subgroups selected in the four farm areas in this study. A1 is subpopulation 1, A2 is subpopulation 2, A3 is subpopulation 3, A4 is subpopulation 4.

### 3.3 Single-trait GWAS

The three genotypic datasets used in this study showed significant SNPs in the single-trait GWAS for the GLM, MLMM, FarmCPU, and BLINK models (**Table 3**). The MLM model did not detect significant SNP in any trait or genotypic datasets. The oil palm genotypic dataset showed a total of 19 significant SNPs in the NW, OC, FM, PFM, and PDM traits in the models GLM, MLMM, FarmCPU, and BLINK (**Supplementary Table 1**). From them, 10 SNPs occurred in genic regions, with the effects ranging from -12.67 to 55.96 (**Table 4**). The de novo genotypic dataset showed a total of 92 significant SNPs in the DBH, NL, NW, LL, OC, FM, HFM, PFM, EFM, KFM, FDM, HDM, PDM, and EDM traits in the models GLM, MLMM, FarmCPU and BLINK (**Supplementary Table 1**), with two occurring in genic regions, effect value of -0.90 and -10.43 (**Table 5**). To the de novo genotypic dataset, many of the significant SNPs were associated more than once in the same statistical model for different traits (**Supplementary Table 2**). The transcriptome genotypic dataset showed one significant SNP in the FDM trait in the MLMM model (**Supplementary Table 1**), and this SNP had a gene annotation in the *Elaeis guineensis* with an effect of 55.96 (**Table 6**).

**Table 3.** Single-trait and multi-trait GWAS in the genotypic dataset used in this work: de novo, oil palm, and transcriptome. The total number of significant SNPs shows the number of significant SNPs associated with the studied traits, considering the duplicated SNPs. Total candidate genes show the number of SNPs similar to genes in different <u>species.</u>

| GWAS | Genotypic dataset | Total of significant SNP | Total of candidate genes |
| --- | --- | --- | --- |
| Single-trait | De novo | 92 | 2 |
| Single-trait | Oil palm | 19 | 10 |
| Single-trait | Transcriptome | 1 | 1 |
| Multi-trait | De novo | 6 | 0 |
| Multi-trait | Oil palm | 11 | 3 |
| Multi-trait | Transcriptome | 2 | 1 |

**Table 4.**
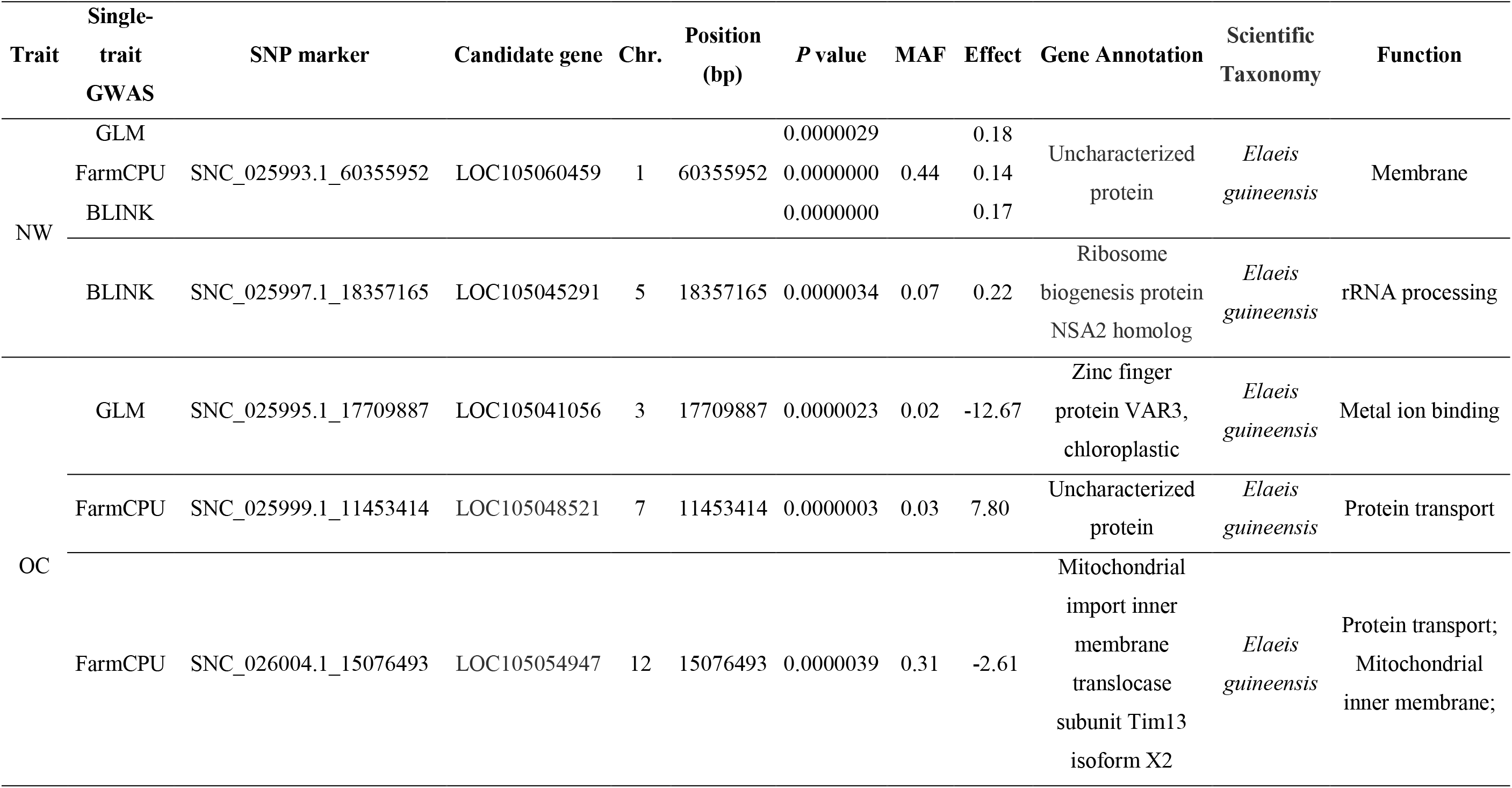
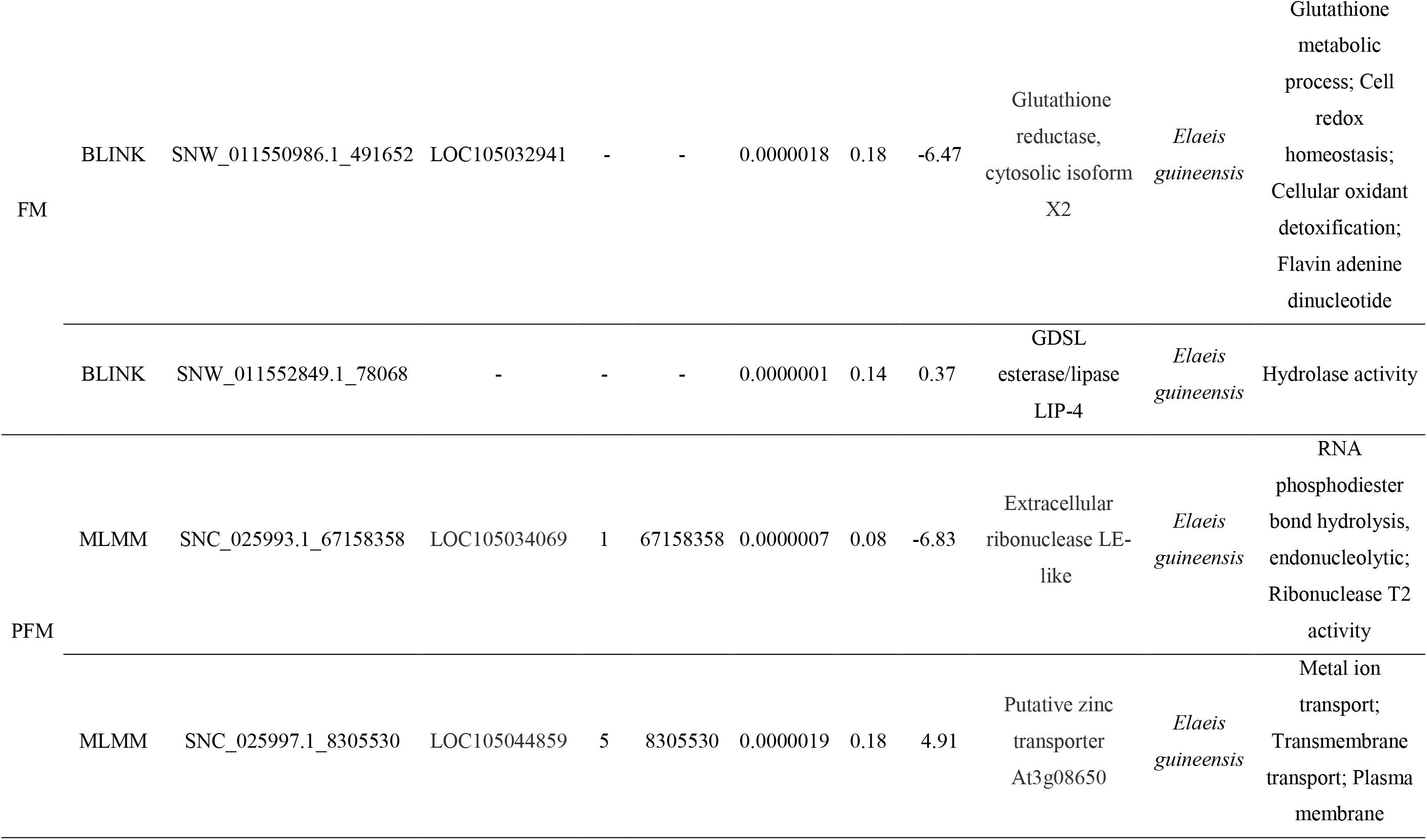
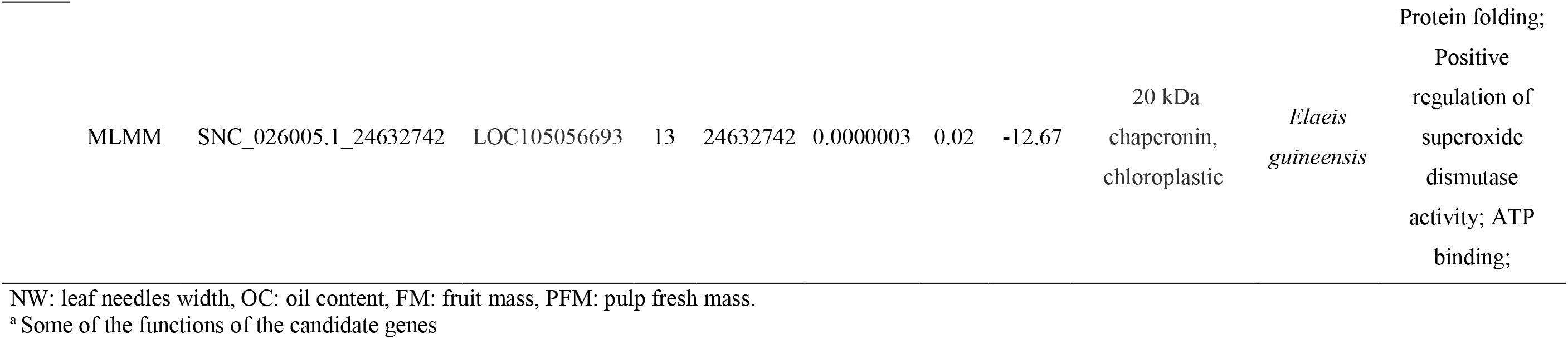
Candidate genes from the oil palm genotypic dataset significantly associated with NW, OC, FM, and PFM traits by different single-trait GWAS models.

| Trait | Single-trait GWAS | SNP marker | Candidate gene | Chr. | Position (bp) | P value | MAF | Effect | Gene Annotation | Scientific Taxonomy | Function |
| --- | --- | --- | --- | --- | --- | --- | --- | --- | --- | --- | --- |
| NW | GLM |  |  |  |  | 0.0000029 |  | 0.18 |  |  |  |
|  | FarmCPU | SNC_025993.1_60355952 | LOC105060459 | 1 | 60355952 | 0.0000000 | 0.44 | 0.14 | Uncharacterized protein | *Elaeis guineensis* | Membrane |
|  | BLINK |  |  |  |  | 0.0000000 |  | 0.17 |  |  |  |
|  | BLINK | SNC_025997.1_18357165 | LOC105045291 | 5 | 18357165 | 0.0000034 | 0.07 | 0.22 | Ribosome biogenesis protein NSA2 homolog | *Elaeis guineensis* | rRNA processing |
| OC | GLM | SNC_025995.1_17709887 | LOC105041056 | 3 | 17709887 | 0.0000023 | 0.02 | -12.67 | Zinc finger protein VAR3, chloroplastic | *Elaeis guineensis* | Metal ion binding |
|  | FarmCPU | SNC_025999.1_11453414 | LOC105048521 | 7 | 11453414 | 0.0000003 | 0.03 | 7.80 | Uncharacterized protein | *Elaeis guineensis* | Protein transport |
|  | FarmCPU | SNC_026004.1_15076493 | LOC105054947 | 12 | 15076493 | 0.0000039 | 0.31 | -2.61 | Mitochondrial import inner membrane translocase subunit Tim13 isoform X2 | *Elaeis guineensis* | Protein transport; Mitochondrial inner membrane; |
| FM | BLINK | SNW_011550986.1_491652 | LOC105032941 | - | - | 0.0000018 | 0.18 | -6.47 | Glutathione reductase, cytosolic isoform X2 | *Elaeis guineensis* | Glutathione metabolic process; Cell redox homeostasis; Cellular oxidant detoxification; Flavin adenine dinucleotide |
|  | BLINK | SNW_011552849.1_78068 | - | - | - | 0.0000001 | 0.14 | 0.37 | GDSL esterase/lipase LIP-4 | *Elaeis guineensis* | Hydrolase activity |
| PFM | MLMM | SNC_025993.1_67158358 | LOC105034069 | 1 | 67158358 | 0.0000007 | 0.08 | -6.83 | Extracellular ribonuclease LE-like | *Elaeis guineensis* | RNA phosphodiester bond hydrolysis, endonucleolytic; Ribonuclease T2 activity |
|  | MLMM | SNC_025997.1_8305530 | LOC105044859 | 5 | 8305530 | 0.0000019 | 0.18 | 4.91 | Putative zinc transporter At3g08650 | *Elaeis guineensis* | Metal ion transport; Transmembrane transport; Plasma membrane |

|  |  |  |  |  |  |  |  |  |  |  |
| --- | --- | --- | --- | --- | --- | --- | --- | --- | --- | --- |
| MLMM | SNC_026005.1_24632742 | LOC105056693 | 13 | 24632742 | 0.0000003 | 0.02 | -12.67 | 20 kDa<br>chaperonin,<br>chloroplastic | *Elaeis*<br>*guineensis* | Protein folding;<br>Positive<br>regulation of<br>superoxide<br>dismutase<br>activity; ATP<br>binding; |

NW: leaf needles width, OC: oil content, FM: fruit mass, PFM: pulp fresh mass.
<sup>a</sup> Some of the functions of the candidate genes

**Table 5.**
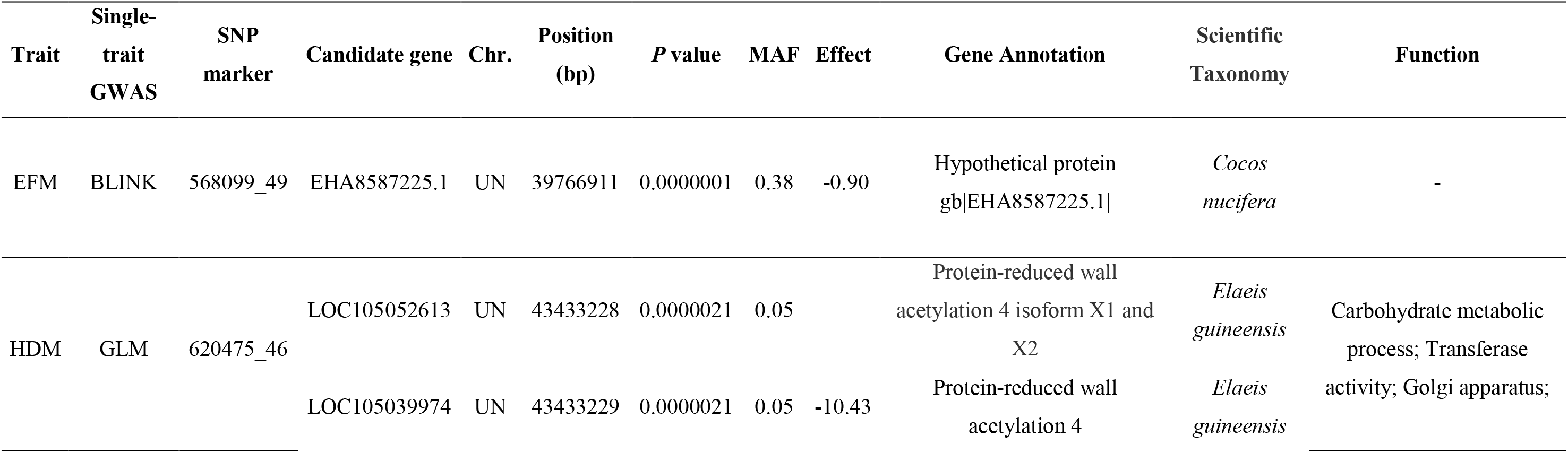
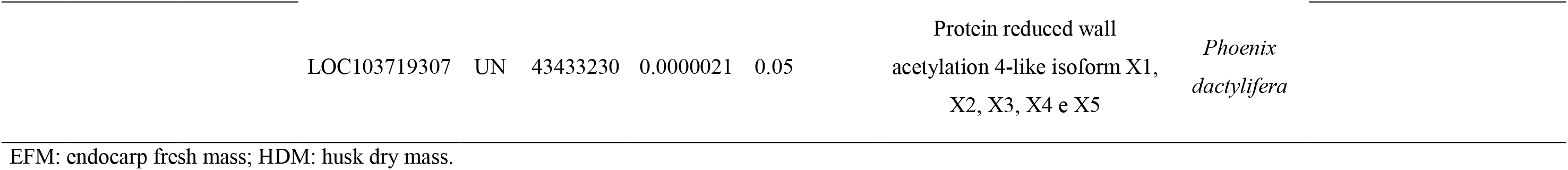
Candidate genes from the de novo genotypic dataset significantly associated with EFM and HDM traits by different single-trait GWAS models

**Table 6.**
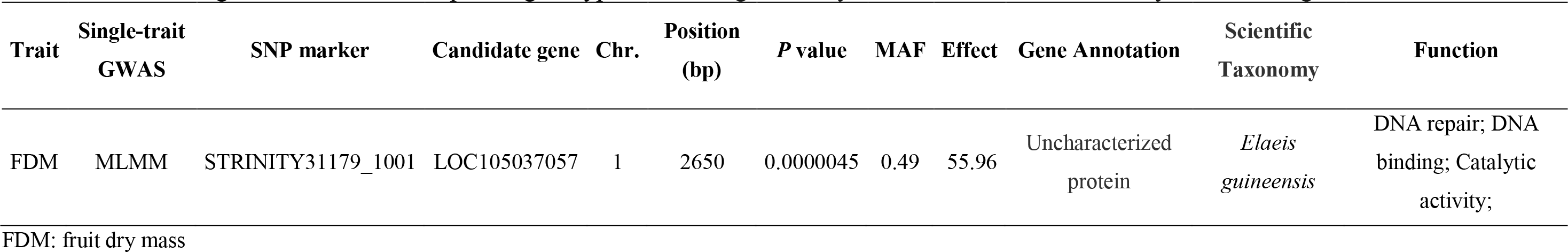
Candidate gene from the transcriptome genotypic dataset significantly associated with FDM trait by MLMM single-trait GWAS model

For the candidate genes, the marker SNC_025993.1_60355952, in the oil palm genotypic dataset, was detected in three single-trait models (GLM, FarmCPU, and BLINK), for the same loci (LOC105060459), which has a membrane function (**Table 4**). Additionally, the SNP 620475_46 in the de novo genotypic dataset was associated with 20 candidate genes in different species, all of them having 100% similarity (data not shown). **Table 5** shows three of these 20 candidate genes, being similar to the *Elaeis guineenses* and *Phoenix dactylifera*, which are phylogenetically closer to macauba. In all the genotypic datasets used, candidate genes showed similarity with the species *Elaeis guineensis* (100%), *Phoenix dactylifera* (100%), and *Cocos nucifera* (91.3%) and presented function as protein transport, DNA repair, metal ion transport, maturation of ribosome subunit and transmembrane transportation.

### 3.5 Multi-trait GWAS

In the multi-trait GWAS, the MSTEP model detected a total of 19 markers associated with traits in all the genotypic datasets studied (**Table 3, Supplementary Table 3**). To the oil palm genotypic dataset, SNC_025993.1_60355952, SNC_025995.1_17709887, and SNC_026000.1_5081018 markers had similarities with *Elaeis guineensis* genes in eight trait combinations. The SNC_025993.1_60355952 showed an association with the trait combinations H-NW and DBH-NW, having membrane function. Furthermore, this marker was also detected in three single-trait GWAS models (**Table 4**, **Figure 5**). The second marker, SNC_025995.1_17709887, had an association with the traits LL-OC and LN-OC, having also been detected in the single-trait GWAS for the OC trait (**Table 4**, **Figure 5**). In addition, the SNP SNC_026000.1_5081018 had an association with the traits LN-HFM, NN-HFM, NN- FM, and NN-HDM. For the transcriptome genotypic dataset, the marker STRINITY9279_4262 was associated with the traits FDM-OC and FDM-HDM and showed similarity with an *Elaeis guineenses* uncharacterized protein (**Table 7**). To visualize the single and multi-trait models simultaneously, we marked in red each of the significant SNPs detected in the multi-trait model in the manhattan plots obtained for each of the traits in the single-trait analysis. Some of these SNPs were also significant in the single trait analysis, while others were not (**Figure 6**). From single-trait and multi-trait GWAS, five candidate genes identified on chromosomes 1 and 7 refer to gene annotation, which proteins have not yet been characterized (**Tables 4, 6, and 7**).

**Figure 5.**
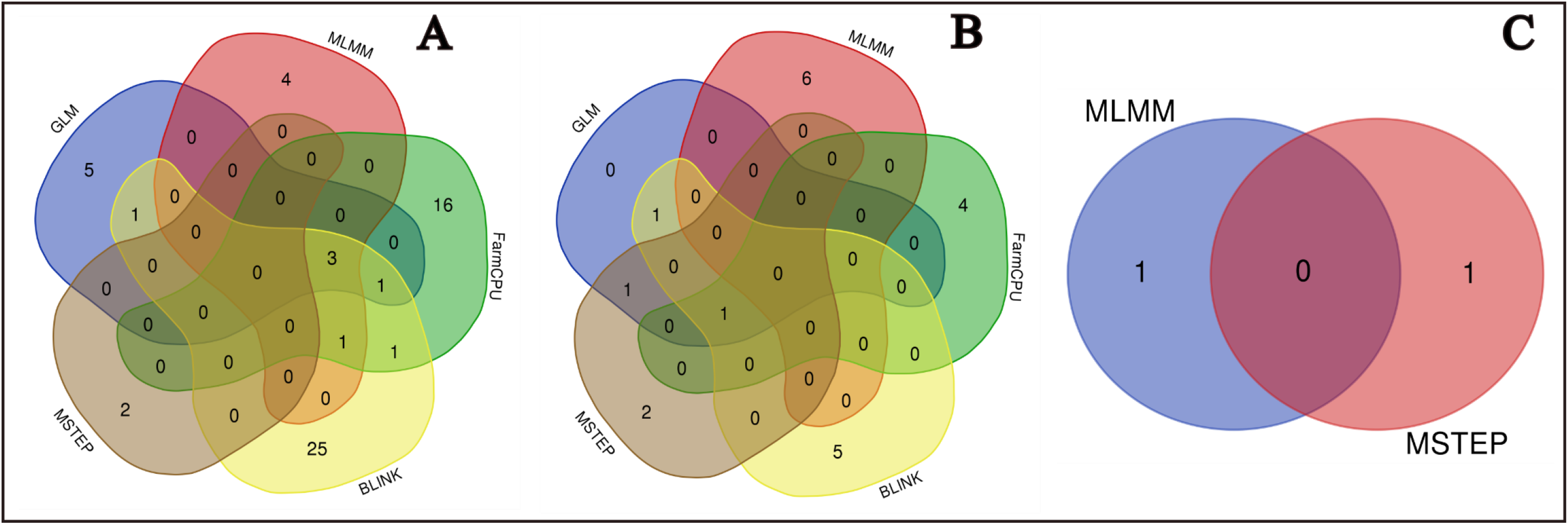
Significant SNPs detected by different single-trait (GLM, MLMM, FarmCPU, BLINK) and multi-trait (MSTEP) GWAS models in each genotypic dataset. Venn Diagram shows the unique and shared significant SNP in the de novo (A), oil palm (B), and transcriptome (C) genotypic datasets. The figure represents the total number of SNP markers in each genotypic dataset without the duplicated SNPs in the de novo genotypic dataset.

**Figure 6.**
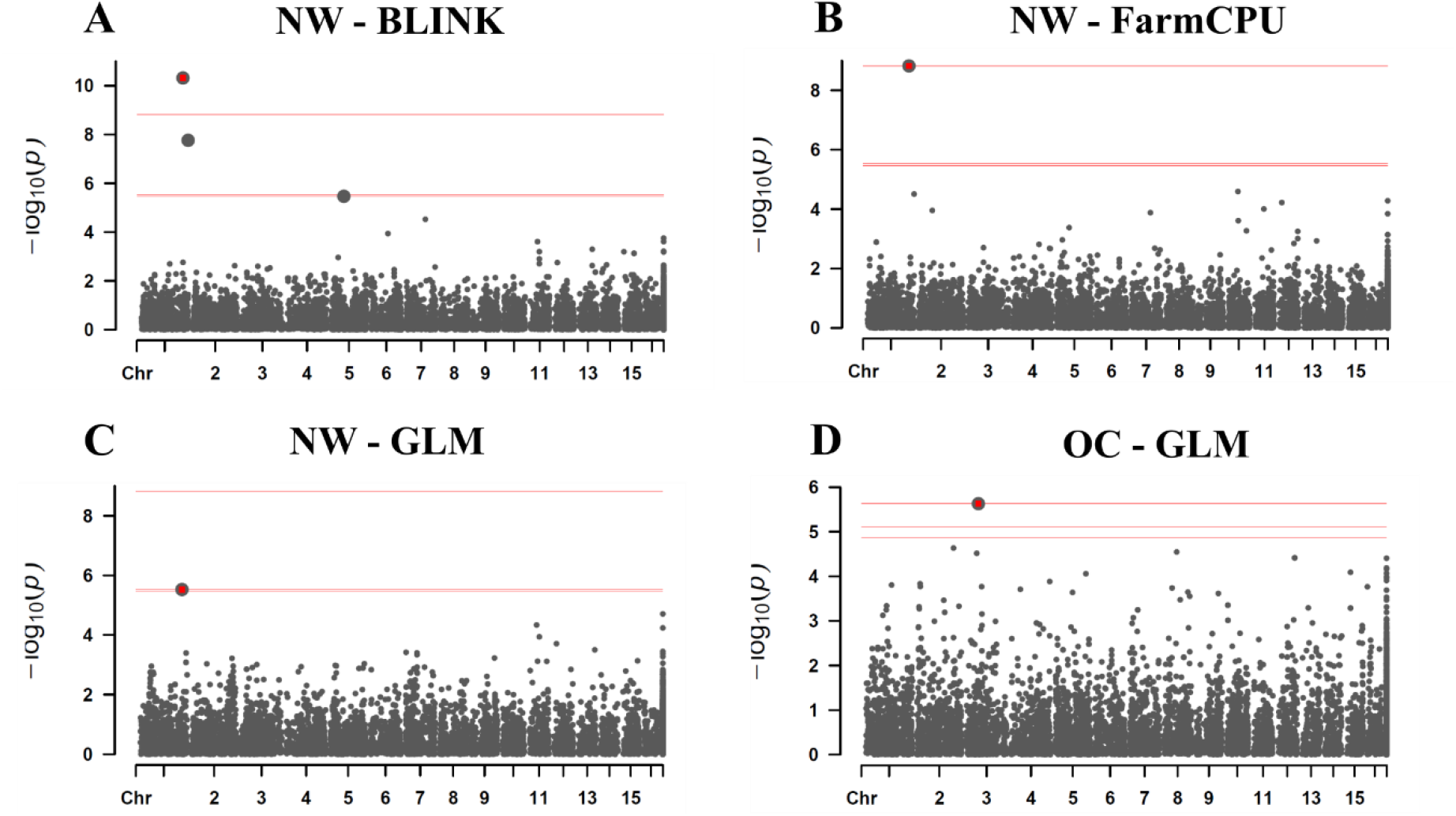
Manhattan plot showing SNPs detected in both single-trait and multi-trait GWAS for oil palm genotypic dataset for NW and OC traits. The red dot represents the significant SNP detected in the multi-trait GWAS, which was also detected in the single-trait GWAS. The red square in graphs A, B, and C are taggint the candidate gene SNC_025993.1, while in graph D, it is tagging the candidate gene SNC_025995.1.

**Table 7.**
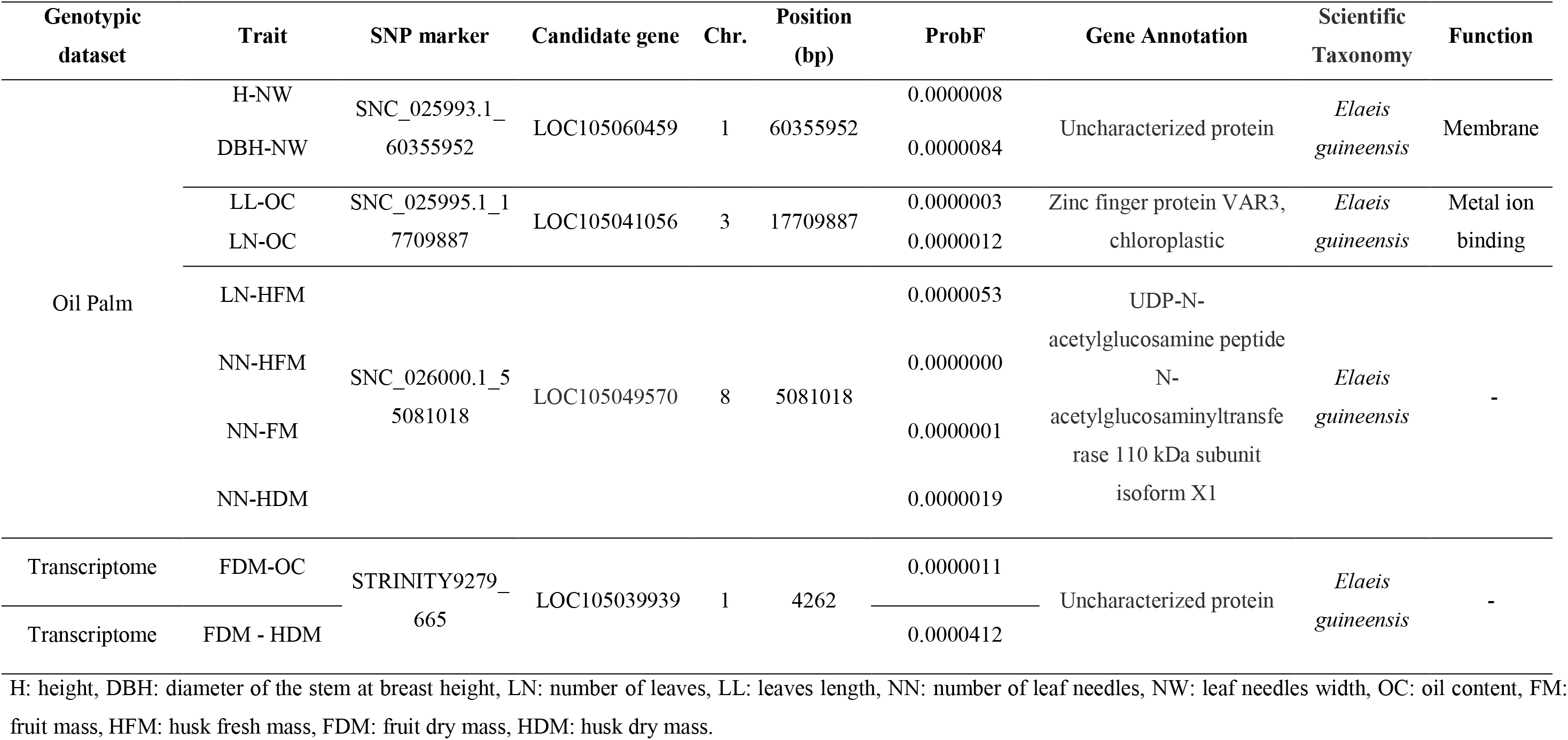
Candidate genes from the oil palm and transcriptome genotypic dataset significantly associated with different trait combinations by the Multi-trait GWAS MSTEP model

## 4. DISCUSSION

*Acrocomia* is a genus from a neotropical palm distributed in almost all tropical and subtropical America, there are nine recognized species in the genus: *A. aculeata*, *A. totai*, *A. hassleri*, *A. glaucescens*., *A. emensis*; *A. intumescens*; *A. media, A. crispa,* and *A. corumbaensis* (Lima et al., 2018; Lorenzi et al., 2010; Diaz et al., 2021). From them, macauba (*Acrocomia aculeata*) stood out in recent years due to its high percentage of oil content in the pulp and kernels of its fruit, with productivity as the oil palm (*Elaeis guineenses*). To our knowledge, this is the first GWAS study in *Acrocomia aculeata*. Understanding macauba genetic architecture is an initial step towards its domestication and commercial planting since markers associated with traits of economic interest can help the selection of promising genotypes in breeding programs.

### 4.1 Importance of the analyzed traits to *Acrocomia aculeata* domestication

The data used in this work were collected from a natural population located inside a private rural property. Obtaining and analyzing macauba phenotypic data allows us to understand the relation between traits and guide future directions in the species’ breeding. In this study, the significance of genetic effects revealed levels of genetic variability within the macauba population evaluated (**Table 1**). Furthermore, we observed a wide range in the phenotypes measured. Given that macauba is a non-domesticated palm, this occurred as a consequence of sampling a macauba native population. The study did not have an experimental design, which means that the age of the macauba genotypes was unknown. For example, the height (H) and diameter of the stem at breast height (DHB) are traits that could not be obtained in plants of the same age. Therefore, it is important to bear in mind that a genotype with low height, compared with another, may be due to its age and not due to its genetic merit. This is one of the great challenges in studying perennial and native populations. In macauba, the first flowering occurs in the fifth or sixth year of the palm tree (Teixeira, 2005). Furthermore, fruit development is slow and can last up to 62 weeks after anthesis, equivalent to 14.3 months (Mazzottini-dos-Santos et al., 2015; Montoya et al., 2016).

Even without an experimental design, the value of the CV ranged from 22.38% to 5.75%, highlighting good accuracy in obtaining phenotypic data. The 60.80% mean of OC in the individuals studied, indicate that superior genotypes for this trait can be selected in this population. Additionally, H, LL, OC, FM, PFM, EFM, HDM, EDM, and KDM showed heritability values ranging from 50 to 80%, evidencing that the performance of breeding cycles in this selected population tends to produce genetic gains in these traits. Other studies have reported high mean heritability values for various morphological traits in macauba (Berton et al., 2013; Domiciano et al., 2015; Coser et al., 2016; Rosado et al., 2019).

Correlation between traits is of great interest in plant breeding since they enable an indirect selection of multi-traits (Fernandes et al., 2018; Costa et al., 2018; Domiciano et al., 2015). In this case, the genotypic correlation is used in data analysis to guide breeding programs because it is inheritable (Cruz, Regazzi, and Carneiro, 2012). Furthermore, high values of genetic correlation may indicate the presence of linked genes or pleiotropy (Fernandes et al., 2021). In this study, we found high genetic correlation values between the fruit traits FM-HFM, PFM-PDM, FDM-PFM, and HDM-PDM (**Figure 3**). In the same way, Costa et al. (2018) found high positive genotypic correlations between macauba fruit traits, such as dry matter from endocarp and epicarp, dry matter from endocarp and kernel, and dry matter from pulp and oil yield per plant. Ciconini et al. (2013) and Concceição et al. (2015) found a positive correlation in pulp oil content with total fruit oil but a negative correlation of pulp oil content with the mesocarp’s proportion of the whole fruit. Negative and low correlation values were found in other macauba studies: i) between stem diameter at breast height with the height of the first spathe (Coser et al., 2016); ii) between oil content and oil yield per plant (Costa et al., 2018); and iii) between biometric fruit traits and oil content (Ciconini et al., 2013). Further information is needed on the selection criteria of the corresponding correlation between traits to assist the breeding of commercial varieties of macauba (Vargas-Carpintero et al. 2021).

The analysis of the genetic diversity of macauba is crucial to guide the selection of the most promising materials for use, maximize genetic gains, and more effectively contribute to the creation of commercial cultivars (Díaz et al., 2021). The three genotypic datasets used in this study showed high values of observed heterozygosity and low values of inbreeding, indicating genetic variability in the population accrediting the population for breeding purposes. An ideotype model of the improved macauba was proposed by Berton (2013): early flowering, low height, high production of fruits and oil content, fewer or absence of thorns, indehiscent fruits, and overlapping bunches. Therefore, the study of macauba traits is important to assist in the selection of potential plants that encompass the traits to be added to the ideotype demanded by the market.

### Absence of a reference genome in *Acrocomia aculeata*

Because *Acrocomia aculeata* does not have a reference genome, different strategies were proposed in this study to perform SNP calling, namely, using the *de novo* pipeline (Catchen et al., 2013), the reference genome of *Elaeis guineensis* var *tenera*, and the transcriptome of *Acrocomia aculeata* (Bazzo et al., 2018). *Elaeis guineensis* reference genome was used in this work due to the phylogenetic proximity to *Acrocomia aculeata* (Bazzo et al., 2018). Phylogenomic studies using chloroplast genome sequences produced a phylogenomic tree in which all nodes had a posterior probability of 1.0 (PP = 1.0). In this tree, *Acrocomia aculeata* had a proximity to *Elaeis guineensis* (Francisconi et al., 2023). The three genotypic datasets used in this study also showed different results in the genetic diversity and PCA analyses, as well as in the GWAS. The number of individuals and SNP markers used in the genetic diversity analyses were different in each genotypic dataset due to the SNP calling and the quality control step (**Table 2**).

Population genetic parameters in macauba individuals from the oil palm and transcriptome genotypic datasets presented higher heterozygosity values than the population from the de novo genotypic dataset (**Table 2**). Therefore, the inbreeding coefficient was higher in the population of the de novo genotypic dataset than in the other two. These values indicate that the heterozygosity and inbreeding coefficient values of the studied macauba populations depended on the strategy used in the SNP calling. Díaz et al. (2021) studied genetic diversity in the *Acrocomia* genus using genome-wide SNP and the de novo genotypic pipeline. They also reported low heterozygosity values (0.031). In our study, the population genetic parameters from the *de novo* genotypic dataset showed low heterozygosity values and higher inbreeding coefficient values when compared to the other genotypic datasets. These results occurred due to the de novo pipeline obtained from the software Stacks, where homologous reads were used to identify the SNPs (Catchen et al., 2011). Conversely, the oil palm and transcriptome dataset had higher heterozygosity values and lower inbreeding coefficient values. This occurred because the *Elaeis guineensis* reference genome used in our study presents higher genetic variability in its alleles than the genome of the de novo pipeline. Even though *Acrocomia* and *Elaeis* are close genera, it is expected that they hold different alleles due to their evolutionary history. The transcriptome dataset was generated by the alignment step with the transcriptome sequencing of *Acrocomia aculeata* (Bazzo et al., 2018). Bazzo and collaborators reported the identification and validation of 145 macauba EST-SSR markers from different tissues and individuals using next-generation sequencing. The study was performed using tissues of leaves, leaf sheaves, roots, bulbs, fruit (mesocarp and endosperm), and male and female flowers for RNA isolation and transcriptome sequencing. The reads originating from mRNA libraries were mapped against the *Elaeis guineensis* reference genome (Singh et al., 2013). Results indicated an 80.7% transferability rate of macauba in oil palm.

PCA revealed that individuals from subgroup 1 are slightly separated from subgroups 2, 3, and 4 (**Figure 3**). However, among the genotypic datasets, it is possible to see that subgroups 2, 3, and 4 presented variations in their groupings, showing that SNP calling strategies may lead to different interpretations of the population subgroups. Results showed here indicate that the SNP calling strategies are important to provide a direction in understanding how the macauba population is behaving in terms of diversity and genetic architecture.

### 4.2 Single-trait and multi-trait GWAS in macauba

Using single-trait and multi-trait GWAS models, we identified SNP markers located in gene regions related to different traits involved in fruit production and fruit pulp oil content of a macauba population. The results related to the genotype-to-phenotype association in these traits could be relevant for understanding the genetic architecture of this neotropical palm. Moreover, the genetic mapping and molecular characterization of these genes that contribute to the variation of complex traits have the potential to facilitate genome-assisted breeding for crop improvement (Stich & Melchinger, 2010). The elucidation of its genetic information could still speed up the implementation of breeding programs to select superior genotypes for traits involved with the oil production of macauba.

The total number of significant SNPs detected in the single-trait and multi-trait GWAS was greater than the number of candidate genes (**Table 3**). This result shows that false positives could have been detected by the statistical models used, and it highlights the importance of obtaining the reference genome for the macauba. The single-trait GWAS revealed significant genomic regions in the three datasets used in this study, associated with the following traits: DBH, LL, NL, NW, OC, FM, HFM, PFM, EFM, KFM, FDM, HDM, PDM and EDM. For the traits H-LN, no genomic regions were found in the search for similarity in the software Blast2GO (**Tables 4, 5, and 6**). This result may have occurred because the phenotypic data came from native palms, whose lifetime is unknown. Therefore, it was not possible to control these traits of the plants taking their age into account, as already discussed in the topic above.

We observed that different single-trait GWAS models detected the same loci (LOC105060459) for the same trait (**Table 4**), or the same SNP markers were detected for different trait combinations (**Supplementary Table 1**). Similar results were observed in the multi-trait models (**Table 7**). Moreover, for the multi-trait model, four candidate genes (SNC_025993.1, SNC_025995.1, SNC_026000.1, STRINITY9279) stood out in the phenotype-genotype association between different combinations of traits. Among these traits, NW, OC, NN, HFM, HDM, and FDM appeared more than once. Furthermore, the SNP markers SNC_025993.1 and SNC_025995.1, detected in the multi-trait model, were also detected in the single-trait models (**Figure 5**). These results demonstrate the certainty of the presence of these loci in the macauba genome, reducing the chance of these SNP markers being false positives. In this context, Fernandes et al. (2021) suggest that single-trait and multi-trait GWAS should both be used to infer whether or not causal mutations underlying peak GWAS associations are pleiotropic. However, the authors discuss the importance of noting that statistical analysis alone cannot distinguish between QTNs in LD and a single pleiotropic QTN, underlying the importance of validating significant SNPs detected by the GWAS.

### 4.3 Candidate genes

In general, candidate genes detected in this work were responsible for RNA maturation, metal ion binding and transport, protein transportation, DNA repair, carbohydrate metabolic process, and other cell regulation biological processes.

From single-trait GWAS, in the NW trait, the candidate gene LOC105045291, which has 260 amino acids polypeptides, is a protein with a gene annotation to ribosome biogenesis protein NSA2 homolog (**Table 4**). NSA2 (Nop seven-associated 2) is a nucleolar protein associated with the ribonucleoprotein complex, involved in cell proliferation and cell cycle regulation. It was identified in a high throughput screening of novel human genes, being evolutionarily conserved across different species (Zhang et al., 2010). For the oil content trait, three candidate genes were detected (**Table 4**). One of them, the loci LOC105041056, is the protein Zinc finger protein VAR3 with 758 amino acids. This protein was also detected in the multi-trait GWAS for the trait combination LL-OC and LN-OC (**Table 7**). Proteins containing the Zinc finger domain play an important role in many cellular functions, including transcriptional regulation, RNA binding, regulation of apoptosis, and protein-protein interactions (Ciftci-Yilmaza & Mittler, 2008). A study in Arabidopsis used a recessive Arabidopsis variegated 3 (var3) mutants to study the VAR3 gene and observed a difference in the metabolic profiling analysis. The pigment profiles are qualitatively similar in wild type and var3. However, var3 accumulates lower levels of chlorophylls and carotenoids. The authors indicated that var3 is part of a protein complex required for normal chloroplast and palisade cell development (Naested et al., 2004).

For the FM trait, the SNP marker SNW_011552849.1_78086 was blasted to the GDSL esterase/lipase LIP-4 gene, which has a hydrolase activity (**Table 4**). GDSL esterase/lipase protein has a variety of lipolytic enzymes that hydrolyze diverse lipidic substrates, including thioesters, aryl esters, and phospholipids (Ding et al., 2019). Cao et al. (2018) studied GDSL esterase/lipase in Rosaceae genomes and found that it participated in fruit development. In macauba, the fruiting is supra-annual, and the fruit growth curve follows a double sigmoidal trend with four stages, observed by Montoya et al. (2016): in the first stage occurs slow growth and negligible differentiation of the fruit inner parts, the second occurs the first growth spurt differentiation of the inner parts; in the third stage the fruit growth slows down, and all structures attained the differentiation; and for last, the second growth spurt and fruit maturation. So, considering this fact, we can suppose that the GDSL esterase/lipase LIP-4 gene is involved in the macauba fruit development. However, to confirm this hypothesis, future research should be implemented to validate this candidate gene in the species.

For the PFM trait, the 20 kDa chaperonin protein located in the chloroplast was detected by the MLMLL single-trait model (**Table 4**). In general, molecular chaperones functionally support protein translocation across membranes, promote complex assembly and disassembly, and participate in many other regulatory processes within the cell (Hartl, 1996 and 2011; Sharma et al., 2010). In *Elaeis guineensis*, a study using proteomics and chemometrics approach provided important information about protein regulation during fruit ripening and oil synthesis (Hassan et al., 2018). In this study, the 20 kDa chaperonin was identified as a folding protein, and the authors observed that it had a downregulated towards fruit ripening.

Candidate genes from the de novo genotypic dataset in the single-trait GWAS showed gene annotation for the Reduced wall acetylation 4 protein in the HDM trait (**Table 5**). The Reduced wall acetylation 4 protein is part of a family protein involved in the O-acetylation of cell wall polysaccharides in the Golgi apparatus. Specifically, the Reduced wall acetylation 4 is thought to be responsible for the translocation of acetyl-CoA across the Golgi membrane and appears to supply the acetyl-donor to both pectins and hemicelluloses (Lee et al., 2011; Manabe et al., 2011). The polysaccharides pectins, hemicelluloses, and celluloses are the main cell wall components in the epidermis, contributing to its protection against xenobiotics, ultraviolet light, and pathogens and providing a waterproof barrier (Nafisi et al., 2015). In this context, the Reduced wall acetylation 4 protein can be involved in the same cell functional route in the epidermis of the husk of macauba fruits.

From multi-trait GWAS, the same candidate gene LOC105049570 was detected for the combination of the traits: LN-HFM, NN-HFM, NN-FM, and NN-HDM (**Table 7**). The UDP-N-acetylglucosamine peptide N-acetylglucosaminyltransferase is the protein blasted with the SNC_026000.1_ 5081018 SNP marker. In cucumber, a study identified and mapped the CsSF4 gene, which encodes for the protein mentioned above (Zhang et al., 2023). The authors observed that the CsSF4 gene was highly expressed in the leaves and male flowers of wild-type cucumbers and that it is required for fruit elongation and development. Since the UDP-N-acetylglucosamine peptide N-acetylglucosaminyltransferase protein was detected for different trait combinations, future studies aimed at validating the candidate gene LOC105049570 are necessary. Through them, it will be possible to confirm the presence of pleiotropy or genes linked to these traits.

## CONCLUSIONS

In our study, genotypes were selected in their natural habitat with the aim of starting to understand the genetic architecture of macauba to open ways for new studies to be carried out. We provide new insights on genomic regions that mapped candidate genes involved in macauba oil production phenotypes. These potential candidate genes need to be confirmed for future targeted functional analyses, and multi-trait associations need to be scrutinized to investigate the presence of pleiotropic or linked genes. Associated markers to the traits of interest may be valuable resources for the development of marker-assisted selection in macauba for its domestication and pre-breeding. This is the first work using GWAS in macauba, and the significant SNPs detected here can be validated in a future study in order to allow marker-assisted selection to support the selection of traits of interest in native plants in other populations.

## Supporting information

Supplementary material_Couto et al.

## ACKNOWLEDGEMENTS

We would like to thank Luiz Henrique Berton, Gabriel Alves, Elivelton Alves, and Evandro Coelho from the Campinas Agronomic Institute and Marcela Barbosa, Laecio Sampaio, and Marcelo Almeida from the Luiz de Queiroz College of Agriculture, for their assistance in obtaining phenotypic data and fruit biometry. We would also like to thank Bárbara Regina Bazzo, Lucas Miguel de Carvalho, and Marcelo Falsarella Carazzolle, who provided the macauba transcripts for the SNP calling procedures. Finally, we would like to thank Saulo Fabrício das Silva Chaves for his valuable suggestions when reviewing the manuscript.

## Data availability

Data will be made available on request.

## AUTHOR CONTRIBUTIONS

The first draft of the manuscript was written by Evellyn Giselly de Oliveira Couto, and all authors read and approved the final manuscript prior to publication. The contribution of each author is presented below:

**Evellyn Giselly de Oliveira Couto**: Data collection, Genotypic data obtation, Methodology, Data mining analyses, Data curation, Conceptualization, Investigation, Writing, Review & Editing.

**Jonathan Morales-Marroquín**: Data collection, Genotypic data obtation, Methodology, Investigation, Review & Editing.

**Alessandro Alves-Pereira**: Bioinformatics analyses, Data mining analyses, Validation.

**Samuel B. Fernandes**: Conceptualization, Methodology, Data mining analyses, Validation, Review & Editing.

**Carlos Augusto Colombo**: Data collection, Resources, Investigation, Funding acquisition.

**Joaquim Adelino de Azevedo Filho**: Data collect, Resources, Investigation.

**Cassia Regina Limonta Carvalho:** Data mining analyses, Data curation.

**Maria Imaculada Zucchi:** Conceptualization, Resources, Investigation, Review & Editing, Funding acquisition, Supervision.

## SUPPLEMENTARY INFORMATION

**Supplementary Table 1.**
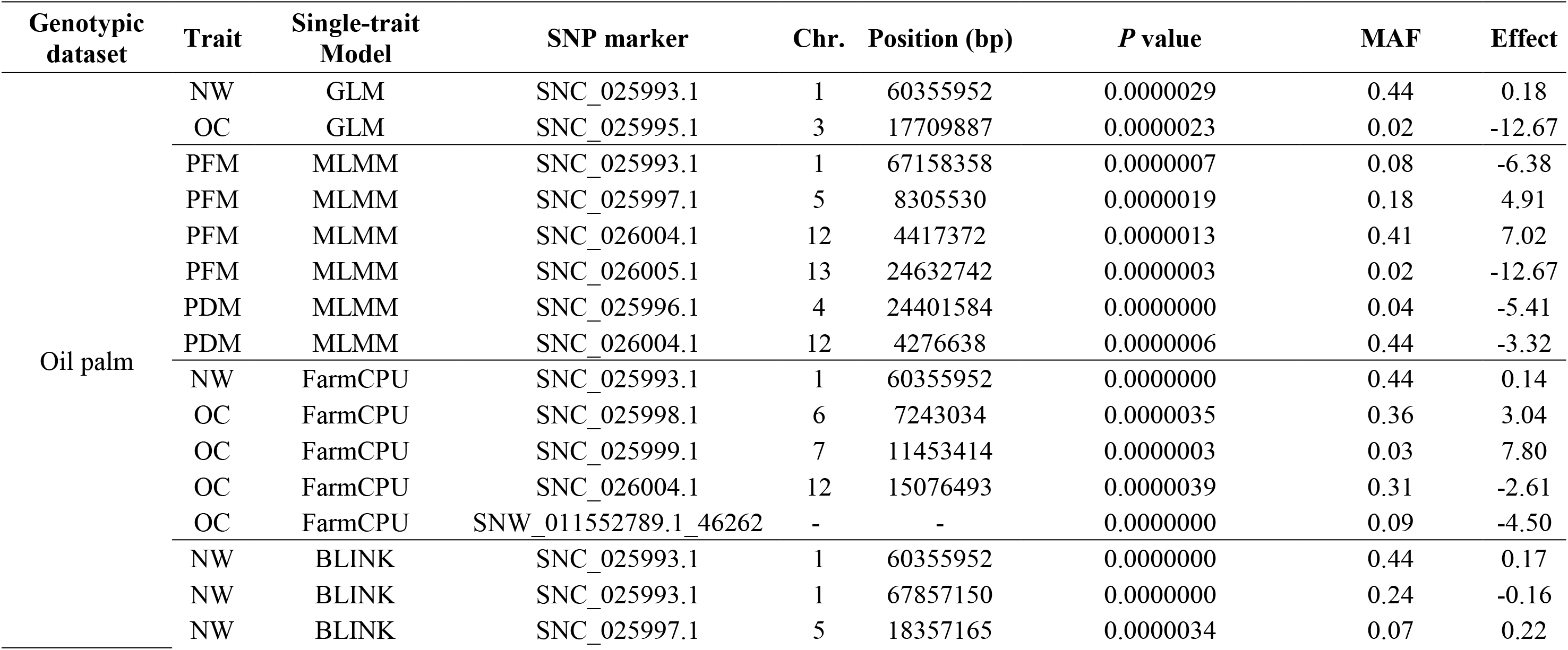
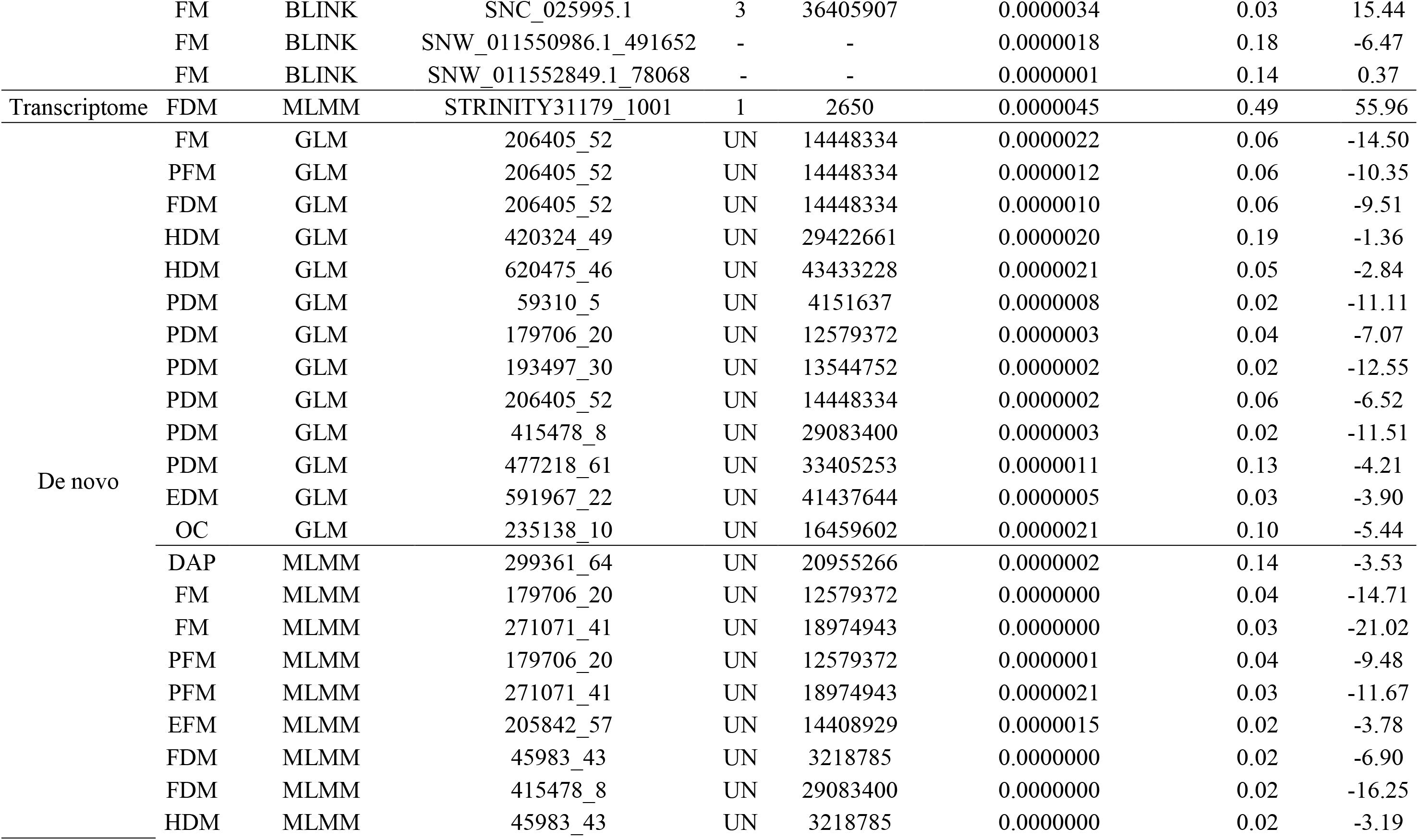
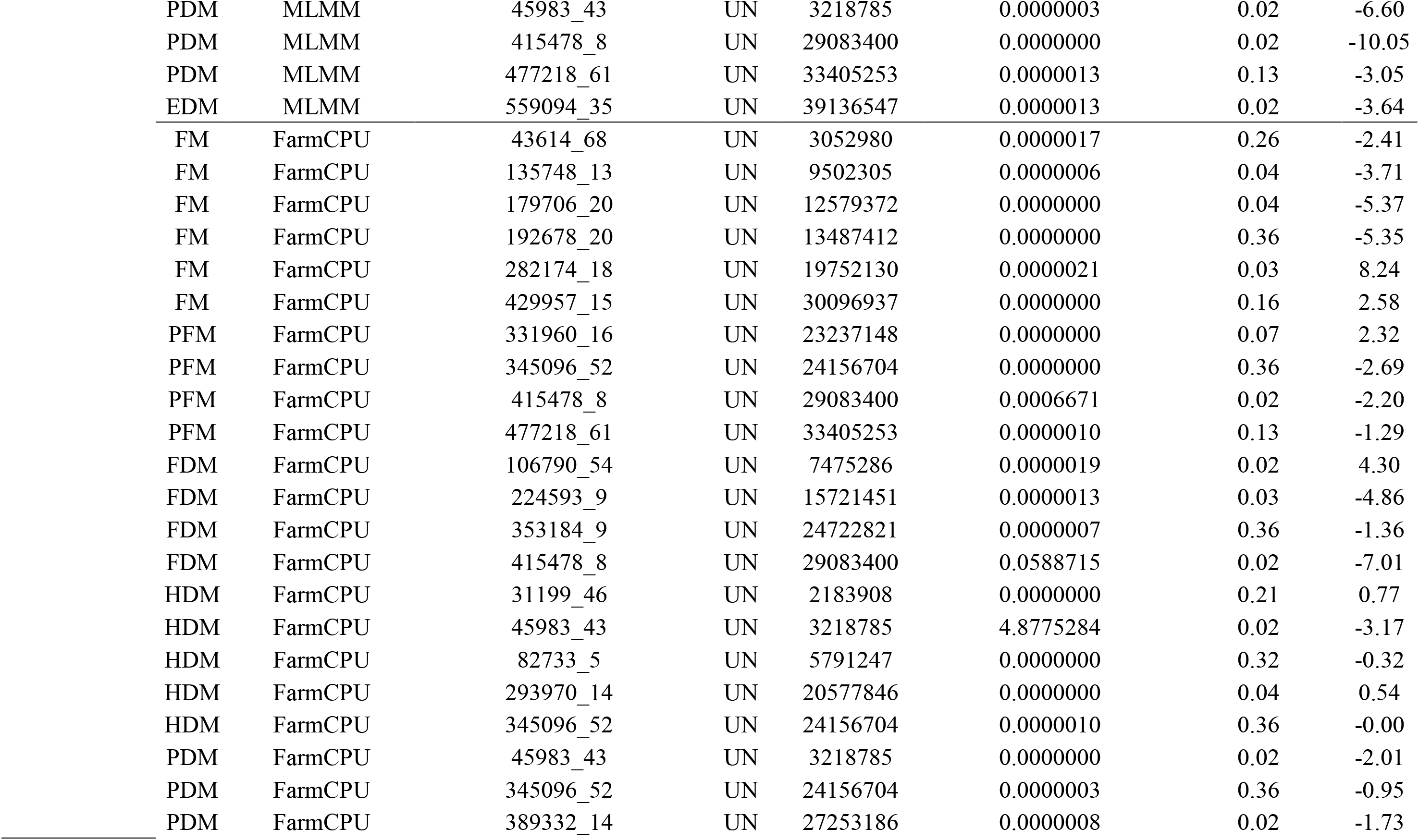
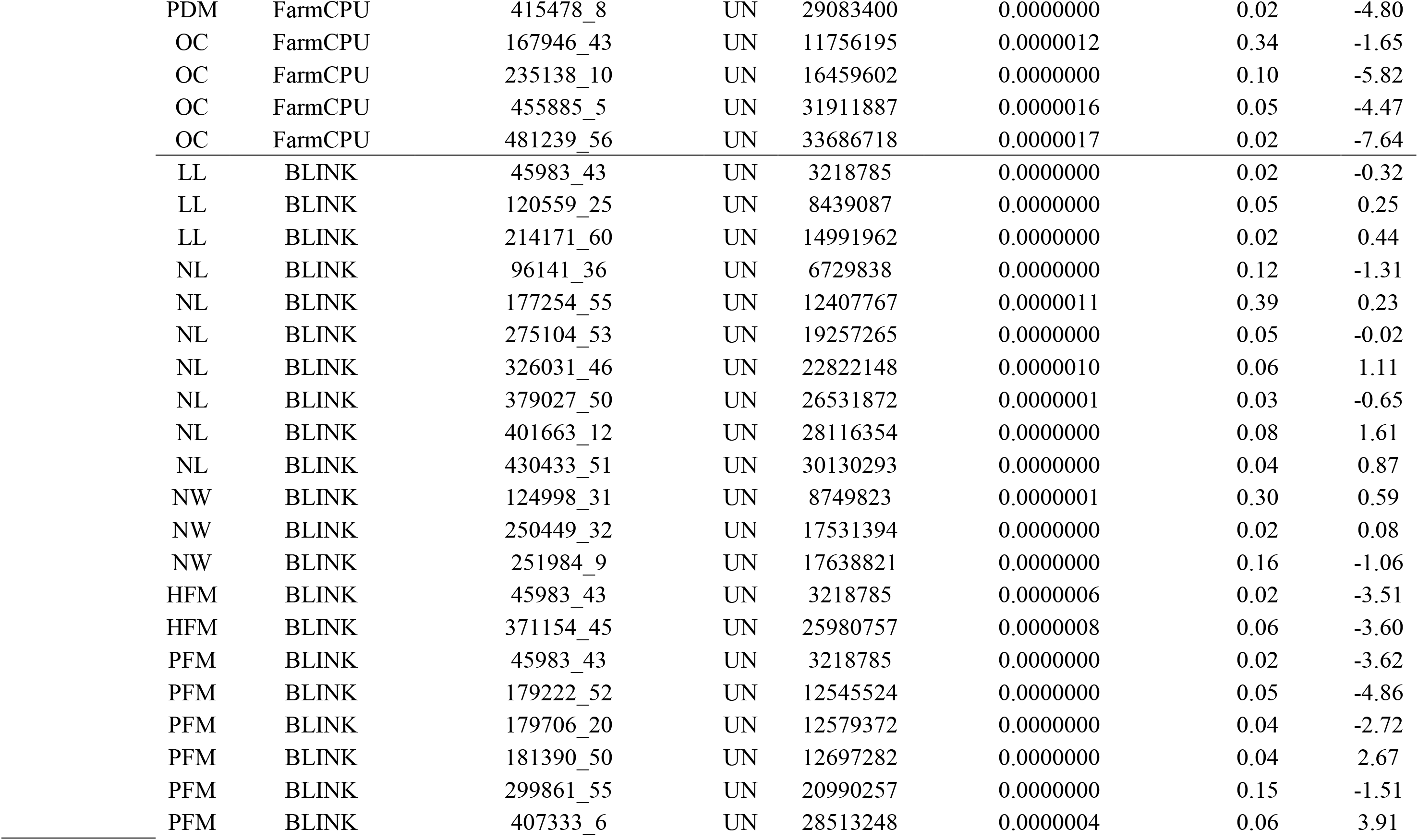
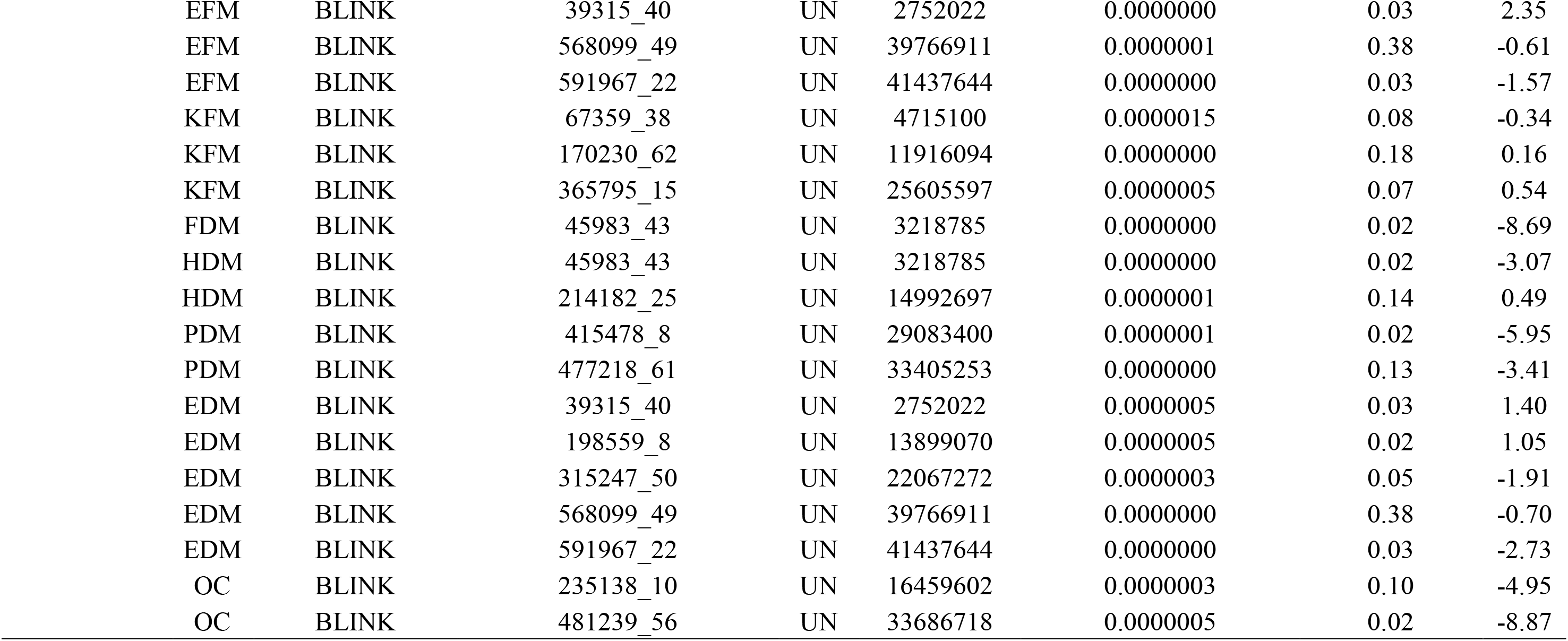
Significant SNP markers detected in the single-trait GWAS models associated with different traits for the three genotypic datasets: oil palm, de novo, and transcriptome. Chromosome (Chr.), position of the marker in base pairs, *p-value*, minor allele frequency (MAF), and effect of each marker.

| Genotypic dataset | Trait | Single-trait Model | SNP marker | Chr. | Position (bp) | *P* value | MAF | Effect |
| --- | --- | --- | --- | --- | --- | --- | --- | --- |
| Oil palm | NW | GLM | SNC_025993.1 | 1 | 60355952 | 0.0000029 | 0.44 | 0.18 |
|  | OC | GLM | SNC_025995.1 | 3 | 17709887 | 0.0000023 | 0.02 | -12.67 |
|  | PFM | MLMM | SNC_025993.1 | 1 | 67158358 | 0.0000007 | 0.08 | -6.38 |
|  | PFM | MLMM | SNC_025997.1 | 5 | 8305530 | 0.0000019 | 0.18 | 4.91 |
|  | PFM | MLMM | SNC_026004.1 | 12 | 4417372 | 0.0000013 | 0.41 | 7.02 |
|  | PFM | MLMM | SNC_026005.1 | 13 | 24632742 | 0.0000003 | 0.02 | -12.67 |
|  | PDM | MLMM | SNC_025996.1 | 4 | 24401584 | 0.0000000 | 0.04 | -5.41 |
|  | PDM | MLMM | SNC_026004.1 | 12 | 4276638 | 0.0000006 | 0.44 | -3.32 |
|  | NW | FarmCPU | SNC_025993.1 | 1 | 60355952 | 0.0000000 | 0.44 | 0.14 |
|  | OC | FarmCPU | SNC_025998.1 | 6 | 7243034 | 0.0000035 | 0.36 | 3.04 |
|  | OC | FarmCPU | SNC_025999.1 | 7 | 11453414 | 0.0000003 | 0.03 | 7.80 |
|  | OC | FarmCPU | SNC_026004.1 | 12 | 15076493 | 0.0000039 | 0.31 | -2.61 |
|  | OC | FarmCPU | SNW_011552789.1_46262 | - | - | 0.0000000 | 0.09 | -4.50 |
|  | NW | BLINK | SNC_025993.1 | 1 | 60355952 | 0.0000000 | 0.44 | 0.17 |
|  | NW | BLINK | SNC_025993.1 | 1 | 67857150 | 0.0000000 | 0.24 | -0.16 |
|  | NW | BLINK | SNC_025997.1 | 5 | 18357165 | 0.0000034 | 0.07 | 0.22 |
|  | FM | BLINK | SNC_025995.1 | 3 | 36405907 | 0.0000034 | 0.03 | 15.44 |
|  | FM | BLINK | SNW_011550986.1_491652 | - | - | 0.0000018 | 0.18 | -6.47 |
|  | FM | BLINK | SNW_011552849.1_78068 | - | - | 0.0000001 | 0.14 | 0.37 |
| Transcriptome | FDM | MLMM | STRINITY31179_1001 | 1 | 2650 | 0.0000045 | 0.49 | 55.96 |
| De novo | FM | GLM | 206405_52 | UN | 14448334 | 0.0000022 | 0.06 | -14.50 |
|  | PFM | GLM | 206405_52 | UN | 14448334 | 0.0000012 | 0.06 | -10.35 |
|  | FDM | GLM | 206405_52 | UN | 14448334 | 0.0000010 | 0.06 | -9.51 |
|  | HDM | GLM | 420324_49 | UN | 29422661 | 0.0000020 | 0.19 | -1.36 |
|  | HDM | GLM | 620475_46 | UN | 43433228 | 0.0000021 | 0.05 | -2.84 |
|  | PDM | GLM | 59310_5 | UN | 4151637 | 0.0000008 | 0.02 | -11.11 |
|  | PDM | GLM | 179706_20 | UN | 12579372 | 0.0000003 | 0.04 | -7.07 |
|  | PDM | GLM | 193497_30 | UN | 13544752 | 0.0000002 | 0.02 | -12.55 |
|  | PDM | GLM | 206405_52 | UN | 14448334 | 0.0000002 | 0.06 | -6.52 |
|  | PDM | GLM | 415478_8 | UN | 29083400 | 0.0000003 | 0.02 | -11.51 |
|  | PDM | GLM | 477218_61 | UN | 33405253 | 0.0000011 | 0.13 | -4.21 |
|  | EDM | GLM | 591967_22 | UN | 41437644 | 0.0000005 | 0.03 | -3.90 |
|  | OC | GLM | 235138_10 | UN | 16459602 | 0.0000021 | 0.10 | -5.44 |
|  | DAP | MLMM | 299361_64 | UN | 20955266 | 0.0000002 | 0.14 | -3.53 |
|  | FM | MLMM | 179706_20 | UN | 12579372 | 0.0000000 | 0.04 | -14.71 |
|  | FM | MLMM | 271071_41 | UN | 18974943 | 0.0000000 | 0.03 | -21.02 |
|  | PFM | MLMM | 179706_20 | UN | 12579372 | 0.0000001 | 0.04 | -9.48 |
|  | PFM | MLMM | 271071_41 | UN | 18974943 | 0.0000021 | 0.03 | -11.67 |
|  | EFM | MLMM | 205842_57 | UN | 14408929 | 0.0000015 | 0.02 | -3.78 |
|  | FDM | MLMM | 45983_43 | UN | 3218785 | 0.0000000 | 0.02 | -6.90 |
|  | FDM | MLMM | 415478_8 | UN | 29083400 | 0.0000000 | 0.02 | -16.25 |
|  | HDM | MLMM | 45983_43 | UN | 3218785 | 0.0000000 | 0.02 | -3.19 |

|  |  |  |  |  |  |  |  |
| --- | --- | --- | --- | --- | --- | --- | --- |
| PDM | MLMM | 45983_43 | UN | 3218785 | 0.0000003 | 0.02 | -6.60 |
| PDM | MLMM | 415478_8 | UN | 29083400 | 0.0000000 | 0.02 | -10.05 |
| PDM | MLMM | 477218_61 | UN | 33405253 | 0.0000013 | 0.13 | -3.05 |
| EDM | MLMM | 559094_35 | UN | 39136547 | 0.0000013 | 0.02 | -3.64 |
| FM | FarmCPU | 43614_68 | UN | 3052980 | 0.0000017 | 0.26 | -2.41 |
| FM | FarmCPU | 135748_13 | UN | 9502305 | 0.0000006 | 0.04 | -3.71 |
| FM | FarmCPU | 179706_20 | UN | 12579372 | 0.0000000 | 0.04 | -5.37 |
| FM | FarmCPU | 192678_20 | UN | 13487412 | 0.0000000 | 0.36 | -5.35 |
| FM | FarmCPU | 282174_18 | UN | 19752130 | 0.0000021 | 0.03 | 8.24 |
| FM | FarmCPU | 429957_15 | UN | 30096937 | 0.0000000 | 0.16 | 2.58 |
| PFM | FarmCPU | 331960_16 | UN | 23237148 | 0.0000000 | 0.07 | 2.32 |
| PFM | FarmCPU | 345096_52 | UN | 24156704 | 0.0000000 | 0.36 | -2.69 |
| PFM | FarmCPU | 415478_8 | UN | 29083400 | 0.0006671 | 0.02 | -2.20 |
| PFM | FarmCPU | 477218_61 | UN | 33405253 | 0.0000010 | 0.13 | -1.29 |
| FDM | FarmCPU | 106790_54 | UN | 7475286 | 0.0000019 | 0.02 | 4.30 |
| FDM | FarmCPU | 224593_9 | UN | 15721451 | 0.0000013 | 0.03 | -4.86 |
| FDM | FarmCPU | 353184_9 | UN | 24722821 | 0.0000007 | 0.36 | -1.36 |
| FDM | FarmCPU | 415478_8 | UN | 29083400 | 0.0588715 | 0.02 | -7.01 |
| HDM | FarmCPU | 31199_46 | UN | 2183908 | 0.0000000 | 0.21 | 0.77 |
| HDM | FarmCPU | 45983_43 | UN | 3218785 | 4.8775284 | 0.02 | -3.17 |
| HDM | FarmCPU | 82733_5 | UN | 5791247 | 0.0000000 | 0.32 | -0.32 |
| HDM | FarmCPU | 293970_14 | UN | 20577846 | 0.0000000 | 0.04 | 0.54 |
| HDM | FarmCPU | 345096_52 | UN | 24156704 | 0.0000010 | 0.36 | -0.00 |
| PDM | FarmCPU | 45983_43 | UN | 3218785 | 0.0000000 | 0.02 | -2.01 |
| PDM | FarmCPU | 345096_52 | UN | 24156704 | 0.0000003 | 0.36 | -0.95 |
| PDM | FarmCPU | 389332_14 | UN | 27253186 | 0.0000008 | 0.02 | -1.73 |
| PDM | FarmCPU | 415478_8 | UN | 29083400 | 0.0000000 | 0.02 | -4.80 |
| OC | FarmCPU | 167946_43 | UN | 11756195 | 0.0000012 | 0.34 | -1.65 |
| OC | FarmCPU | 235138_10 | UN | 16459602 | 0.0000000 | 0.10 | -5.82 |
| OC | FarmCPU | 455885_5 | UN | 31911887 | 0.0000016 | 0.05 | -4.47 |
| OC | FarmCPU | 481239_56 | UN | 33686718 | 0.0000017 | 0.02 | -7.64 |
| LL | BLINK | 45983_43 | UN | 3218785 | 0.0000000 | 0.02 | -0.32 |
| LL | BLINK | 120559_25 | UN | 8439087 | 0.0000000 | 0.05 | 0.25 |
| LL | BLINK | 214171_60 | UN | 14991962 | 0.0000000 | 0.02 | 0.44 |
| NL | BLINK | 96141_36 | UN | 6729838 | 0.0000000 | 0.12 | -1.31 |
| NL | BLINK | 177254_55 | UN | 12407767 | 0.0000011 | 0.39 | 0.23 |
| NL | BLINK | 275104_53 | UN | 19257265 | 0.0000000 | 0.05 | -0.02 |
| NL | BLINK | 326031_46 | UN | 22822148 | 0.0000010 | 0.06 | 1.11 |
| NL | BLINK | 379027_50 | UN | 26531872 | 0.0000001 | 0.03 | -0.65 |
| NL | BLINK | 401663_12 | UN | 28116354 | 0.0000000 | 0.08 | 1.61 |
| NL | BLINK | 430433_51 | UN | 30130293 | 0.0000000 | 0.04 | 0.87 |
| NW | BLINK | 124998_31 | UN | 8749823 | 0.0000001 | 0.30 | 0.59 |
| NW | BLINK | 250449_32 | UN | 17531394 | 0.0000000 | 0.02 | 0.08 |
| NW | BLINK | 251984_9 | UN | 17638821 | 0.0000000 | 0.16 | -1.06 |
| HFM | BLINK | 45983_43 | UN | 3218785 | 0.0000006 | 0.02 | -3.51 |
| HFM | BLINK | 371154_45 | UN | 25980757 | 0.0000008 | 0.06 | -3.60 |
| PFM | BLINK | 45983_43 | UN | 3218785 | 0.0000000 | 0.02 | -3.62 |
| PFM | BLINK | 179222_52 | UN | 12545524 | 0.0000000 | 0.05 | -4.86 |
| PFM | BLINK | 179706_20 | UN | 12579372 | 0.0000000 | 0.04 | -2.72 |
| PFM | BLINK | 181390_50 | UN | 12697282 | 0.0000000 | 0.04 | 2.67 |
| PFM | BLINK | 299861_55 | UN | 20990257 | 0.0000000 | 0.15 | -1.51 |
| PFM | BLINK | 407333_6 | UN | 28513248 | 0.0000004 | 0.06 | 3.91 |
| EFM | BLINK | 39315_40 | UN | 2752022 | 0.0000000 | 0.03 | 2.35 |
| EFM | BLINK | 568099_49 | UN | 39766911 | 0.0000001 | 0.38 | -0.61 |
| EFM | BLINK | 591967_22 | UN | 41437644 | 0.0000000 | 0.03 | -1.57 |
| KFM | BLINK | 67359_38 | UN | 4715100 | 0.0000015 | 0.08 | -0.34 |
| KFM | BLINK | 170230_62 | UN | 11916094 | 0.0000000 | 0.18 | 0.16 |
| KFM | BLINK | 365795_15 | UN | 25605597 | 0.0000005 | 0.07 | 0.54 |
| FDM | BLINK | 45983_43 | UN | 3218785 | 0.0000000 | 0.02 | -8.69 |
| HDM | BLINK | 45983_43 | UN | 3218785 | 0.0000000 | 0.02 | -3.07 |
| HDM | BLINK | 214182_25 | UN | 14992697 | 0.0000001 | 0.14 | 0.49 |
| PDM | BLINK | 415478_8 | UN | 29083400 | 0.0000001 | 0.02 | -5.95 |
| PDM | BLINK | 477218_61 | UN | 33405253 | 0.0000000 | 0.13 | -3.41 |
| EDM | BLINK | 39315_40 | UN | 2752022 | 0.0000005 | 0.03 | 1.40 |
| EDM | BLINK | 198559_8 | UN | 13899070 | 0.0000005 | 0.02 | 1.05 |
| EDM | BLINK | 315247_50 | UN | 22067272 | 0.0000003 | 0.05 | -1.91 |
| EDM | BLINK | 568099_49 | UN | 39766911 | 0.0000000 | 0.38 | -0.70 |
| EDM | BLINK | 591967_22 | UN | 41437644 | 0.0000000 | 0.03 | -2.73 |
| OC | BLINK | 235138_10 | UN | 16459602 | 0.0000003 | 0.10 | -4.95 |
| OC | BLINK | 481239_56 | UN | 33686718 | 0.0000005 | 0.02 | -8.87 |

**Supplementary Table 2.**
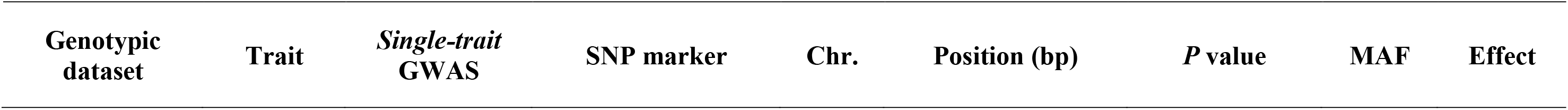
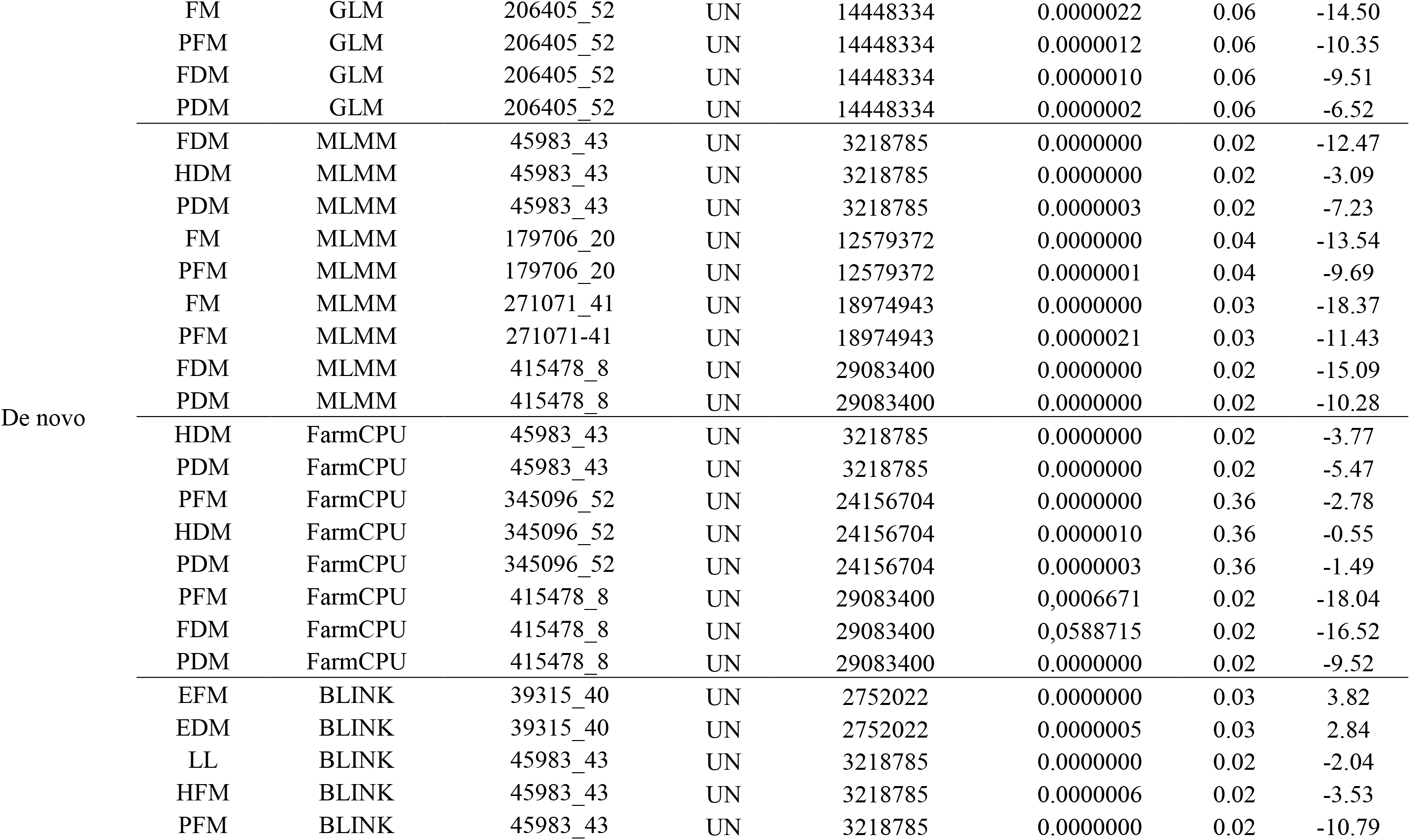
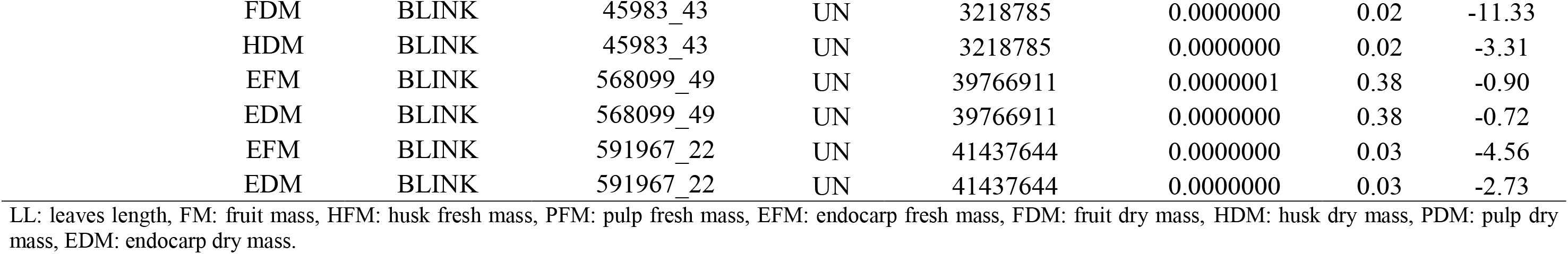
The same SNP markers associated with different traits in the same single-trait GWAS model for the de novo genotypic dataset. Chromosome (Chr.), position of the marker in base pairs, *p-value*, minor allele frequency (MAF), and effect of each marker.

**Supplementary Table 3.** Significant SNP markers detected in the multi-trait GWAS MSTEP model associated with different trait combinations for the three genotypic datasets: oil palm, de novo, and transcriptome. Chromosome (Chr.), position of the marker in base pairs, of each marker

| Genotypic dataset | Trait 1 | Trait 2 | SNP marker | Chr. | Position (bp) | approx_F | num_df | den_df | probF |
| --- | --- | --- | --- | --- | --- | --- | --- | --- | --- |
| Oil palm | H | EFM | SNW_011557734.1_1319 | - | - | 24.46 | 2 | 155 | 0.0000000 |
|  | H | NW | SNC_025993.1 | 1 | 60355952 | 8.87 | 4 | 304 | 0.0000008 |
|  | LL | OC | SNC_025995.1 | 3 | 17709887 | 16.53 | 2 | 153 | 0.0000003 |
|  | DAP | NW | SNC_025993.1 | 1 | 60355952 | 7.53 | 4 | 304 | 0.0000084 |
|  | NW | HFM | SNW_011550905.1_990562 | - | - | 8.68 | 4 | 304 | 0.0000012 |
|  | NW | HDM | SNW_011550905.1_990562 | - | - | 8.09 | 4 | 304 | 0.0000032 |
|  | LN | HFM | SNC_026000.1 | 8 | 5081018 | 13.14 | 2 | 155 | 0.0000053 |
|  | LN | OC | SNC_025995.1 | 3 | 17709887 | 14.85 | 2 | 153 | 0.0000012 |
|  | NN | HFM | SNC_026000.1 | 8 | 5081018 | 20.11 | 2 | 152 | 0.0000000 |
|  | NN | FM | SNC_026000.1 | 8 | 5081018 | 17.19 | 2 | 155 | 0.0000001 |
|  | NN | HDM | SNC_026000.1 | 8 | 5081018 | 14.31 | 2 | 155 | 0.0000019 |
| De novo | HDM | PDM | 408997_47 | UN | 28629769 | 9.36 | 4 | 294 | 0.0000003 |
|  | HDM | PDM | 333769_9 | UN | 23363771 | 8.24 | 4 | 294 | 0.0000025 |
|  | FDM | OC | 333769_9 | UN | 23363771 | 9.05 | 4 | 298 | 0.0000006 |
|  | FDM | HDM | 333769_9 | UN | 23363771 | 9.83 | 4 | 294 | 0.0000001 |
|  | FDM | PDM | 408997_47 | UN | 28629769 | 8.34 | 4 | 298 | 0.0000021 |
|  | FDM | KDM | 477218_61 | UN | 33405253 | 13.13 | 4 | 290 | 0.0000000 |
| Transcriptome | FDM | OC | STRINITY9279_665 | 1 | 4262 | 8.68 | 4 | 322 | 0.0000011 |
|  | FDM | HDM | STRINITY9279_665 | 1 | 4262 | 6.59 | 4 | 326 | 0.0000412 |

## REFERENCES

1. Abreu A.G., Priolli R.H.G, Azevedo-Filho J.A., Nucci S.M., Zucchi M.I., Coelho R.M., Colombo A.C. The Genetic Structure and Mating System of Acrocomia Aculeata (Arecaceae) (2012). Genetics and Molecular Biology, 35(1), 116–121.

2. Andrews S. FASTQC. A quality control tool for high throughput sequence data (2010). https://www.bioinformatics.babraham.ac.uk/projects/fastqc/.

3. Bates D, Mächler M, Bolker B, Walker S. Fitting Linear Mixed-Effects Models Using lme4 (2015). Journal of Statistical Software, 67(1), 1–48.

4. Bazzo B.R., Carvalho L.M., Carazzolle M.F., Pereira G.A.G., Colombo C.A. Development of Novel EST-SSR Markers in the Macaúba Palm (Acrocomia Aculeata) Using Transcriptome Sequencing and Cross-Species Transferability in Arecaceae Species (2018). BMC Plant Biology, 8(1).

5. Berton, Luiz Henrique Chorfi. Avaliação de populações naturais, estimativas de parâmetros genéticos e seleção de genótipos elite de macaúba (*Acrocomia aculeata*) (2013) Tese. Campinas.

6. Bradbury P.J., Zhang Z., Kroon D.E., Casstevens T.M., Ramdoss Y., & Buckler E.S. TASSEL: Software for association mapping of complex traits in diverse samples (2007). Bioinformatics, 23, 2633–2635.

7. Browning, B.L., Tian X., Zhou Y., Browning S.R. Fast two-stage phasingof large-scale sequence data (2021). American Journal of Human Genetics108(10), 1880-1890.

8. Cao, Y.; Han, Y.; Meng, D.; Abdullah, M.; Yu, J.; Li, D.; Jin, Q.; Lin, Y.; Cai, Y. Expansion and evolutionary patterns of GDSL-type esterases/lipases in Rosaceae genomes (2018). Funct. Integr. Genom., 18, 673–684.

9. Catchen J.M., Amores A., Hohenlohe P., Cresko W., Postlethwait J.H. Stacks: Building and genotyping loci de novo from shortread sequences (2011). G3: Genes, Genomes, Genetics, 1(3).

10. Cichonska A., Rousu J., Marttinen P., Kangas A. J., Soininen P., Lehtimäki T., et al. metaCCA: summary statistics-based multivariate metaanalysis of genome-wide association studies using canonical correlation analysis (2016). Bioinformatics 32, 1981–1989.

11. Ciconini G., Favaro S.P., Roscoe R., Miranda C.H.B., Tapeti C.F., Miyahira M.A.M., Bearari L., Galvani F., Borsato A.V., Colnago L.A., Naka M.H. Biometry and Oil Contents of Acrocomia Aculeata Fruits from Cerrados and Pantanal Biomes in Mato Grosso Do Sul, Brazil (2013). Industrial Crops and Products, 45, 208-214,

12. Ciftci-Yilmaza S., and Mittler R. The zinc finger network of plants (2008). Cellular and Molecular Life Sciences. 65, 1150 – 1160.

13. Coimbra M.C., Jorge N. Fatty Acids and Bioactive Compounds of the Pulps and Kernels of Brazilian Palm Species, Guariroba (Syagrus Oleraces), Jerivá (Syagrus Romanzoffiana) and Macaúba (Acrocomia Aculeata) (2012). Journal of the Science of Food and Agriculture, 92(3),679–684.

14. Colombo C.A., Berton L.H.C., Diaz B.G., Ferrari R.A. Macauba: A Promising Tropical Palm for the Production of Vegetable Oil (2018). OCL, 25(1), 1–9.

15. Conesa A., Gotz S., Garcia-Gomez J.M., Terol J., Talon M., Robles M. Blast2GO: a universal tool for annotation, visualization and analysis in functional genomics research (2005). Bioinformatics, 21: 3674–3676.

16. Coser S.M., Motoike S.Y., Corrêa T.R., Pires T.P., Resende M.D.V. Breeding of Acrocomia aculeata using genetic diversity parameters and correlations to select accessions based on vegetative, phenological, and reproductive characteristics (2016). Genetic Molecular Research, 15(4).

17. Covarrubias-Pazaran G. Genome assisted prediction of quantitative traits using the R package sommer (2016). PLoS ONE, 11, 1–15.

18. Cruz C.D., Regazzi A.J, Carneiro P.C.S. Modelos biométricos aplicados ao melhoramento genético. 4. ed. Viçosa, MG: Ed. UFV, 2012.

19. Cruz CD, Ferreira FM, Pessoni LA. Biometria aplicada ao estudo da diversidade genética (2011). 1a ed. Visconde do Rio Branco-MG: Suprema. 620 p.

20. Diaz B.G., Zucchi M.I., Alves-Pereira A., Almeida C.P.A., Moraes A.C.L., Vianna S.A., Azevedo-Filho J., Colombo C.A. Genome-wide SNP analysis to assess the genetic population structure and diversity of Acrocomia species (2021). PLoS ONE, 16(7): 1–24.

21. Ding, L.N.; Wang, W.J.; Cao, J.; Wang, Z.; Zhu, K.M.; Yang, Y.H.; Li, Y.L.; Tan, X.L. Advances in plant GDSL lipases: From sequences to functional mechanisms (2019). Acta Physiol. Plant. 2019, 41, 151.

22. Domiciano G.P., Alves A.A., Laviola B.G., da Conceição L.D.H.C.S. Genetic parameters and diversity in progenies from macaw palm based on morphological and physiological traits [Parâmetros genéticos e diversidade em progênies de macaúba com base em características morfológicas e fsiológicas] (2015). Ciência Rural 45(9), 159–1605.

23. Doyle J.J., Doyle J.L. Isolation ofplant DNA from fresh tissue (1990). Focus, 12(13), 39–40.

24. Falconer D.S., Mackay T.F.C. Introduction to quantitative genetics. 4 ed. ed. New York, NY: Longman Group Limited: Edinburgh, 1996.

25. Farias Neto JT, Clement CR, Resende MDV. Estimativas de parâmetros genéticos e ganho de seleção para produção de frutos em progênies de polinização aberta de pupunheira no estado do Pará, Brasil (2013). Bragantia. 32, 122-126.

26. Francisconi A.F., Marroquín J.A.M., Cauz-Santos L.A. et al. Complete chloroplast genomes of six neotropical palm species, structural comparison, and evolutionary dynamic patterns (2023). Scientific Reports, 13, 20635.

27. François O., Caye K. Naturalgwas: An R package for evaluating genomewide association methods with empirical data (2018). Molecular Ecology Resources, Special issue: Association mapping in natural populations.

28. Fernandes, S.B., Dias, K.O.G., Ferreira, D.F. et al. Efficiency of multi-trait, indirect, and trait-assisted genomic selection for improvement of biomass sorghum (2018). Theoretical and Applied Genetics, 131, 747–755.

29. Fernandes S.B., Zhang K.S., Jamann T.M. and Lipka A.E. How Well Can Multivariate and Univariate GWAS Distinguish Between True and Spurious Pleiotropy? (2021). Frontiers in Genetics, 11:602526.

30. Fernandes S.B., Casstevens T.M., Bradbury P.J., Lipka A.E. A multi-trait multi-locus stepwise approach for conducting GWAS on correlated traits (2022). Plant Genome, 15:e20200

31. Filzmoser P., Ruiz-Gazen A., and Thomas-Agnan C.: Identification of local multivariate outliers. Submitted for publication, 2012.

32. Furlotte N.A., & Eskin E. Efficient multiple-trait association and estimation of genetic correlation using the matrix-variate linear mixed model (2015). Genetics, 200, 59–68.

33. Garrison E., Marth G. Haplotype-based variant detection from short-read sequencing (2012). arXiv preprint arXiv:1207.3907.

34. Goudet J. Hierfstat, a package for R to compute and test hierarchical F-statistics (2005). Molecular Ecology Notes, 5: 184–186.

35. Hassan H., Amiruddin M.D, Weckwerth W., Ramli U.S. Deciphering key proteins of oil palm (Elaeis guineensis Jacq.) fruit mesocarp development by proteomics and chemometrics (2019). Electrophoresis 40 (2), 254-265.

36. Hartl F.U. Molecular chaperones in cellular protein folding (1996). Nature, 381, 571–579

37. Hartl, F. Chaperone-assisted protein folding: the path to discovery from a personal perspective (2011). Nat Med 17, 1206–1210.

38. Heberle, H., Meirelles, G.V., da Silva, F.R. et al. InteractiVenn: a web-based tool for the analysis of sets through Venn diagrams (2015). BMC Bioinformatics 16, 169.

39. Hiane P.A., Filho M.M.R., Ramos M.I.L, Macedo M.L. Bocaiuva, Acrocomia aculeata (Jacq.) Lodd., pulp and kernel oils: characterization and fatty acid composition (2005). Brazilian journal of food technology, p. 256–259.

40. Huang M., Liu X., Zhou Y., Summers R.M., Zhang Z. BLINK: a package for the next level of genome-wide association studies with both individuals and markers in the millions (2019). Gigascience, 8, 1–12.

41. Jombart T. “adegenet: a R package for the multivariate analysis of genetic markers.” (2008). Bioinformatics, 24, 1403-1405.

42. Jombart T., Ahmed I. “adegenet 1.3-1: new tools for the analysis of genome-wide SNP data.” (2011). Bioinformatics, 27(21), 3070-1.

43. Joo J.W.J., Kang E.Y., Org E., Furlotte N., Parks B., Hormozdiari F., et al. Efficient and accurate multiple-phenotype regression method for high dimensional data considering population structure (2016). Genetics 204, 1379–1390.

44. Lee C., Teng Q., Zhong R., Ye Z.H. The four Arabidopsis reduced wall acetylation genes are expressed in secondary wall-containing cells and required for the acetylation of xylan (2011). Plant Cell Physiol 52, 1289–1301.

45. Leuenberger D., Bally N. A, Schatz G., Koehler C.M. Different import pathways through themitochondrial intermembrane space for innermembrane proteins (1999). The EMBO Journal, 18 (17), 4816–4822.

46. Li H., Handsaker B., Wysoker A., Fennell T., Ruan J., Homer N., Marth G., Abecasis G., Durbin R., Subgroup 1000 Genome Project Data Processing. The Sequence Alignment/Map format and SAMtools (2009). Bioinformatics 25, 2078-2079.

47. Li H. Aligning sequence reads, clone sequences and assembly contigs with BWA-MEM 2013. arXiv:1303.3997v2.

48. Lima N.E., Carvalho A.A., Meerow A.W. et al. A review of the palm genus *Acrocomia*: Neotropical green gold (2018). Organisms Diversity & Evolution, 18, 151–161.

49. Liu X., Huang M., Fan B., Buckler E.S., Zhang Z. Iterative usage of fixed and random effect models for powerful and efficient genome- wide association studies (2016). PLoS Genet,12:e1005767

50. Lorenzi H., Noblick L., Kahn F., & Ferreira, E. Flora Brasileira: Arecaceae (palmeiras). Nova Odessa, SP: Instituto plantarum, 2010.

51. Mackay T.F.C. Quantitative Trait Loci in Drosophila (2001). Nature Reviews Genetics, 2(1), 11–20.

52. Malosetti M., Ribaut J.M., Vargas M. et al. A Multi-Trait Multi-Environment QTL Mixed Model with an Application to Drought and Nitrogen Stress Trials in Maize (Zea Mays L.) (2008). Euphytica, 161, 241-257.

53. Manabe Y., Nafisi M., Verhertbruggen Y., Orfila C., Gille S., Rautengarten C., Cherk C., Marcus S.E., Somerville S., Pauly M., et al. Loss-of-function mutation of REDUCED WALL ACETYLATION2 in Arabidopsis leads to reduced cell wall acetylation and increased resistance to Botrytis cinérea (2011). Plant Physiol 155, 1068– 1078

54. Mazzottini-dos-Santos H.C., Ribeiro L.M., Mercadante-Simões M.O. et al. Floral structure in Acrocomia aculeata (Arecaceae): evolutionary and ecological aspects (2015). Plant Syst Evol 301, 1425–1440.

55. Montoya, S.G., Motoike, S.Y., Kuki, K.N. et al. Fruit development, growth, and stored reserves in macauba palm (Acrocomia aculeata), an alternative bioenergy crop (2016). Planta 244, 927–938.

56. Naested H., Holm A., Jenkins T., Nielsen H.B., Harris C.A., Beale M.H., Andersen M., Mant A., Scheller H., Camara B., Mattsson O., Mundy J. Arabidopsis VARIEGATED 3 encodes a chloroplast-targeted, zinc-finger protein required for chloroplast and palisade cell development (2004). J Cell Sci, 15 (117), 4807–18.

57. Nafisi M., Stranne M., Fimognari L., Atwell S. , Martens H. J., Pedas P. R., Hansen S. F., Nawrath C., Scheller H. V., Kliebenstein D. J., Sakuragi Y. Acetylation of cell wall is required for structural integrity of the leaf surface and exerts a global impact on plant stress responses (2015). Frontiers in Plant Science, 6.

58. Oraguzie N.C. et al. Association Mapping in Plants. New York, NY: Springer New York, 2007.

59. Peterson R.A., Cavanaugh J.E. “Ordered quantile normalization: a semiparametric transformation built for the cross-validation era.” (2020). Journal of Applied Statistics, 47(13-15), 2312-2327.

60. Poland J.A., Rife T.W. Genotyping-by-Sequencing for Plant Breeding and Genetics (2012). The Plant Genome, 5 (3).

61. Poland J.A., Brown P.J., Sorrells M.E., Jannink J.L. Development of high-density genetic maps for barley and wheat using a novel two-enzyme genotyping-by-sequencing approach (2012). PloS one, 7(2), e32253.

62. Porter H.F., and O’Reilly P. F. Multivariate simulation framework reveals performance of multi-trait GWAS methods (2017). Scientific Reports, 7, 38837.

63. Pritikin J.N., Neale M.C., Prom-Wormley, E.C., Clark, S.L., & Verhulst, B. GW-SEM 2.0: Efficient, flexible, and accessible multivariate GWAS (2021). Behavior Genetics, 51, 343–357.

64. Quinlan A.R. and Hall I.M. BEDTools: a flexible suite of utilities for comparing genomic features (2012). Bioinformatics. 26(6), 841–842.

65. R Core Team. R: A language and environment for statistical computing (2021). R Foundation for Statistical Computing, Vienna, Austria. URL https://www.R-project.org/.

66. Reis SB, Mercadante-Simoes MO, Ribeiro LM. Pericarp development in the macaw palm *Acrocomia aculeata* (Arecaceae) (2012). Rodriguesia 63(3), 541–549.

67. Rosado RDS, Rosado TB, Cruz CD, Ferraz AG, da Conceição LDHCS, Laviola BG Genetic parameters and simultaneous selection for adaptability and stability of macaw palm (2019). Scie HorticAmsterdam 248, 291-296

68. Scariot A., Lleras E., HAY J.D. Flowering and Fruiting Phenologies of the Palm Acrocomia aculeata: Patterns and Consequences (1995). Biotropica, 27(2), 168.

69. Sharma S.K., Rios P.D., Christen P., Lustig A., Goloubinoff P. The kinetic parameters and energy cost of the Hsp70 chaperone as a polypeptide unfoldase (2010) Nat. Chem. Biol., 6, 914–920

70. Silva J.C., Barrichelo L.E.G., Brito J.O. Endocarpos de Macaúba e de Babaçu comparados a madeira de Eucaliptus grandis na produção de carvão vegetal (1986). IPEF, 34, 31–34.

71. Singh R., Ong-Abdullah M., Low E-TL., Manaf M.A.A., Rosli R., Nookiah R., et al. Oil palm genome sequence reveals divergence of interfertile species in old and new worlds (2013). Nature, 500:335–9.

72. Stich B., Melchinger A.E., An introduction to association mapping in plants (2010). CABI Reviews, 5, No. 039

73. Teixeira L.C. Potencialidades de oleaginosas para produção de biodiesel (2005). Informe Agropecuário, 26, 18–27.

74. Vargas-Carpintero, R., Hilger T., Mössinger J., Souza R.F., Armas J.C.B.A., Tiede K., Lewandowski I. *Acrocomia* spp.; negleted crop, ballyhooed multipurpose palm or fit for the bioeconomy? A review (2021). Agronomy for sustainable development, 41 (6), 75.

75. Wang Q., Tian F., Pan Y., Buckler E. S., and Zhang Z. A SUPER powerful method for genome wide association study (2014). PLoS ONE, 9, e107684.

76. Wang, J. & Zhang, Z. GAPIT Version 3: Boosting Power and Accuracy for Genomic Association and Prediction (2021). Genomics Proteomics Bioinformatics 19, 1–12.

77. Yu J. et al. A Unified Mixed-Model Method for Association Mapping That Accounts for Multiple Levels of Relatedness (2006). Nature Genetics, 38(2), 203–208.

78. Zhang H., Ma X., Shi T., Song Q., Zhao H., Ma D. NSA2, a novel nucleolus protein regulates cell proliferation and cell cycle (2010). Biochemical and Biophysical Research Communications, 391(1), 651–658,

79. Zhang K., Yao D., Chen Y., Wen H., Pan J., Xiao T., Lv D., He H., Pan J., Cai R., Wang G. Mapping and identification of CsSF4, a gene encoding a UDP-N-acetyl glucosamine-peptide N-acetylglucosaminyltransferase required for fruit elongation in cucumber (Cucumis sativus L.) (2023). Theor Appl Genet, 13, 136(3):54.

80. Zhang Z., Ersoz E., Lai C. Q., Todhunter R. J., Tiwari H. K., Gore M. A., et al. Mixed linear model approach adapted for genome-wide association Studies (2010). Nature Genetics, 42, 355–360.

81. Zhou X., Stephens M. Efficient Multivariate Linear Mixed Model Algorithms for Genome-Wide Association Studies (2014). Nature Methods, 11(4), 407–409.

82. Zhu C. et al. Status and Prospects of Association Mapping in Plants (2008). The Plant Genome, 1(1), 5.

