## Supplementary material_Couto et al. for "Genome-Wide Association Insights into the Genomic Regions Controlling Oil Production Traits in *Acrocomia aculeata* (neotropical native palm)"

### GWAS on Macauba

#Multiple trait GWAS figures

---

#### ## GLM Dende

```
##  Rectangular-Manhattan Plotting Altura.
```

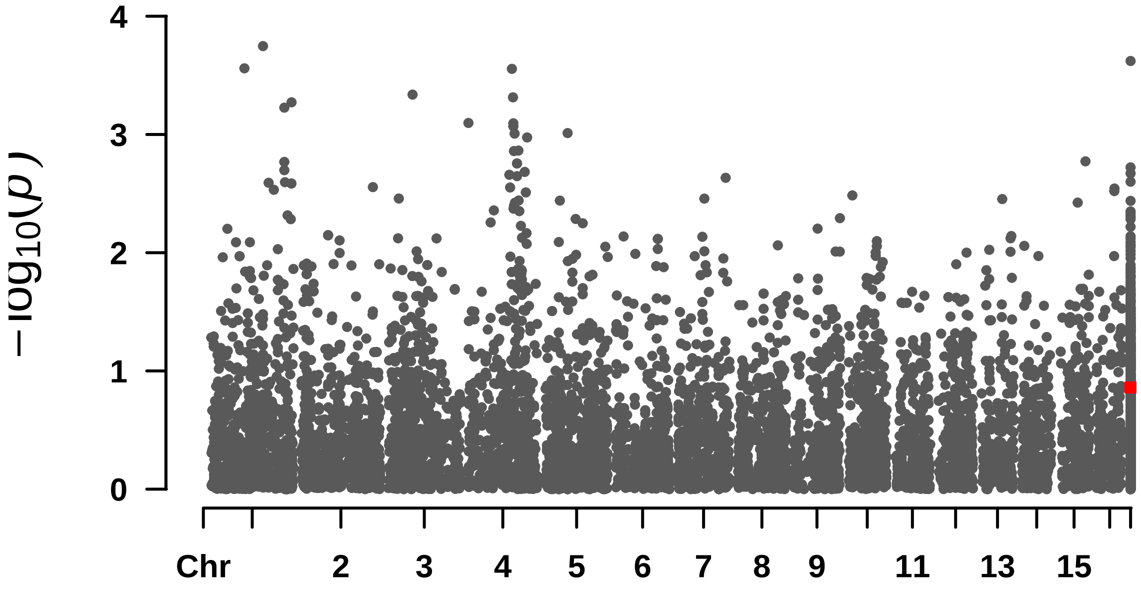


```
##  Rectangular-Manhattan Plotting F_Endo.
```

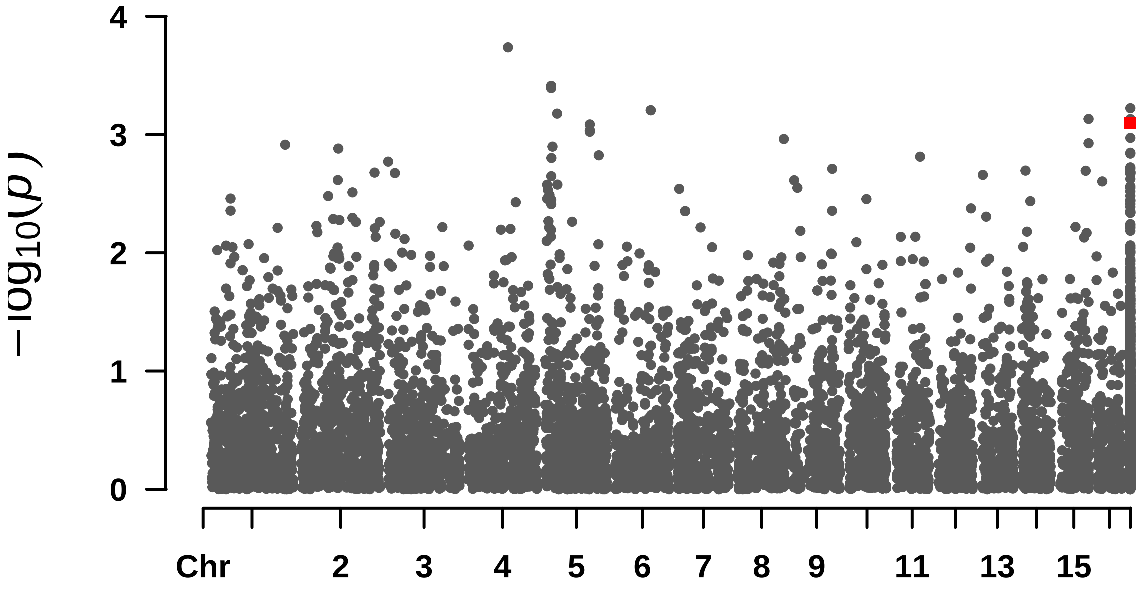

```
##
##
## [1] "===================================================================="
##  Rectangular-Manhattan Plotting Altura.
```

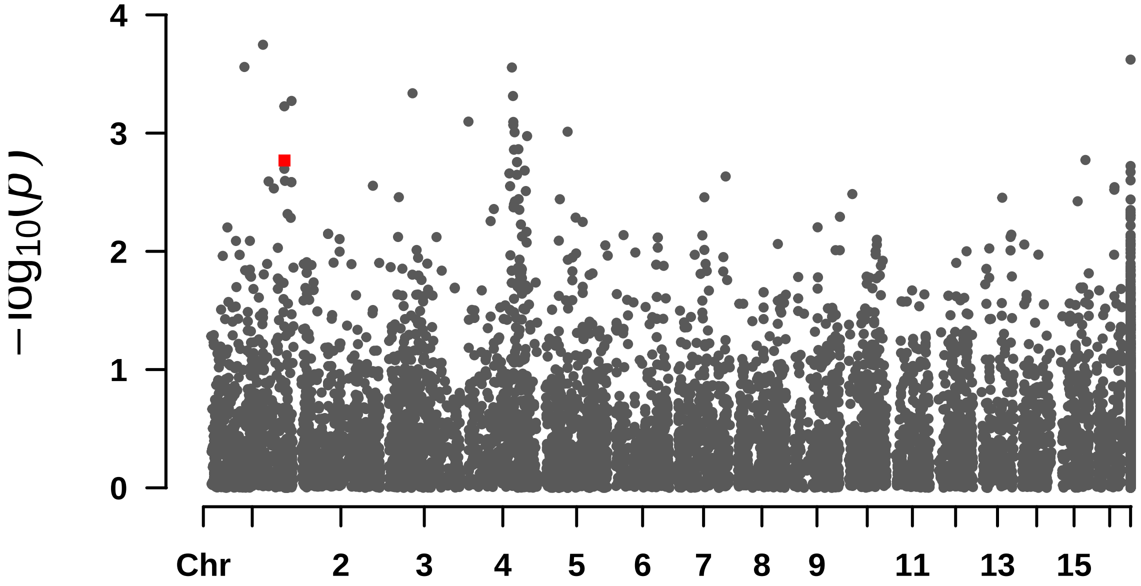


```
##  Rectangular-Manhattan Plotting DAP.
```

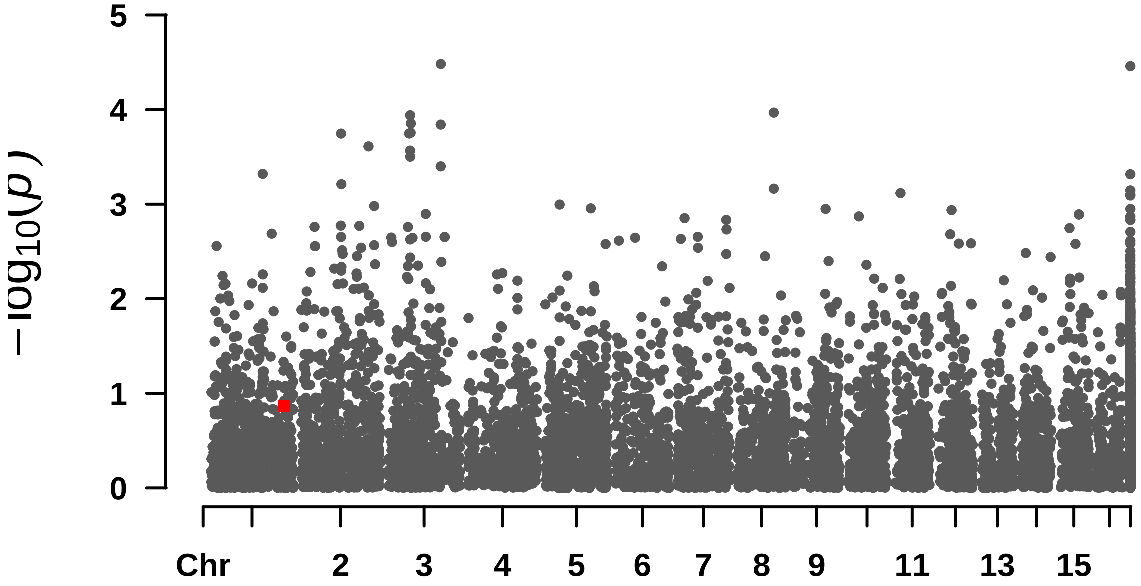


```
##  Rectangular-Manhattan Plotting Largura_pina.
```

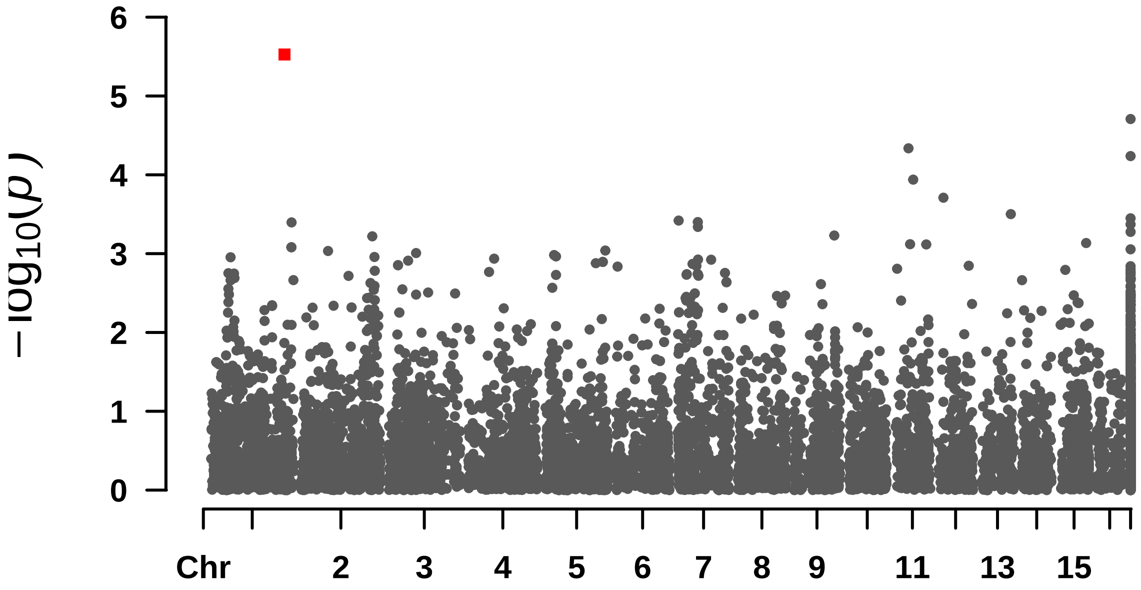


```
##
##
## [1] "=================================================================="
##  Rectangular-Manhattan Plotting Compri_folha.
```

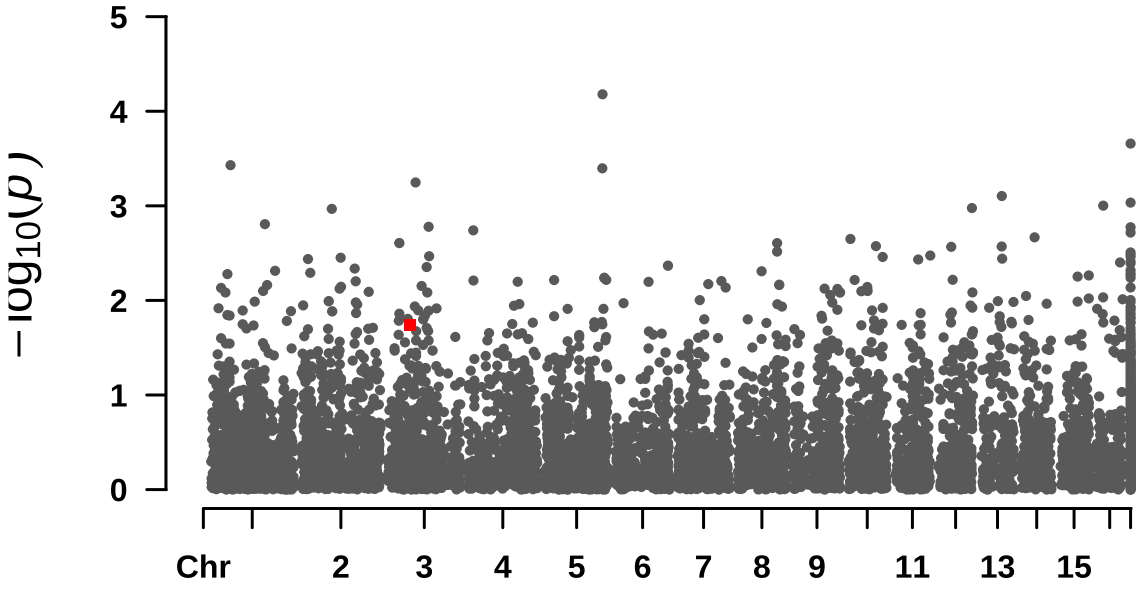


```
##  Rectangular-Manhattan Plotting N_folhas.
```

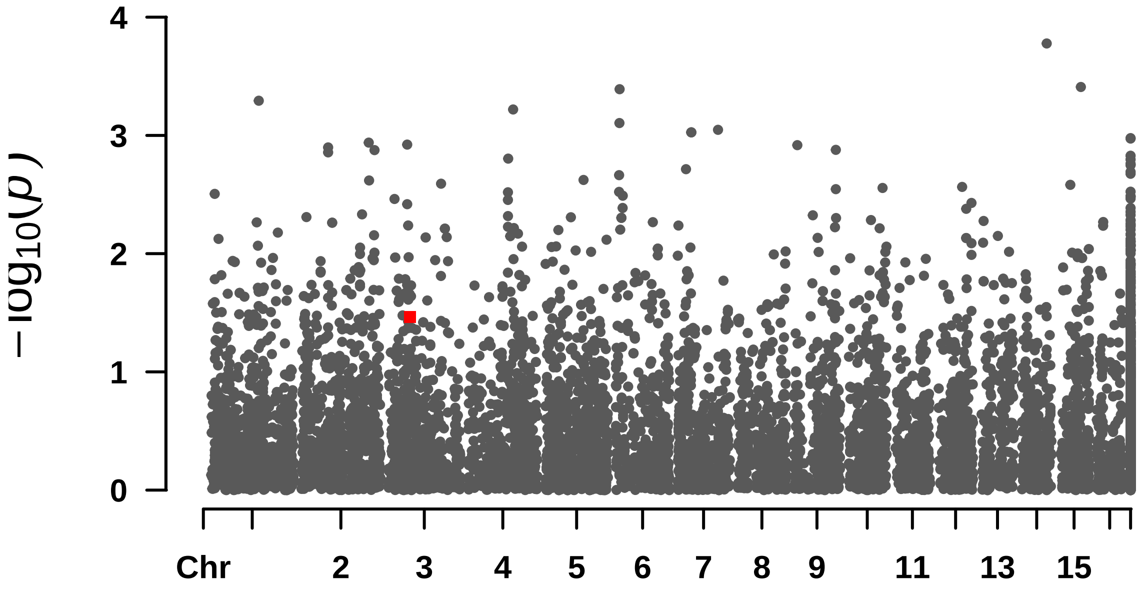


```
##  Rectangular-Manhattan Plotting Oil_content.
```

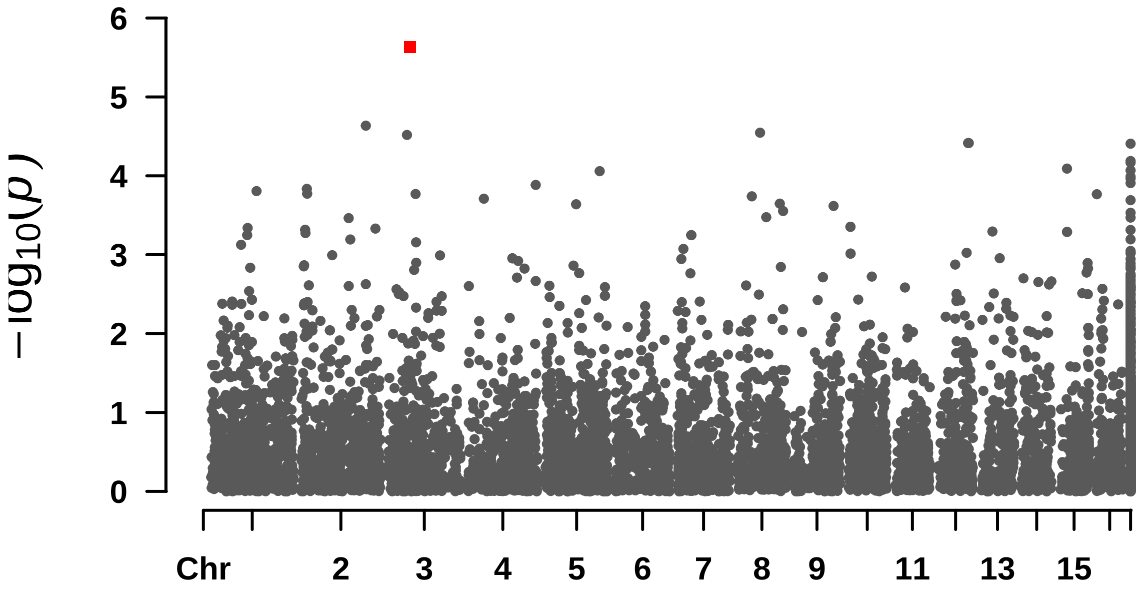


```
##
##
## [1] "=================================================================="
##  Rectangular-Manhattan Plotting Largura_pina.
```

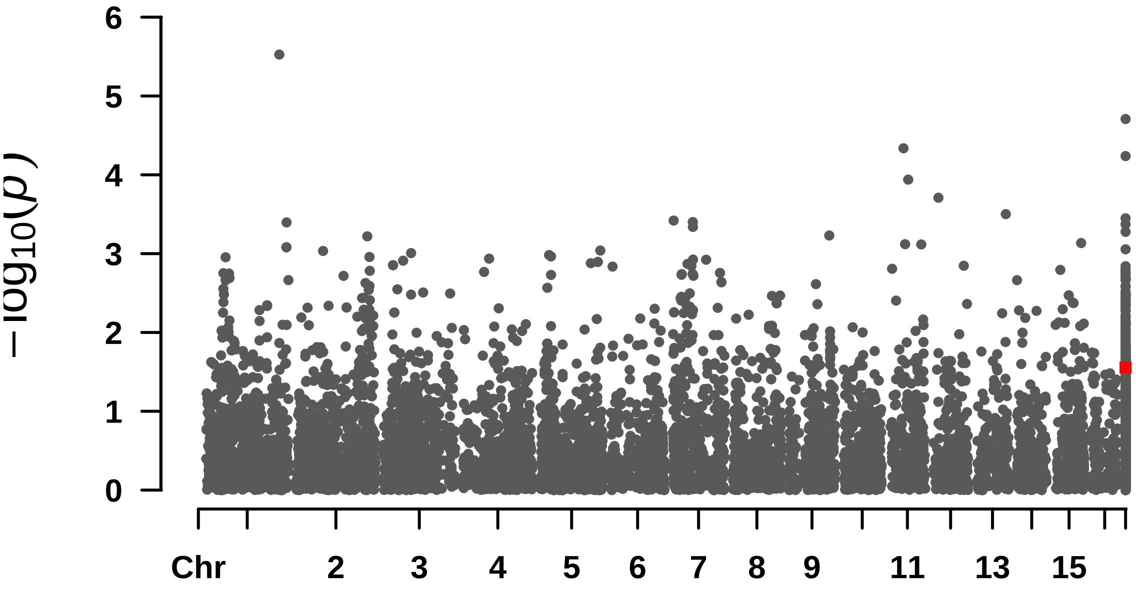


####  Rectangular-Manhattan Plotting F_Casca.

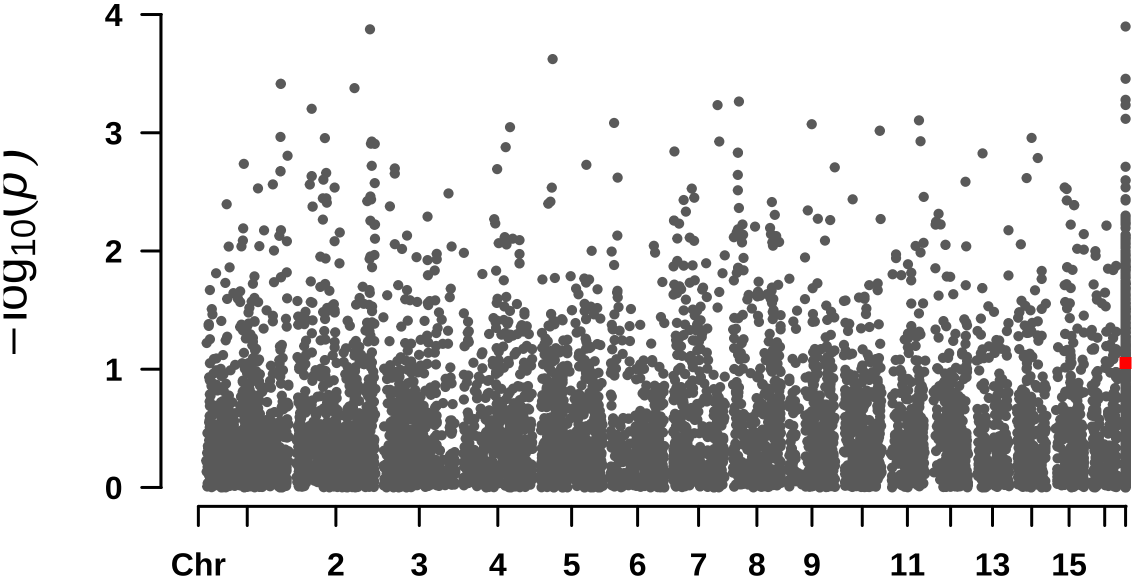


####  Rectangular-Manhattan Plotting S_Casca.

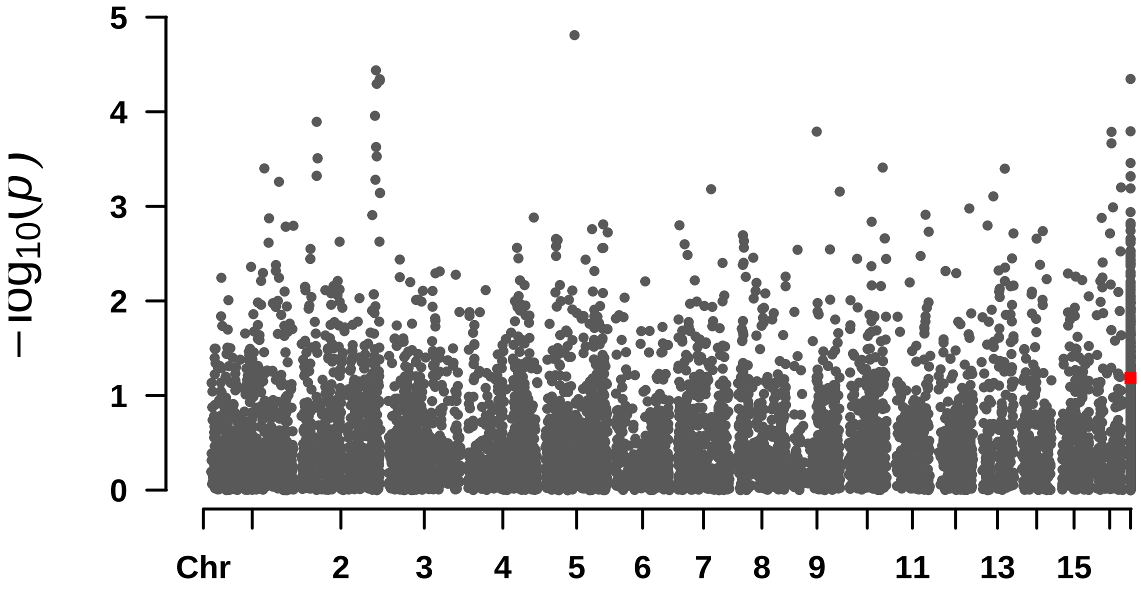


```
##
##
## [1] "=================================================================="
##  Rectangular-Manhattan Plotting N_folhas.
```

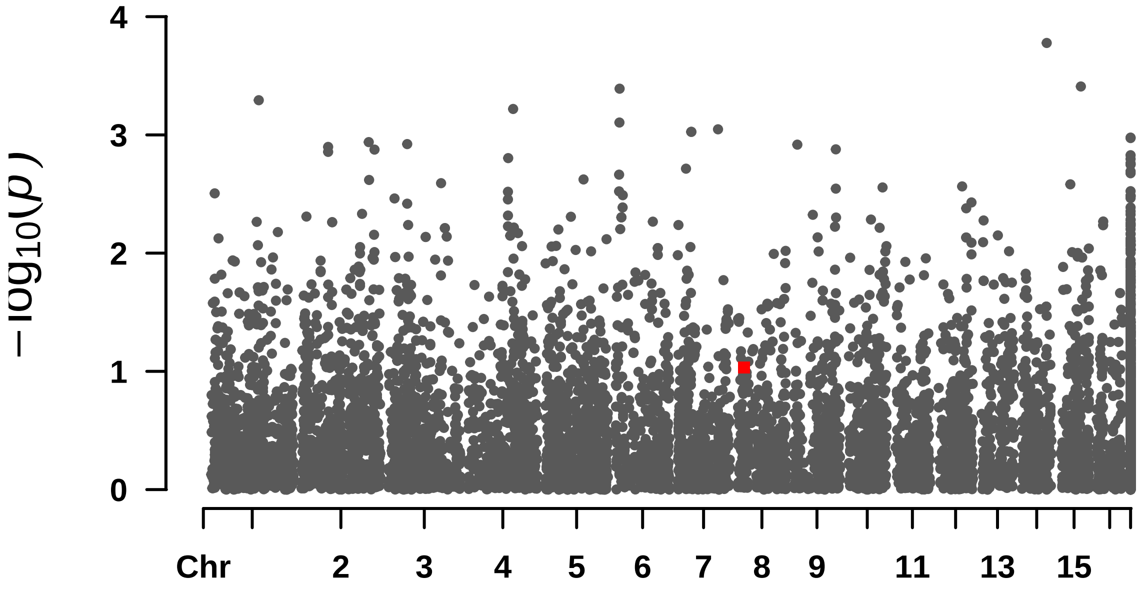


```
##  Rectangular-Manhattan Plotting N_pinas.
```

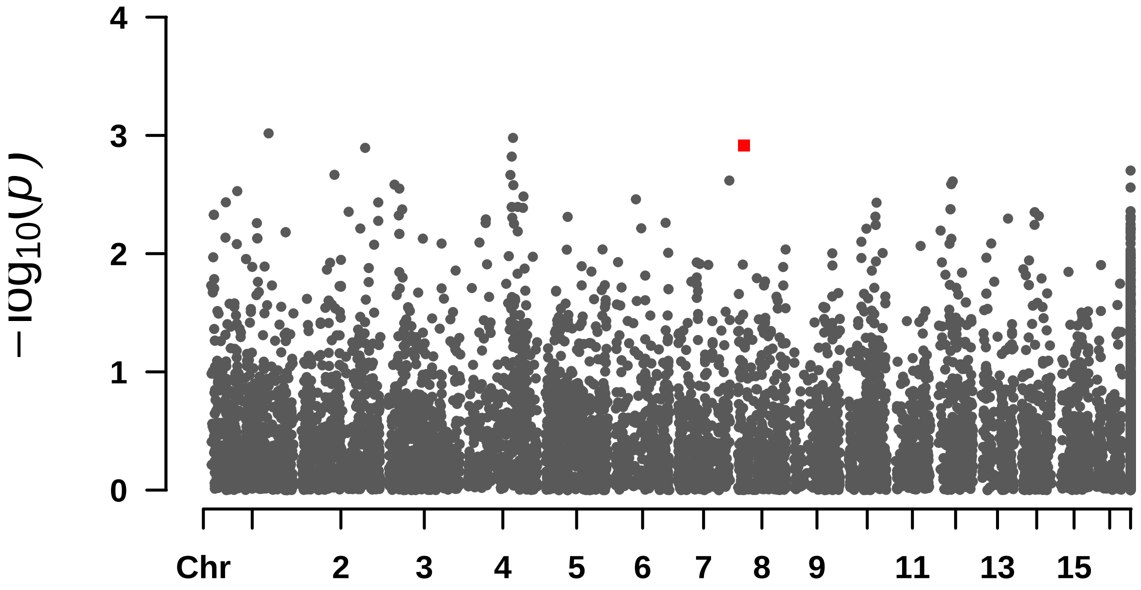


#### Rectangular-Manhattan Plotting F_Casca.

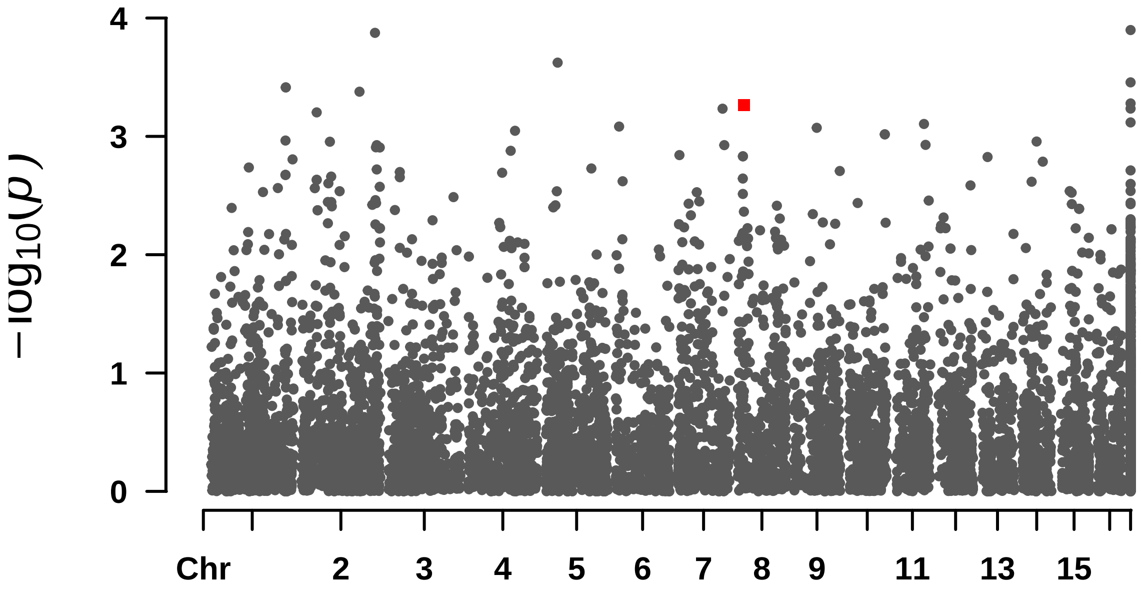


#### Rectangular-Manhattan Plotting F_T.

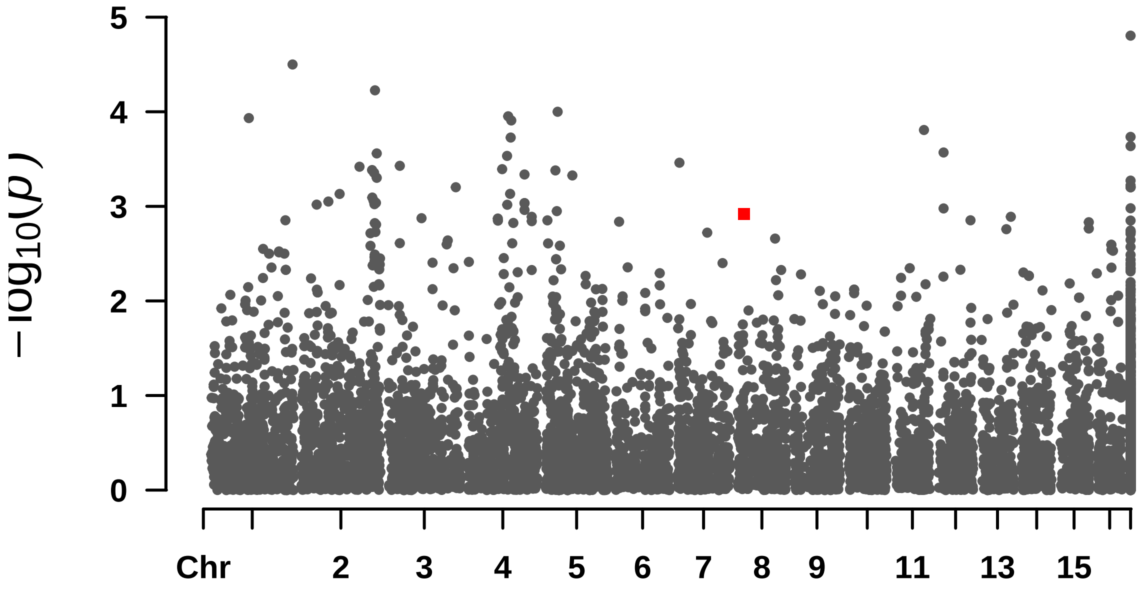


```
##  Rectangular-Manhattan Plotting S_Casca.
```

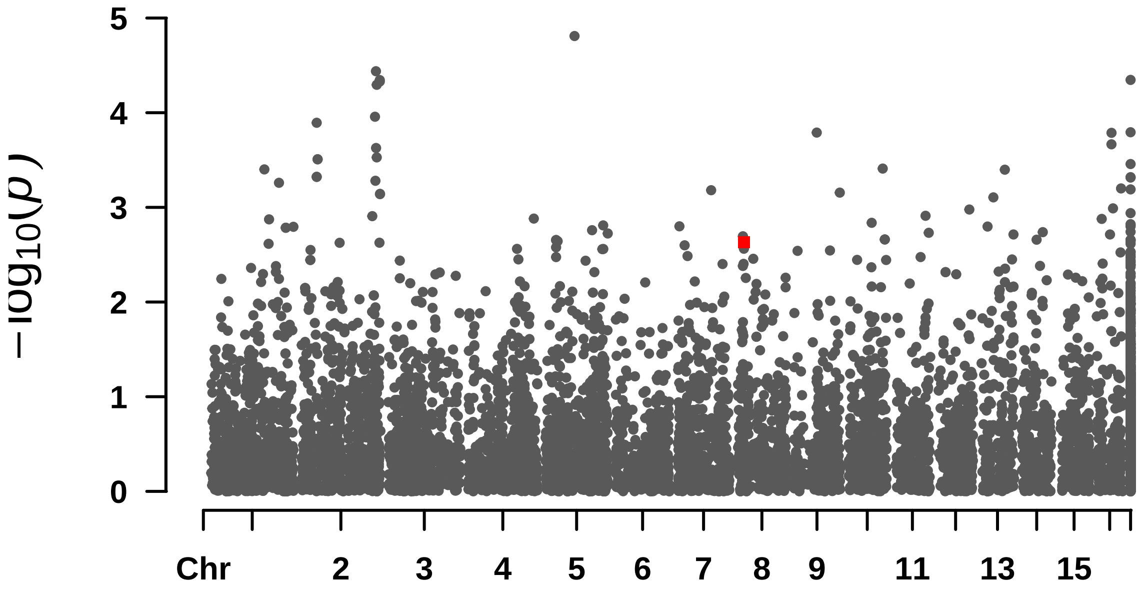


```
##
##
## [1] "====================================================================="
```

---

#### ## GLM Denovo

```
##  Rectangular-Manhattan Plotting S_Casca.
```

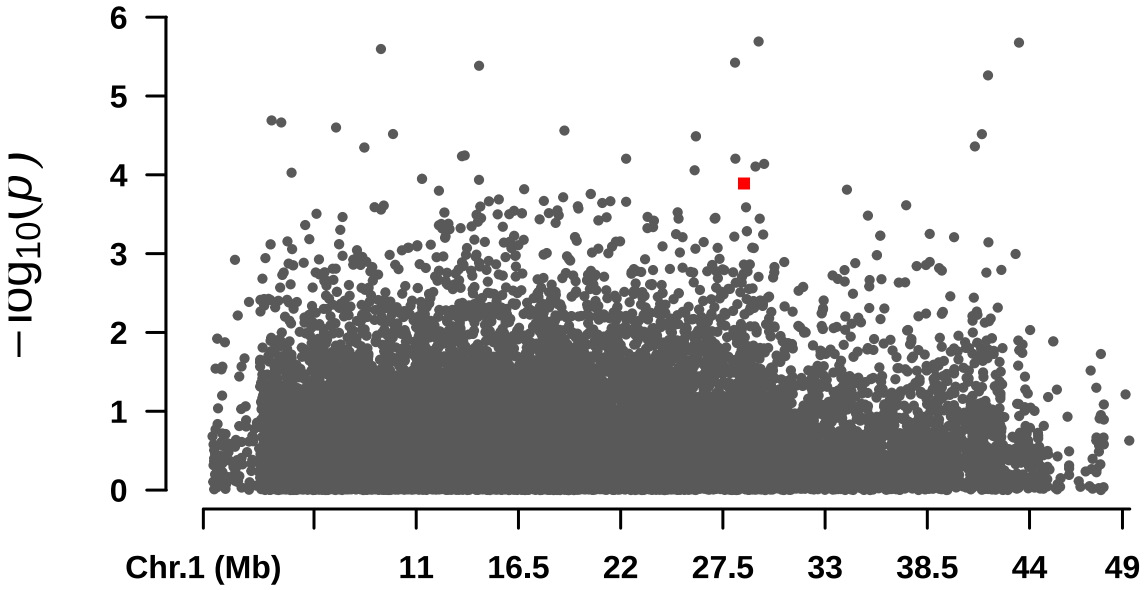


```
##  Rectangular-Manhattan Plotting S_T.
```

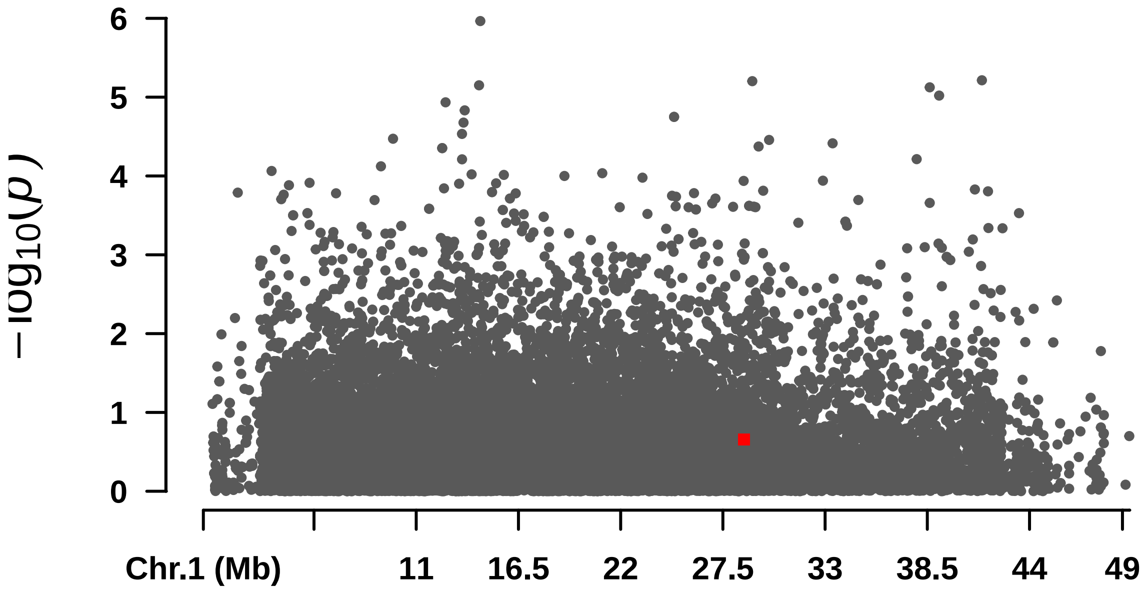


```
##  Rectangular-Manhattan Plotting S_Polpa.
```

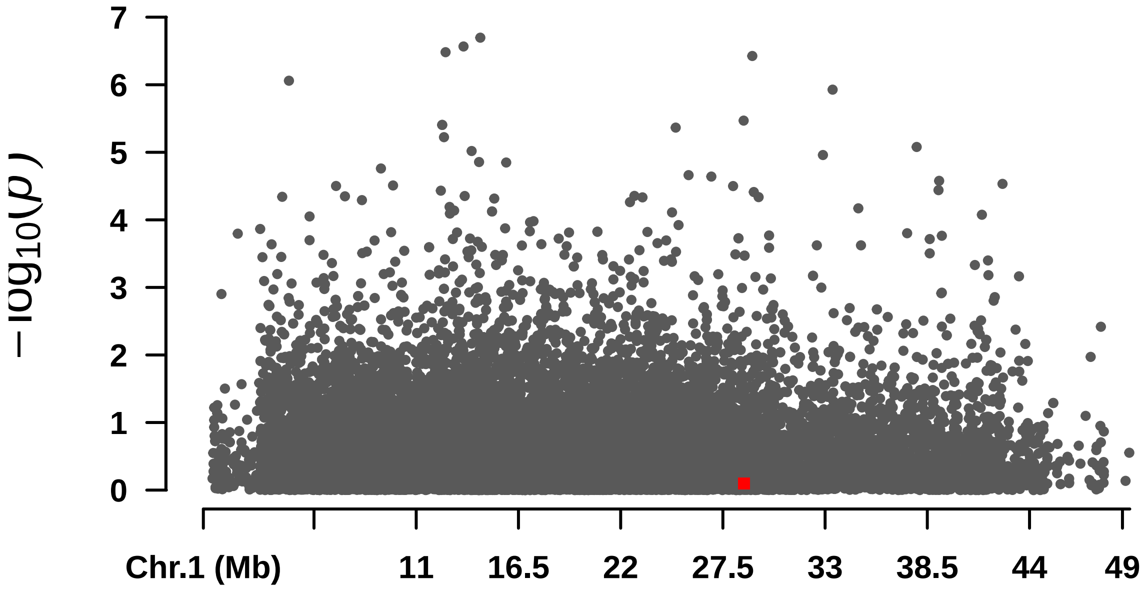


```
##
##
## [1] "=================================================================="
##  Rectangular-Manhattan Plotting S_Casca.
```

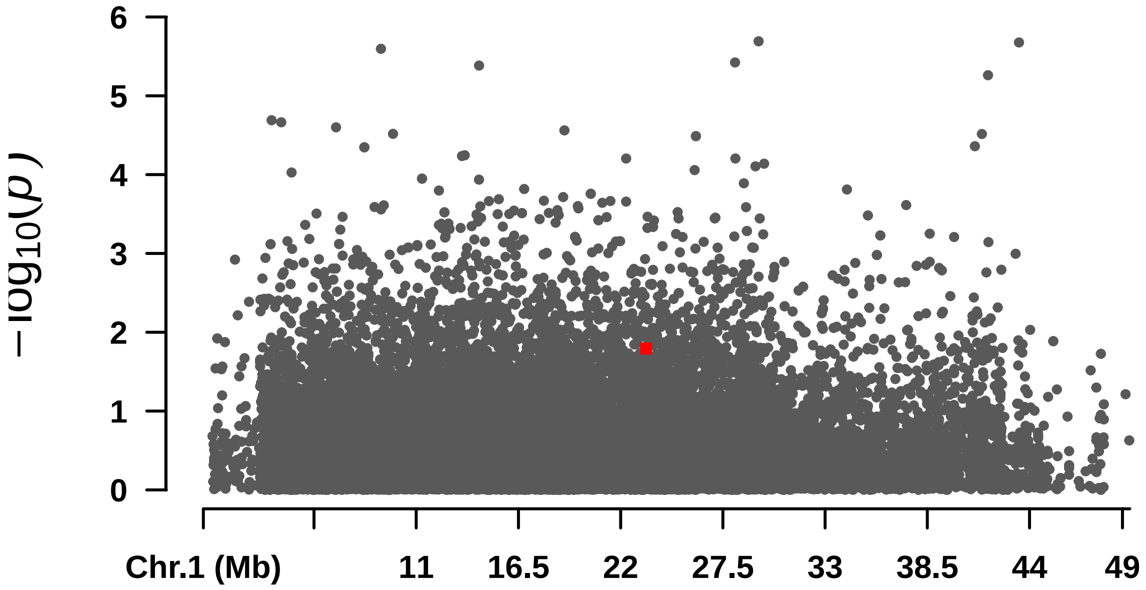


```
##  Rectangular-Manhattan Plotting S_T.
```

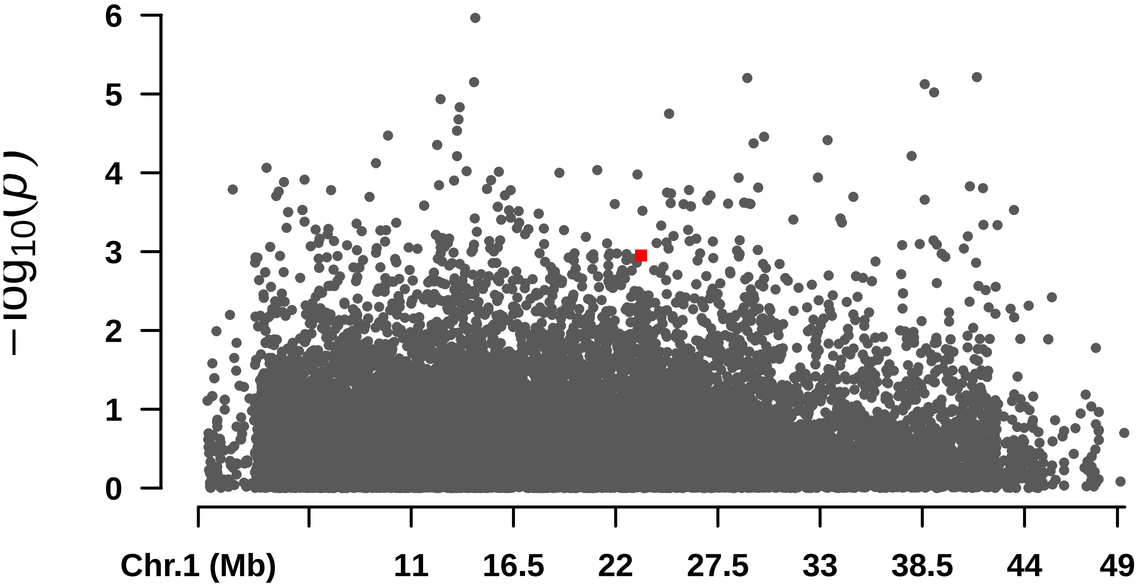


#### Rectangular-Manhattan Plotting S_Polpa.

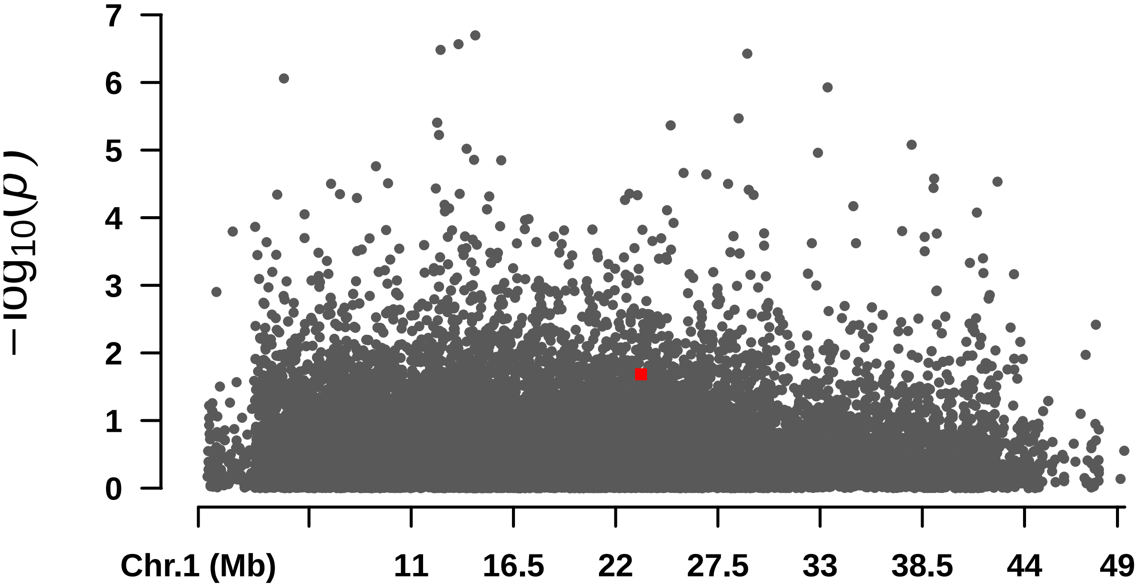


#### Rectangular-Manhattan Plotting Oil_content.

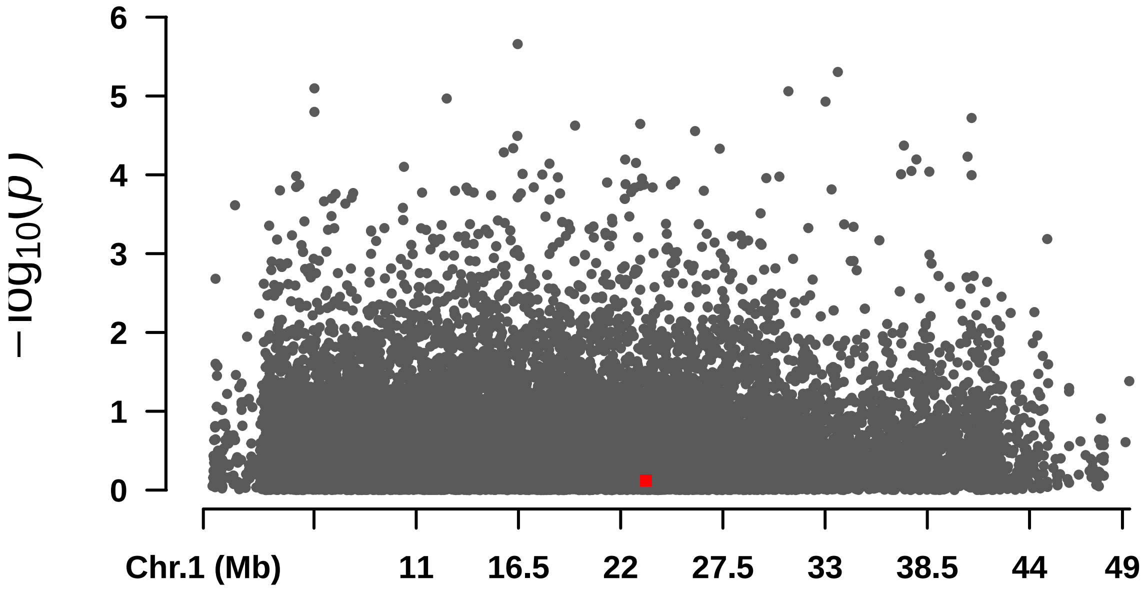


```
##
##
## [1] "=================================================================="
##  Rectangular-Manhattan Plotting S_T.
```

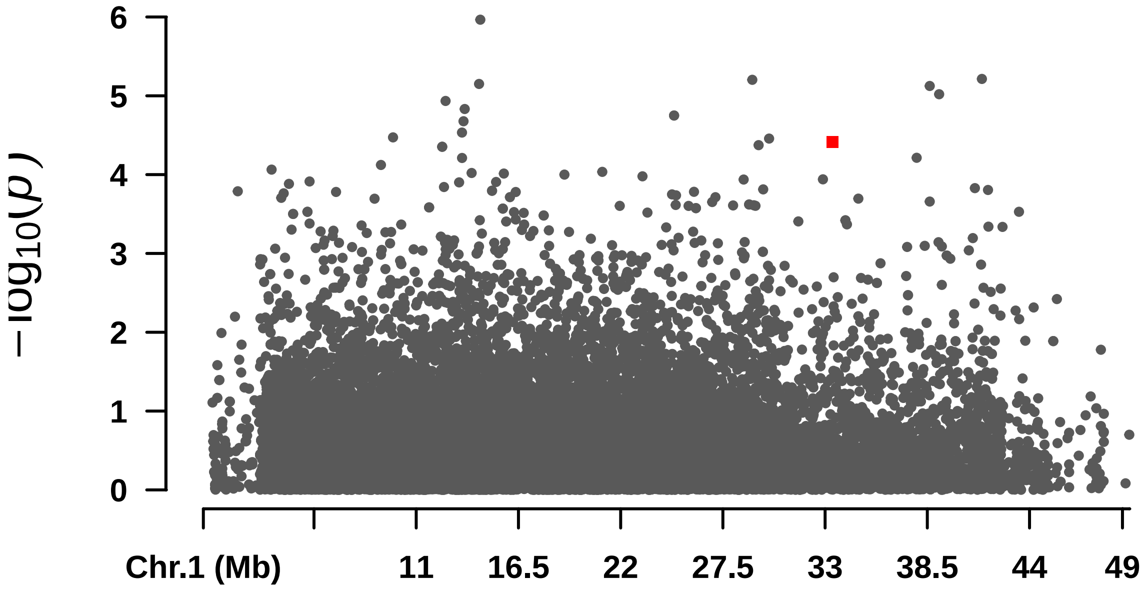


```
##  Rectangular-Manhattan Plotting S_Sem.
```

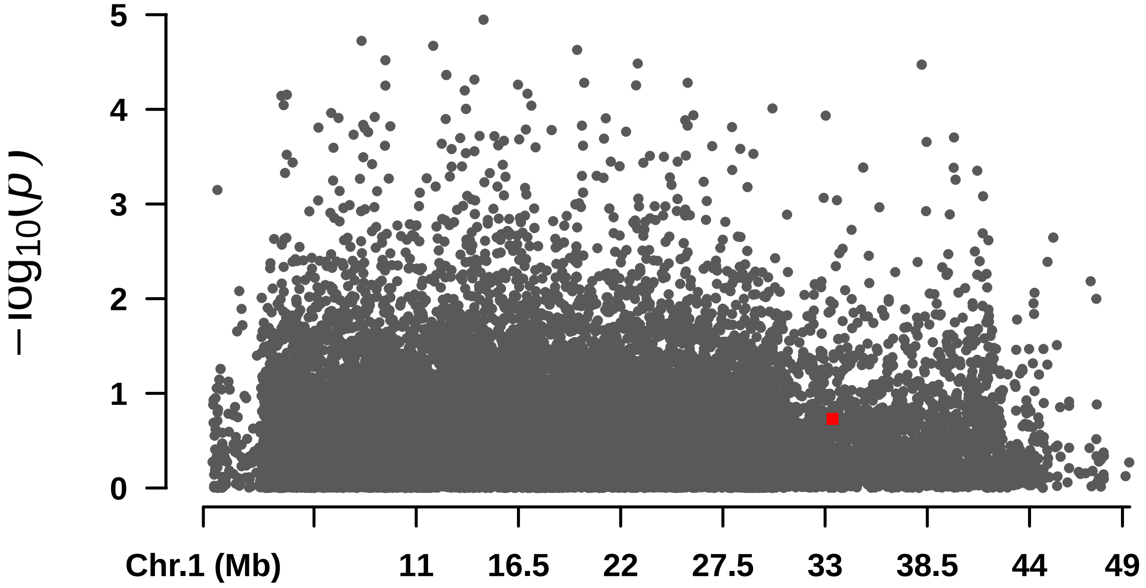


```
##
##
## [1] "======================================================================"
```

---

#### ## GLM trans

```
##  Rectangular-Manhattan Plotting S_T.
```

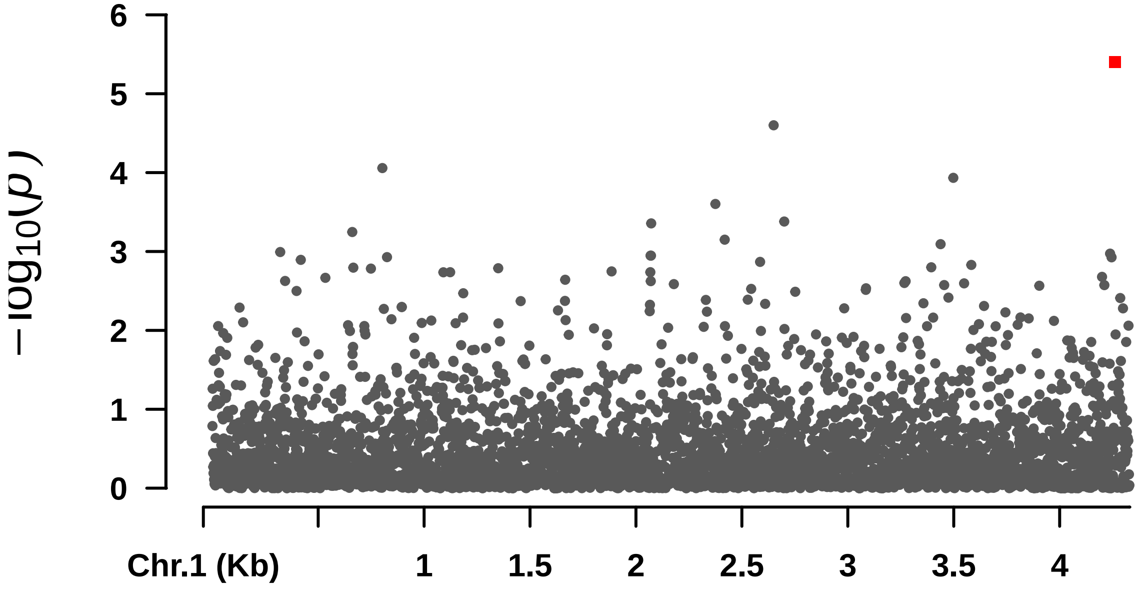


```
##  Rectangular-Manhattan Plotting Oil_content.
```

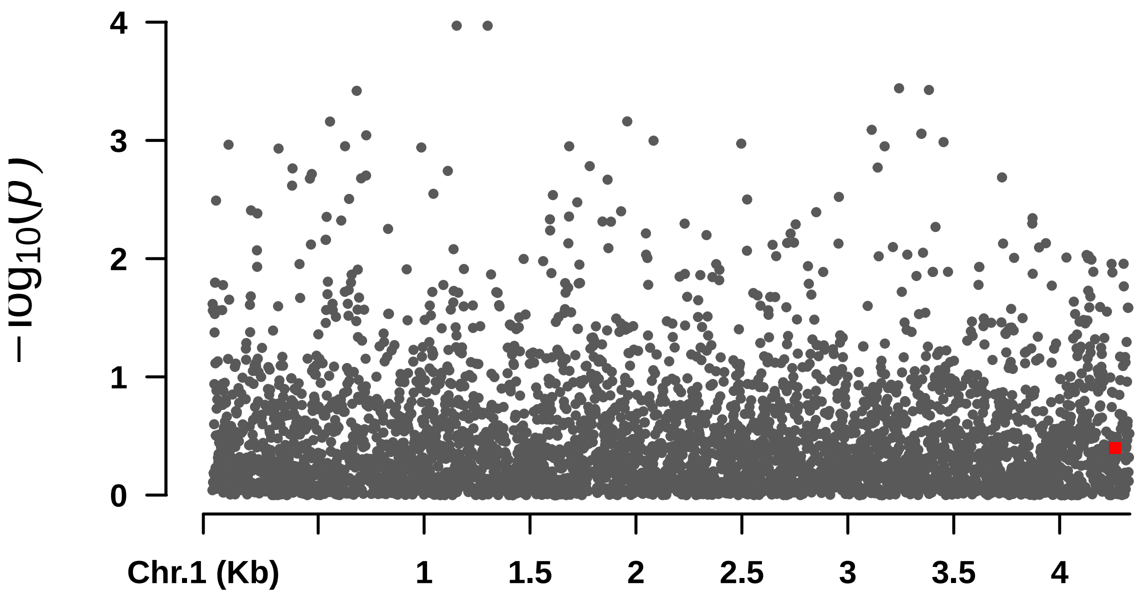


```
##  Rectangular-Manhattan Plotting S_Casca.
```

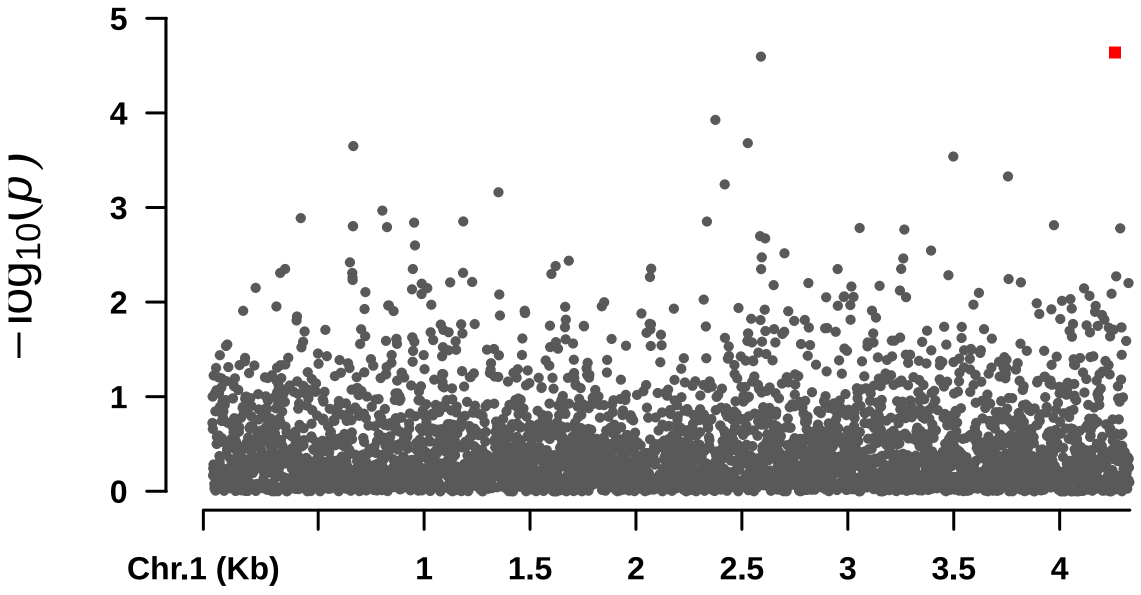


```
##
##
## [1] "==================================================================="
```

#Single trait GWAS figures - DENDE

#### Altura

#### numeric(0)

####  Rectangular-Manhattan Plotting BLINK.

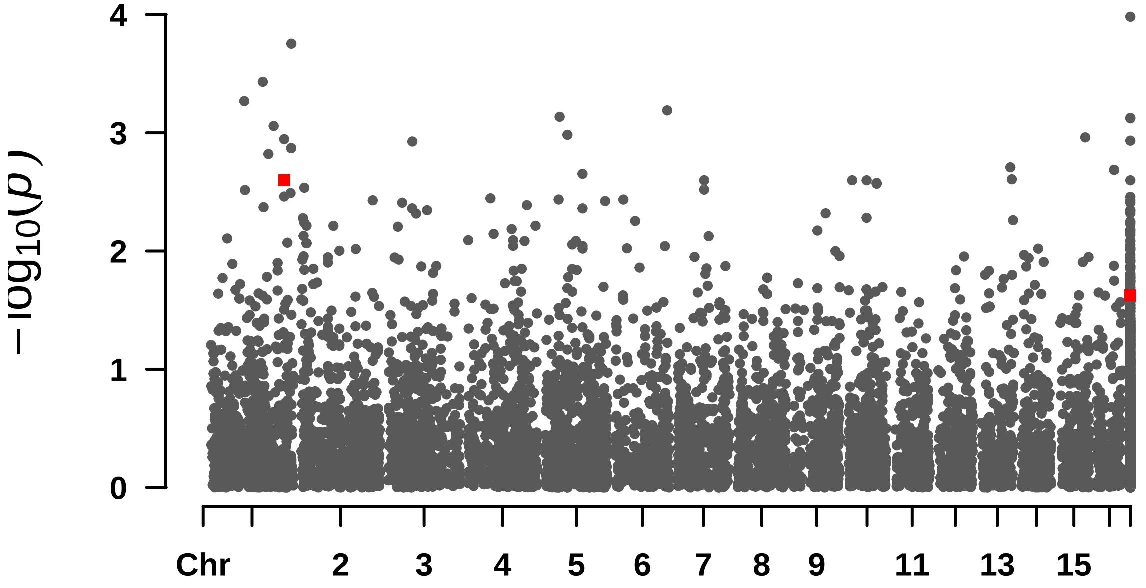


####  Rectangular-Manhattan Plotting FarmCPU.

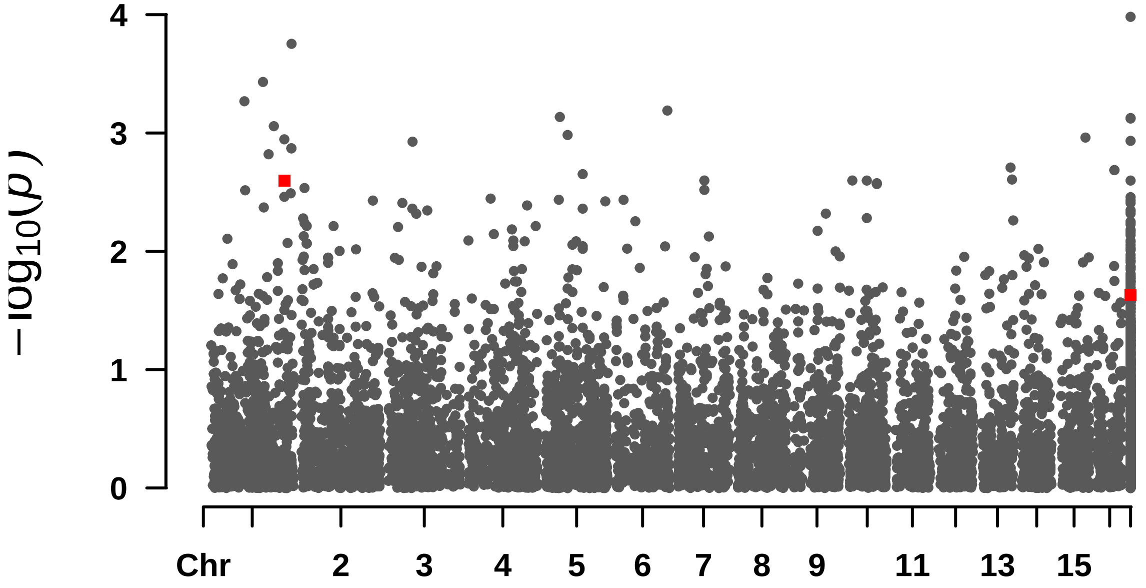


####  Rectangular-Manhattan Plotting GLM.

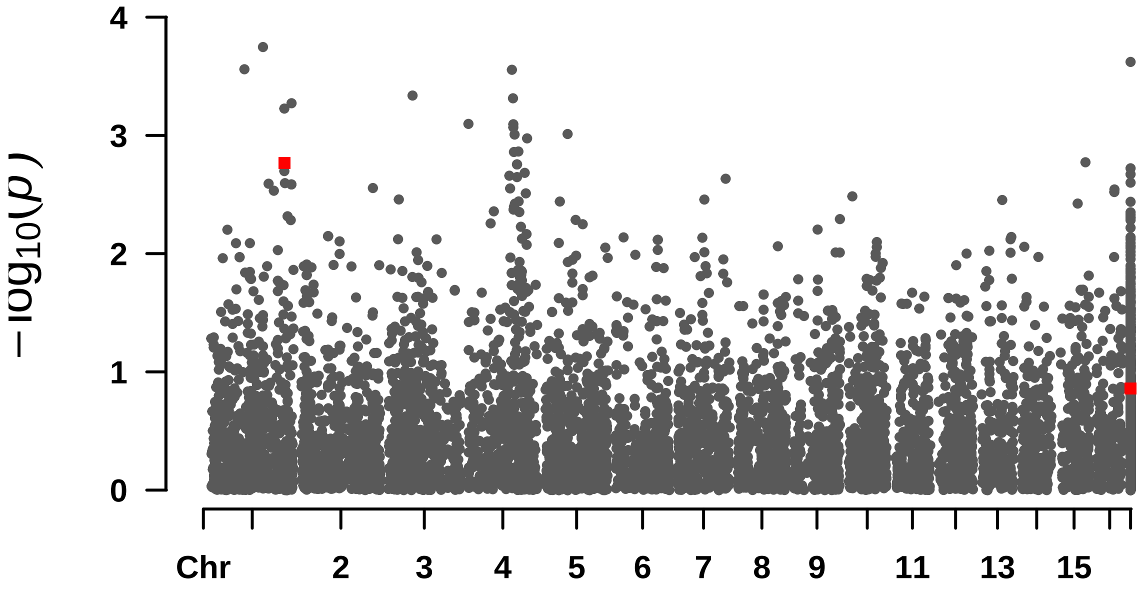


#### Rectangular-Manhattan Plotting MLM.

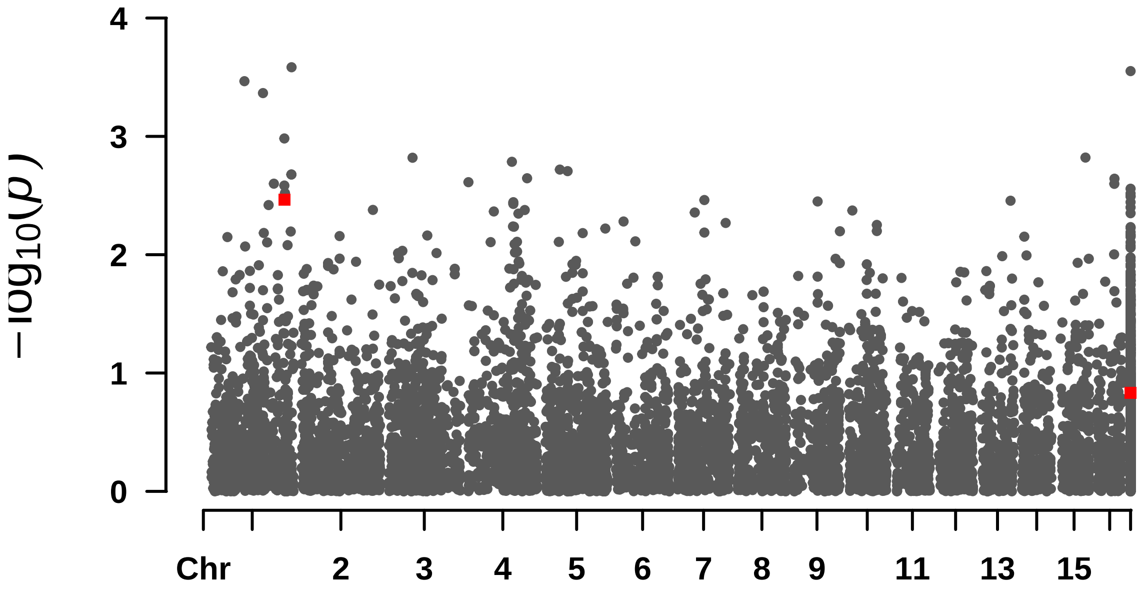


#### Rectangular-Manhattan Plotting MLMM.

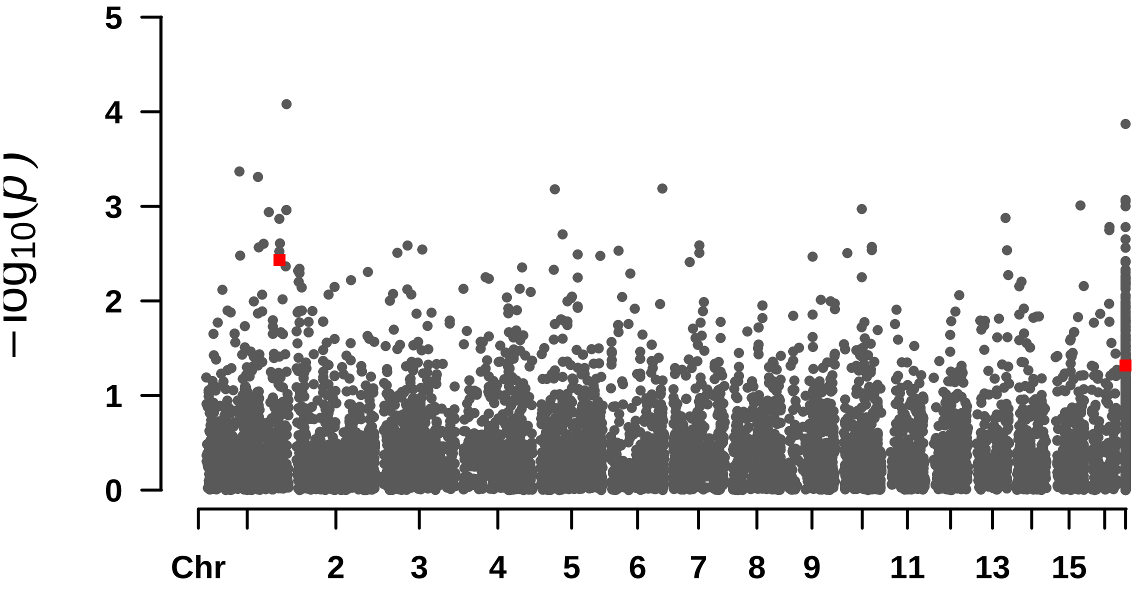

$-\log_{10}(p)$
5
4
3
2
1
0
Chr
2
3
4
5
6
7
8
9
11
13
15

#### Compri_folha

```
## numeric(0)
```

```
##  Rectangular-Manhattan Plotting BLINK.
```

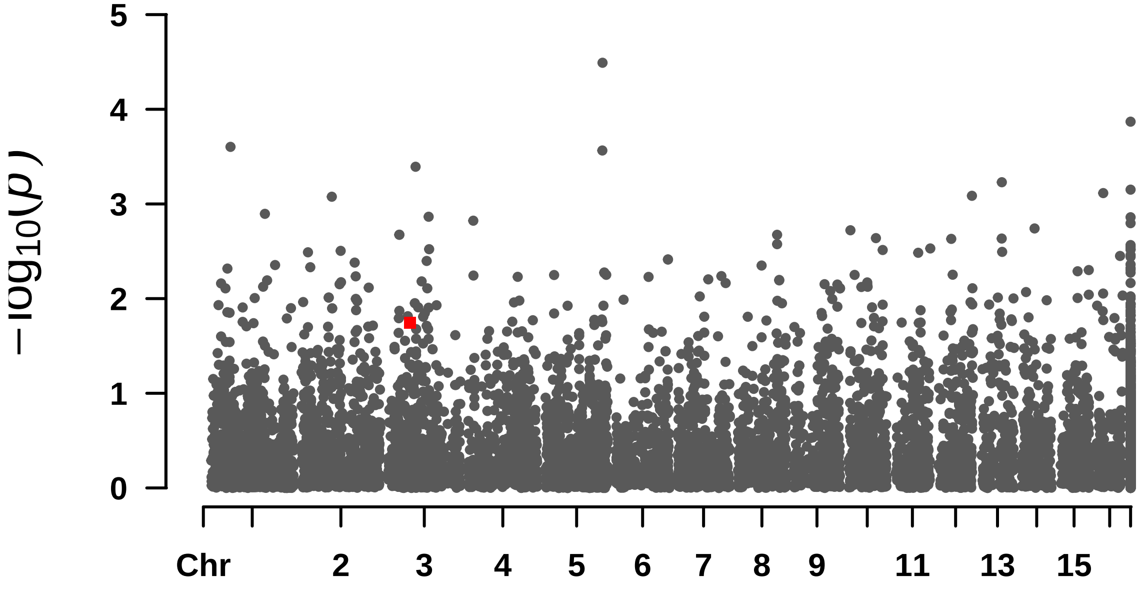


```
##  Rectangular-Manhattan Plotting FarmCPU.
```

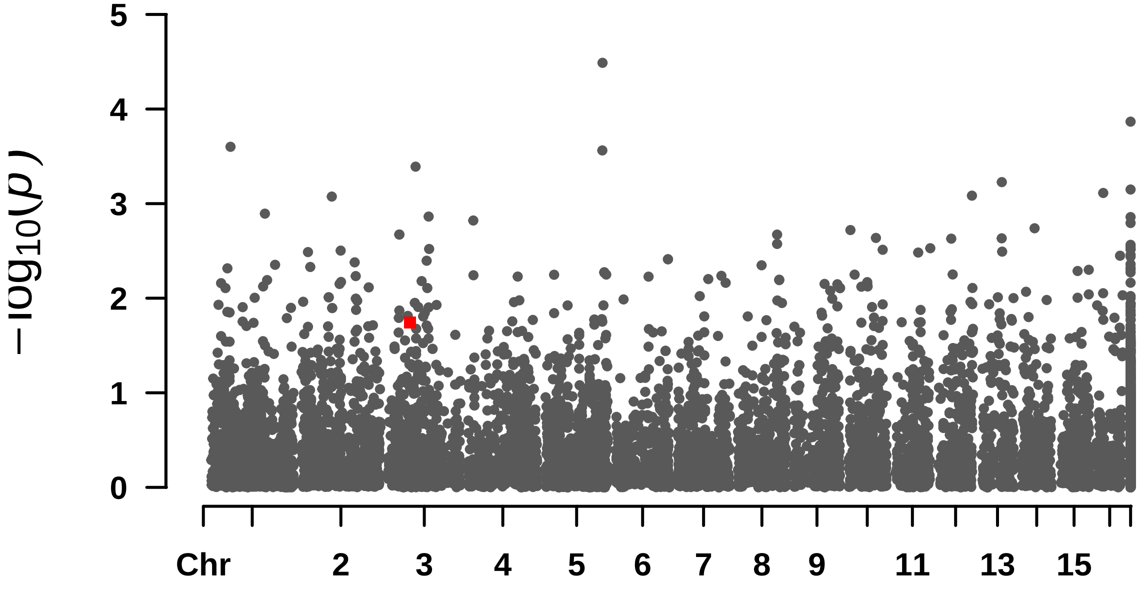


```
##  Rectangular-Manhattan Plotting GLM.
```

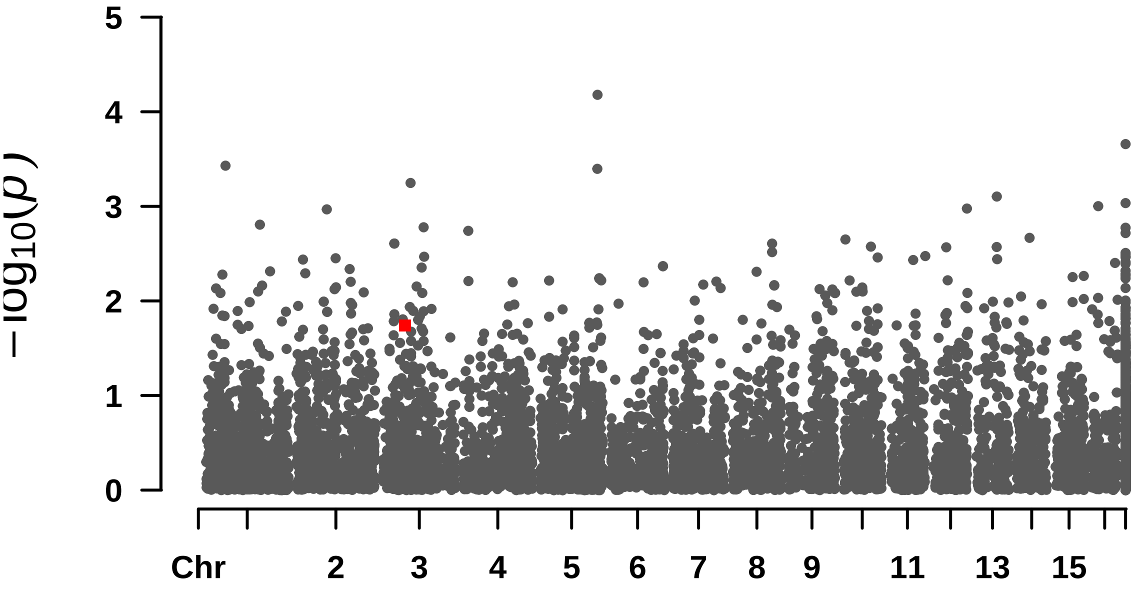


#### Rectangular-Manhattan Plotting MLM.

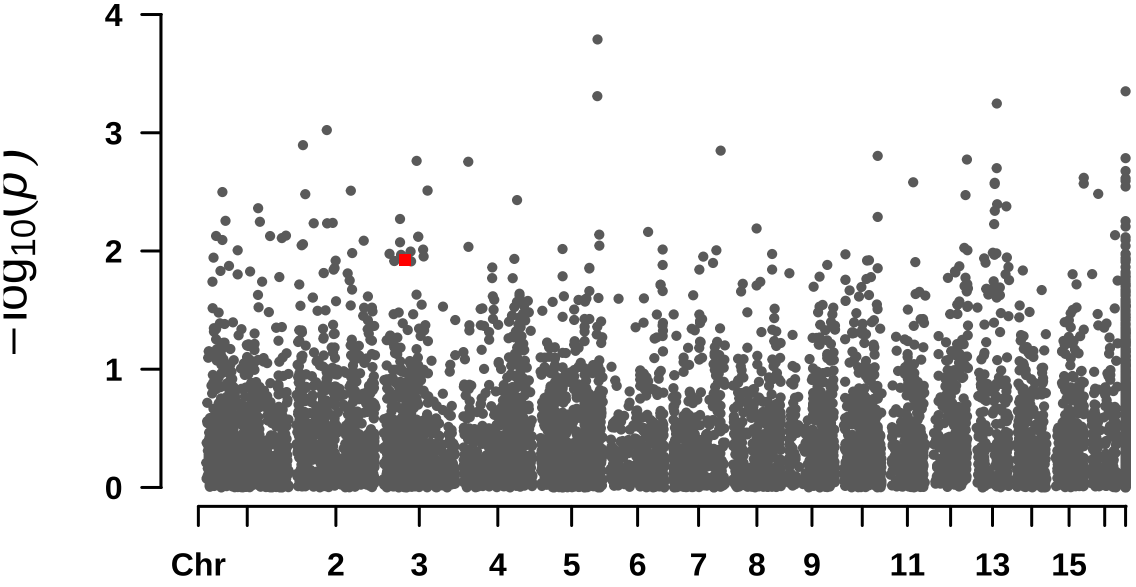


#### Rectangular-Manhattan Plotting MLMM.

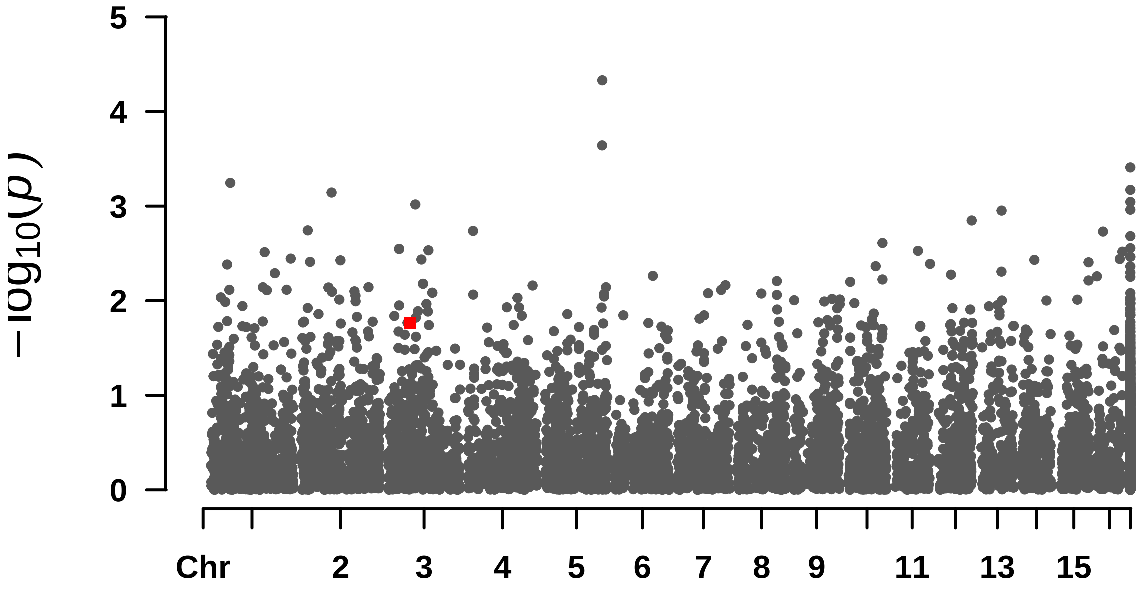

$-\log_{10}(p)$
5
4
3
2
1
0
Chr
2
3
4
5
6
7
8
9
11
13
15

#### DAP

```
## numeric(0)
```

```
##  Rectangular-Manhattan Plotting BLINK.
```

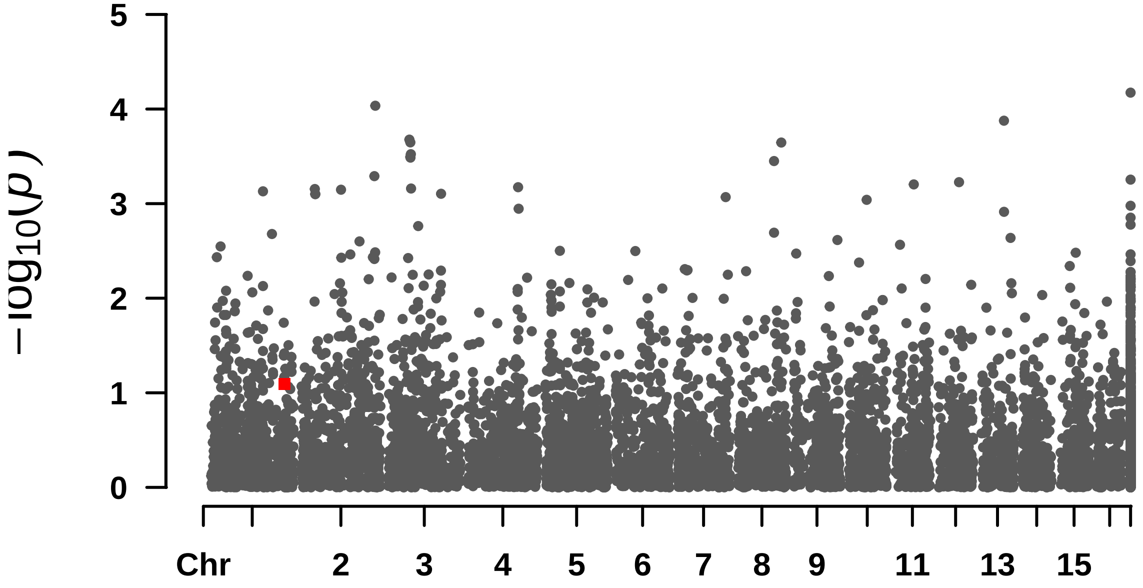


```
##  Rectangular-Manhattan Plotting FarmCPU.
```

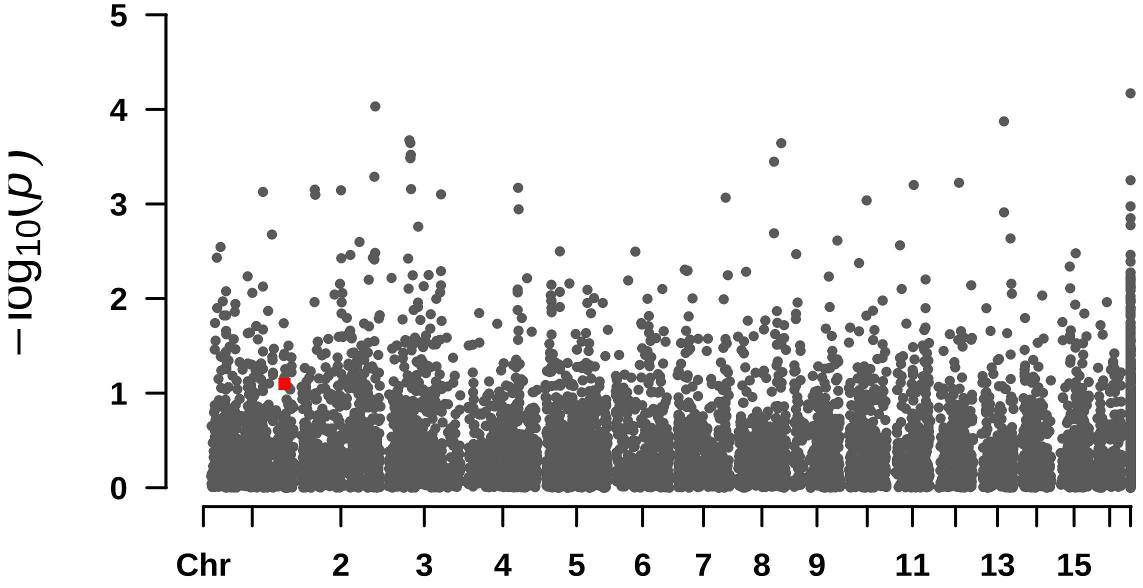


```
##  Rectangular-Manhattan Plotting GLM.
```

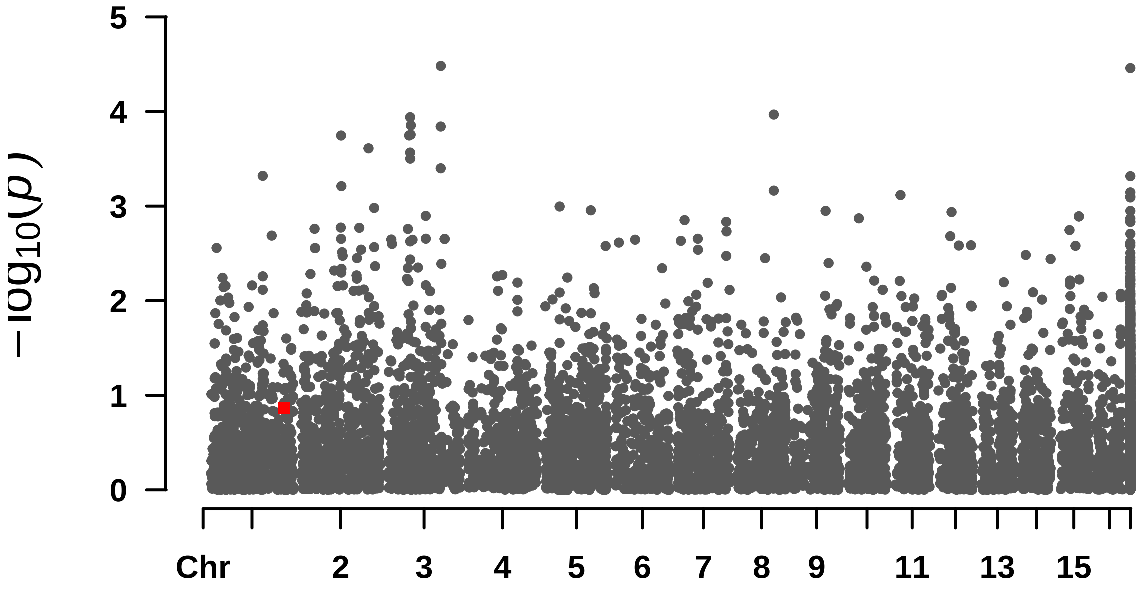


```
##  Rectangular-Manhattan Plotting MLM.
```

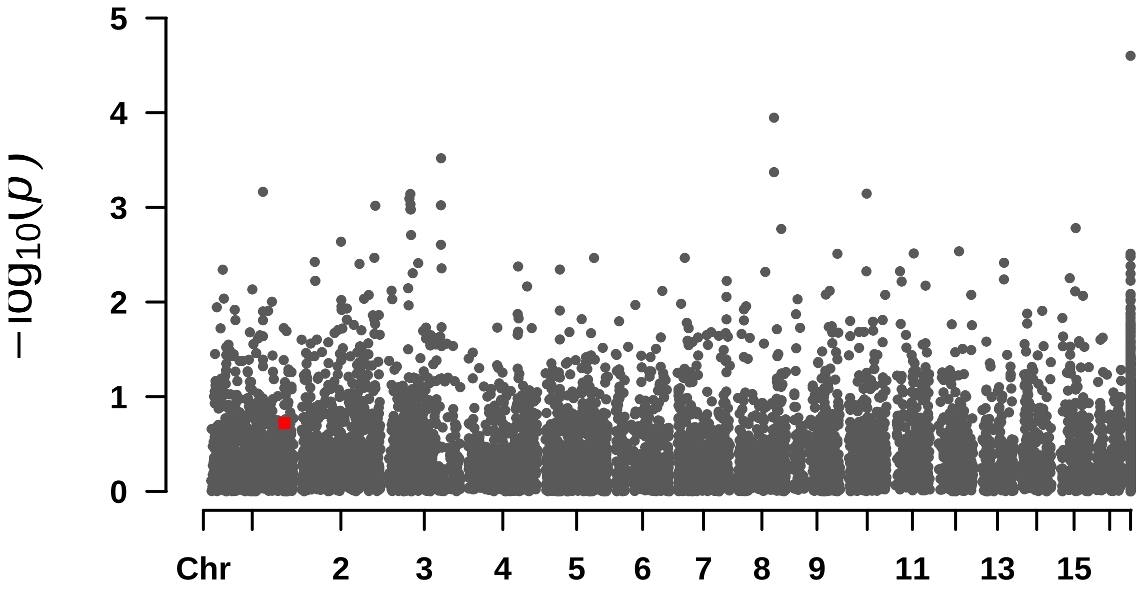


```
##  Rectangular-Manhattan Plotting MLMM.
```

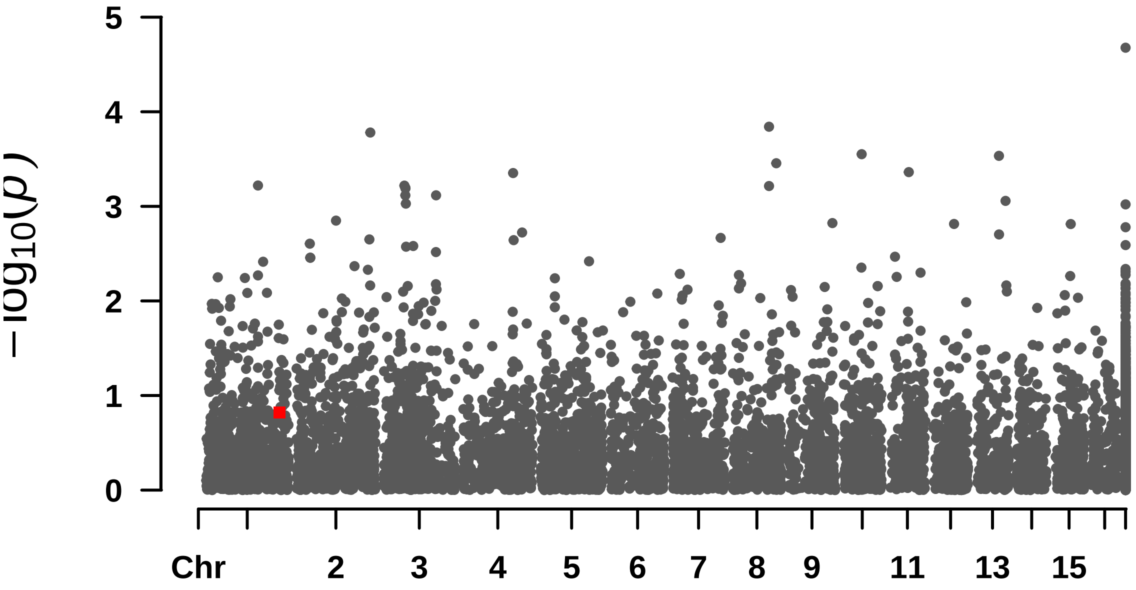
5
4
3
2
1
0
−log10(p)
Chr
2
3
4
5
6
7
8
9
11
13
15

—

#### F_Casca

#### numeric(0)

####  Rectangular-Manhattan Plotting BLINK.

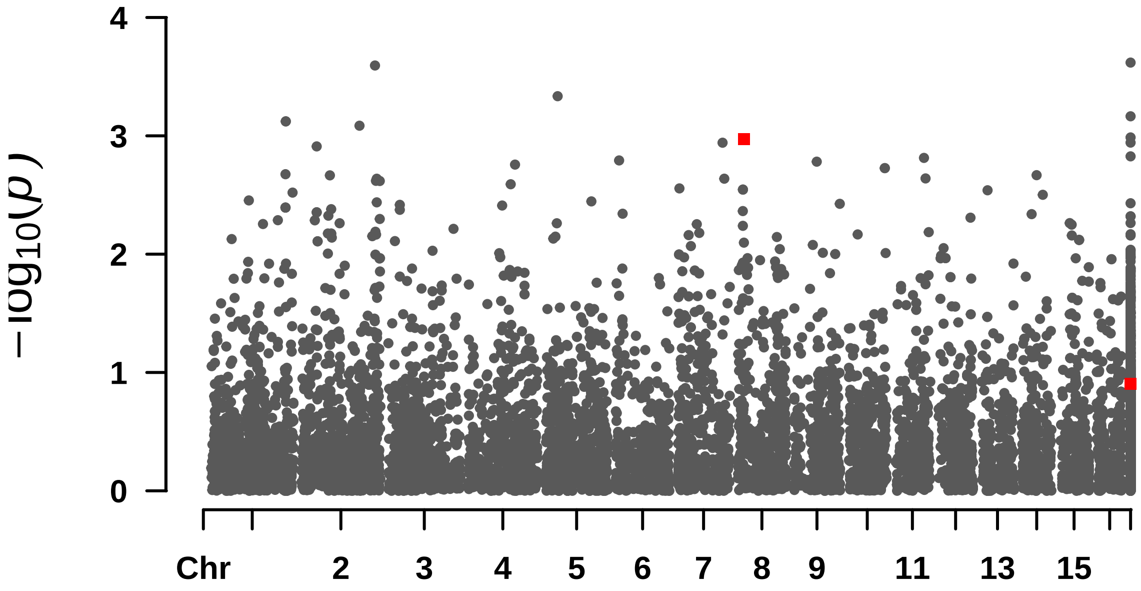


####  Rectangular-Manhattan Plotting FarmCPU.

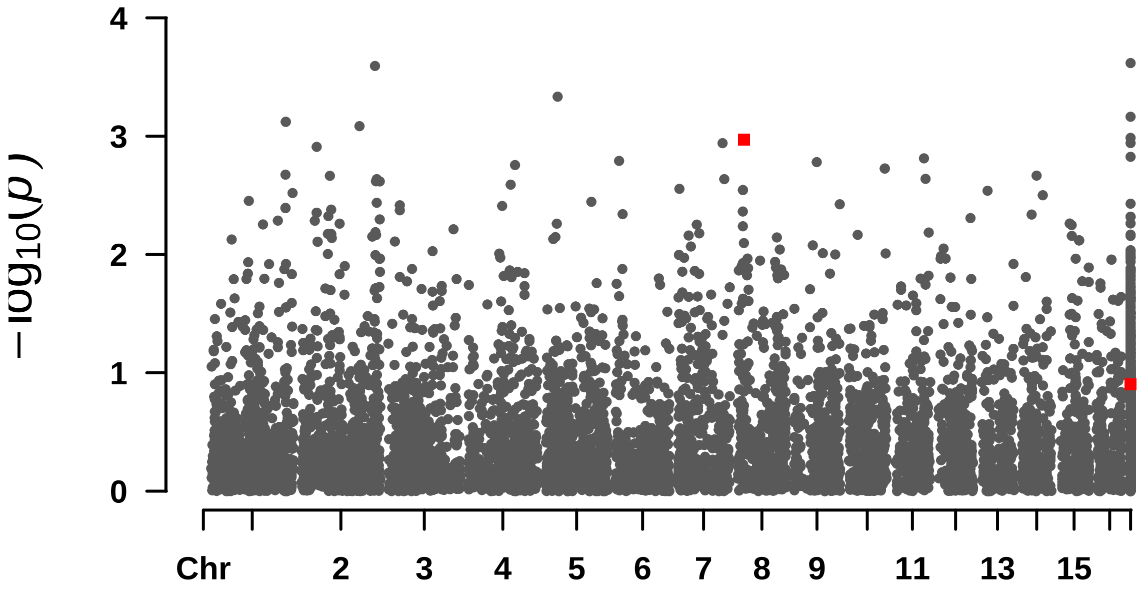


####  Rectangular-Manhattan Plotting GLM.

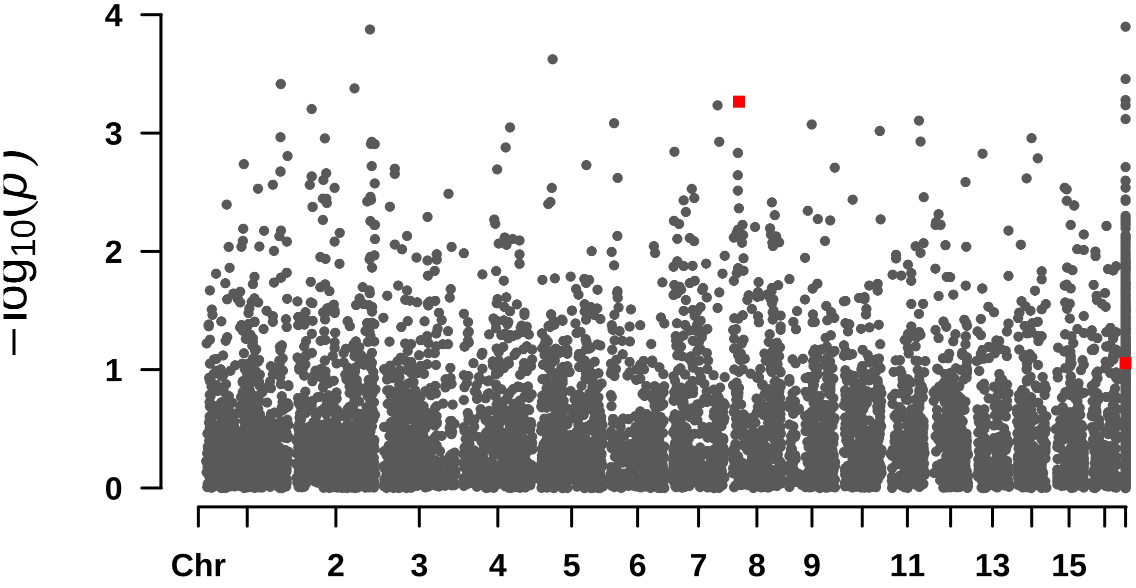


#### Rectangular-Manhattan Plotting MLM.

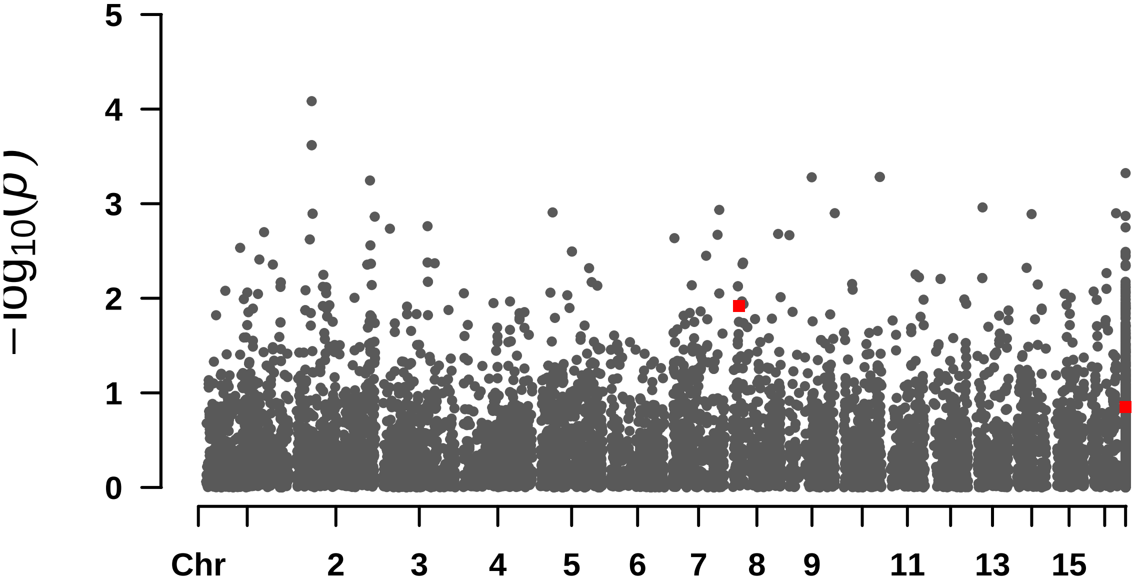


#### Rectangular-Manhattan Plotting MLMM.

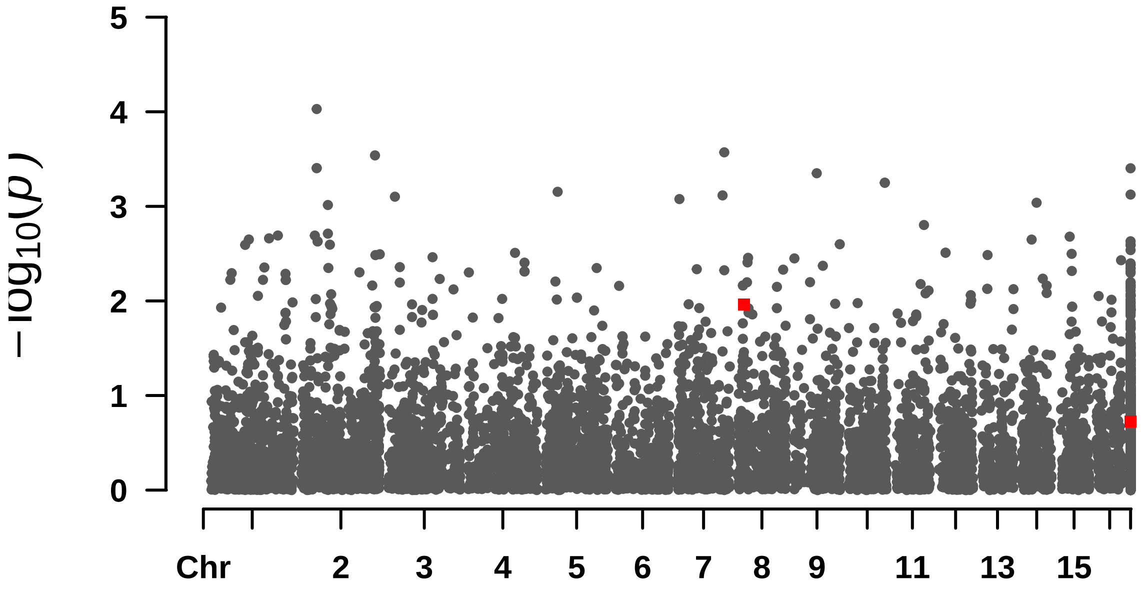

5
4
3
2
1
0
$-\log_{10}(p)$
Chr
2
3
4
5
6
7
8
9
11
13
15

—

#### F_Endo

```
## numeric(0)
```

```
##  Rectangular-Manhattan Plotting BLINK.
```

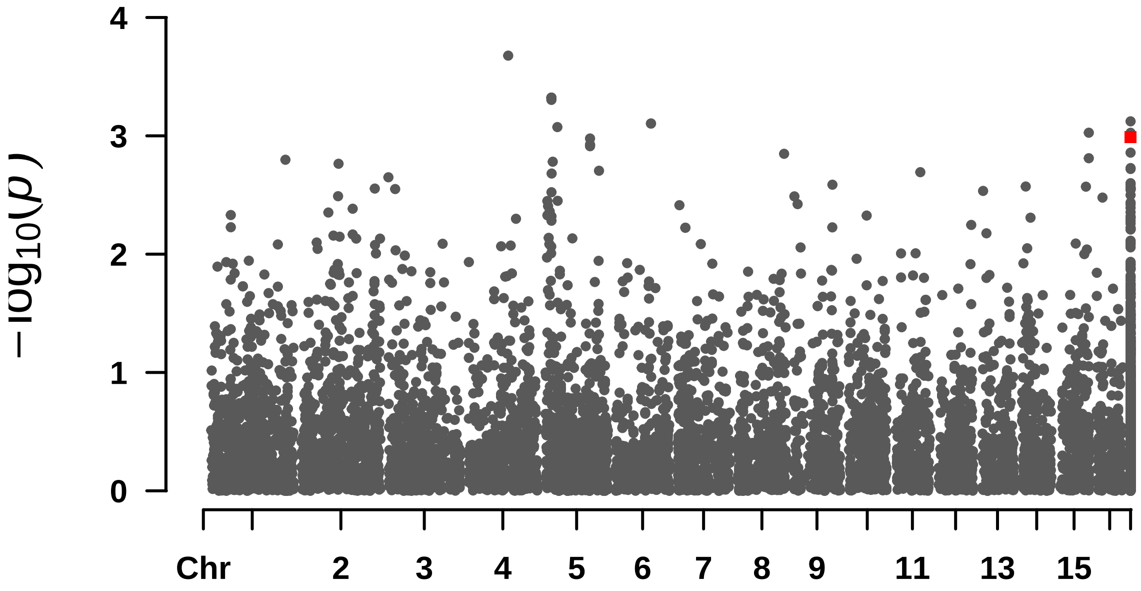


```
##  Rectangular-Manhattan Plotting FarmCPU.
```

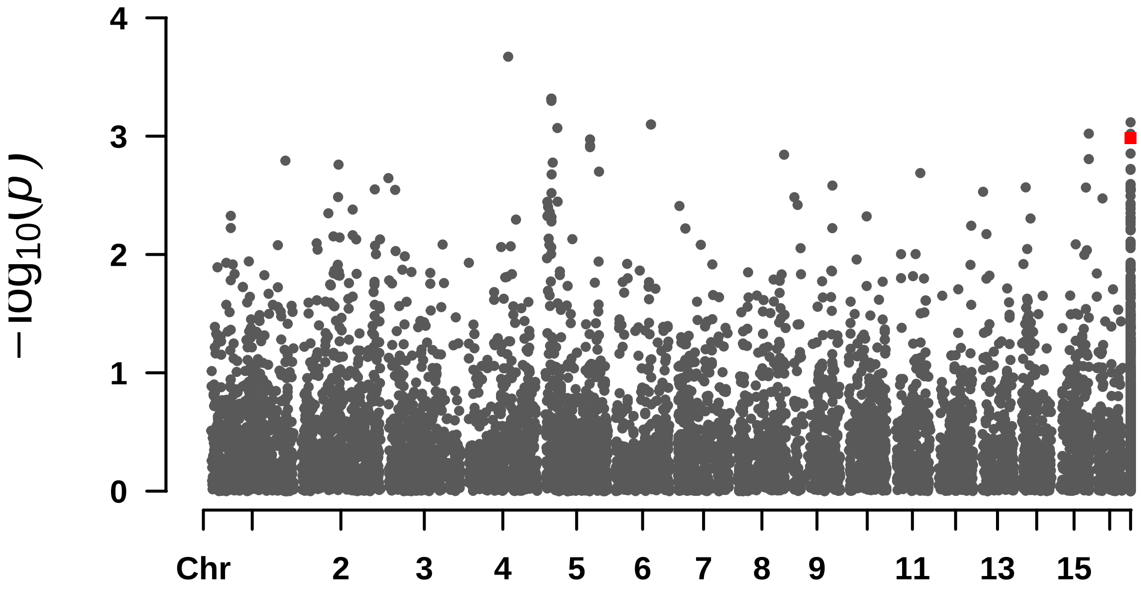


```
##  Rectangular-Manhattan Plotting GLM.
```

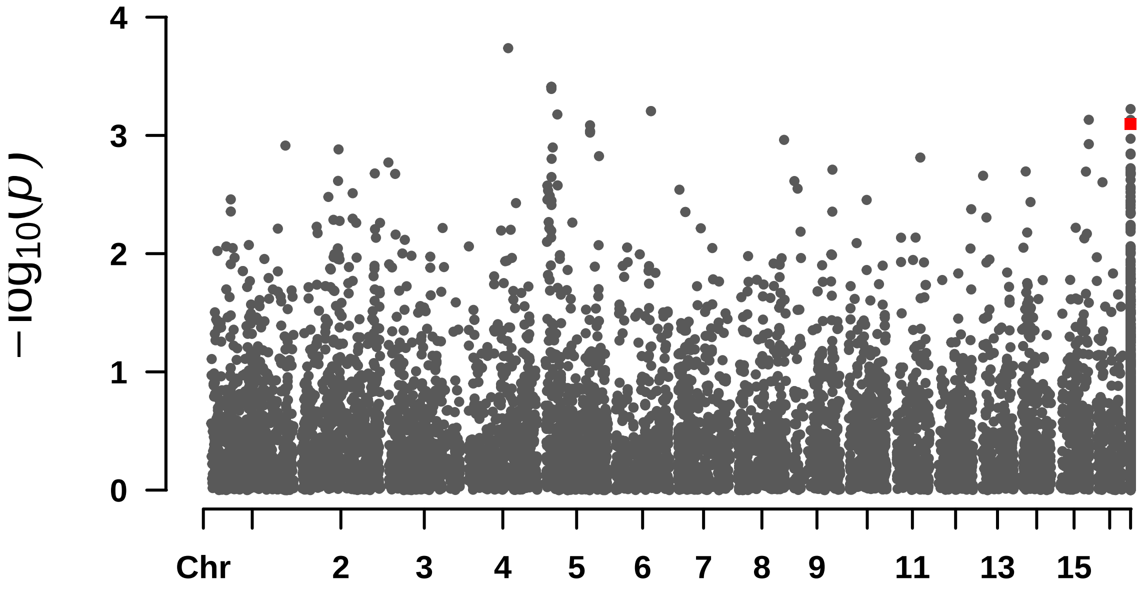


```
##  Rectangular-Manhattan Plotting MLM.
```

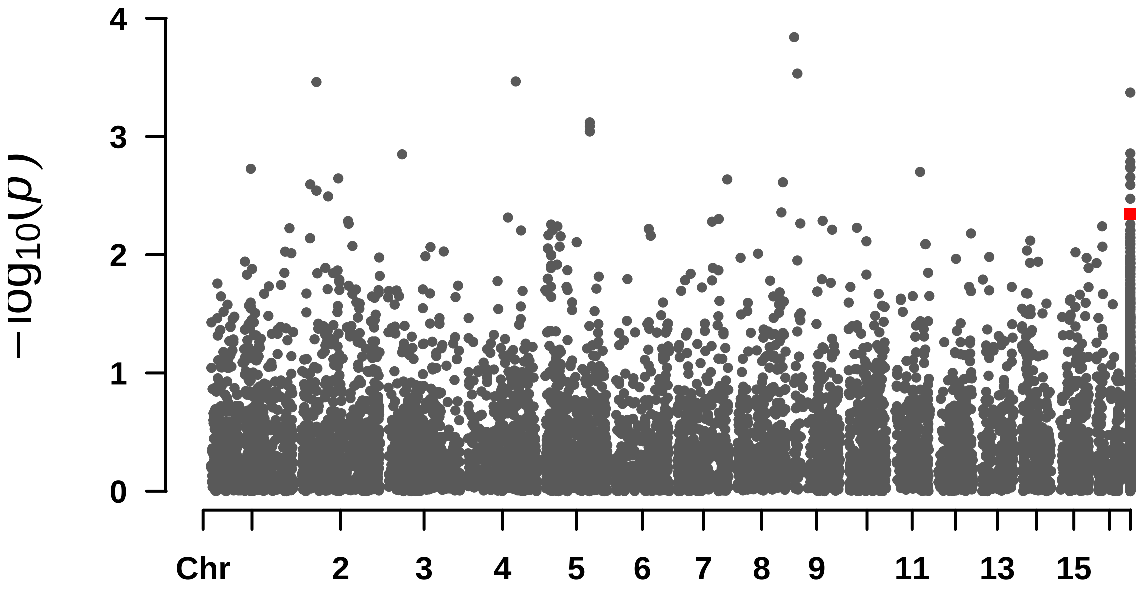


```
##  Rectangular-Manhattan Plotting MLMM.
```

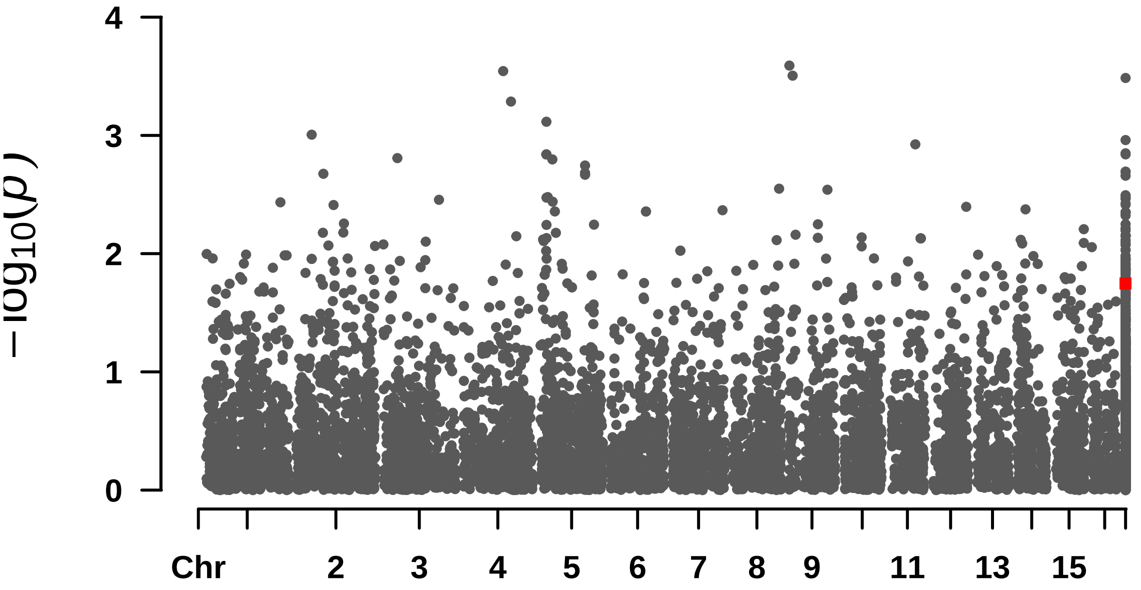
$-\log_{10}(p)$
0
1
2
3
4
Chr
2
3
4
5
6
7
8
9
11
13
15

—

#### F_Polpa

```
## $BLINK
## numeric(0)
##
## $FarmCPU
## numeric(0)
##
## $GLM
## numeric(0)
##
## $MLM
## numeric(0)
##
## $MLMM
## [1] 7.762472e-07 1.902984e-06 1.353202e-06 3.484940e-07
```

```
##  Rectangular-Manhattan Plotting BLINK.
```

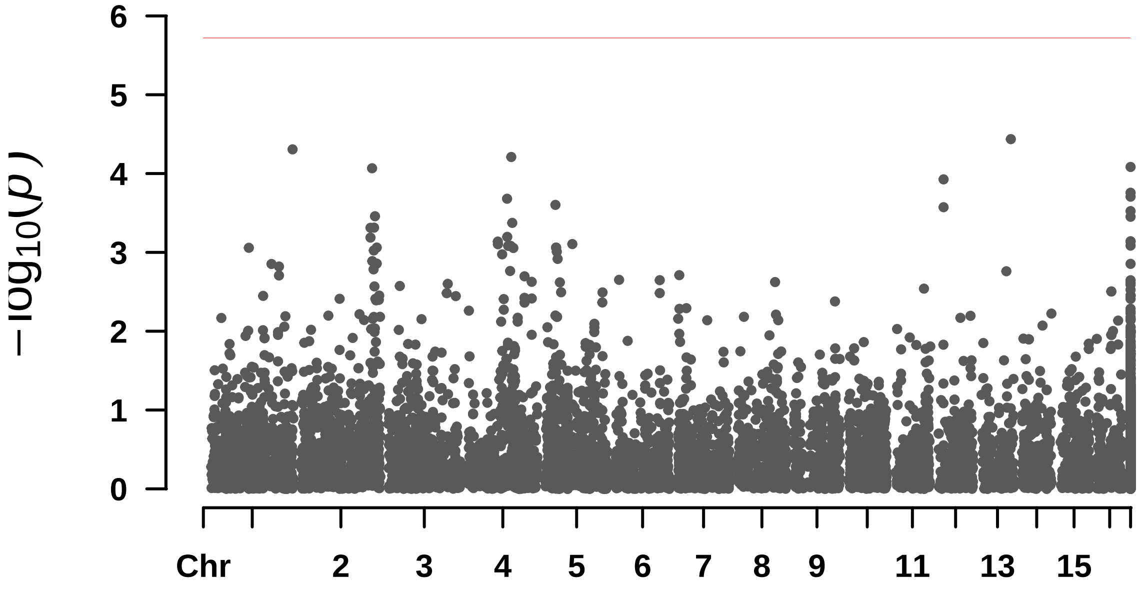


```
##  Rectangular-Manhattan Plotting FarmCPU.
```

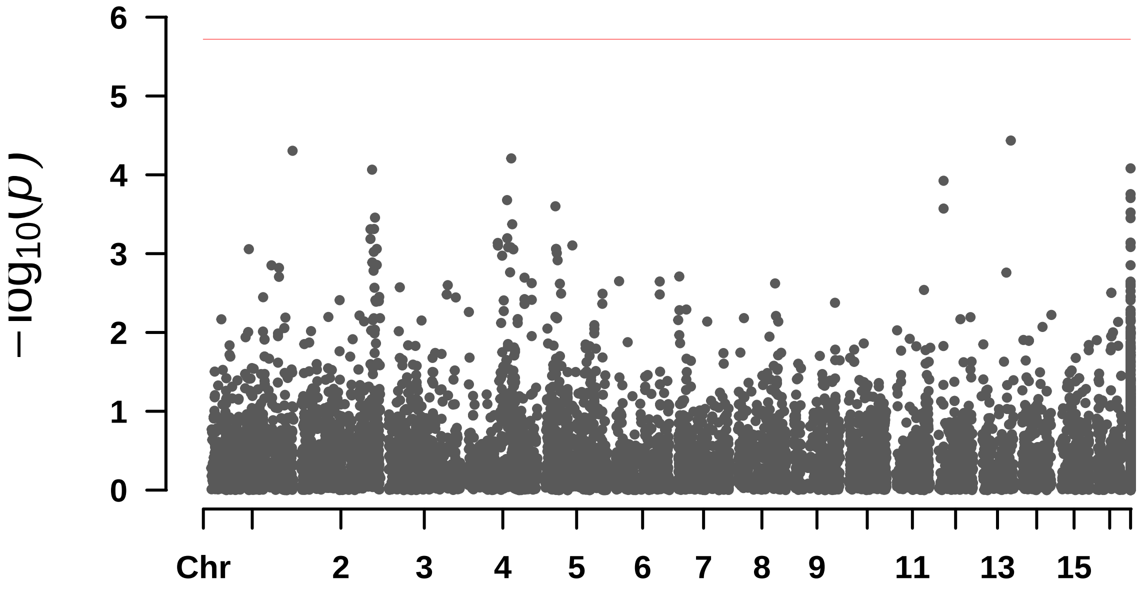


####  Rectangular-Manhattan Plotting GLM.

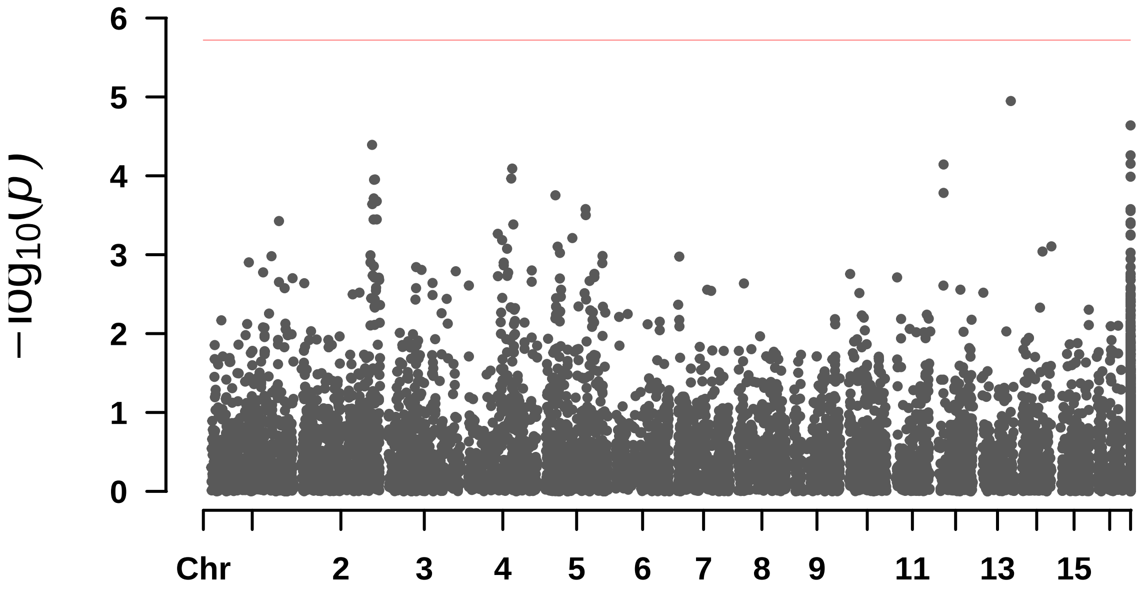


####  Rectangular-Manhattan Plotting MLM.

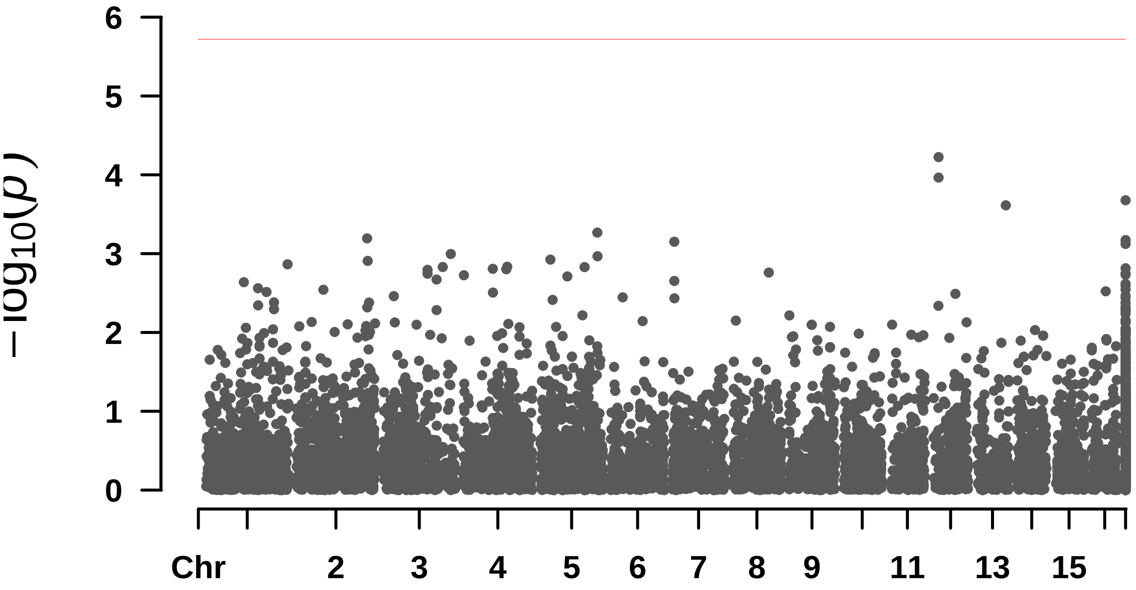


```
##  Rectangular-Manhattan Plotting MLMM.
```

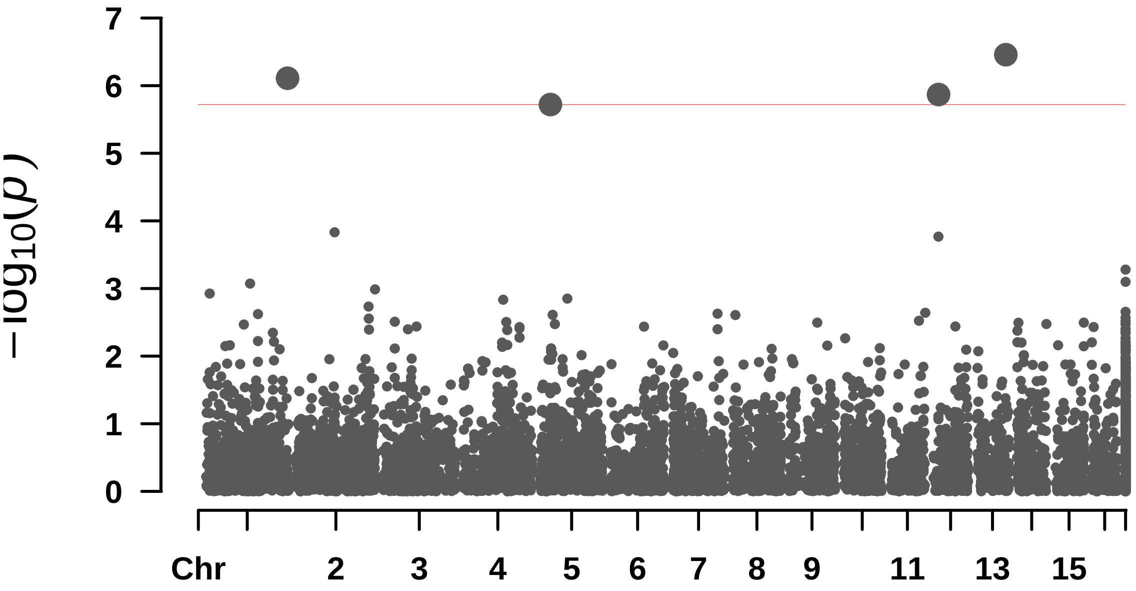

—

## ## F_T

```
## $BLINK
## [1] 3.467431e-06 1.899594e-06 1.232049e-07
##
## $FarmCPU
## numeric(0)
##
## $GLM
## numeric(0)
##
## $MLM
## numeric(0)
##
## $MLMM
## numeric(0)
```

```
##  Rectangular-Manhattan Plotting BLINK.
```

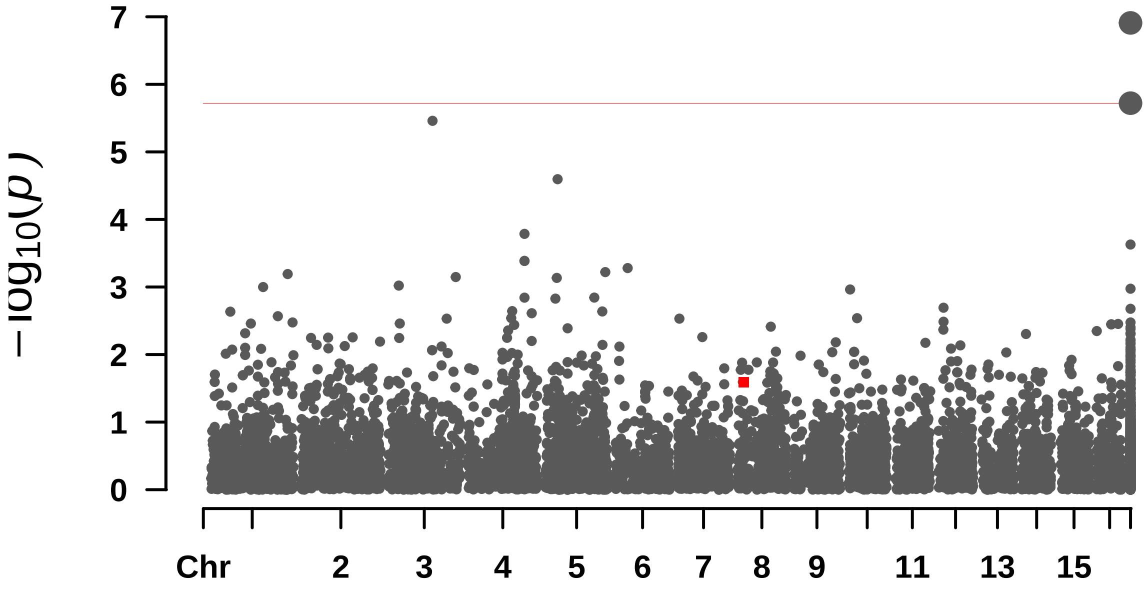


```
##  Rectangular-Manhattan Plotting FarmCPU.
```

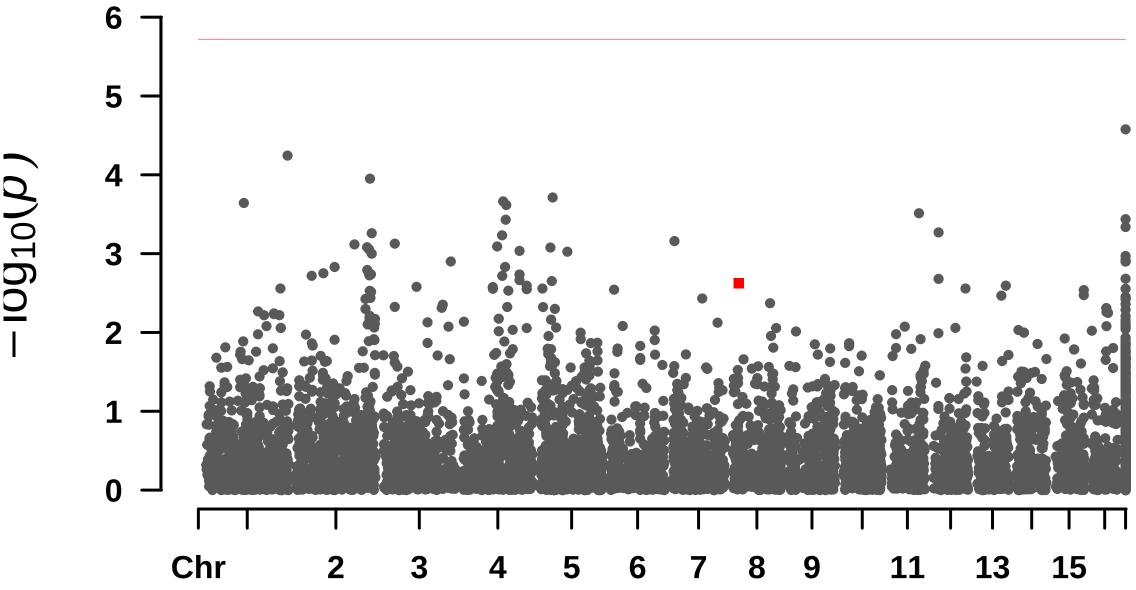


```
##  Rectangular-Manhattan Plotting GLM.
```

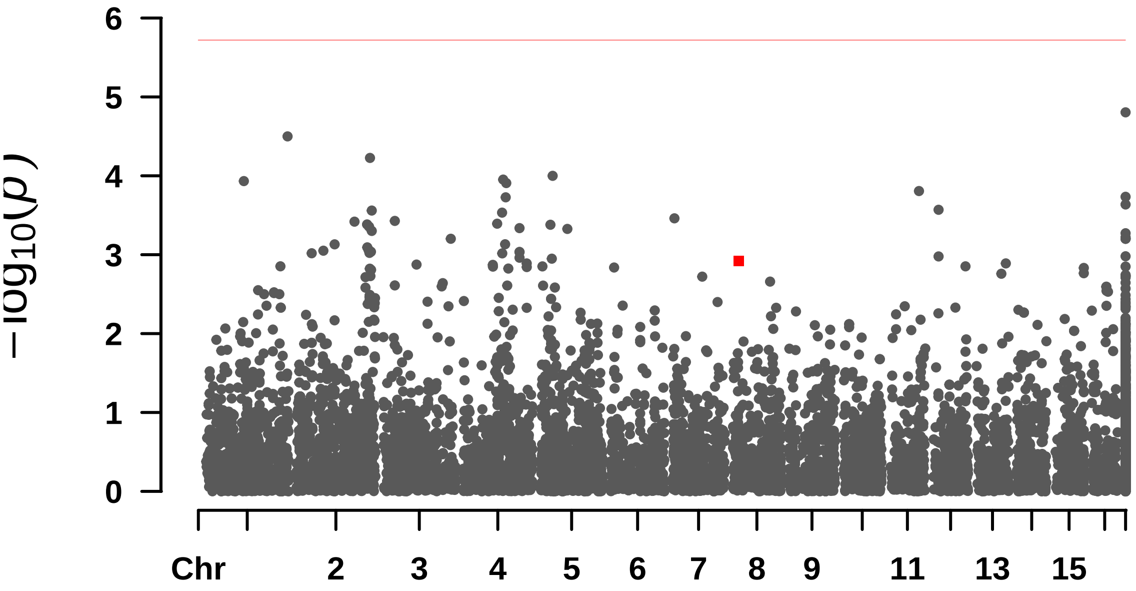


```
##  Rectangular-Manhattan Plotting MLM.
```

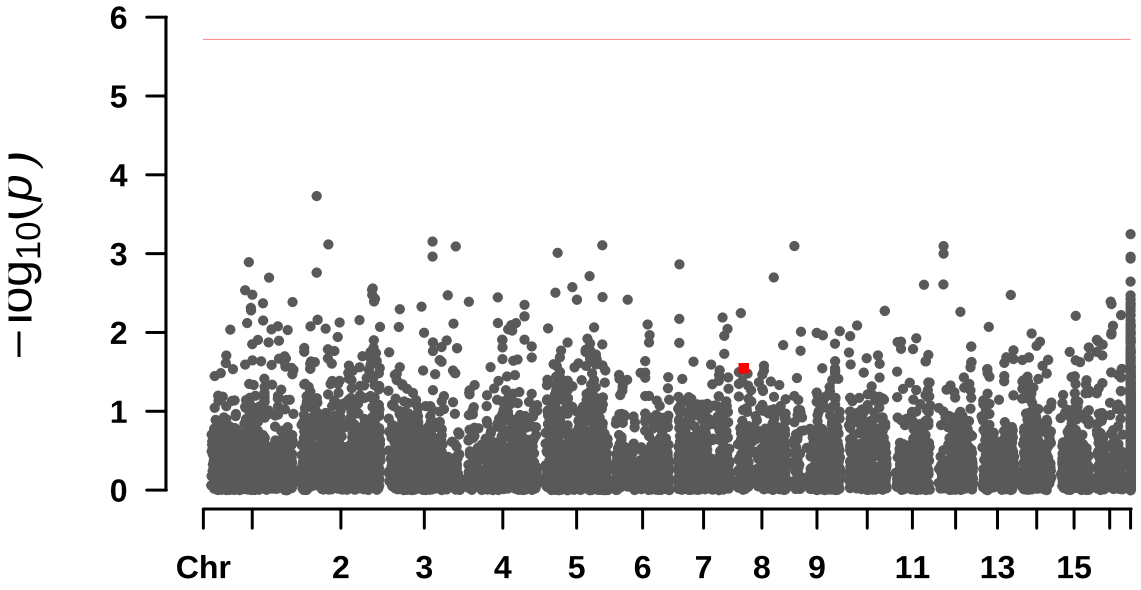


```
##  Rectangular-Manhattan Plotting MLMM.
```

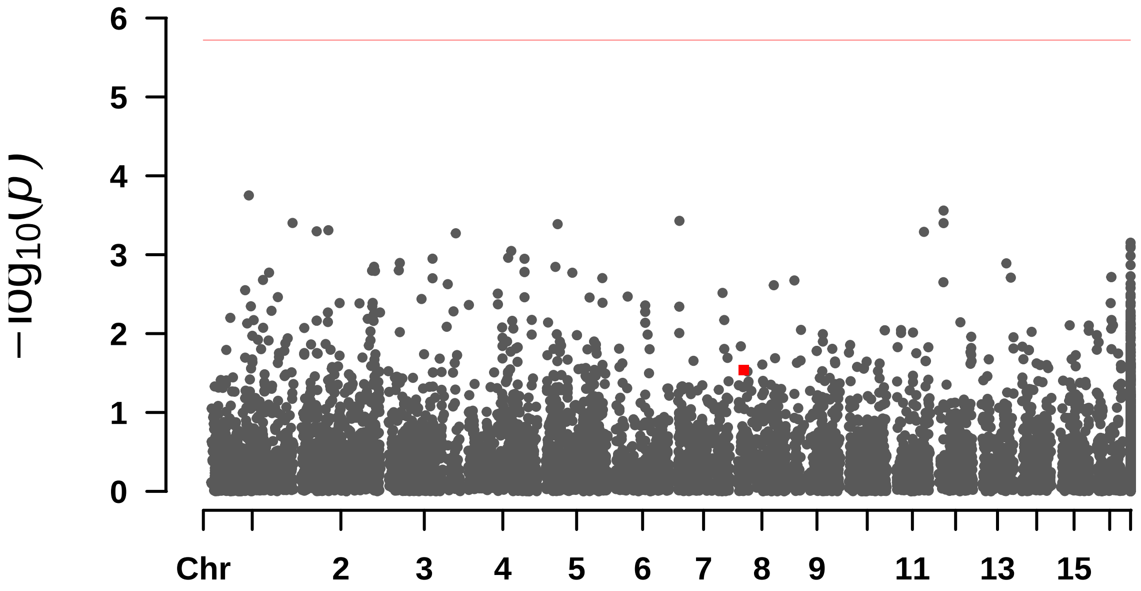

—

#### Largura_pina

```
## $BLINK
## [1] 4.822903e-11 1.735783e-08 3.431180e-06
##
## $FarmCPU
## [1] 1.498494e-09
##
## $GLM
## [1] 2.982141e-06
##
## $MLM
## numeric(0)
##
## $MLMM
## numeric(0)
```

```
##  Rectangular-Manhattan Plotting BLINK.
```

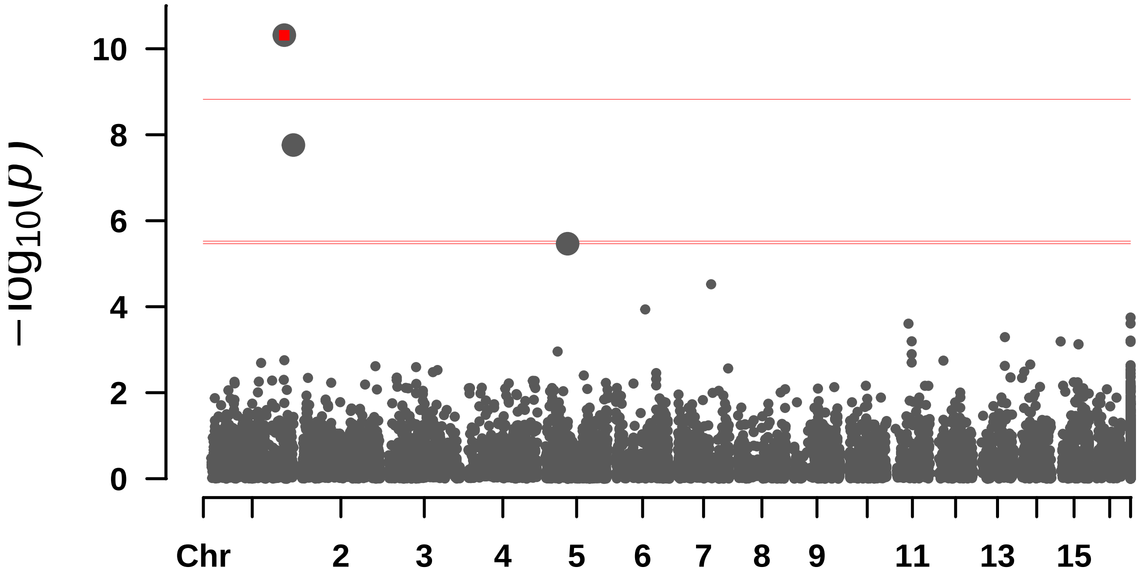


```
##  Rectangular-Manhattan Plotting FarmCPU.
```

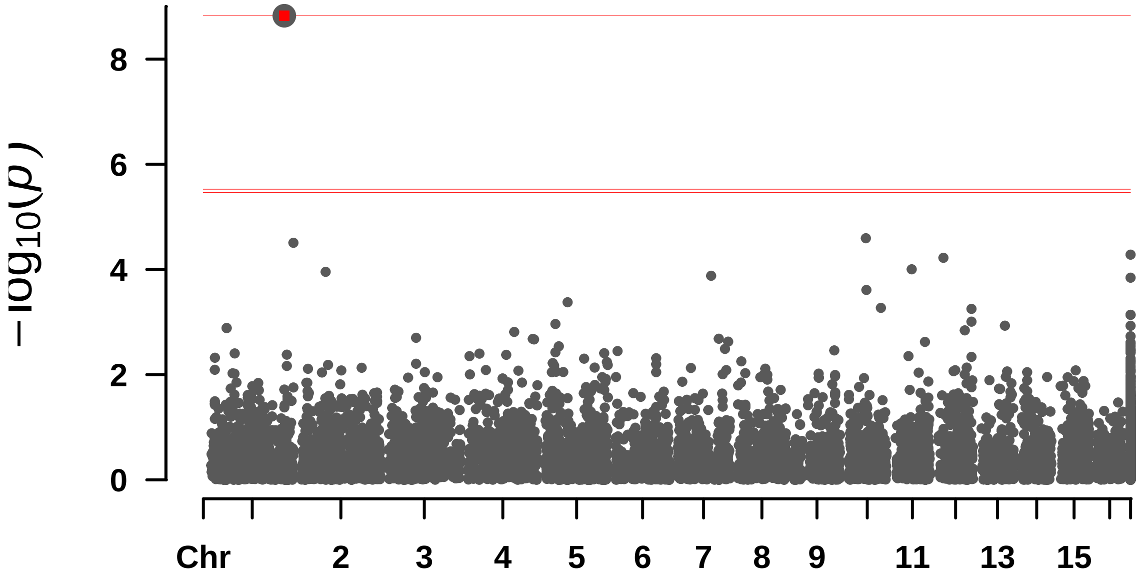


#### Rectangular-Manhattan Plotting GLM.

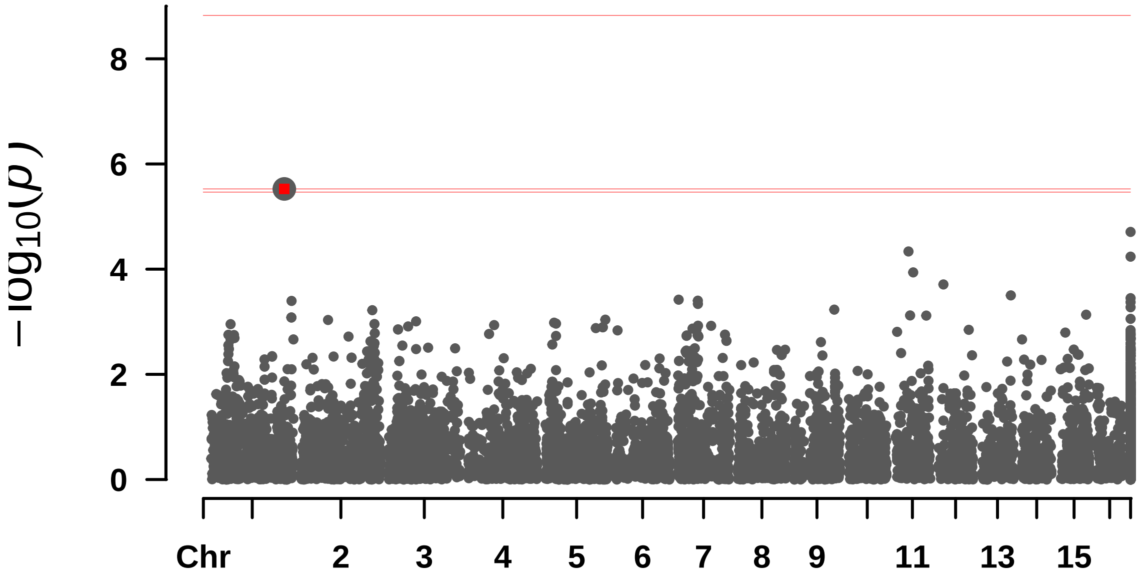


#### Rectangular-Manhattan Plotting MLM.

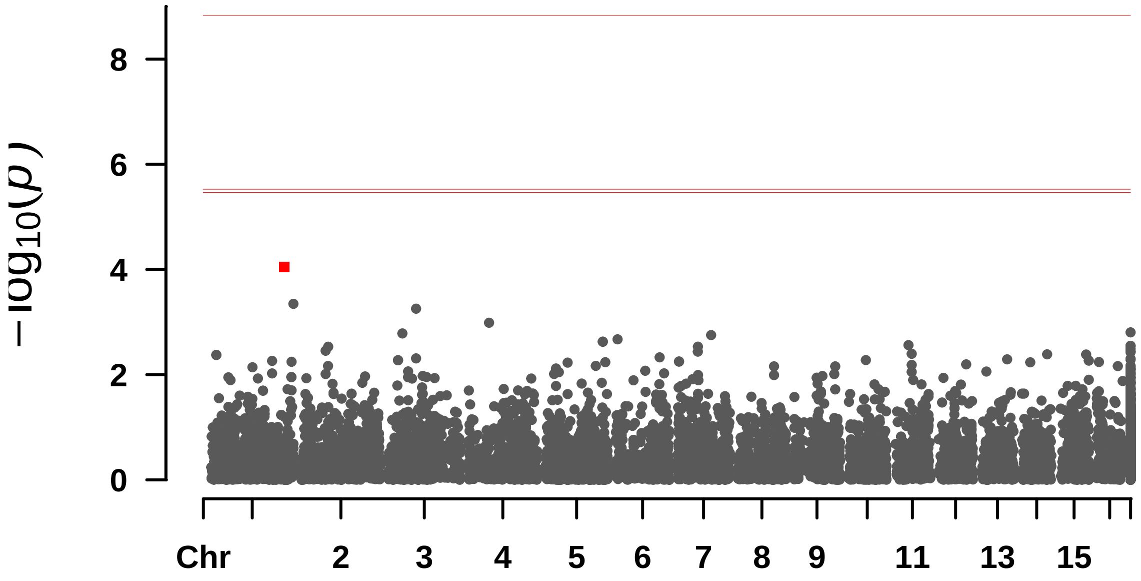


```
##  Rectangular-Manhattan Plotting MLMM.
```

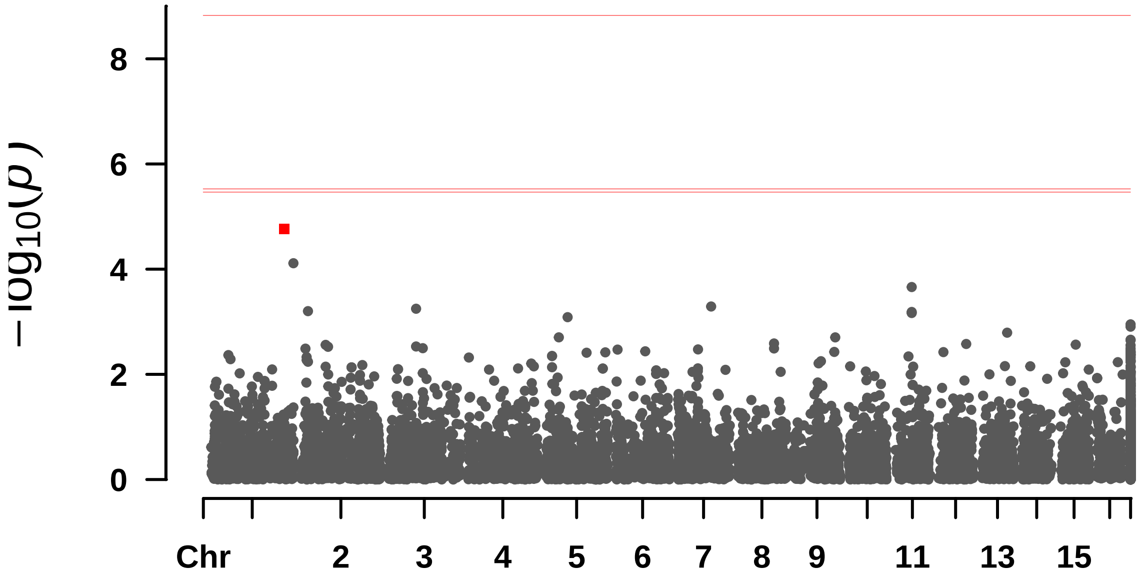

—

#### N_folhas

```
## numeric(0)
```

```
##  Rectangular-Manhattan Plotting BLINK.
```

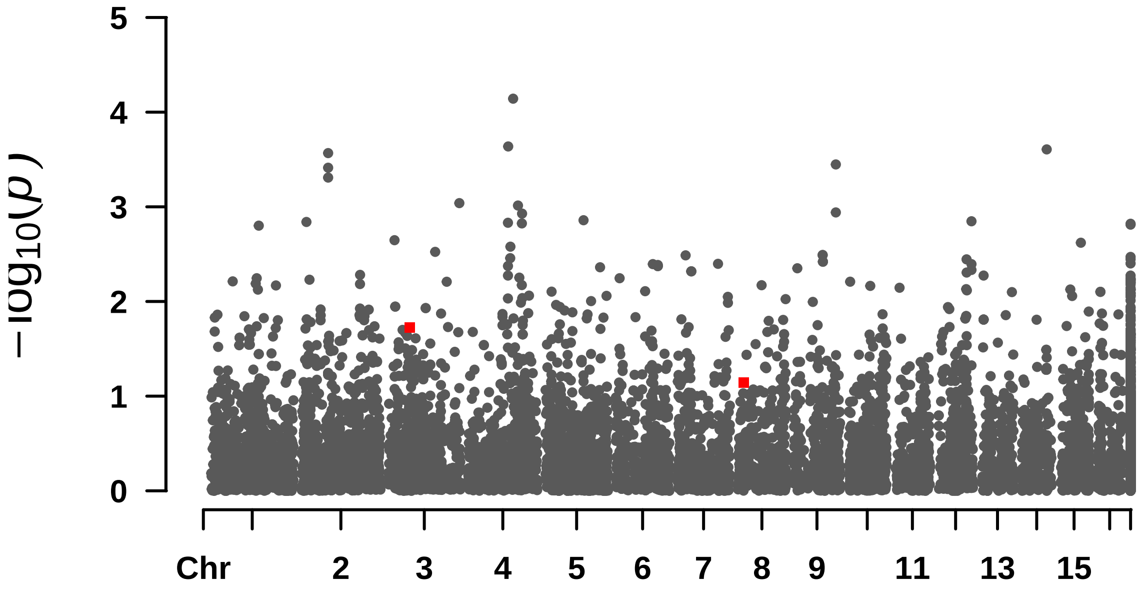


```
##  Rectangular-Manhattan Plotting FarmCPU.
```

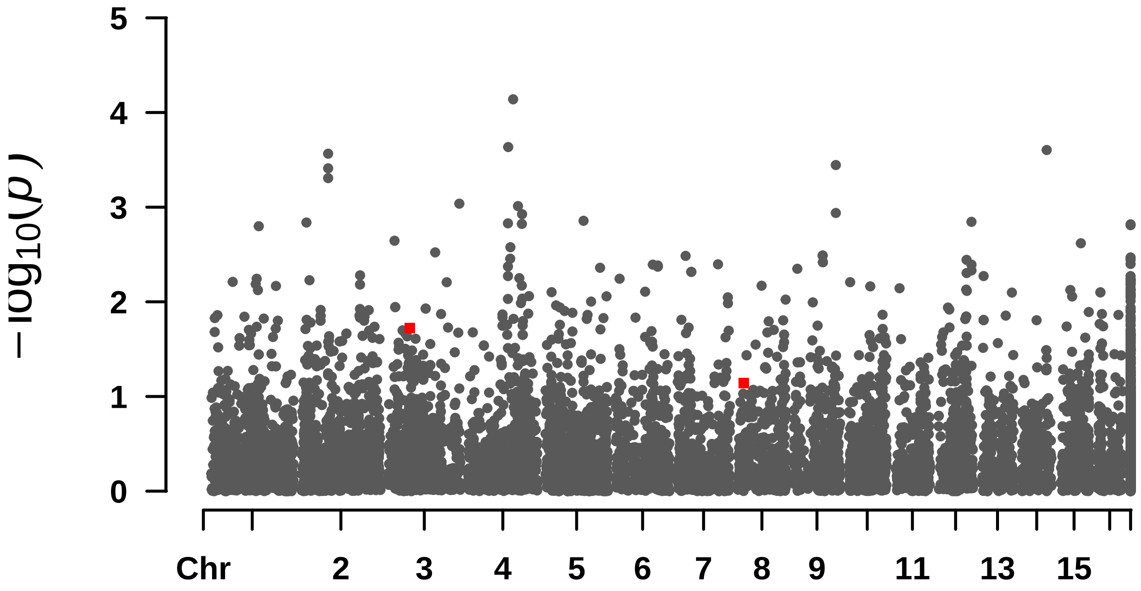


```
##  Rectangular-Manhattan Plotting GLM.
```

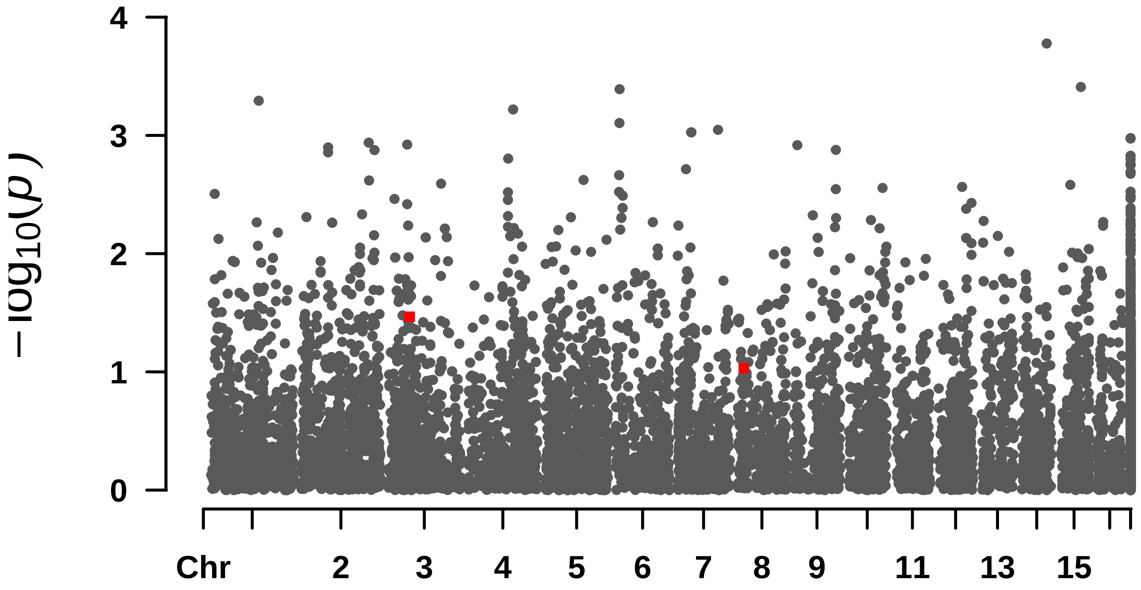


```
##  Rectangular-Manhattan Plotting MLM.
```

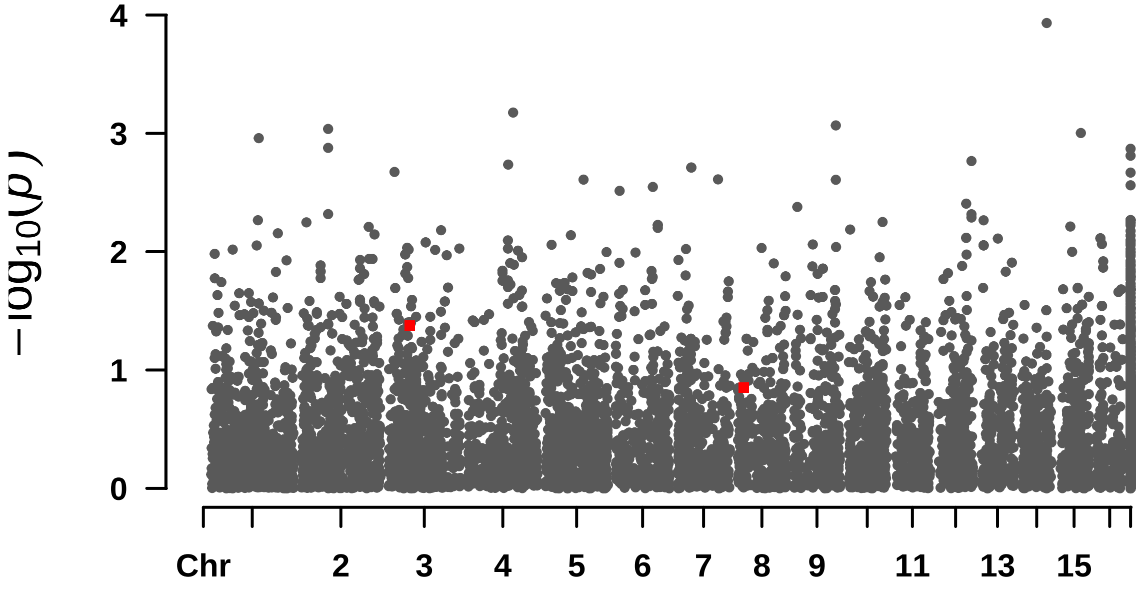


```
##  Rectangular-Manhattan Plotting MLMM.
```

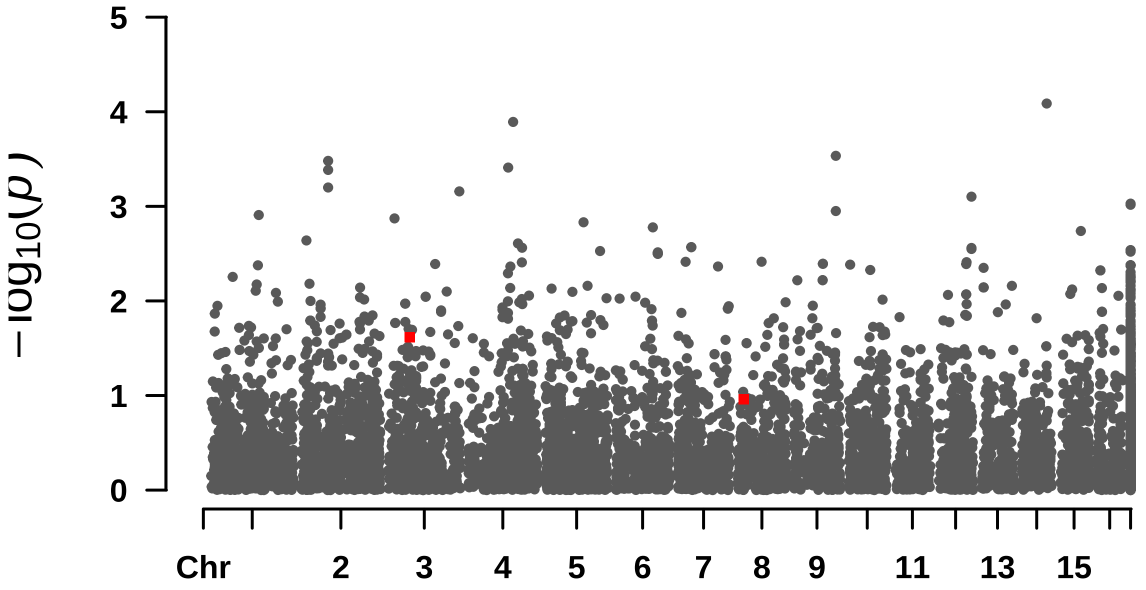

5
4
3
2
1
0
$-\log_{10}(p)$
Chr
2
3
4
5
6
7
8
9
11
13
15

—

#### N_pinas

```
## numeric(0)
```

```
##  Rectangular-Manhattan Plotting BLINK.
```

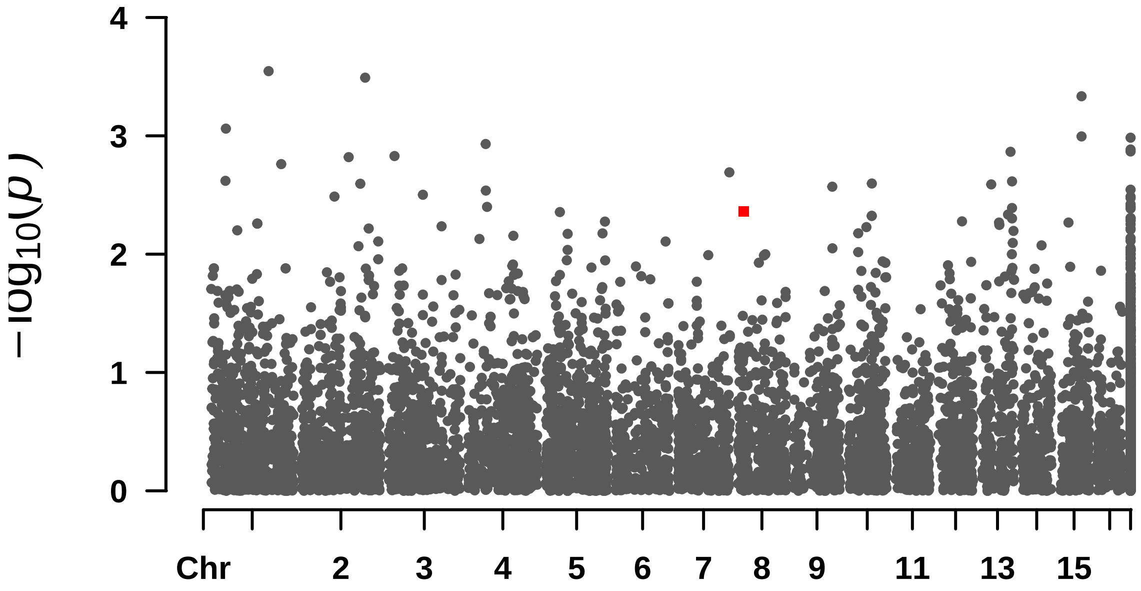


```
##  Rectangular-Manhattan Plotting FarmCPU.
```

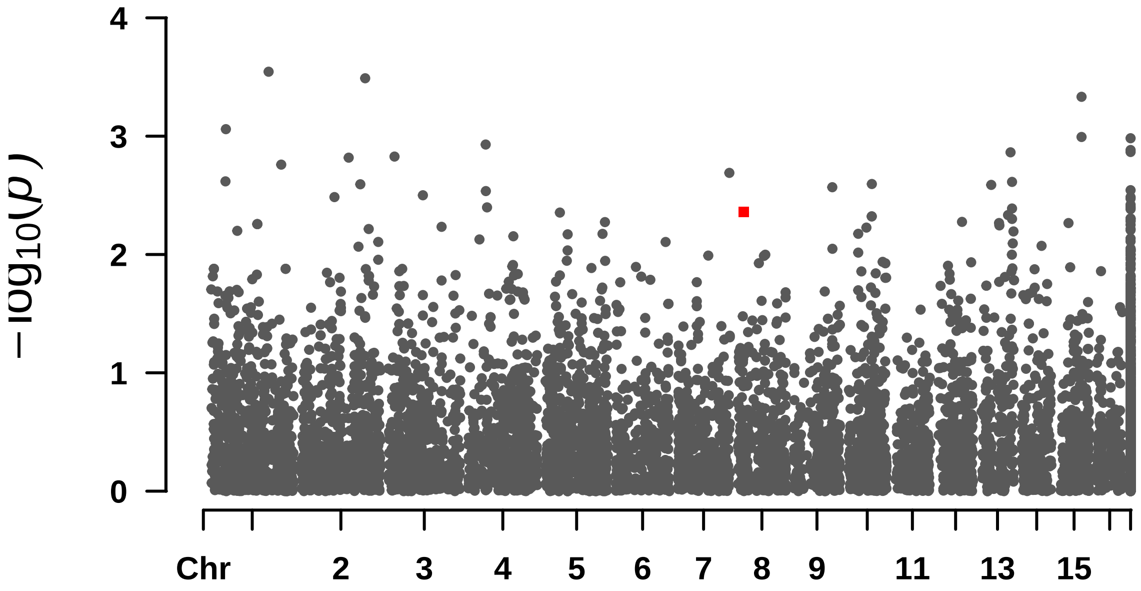


```
##  Rectangular-Manhattan Plotting GLM.
```

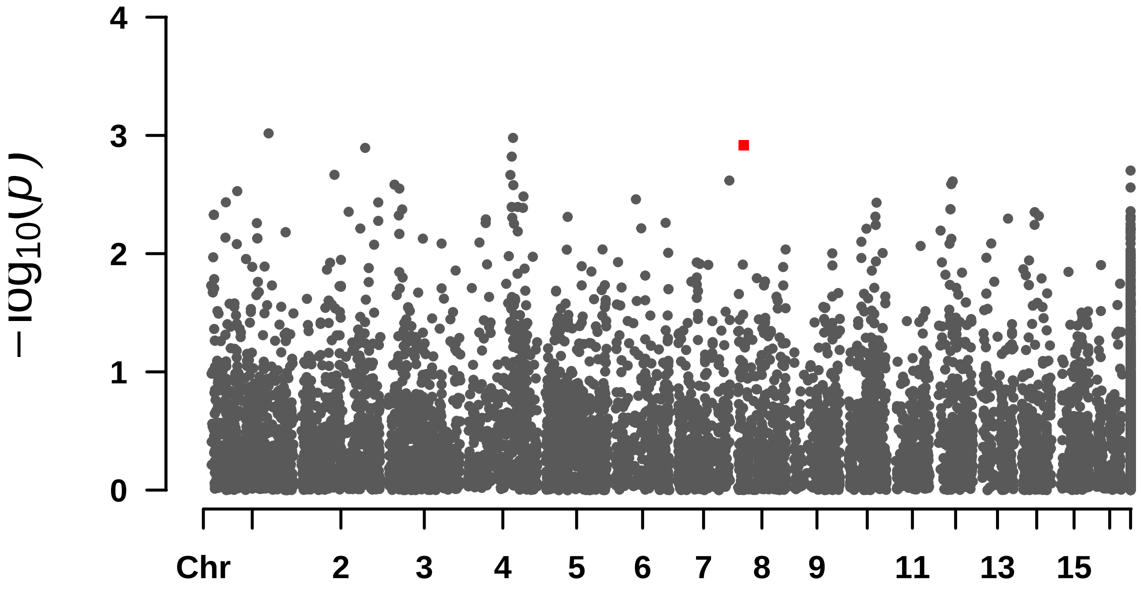


#### Rectangular-Manhattan Plotting MLM.

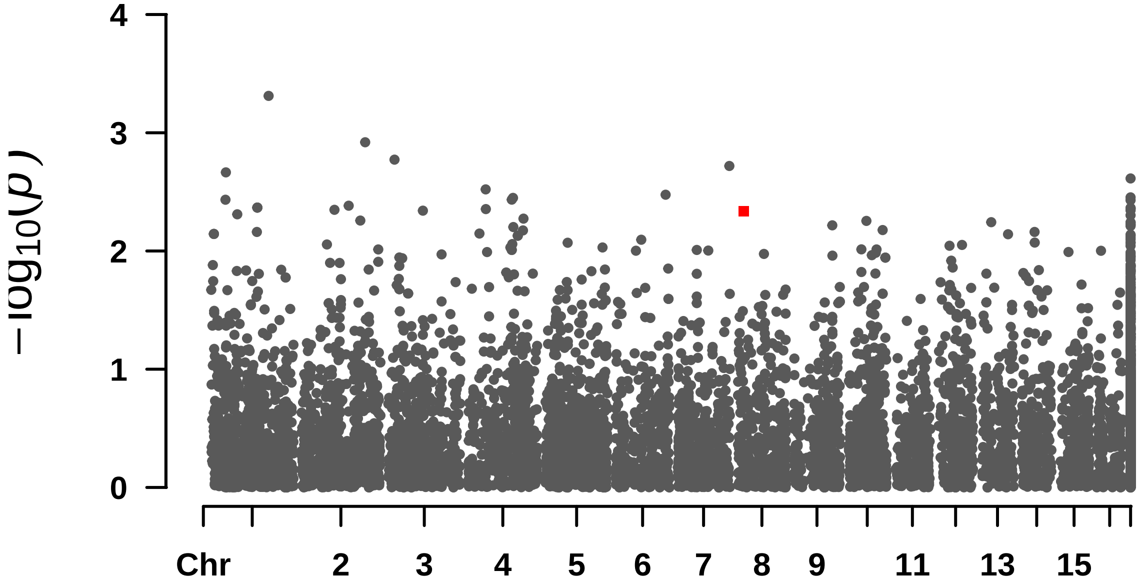


#### Rectangular-Manhattan Plotting MLMM.

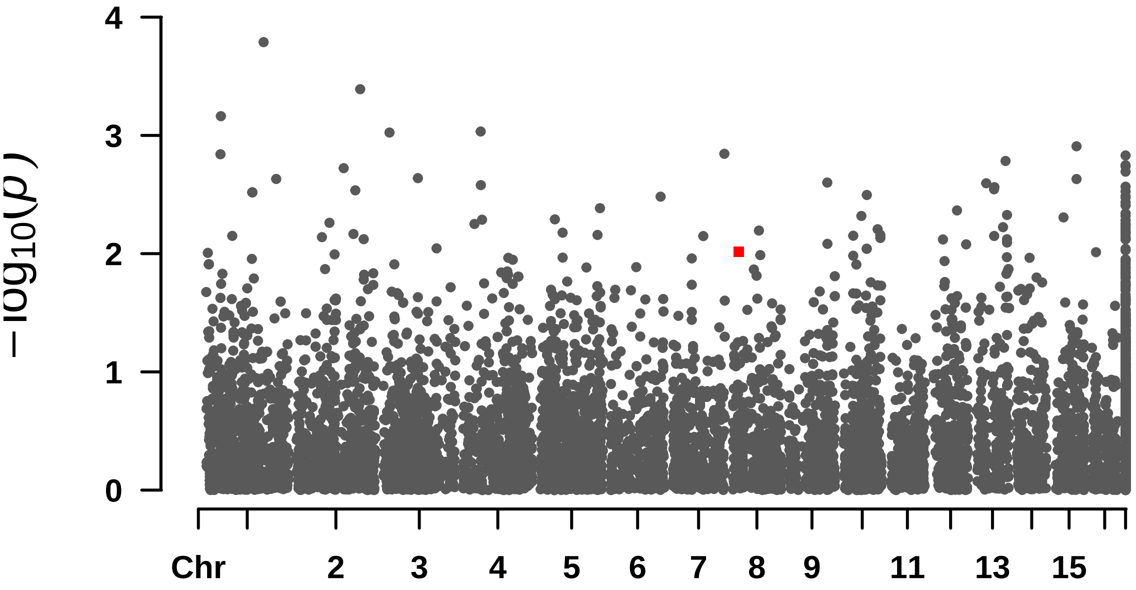

$-\log_{10}(p)$
0
1
2
3
4
Chr
2
3
4
5
6
7
8
9
11
13
15

—

#### ## Oil_content

```
## $BLINK
## [1] 1.404943e-05 1.174886e-05 8.391095e-06 1.291354e-05
##
## $FarmCPU
## [1] 3.594279e-06 3.213166e-07 3.992582e-06 7.674756e-06 3.096688e-09
##
## $GLM
## [1] 2.342878e-06
##
## $MLM
## numeric(0)
##
## $MLMM
## numeric(0)
```

```
##  Rectangular-Manhattan Plotting BLINK.
```

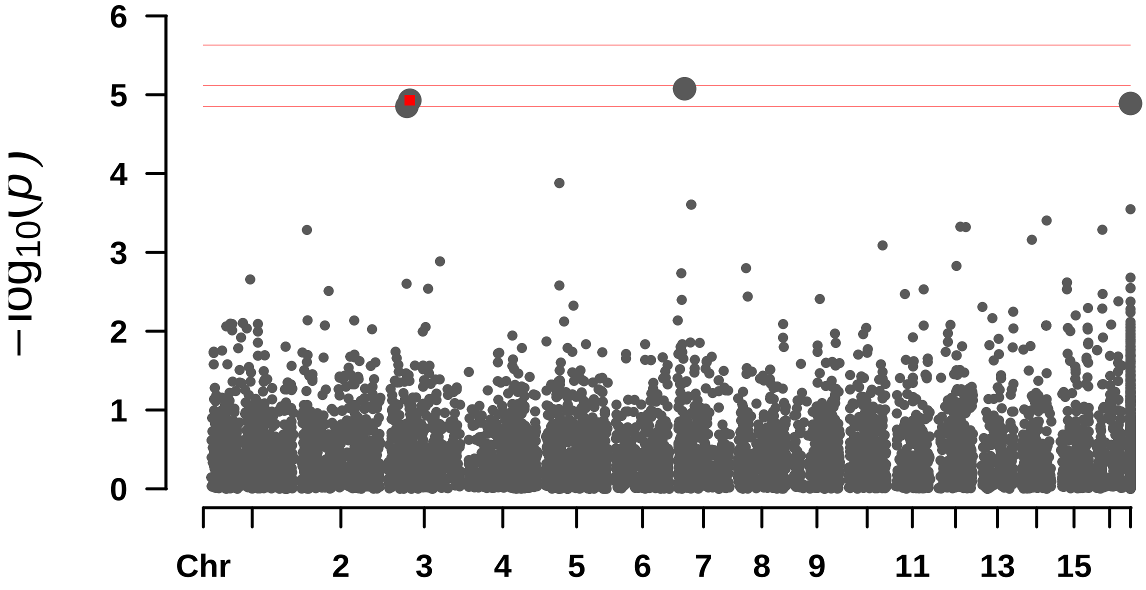


```
##  Rectangular-Manhattan Plotting FarmCPU.
```

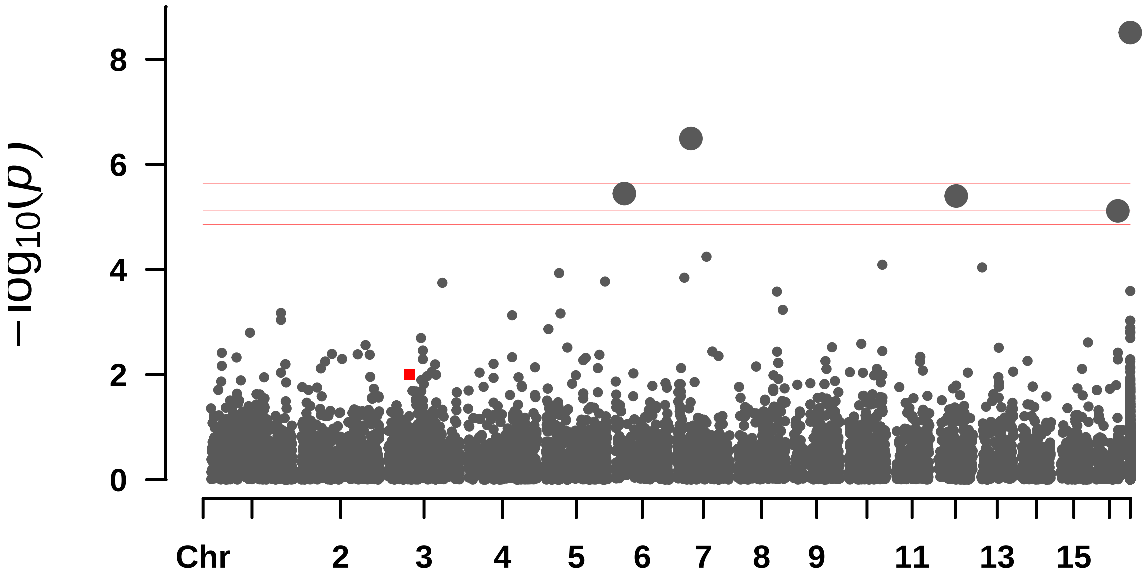


```
##  Rectangular-Manhattan Plotting GLM.
```

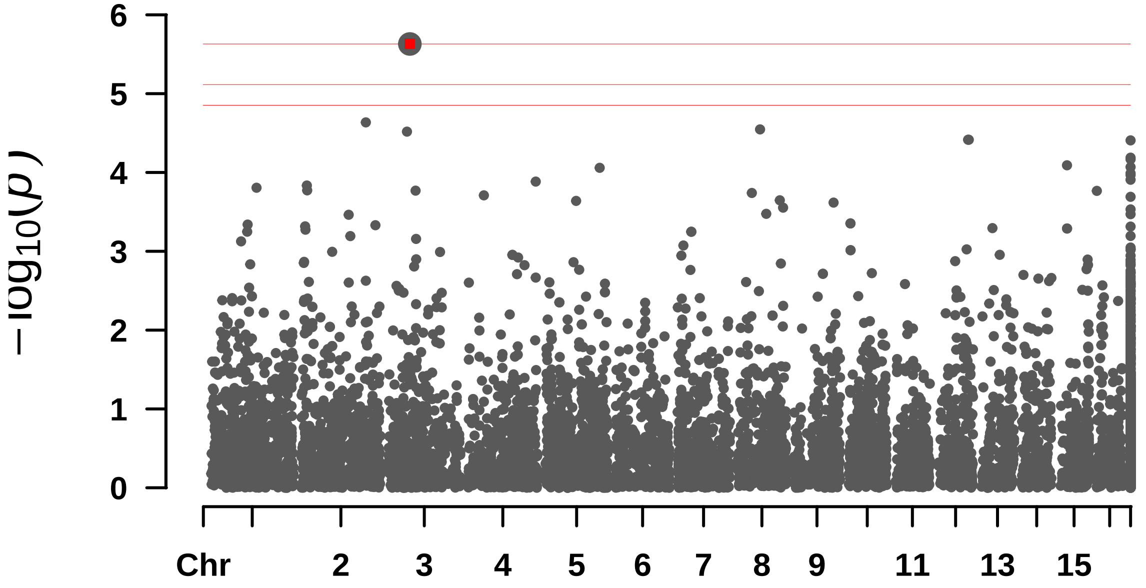


```
##  Rectangular-Manhattan Plotting MLM.
```

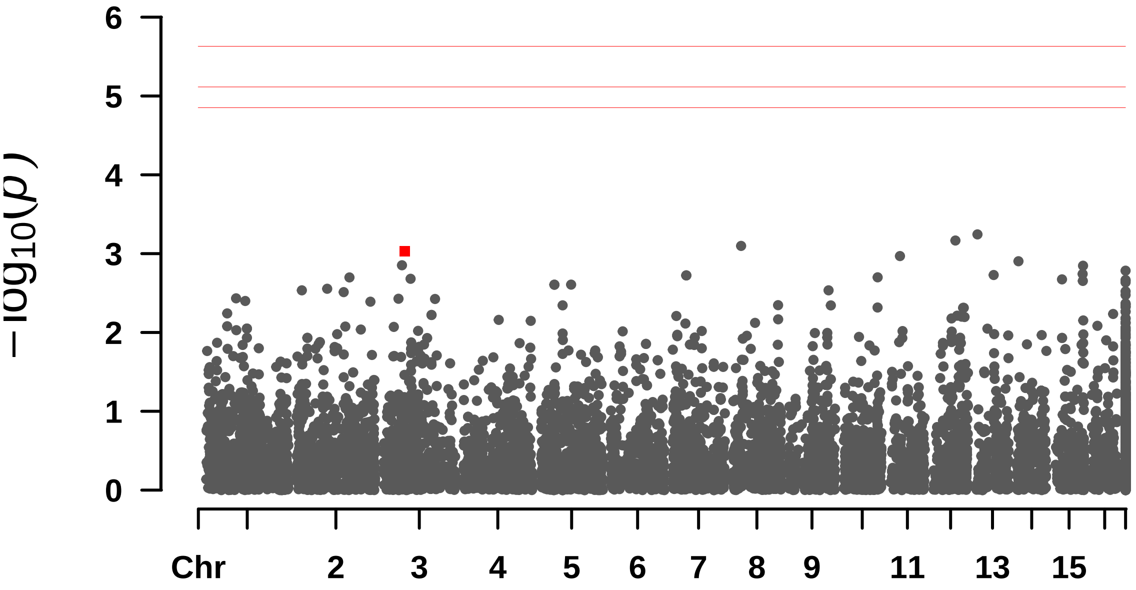


```
## Rectangular-Manhattan Plotting MLMM.
```

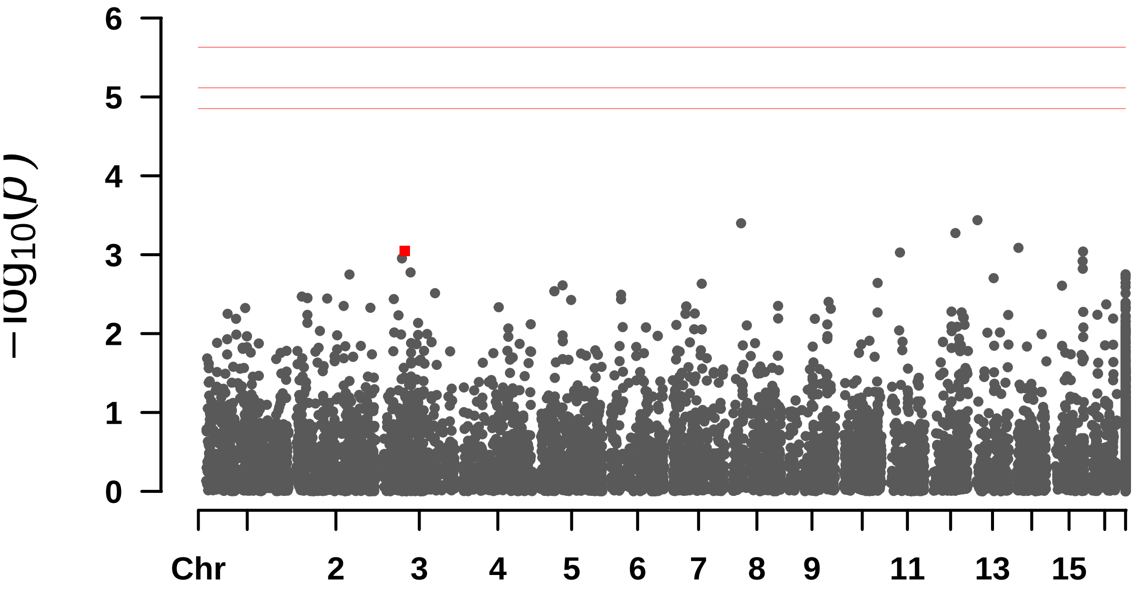

—

#### S_Casca

```
## numeric(0)
```

```
##  Rectangular-Manhattan Plotting BLINK.
```

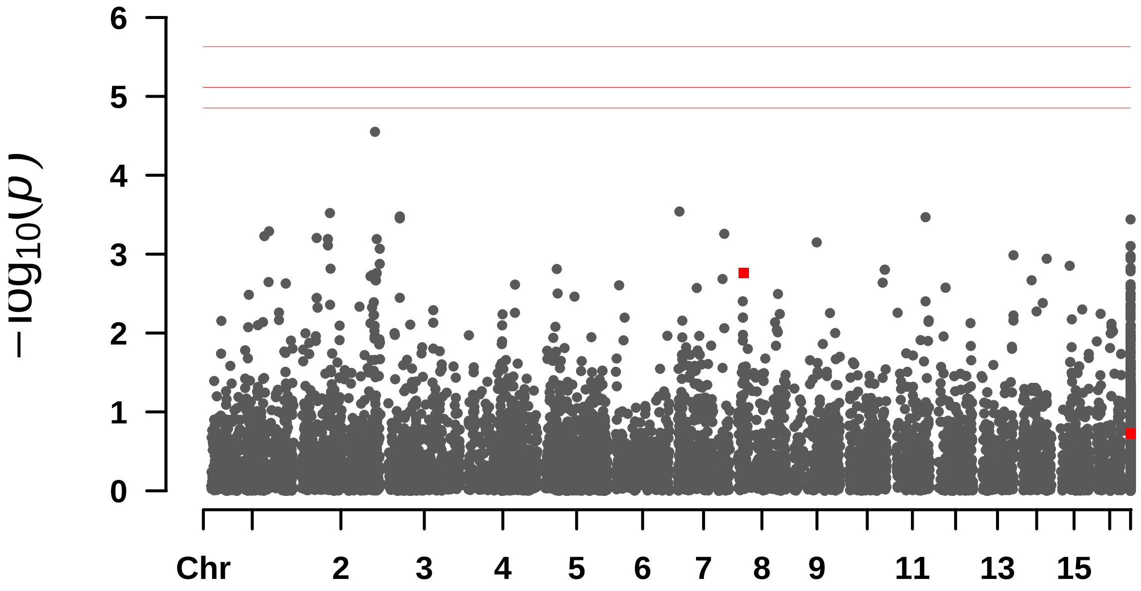


```
##  Rectangular-Manhattan Plotting FarmCPU.
```

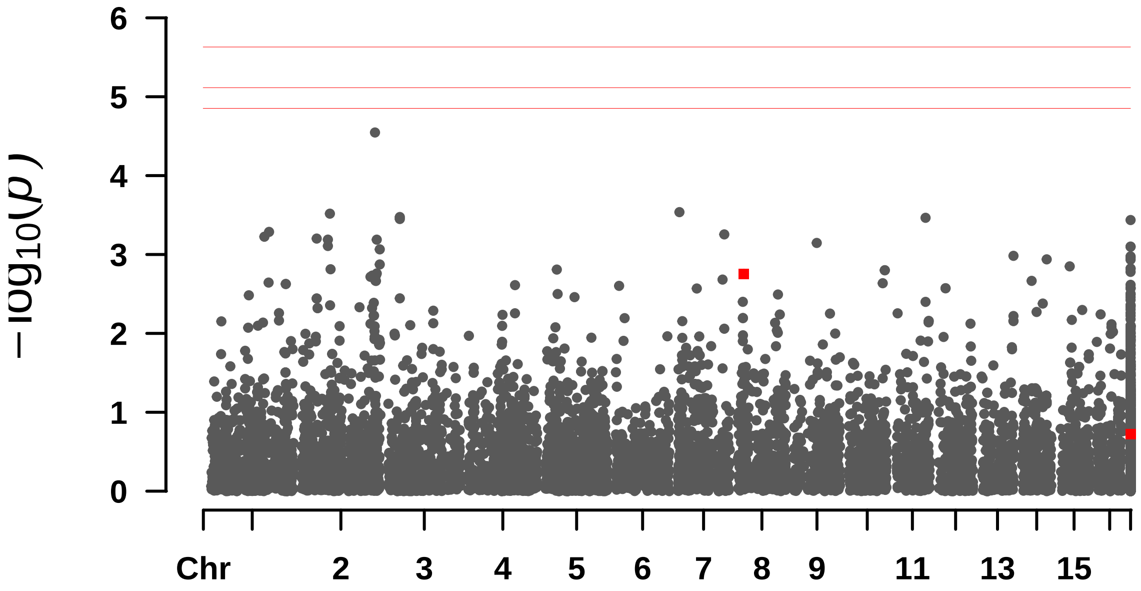


```
##  Rectangular-Manhattan Plotting GLM.
```

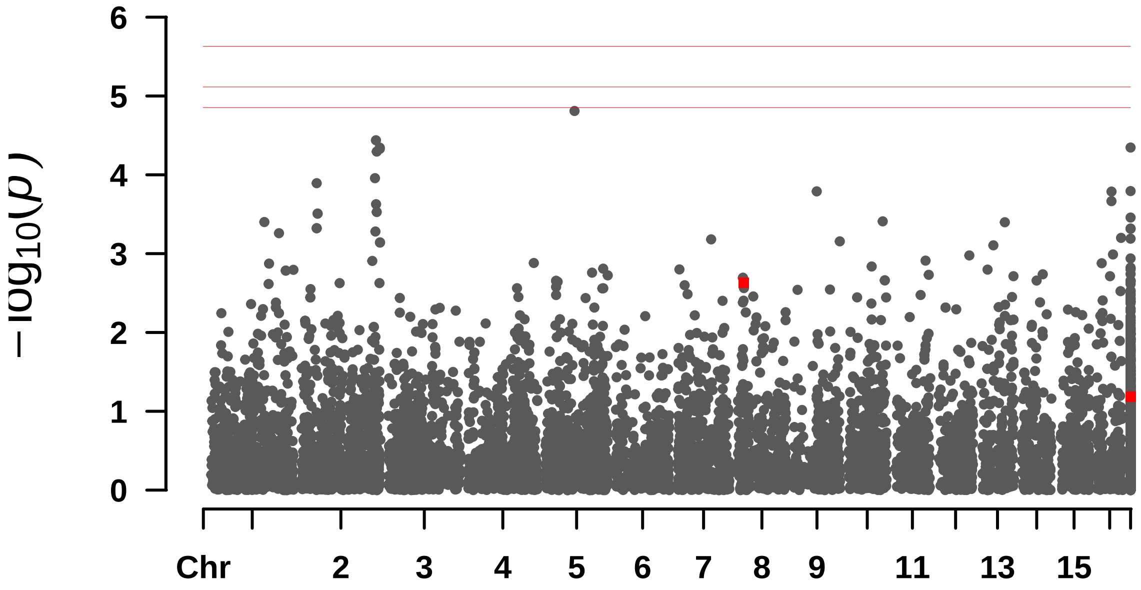


#### Rectangular-Manhattan Plotting MLM.

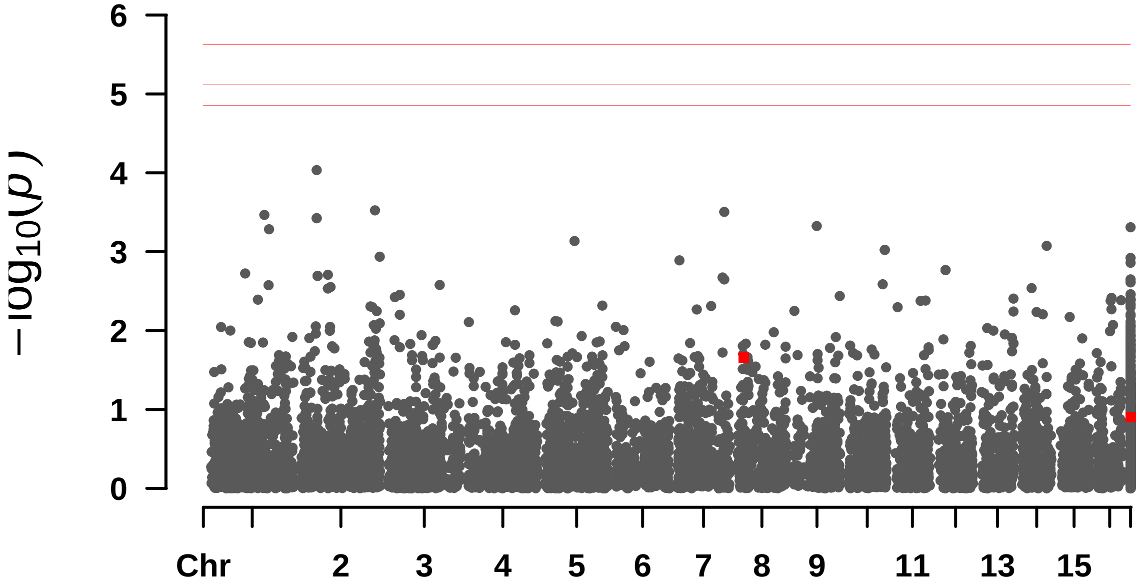


#### Rectangular-Manhattan Plotting MLMM.

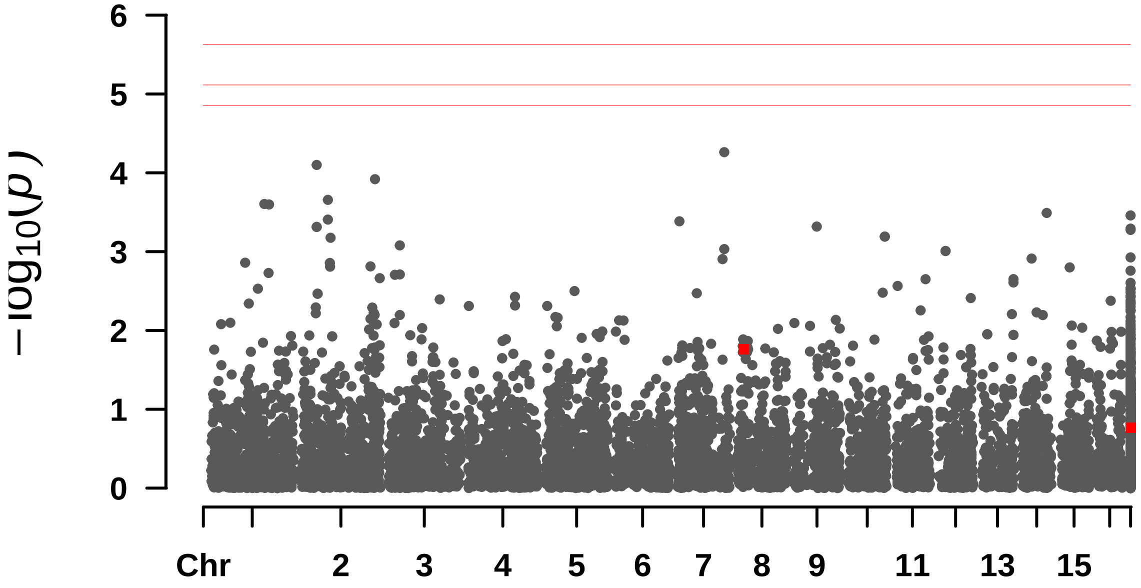
6
5
4
3
2
1
0
$-\log_{10}(p)$
Chr
2
3
4
5
6
7
8
9
11
13
15

#### ## S__Polpa

```
## $BLINK
## numeric(0)
##
## $FarmCPU
## numeric(0)
##
## $GLM
## numeric(0)
##
## $MLM
## numeric(0)
##
## $MLMM
## [1] 9.733702e-08 6.474075e-07
```

```
##  Rectangular-Manhattan Plotting BLINK.
```

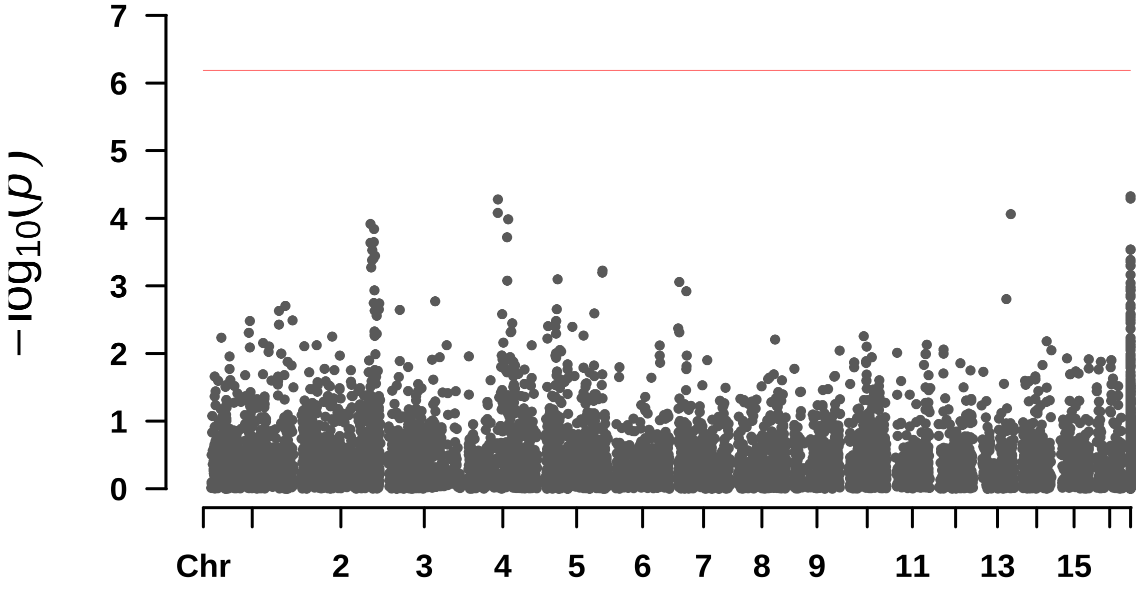


```
##  Rectangular-Manhattan Plotting FarmCPU.
```

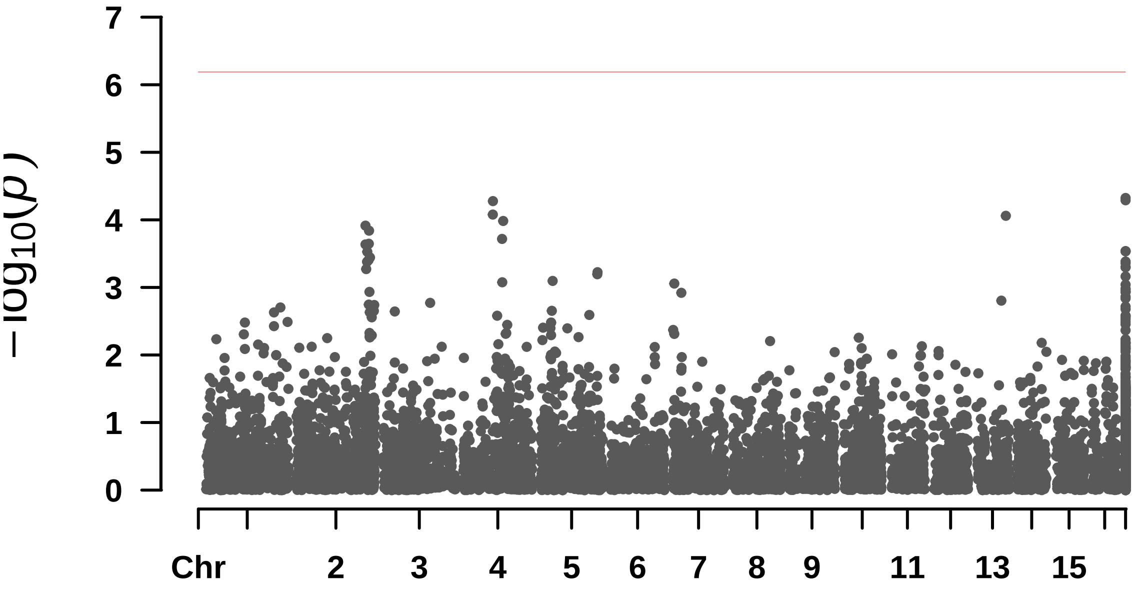


#### Rectangular-Manhattan Plotting GLM.

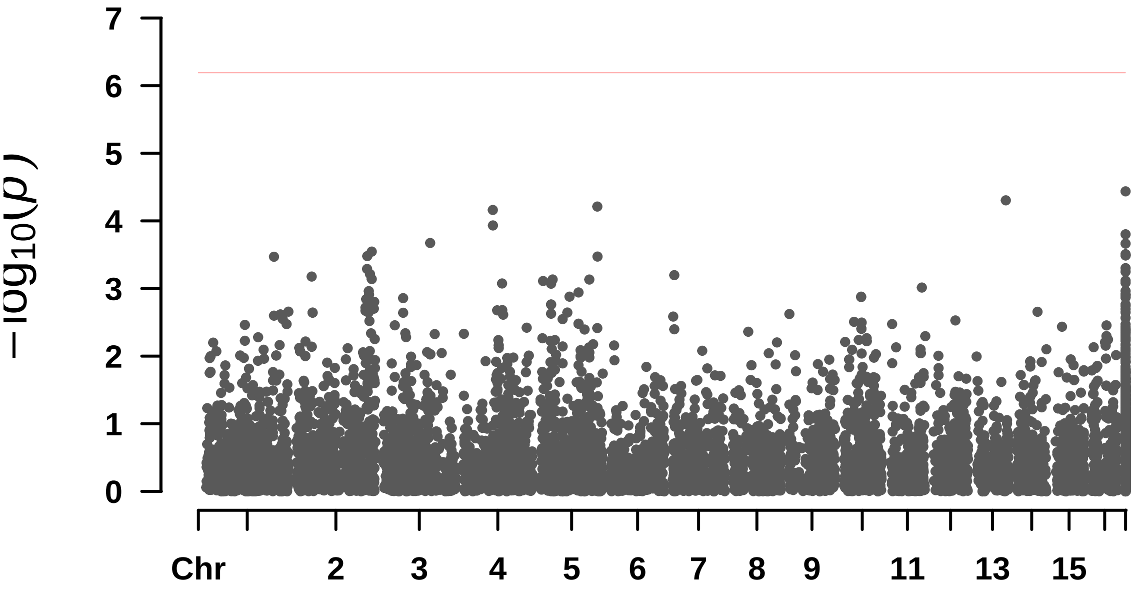


#### Rectangular-Manhattan Plotting MLM.

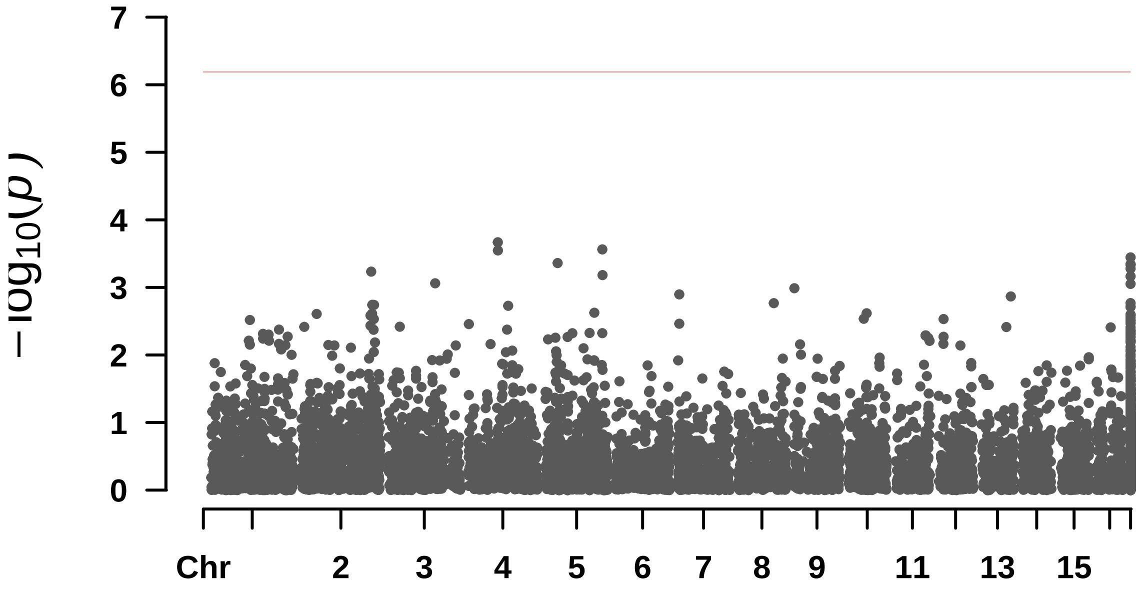


```
##  Rectangular-Manhattan Plotting MLMM.
```

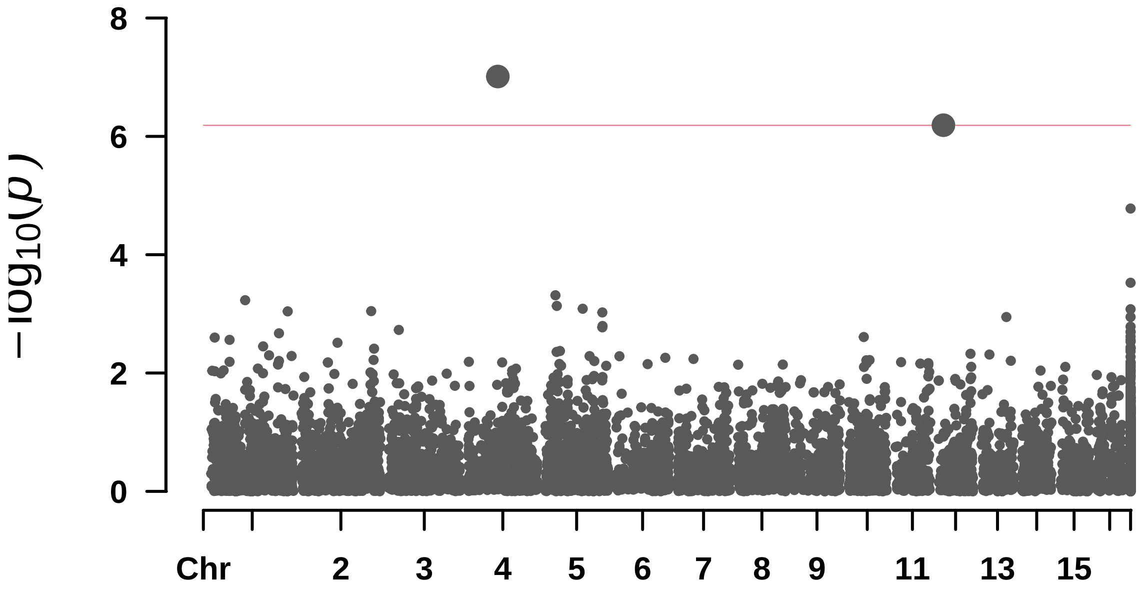


### Single trait GWAS figures - DENOVO

#### Compri_folha

```
## $BLINK
## [1] 1.281864e-09 4.265844e-08 1.075509e-11
##
## $FarmCPU
## numeric(0)
##
## $GLM
## numeric(0)
##
## $MLM
## numeric(0)
```

```
##
## $MLMM
## numeric(0)
```

```
##  Rectangular-Manhattan Plotting BLINK.
```

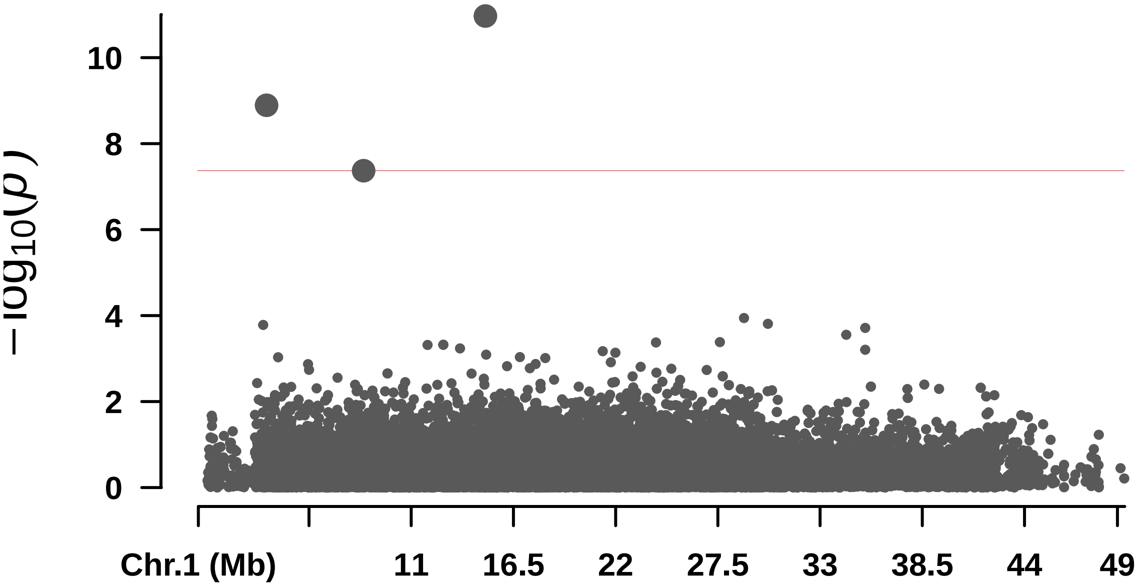


```
##  Rectangular-Manhattan Plotting FarmCPU.
```

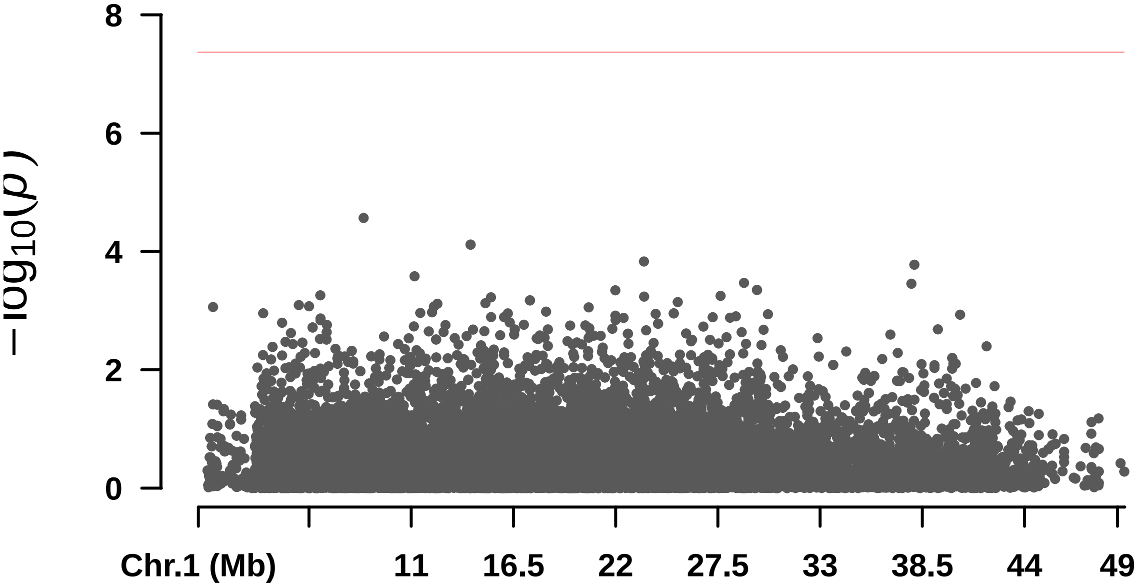


```
##  Rectangular-Manhattan Plotting GLM.
```

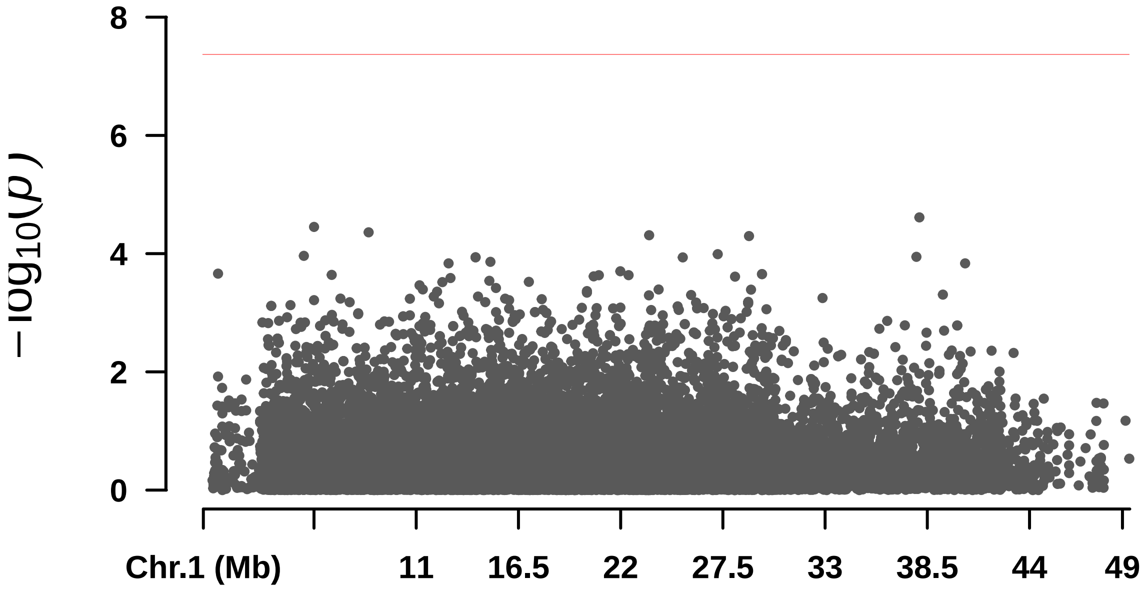


```
##  Rectangular-Manhattan Plotting MLM.
```

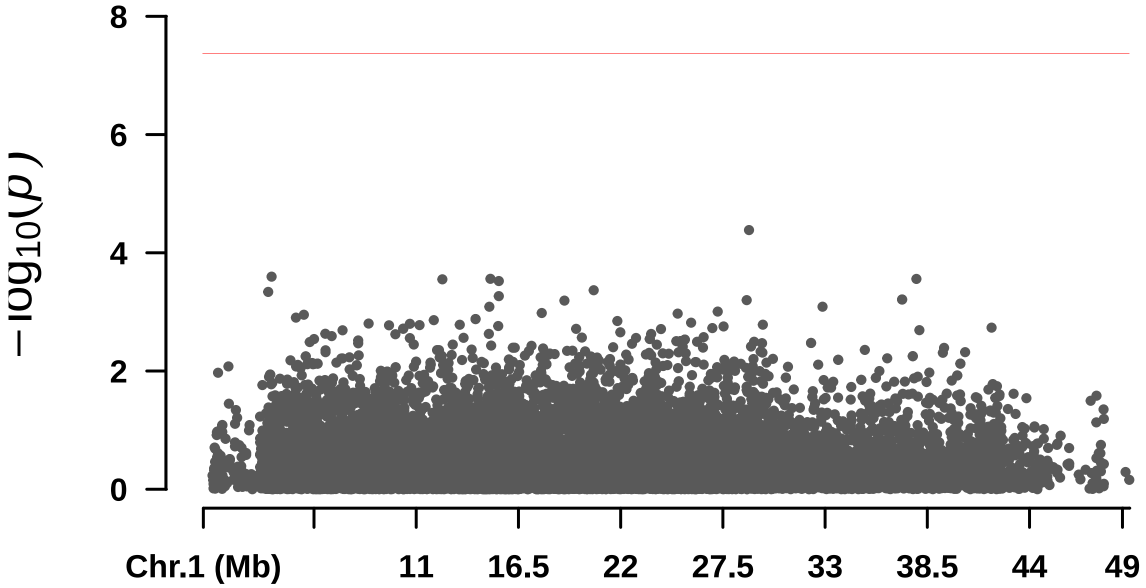


```
##  Rectangular-Manhattan Plotting MLMM.
```

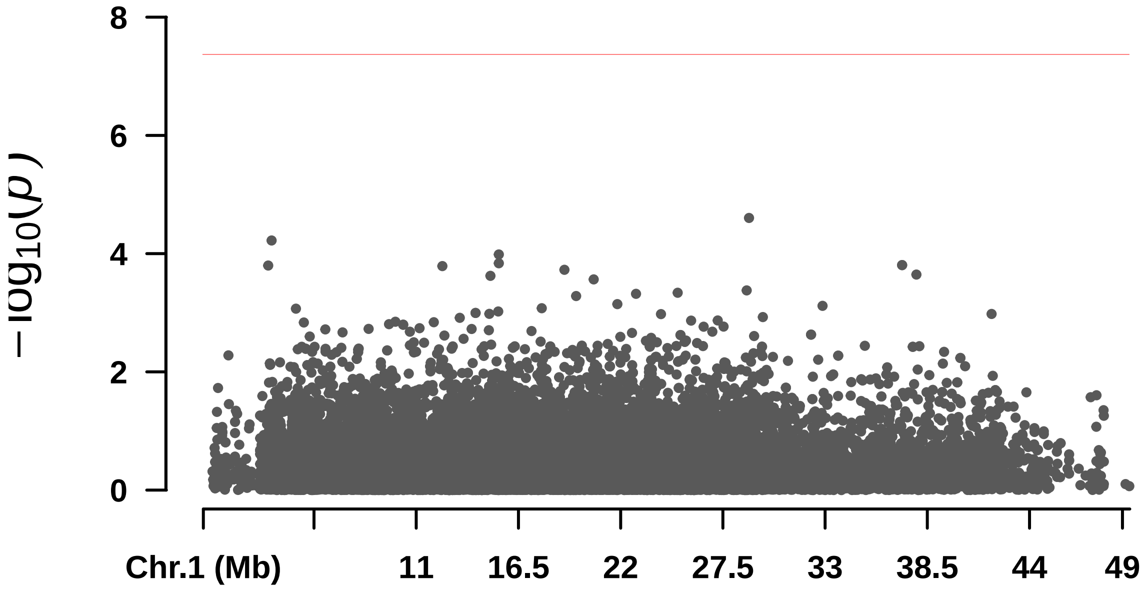
8
6
4
2
0
$-\log_{10}(p)$
Chr.1 (Mb)
11
16.5
22
27.5
33
38.5
44
49

#### Compri_pina

```
## $BLINK
## [1] 4.183538e-08 1.172187e-06 5.556791e-10 1.071399e-06 1.257862e-07
## [6] 8.912329e-14 6.782410e-10
##
## $FarmCPU
## numeric(0)
##
## $GLM
## numeric(0)
##
## $MLM
## numeric(0)
##
## $MLMM
## numeric(0)
```

```
##  Rectangular-Manhattan Plotting BLINK.
```

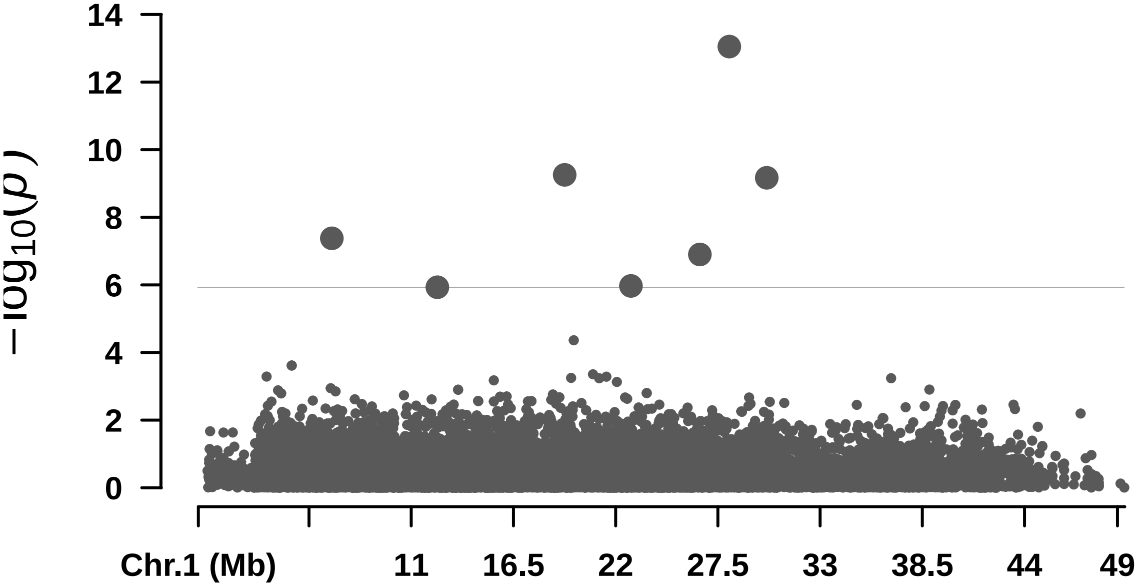


```
##  Rectangular-Manhattan Plotting FarmCPU.
```

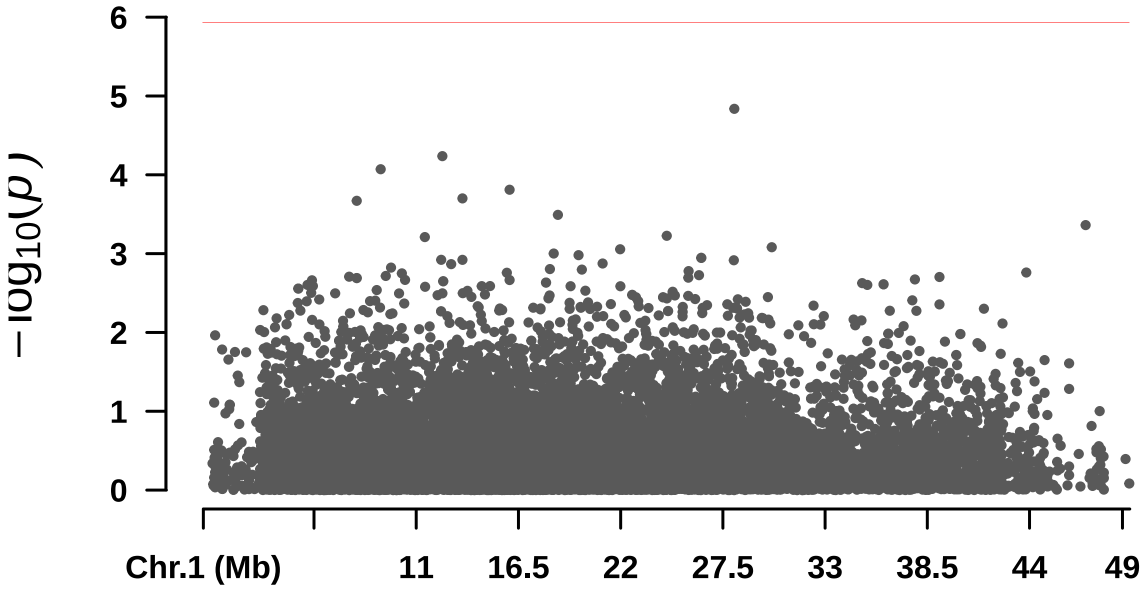


#### Rectangular-Manhattan Plotting GLM.

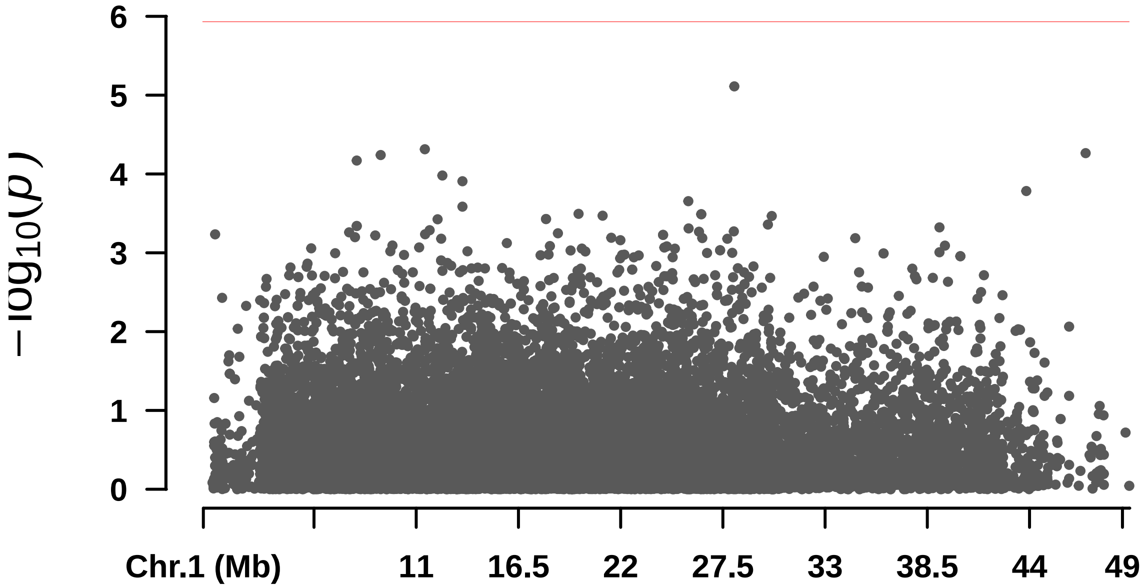


#### Rectangular-Manhattan Plotting MLM.

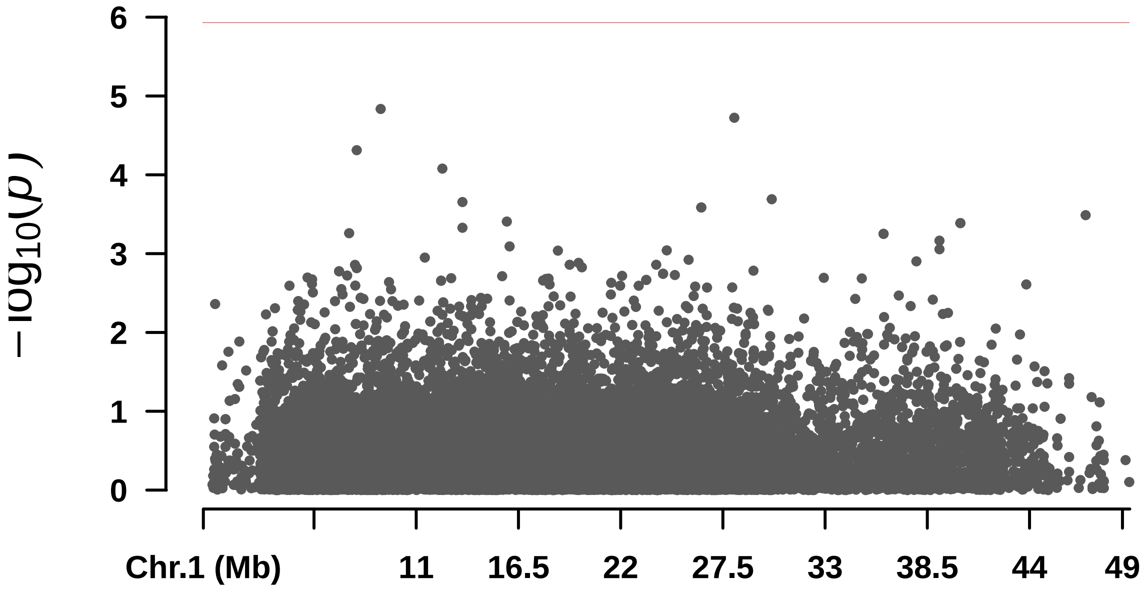


```
##  Rectangular-Manhattan Plotting MLMM.
```

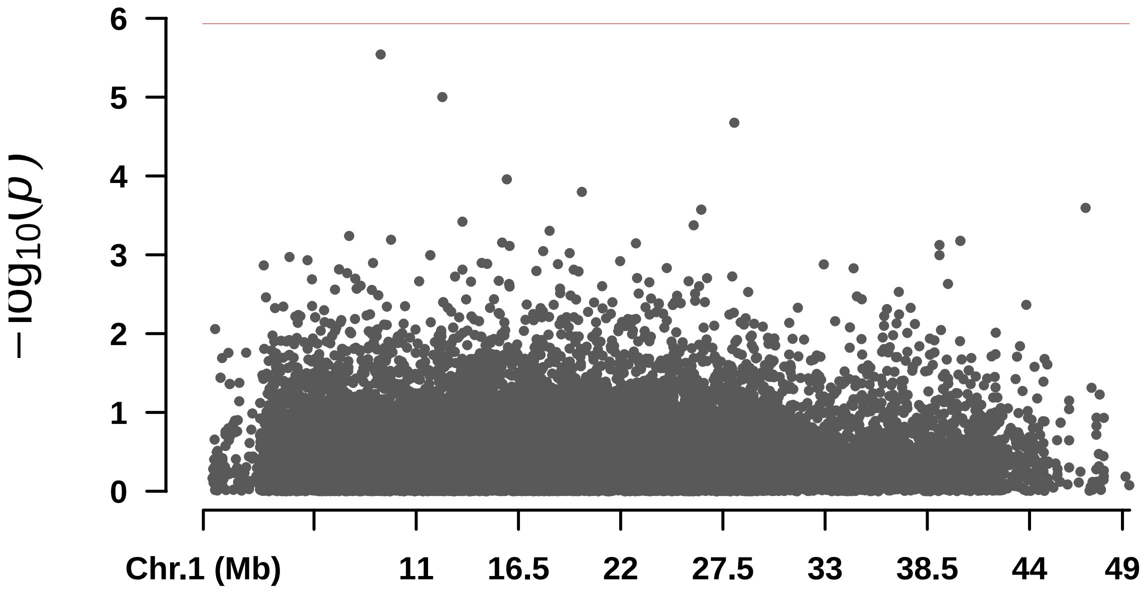

—

#### ## DAP

```
## $BLINK
## numeric(0)
##
## $FarmCPU
## numeric(0)
##
## $GLM
## [1] 5.958571e-06 4.512024e-06 6.536304e-06
##
## $MLM
## numeric(0)
##
## $MLMM
## [1] 2.683418e-07
```

```
##  Rectangular-Manhattan Plotting BLINK.
```

```
##  Rectangular-Manhattan Plotting FarmCPU.
```

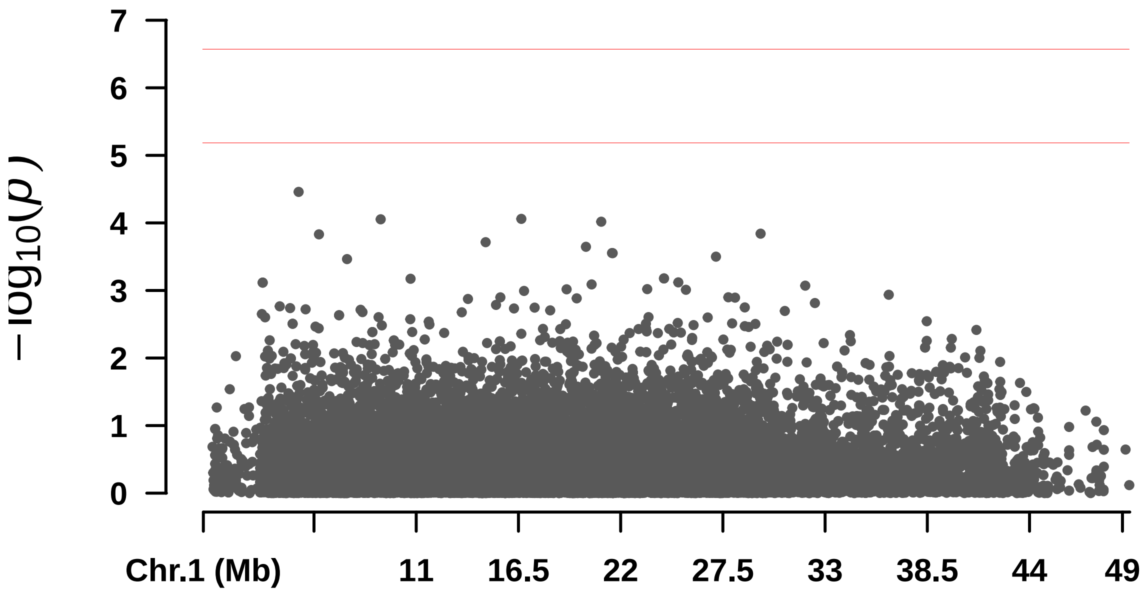


```
##  Rectangular-Manhattan Plotting GLM.
```

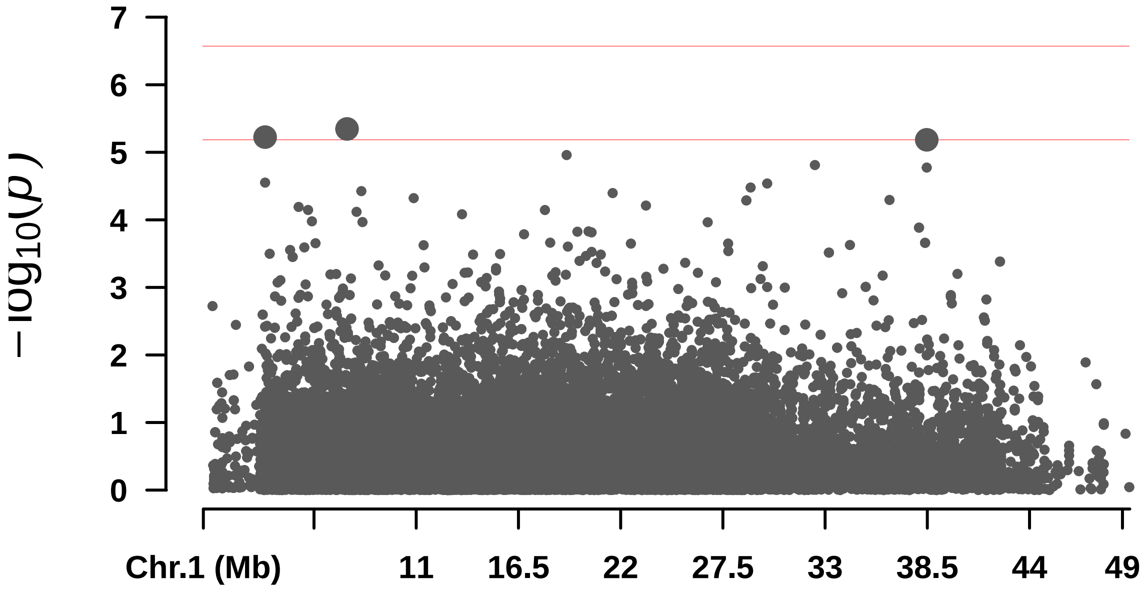


#### Rectangular-Manhattan Plotting MLM.

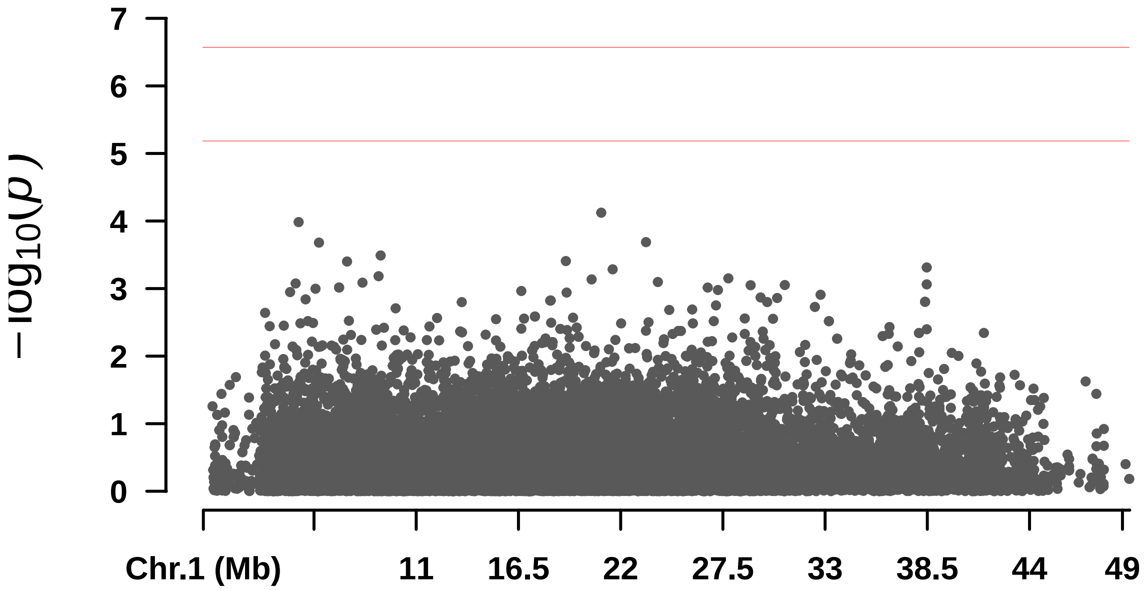


#### Rectangular-Manhattan Plotting MLMM.

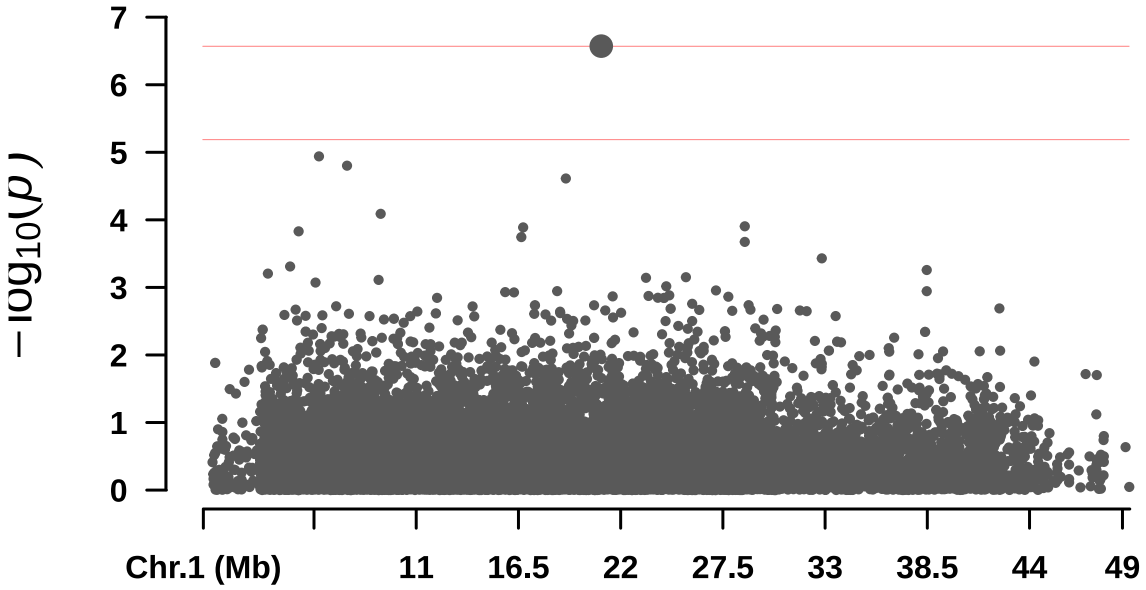

7
6
5
4
3
2
1
0
$-\log_{10}(p)$
Chr.1 (Mb)
11
16.5
22
27.5
33
38.5
44
49

—

#### ## F_Casca

```
## $BLINK
## [1] 6.763140e-07 6.477370e-06 8.437933e-07
##
## $FarmCPU
## numeric(0)
##
## $GLM
## numeric(0)
##
## $MLM
## numeric(0)
##
## $MLMM
## numeric(0)
```

```
## Rectangular-Manhattan Plotting BLINK.
```

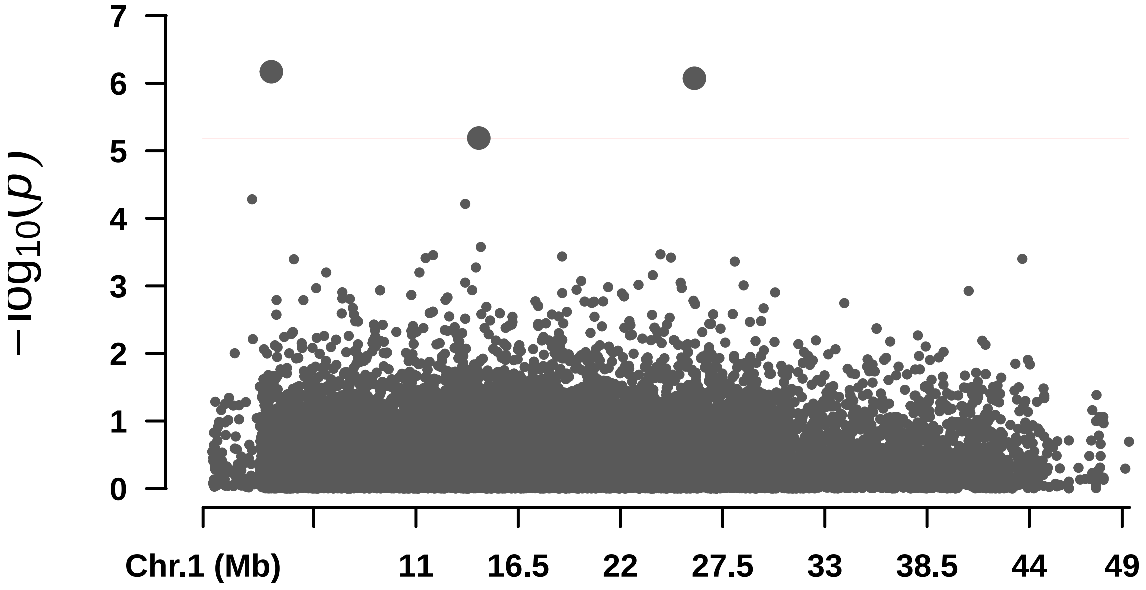


```
## Rectangular-Manhattan Plotting FarmCPU.
```

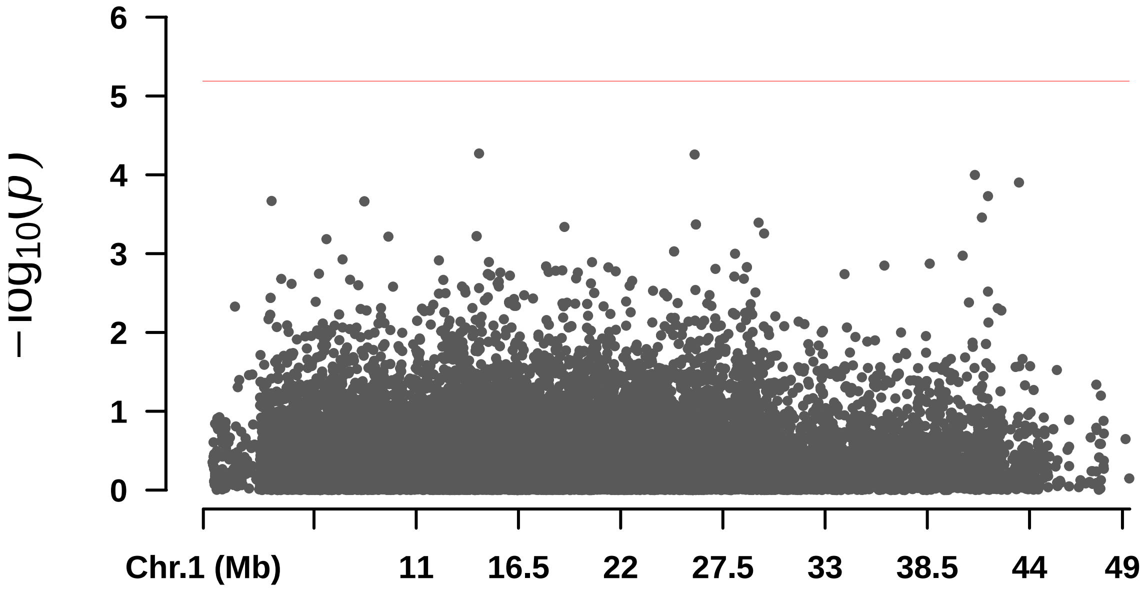


```
##  Rectangular-Manhattan Plotting GLM.
```

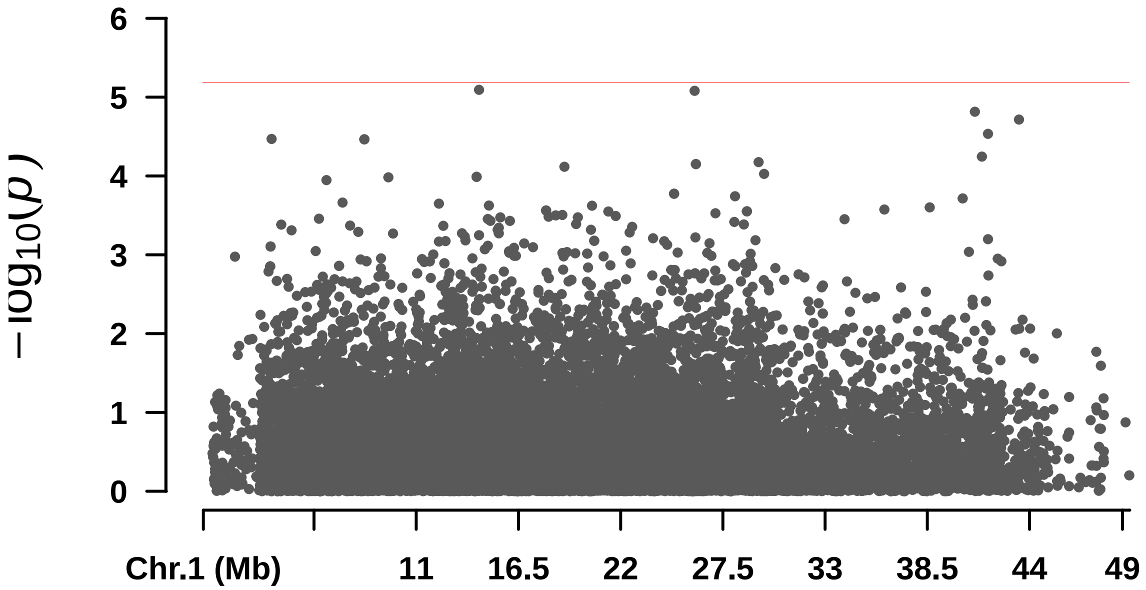


```
##  Rectangular-Manhattan Plotting MLM.
```

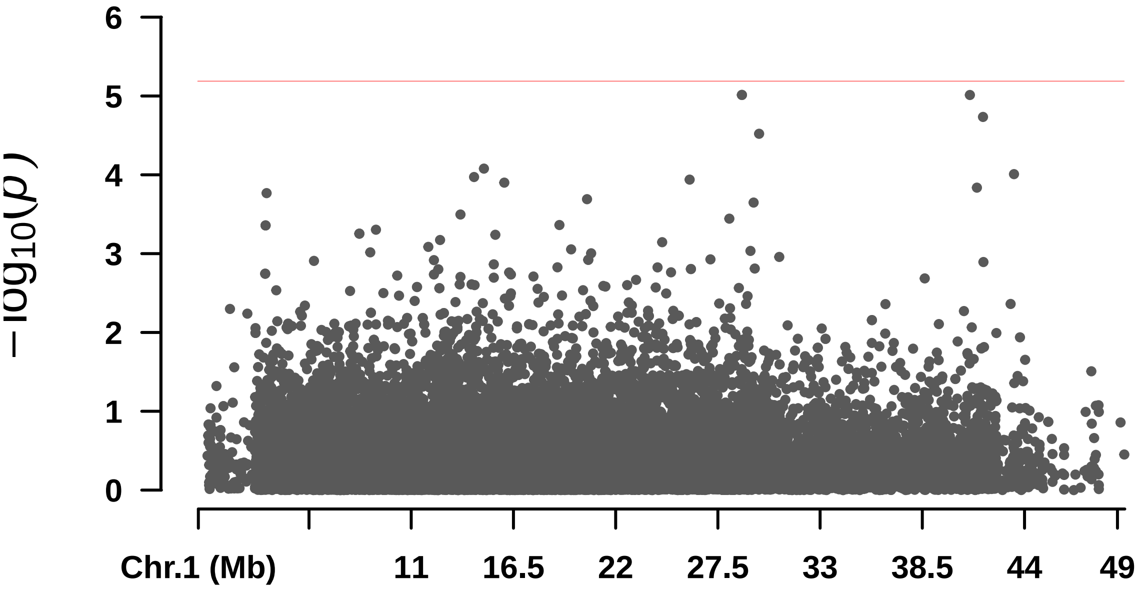


```
##  Rectangular-Manhattan Plotting MLMM.
```

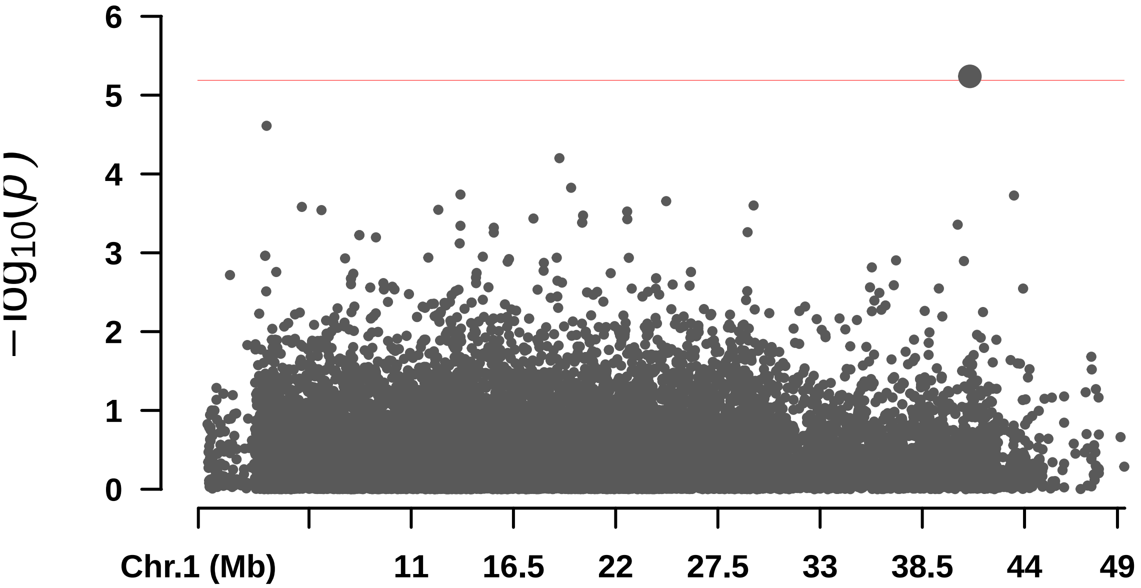

—

#### ## F_Endo

```
## $BLINK
## [1] 6.843718e-08 7.798951e-06 1.233460e-07 3.027799e-12
##
## $FarmCPU
## numeric(0)
##
## $GLM
## numeric(0)
##
## $MLM
## [1] 4.794505e-06 5.354354e-06 6.889056e-06
##
## $MLMM
## [1] 1.512983e-06
```

```
##  Rectangular-Manhattan Plotting BLINK.
```

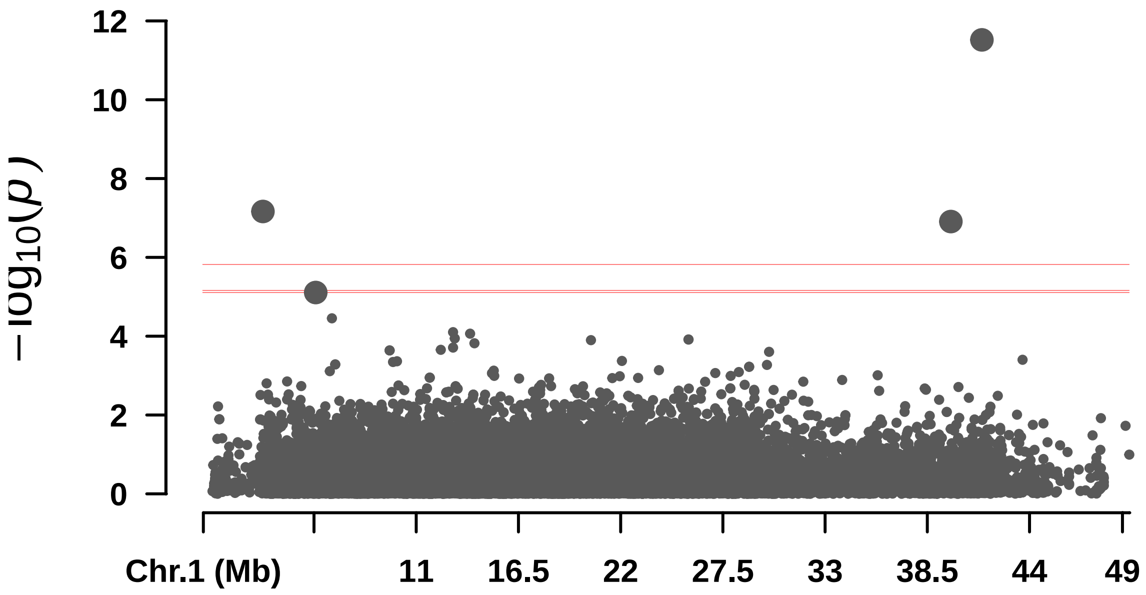


```
##  Rectangular-Manhattan Plotting FarmCPU.
```

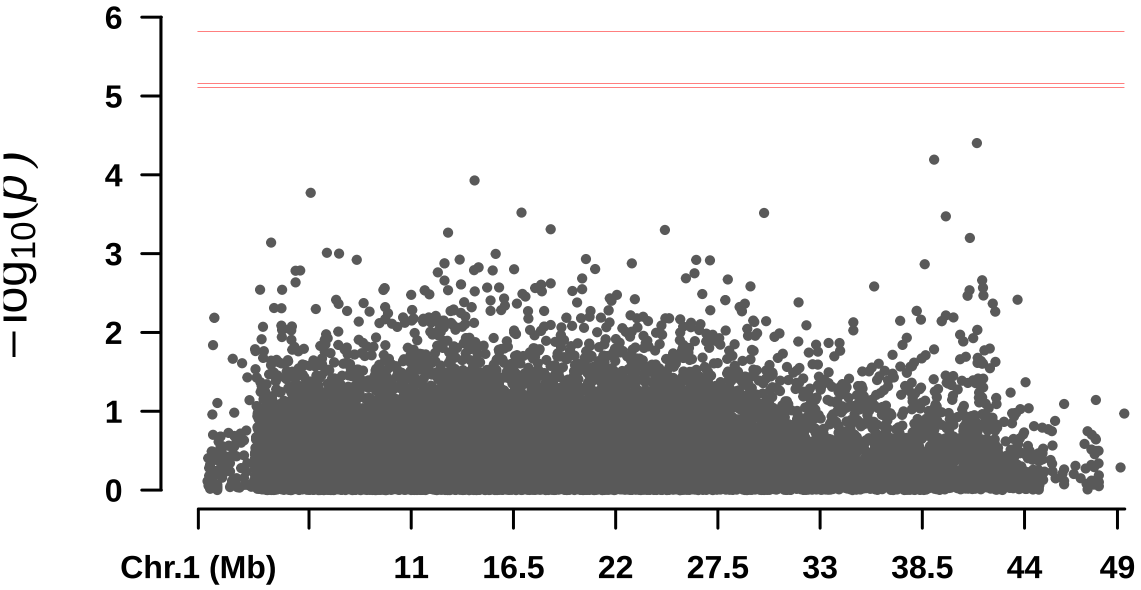


```
##  Rectangular-Manhattan Plotting GLM.
```

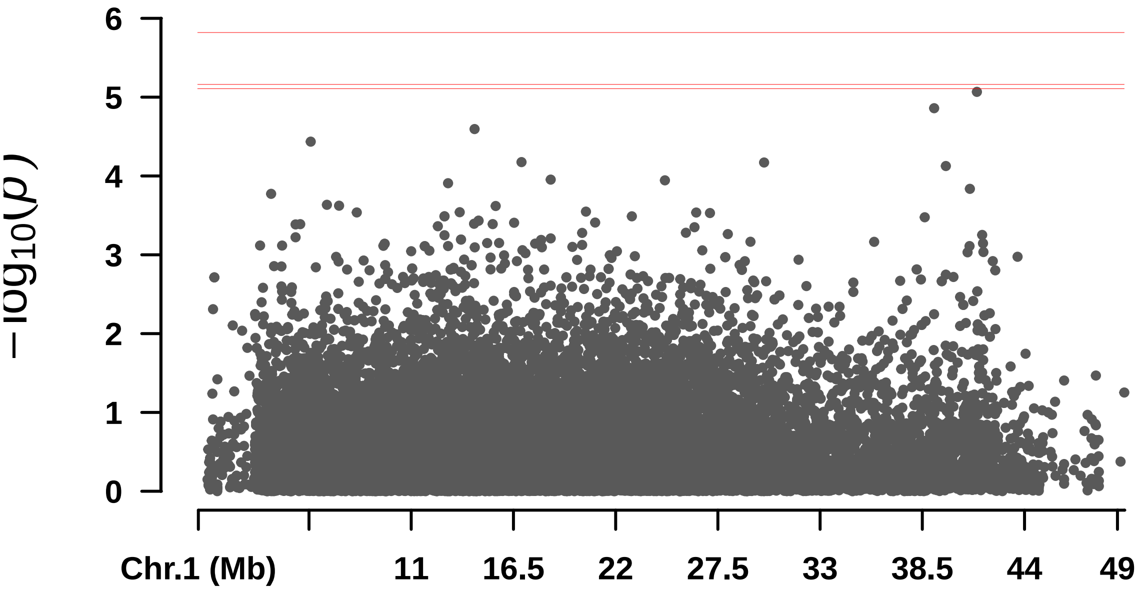


```
##  Rectangular-Manhattan Plotting MLM.
```

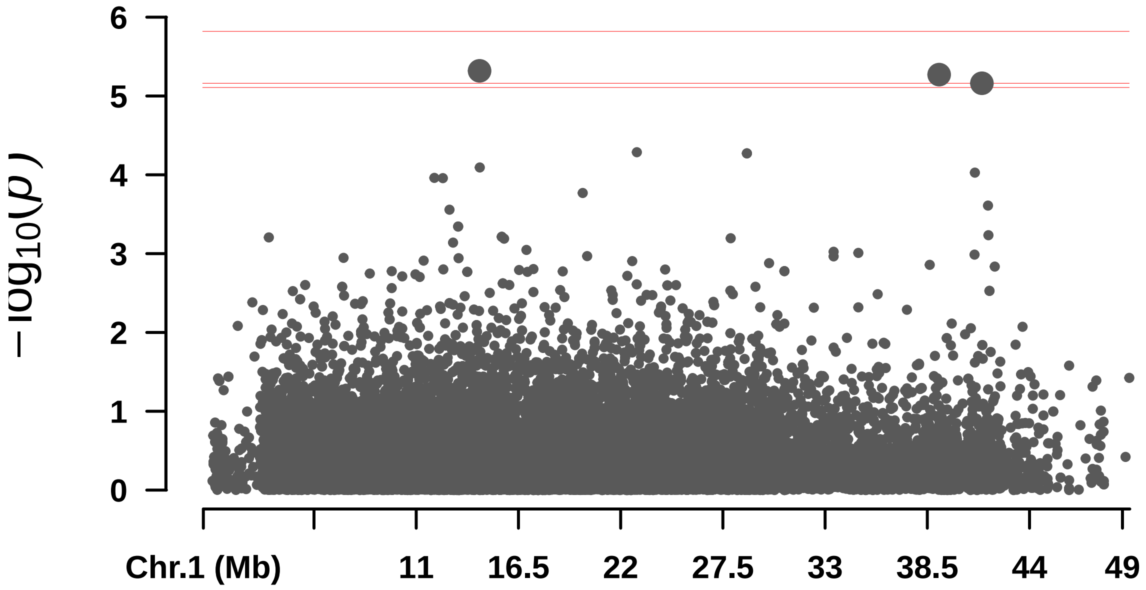


```
##  Rectangular-Manhattan Plotting MLMM.
```

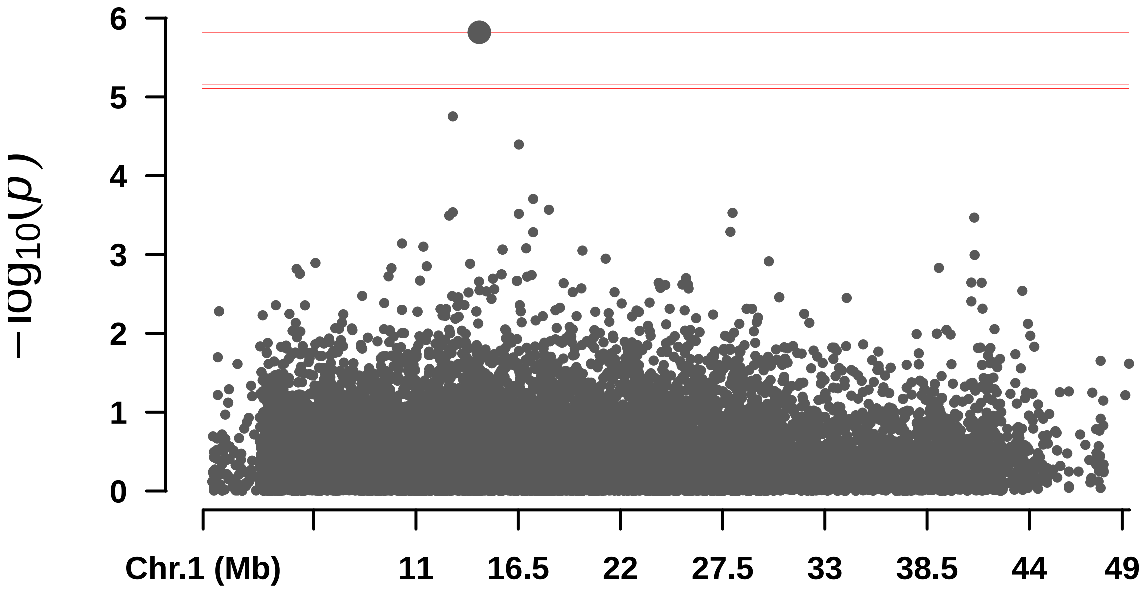

—

#### ## F__Polpa

```
## Warning in min(x[m <= 0.05]): no non-missing arguments to min; returning Inf
```

```
##        BLINK      FarmCPU          GLM          MLM         MLMM
## 5.239869e-13 6.671949e-17 1.293244e-06          Inf 1.920232e-07
```

```
##  Rectangular-Manhattan Plotting BLINK.
```

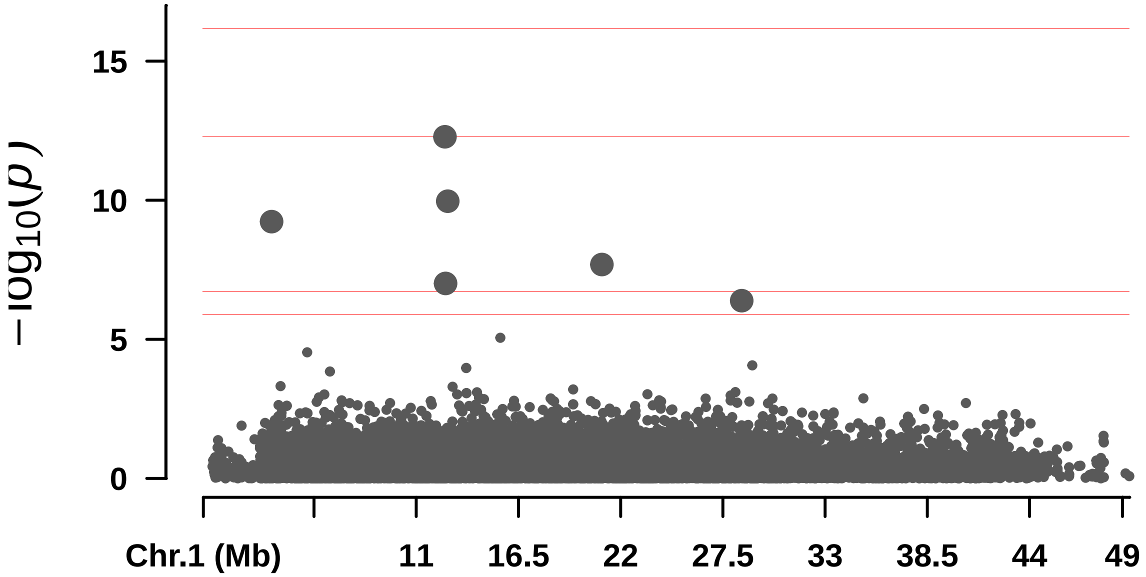


```
##  Rectangular-Manhattan Plotting FarmCPU.
```

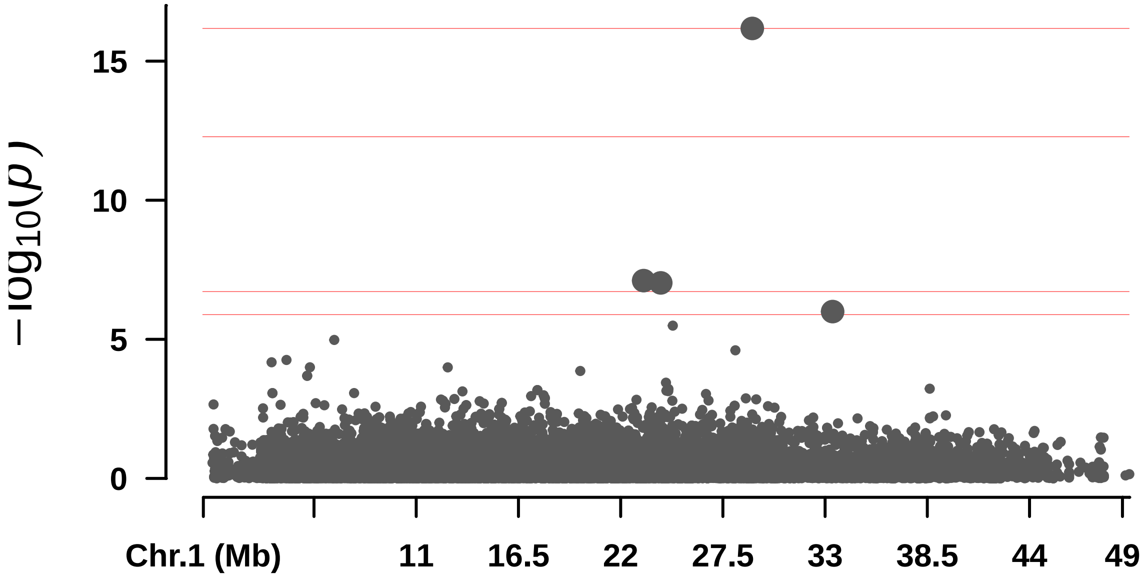


```
##  Rectangular-Manhattan Plotting GLM.
```

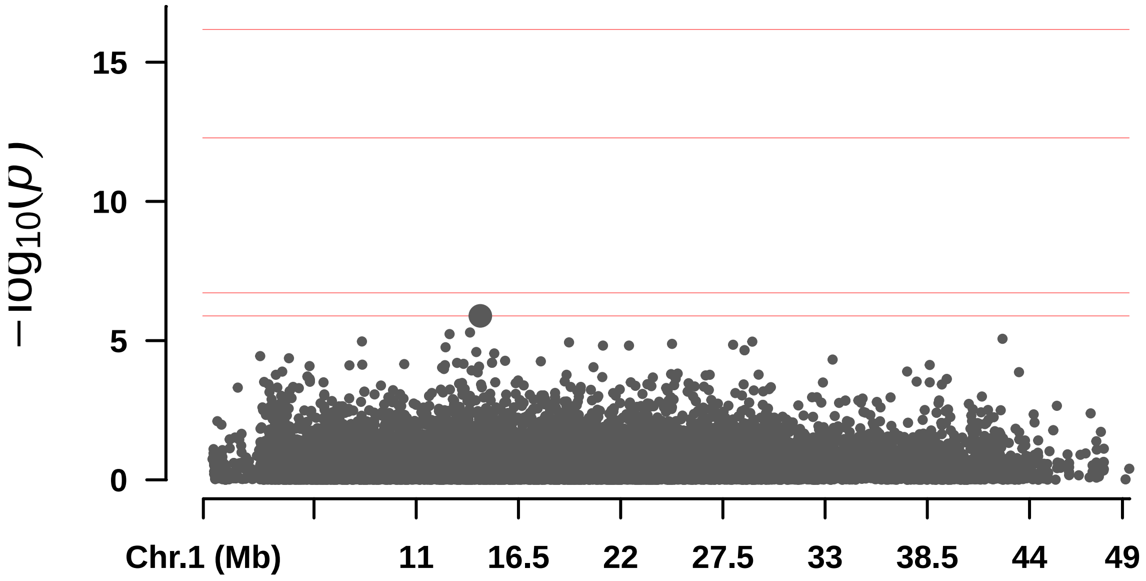


####  Rectangular-Manhattan Plotting MLM.

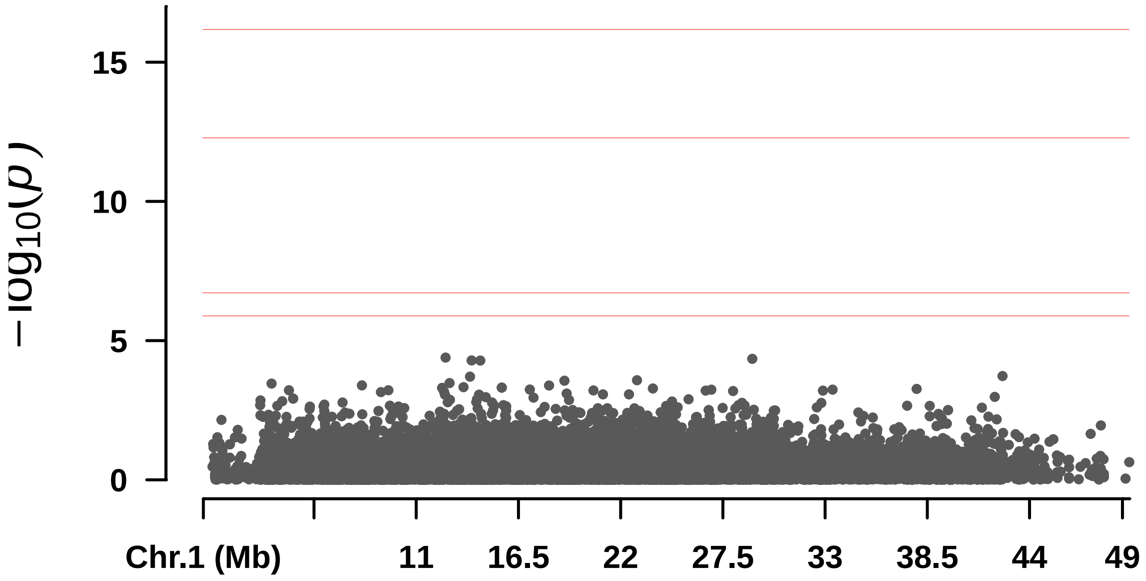


####  Rectangular-Manhattan Plotting MLMM.

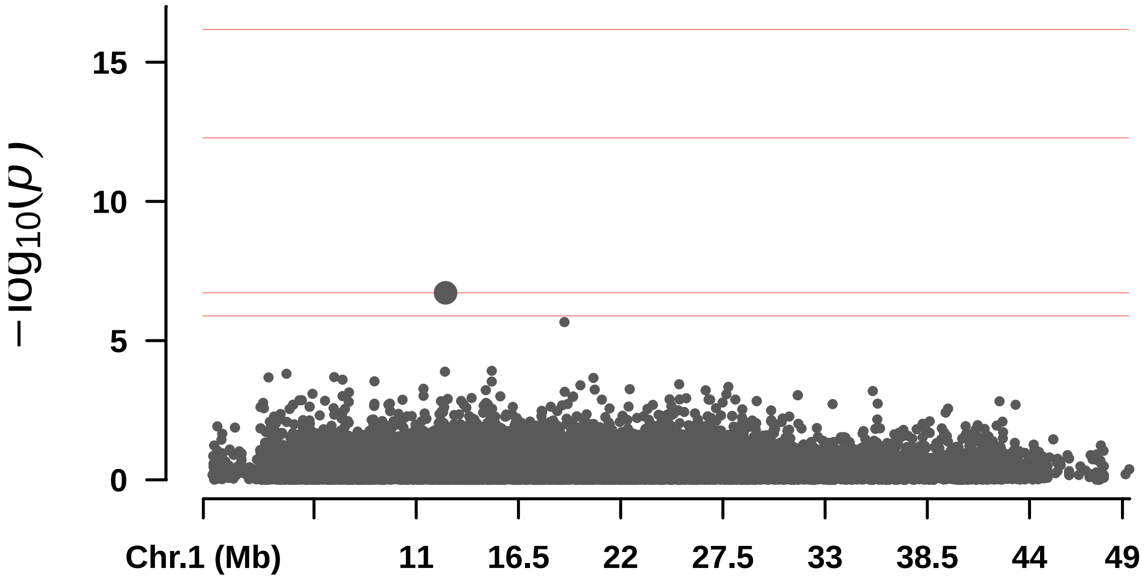
−log10(p)
15
10
5
0
Chr.1 (Mb)
11
16.5
22
27.5
33
38.5
44
49

—

#### ## F_Sem

```
## Warning in max(x[m <= 0.05]): no non-missing arguments to max; returning -Inf

## Warning in max(x[m <= 0.05]): no non-missing arguments to max; returning -Inf

## Warning in max(x[m <= 0.05]): no non-missing arguments to max; returning -Inf

##        BLINK      FarmCPU          GLM          MLM         MLMM
## 1.549935e-06         -Inf 1.040112e-04         -Inf         -Inf

##  QQ Plotting BLINK.
```

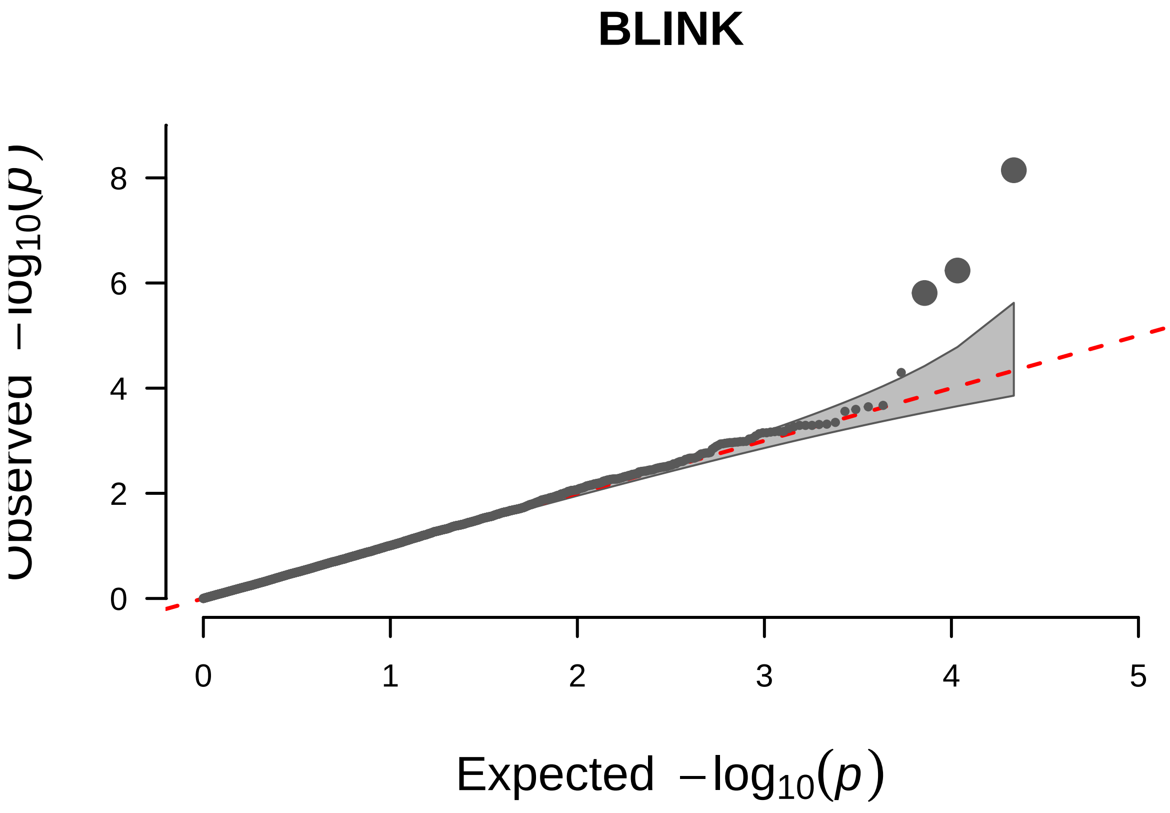


```
##  QQ Plotting FarmCPU.
```

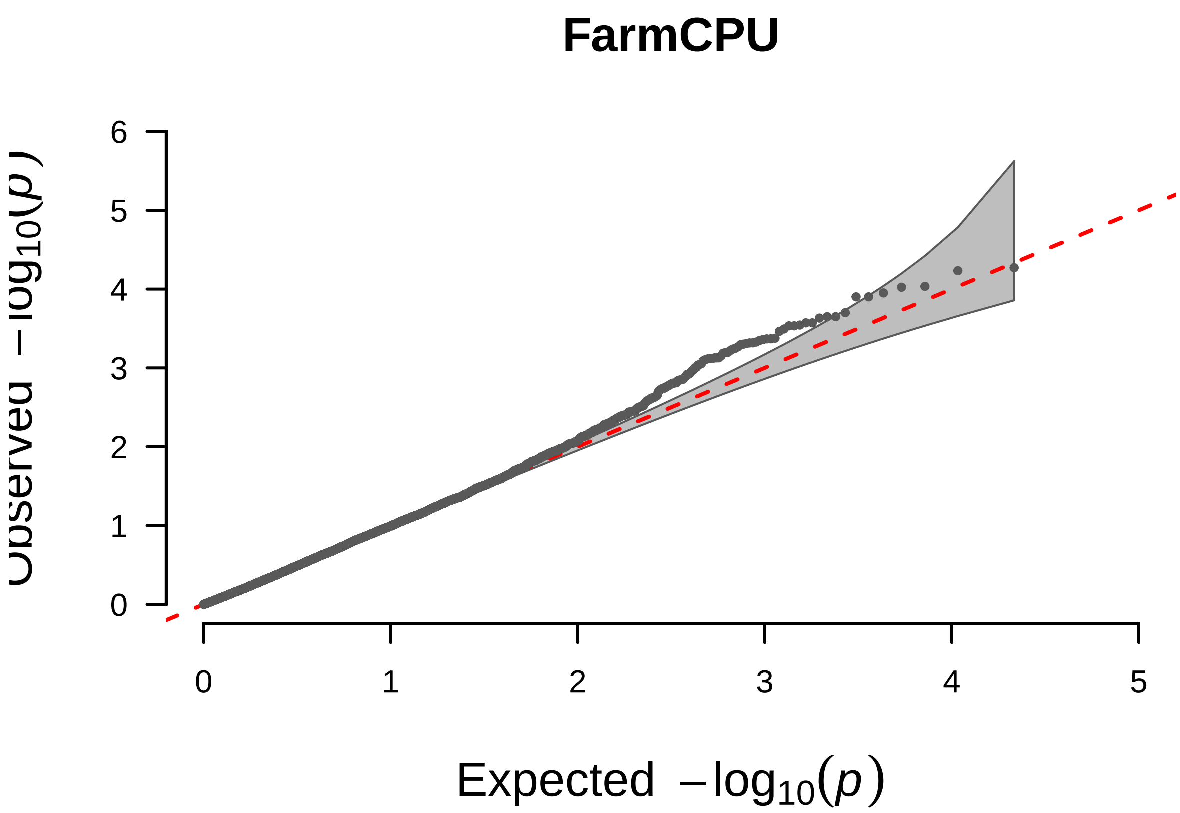


#### QQ Plotting GLM.

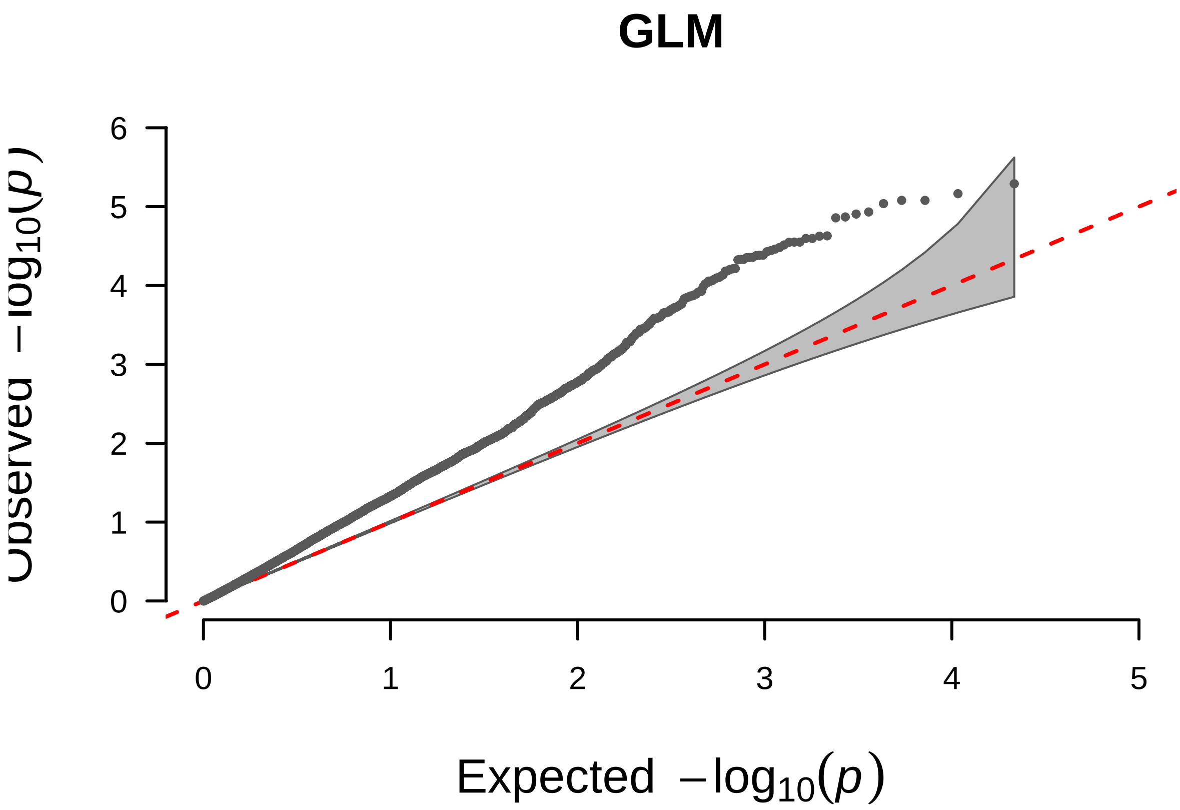


#### QQ Plotting MLM.

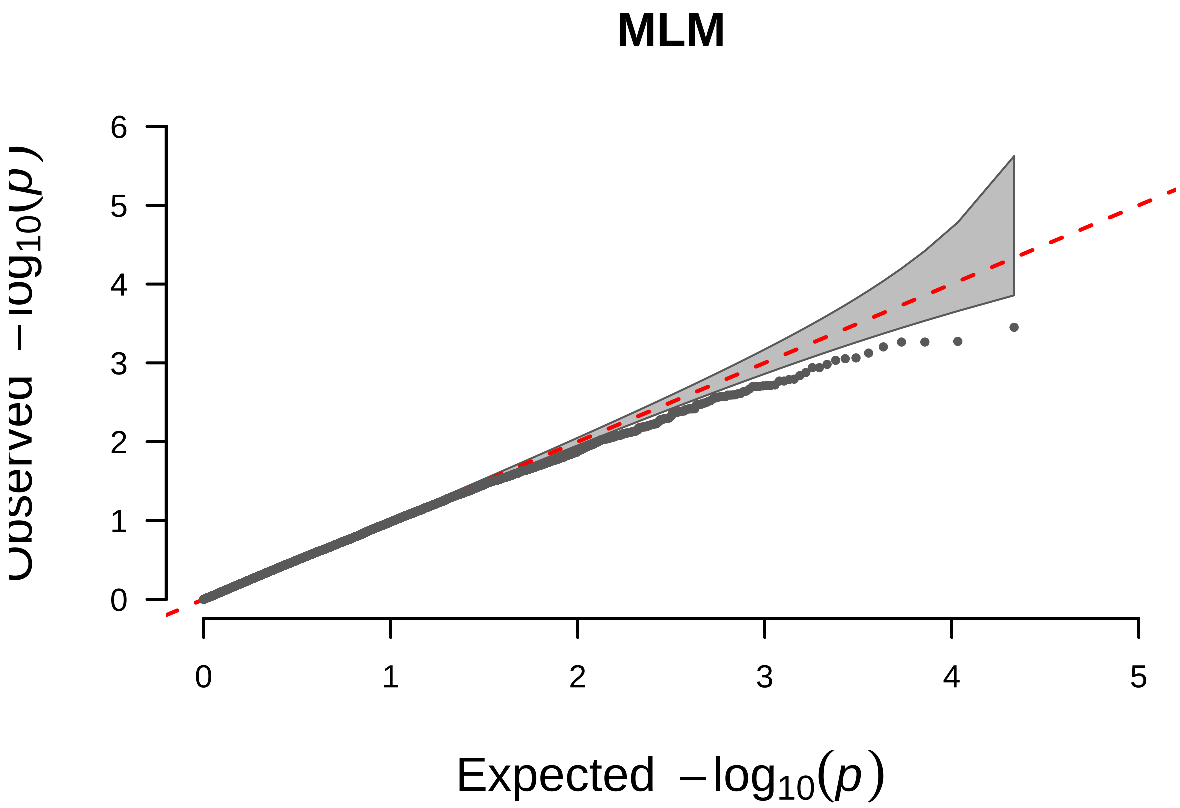


#### QQ Plotting MLMM.

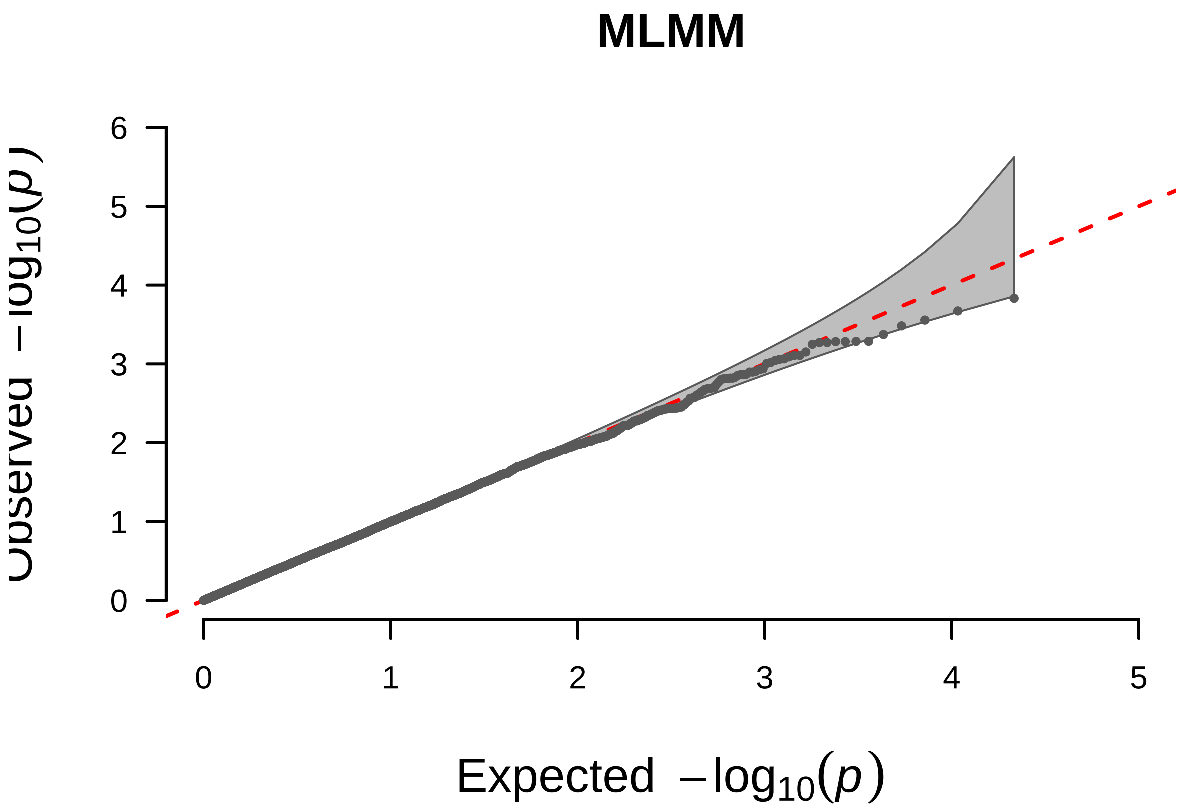

—

## F_T

```
## Warning in max(x[m <= 0.05]): no non-missing arguments to max; returning -Inf

## Warning in max(x[m <= 0.05]): no non-missing arguments to max; returning -Inf

##        BLINK      FarmCPU          GLM          MLM         MLMM
##         -Inf 4.261708e-06 2.263137e-06         -Inf 6.003165e-08

##  Rectangular-Manhattan Plotting BLINK.
```

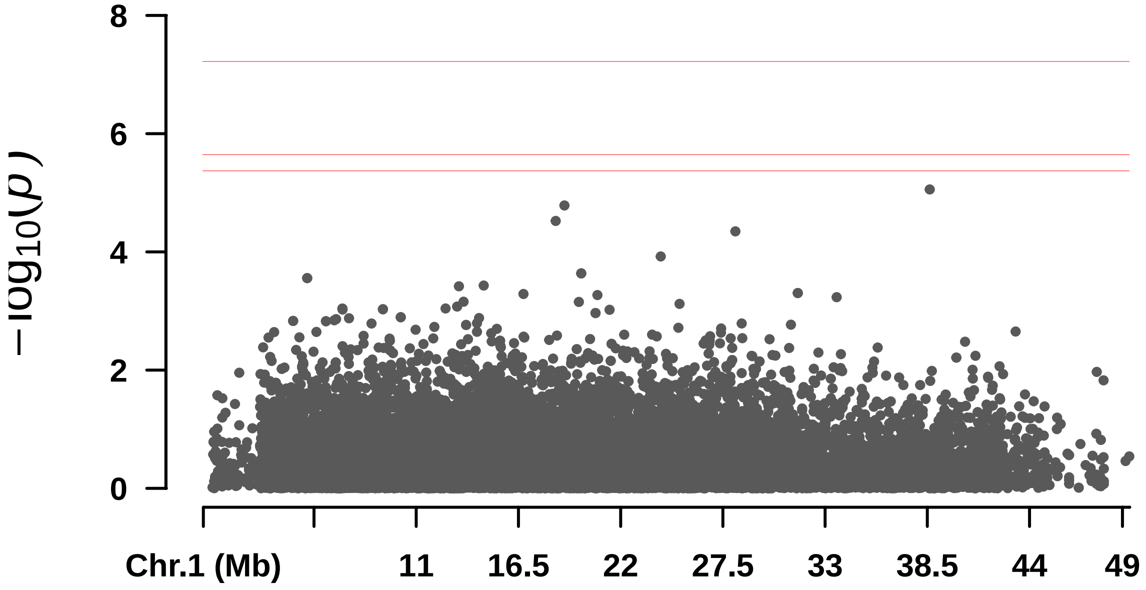


```
##  Rectangular-Manhattan Plotting FarmCPU.
```

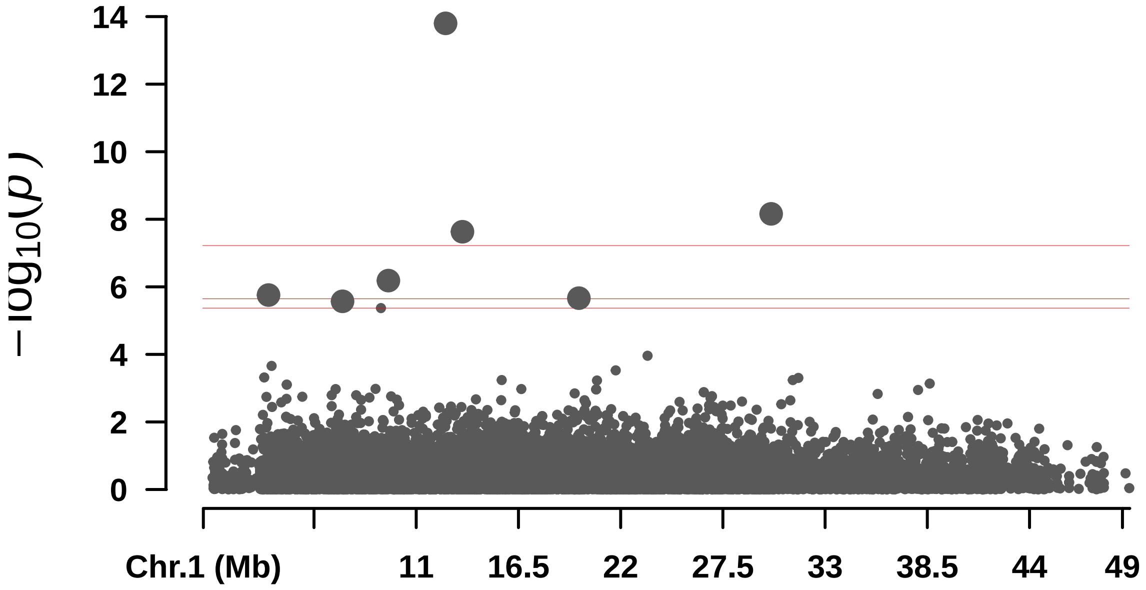


```
##  Rectangular-Manhattan Plotting GLM.
```

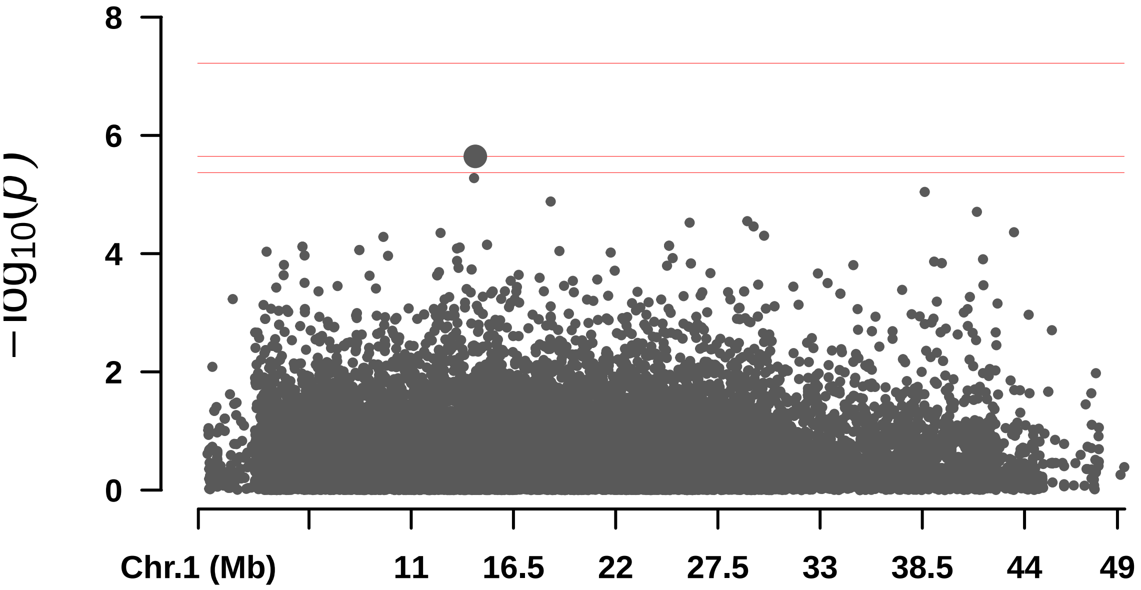


```
##  Rectangular-Manhattan Plotting MLM.
```

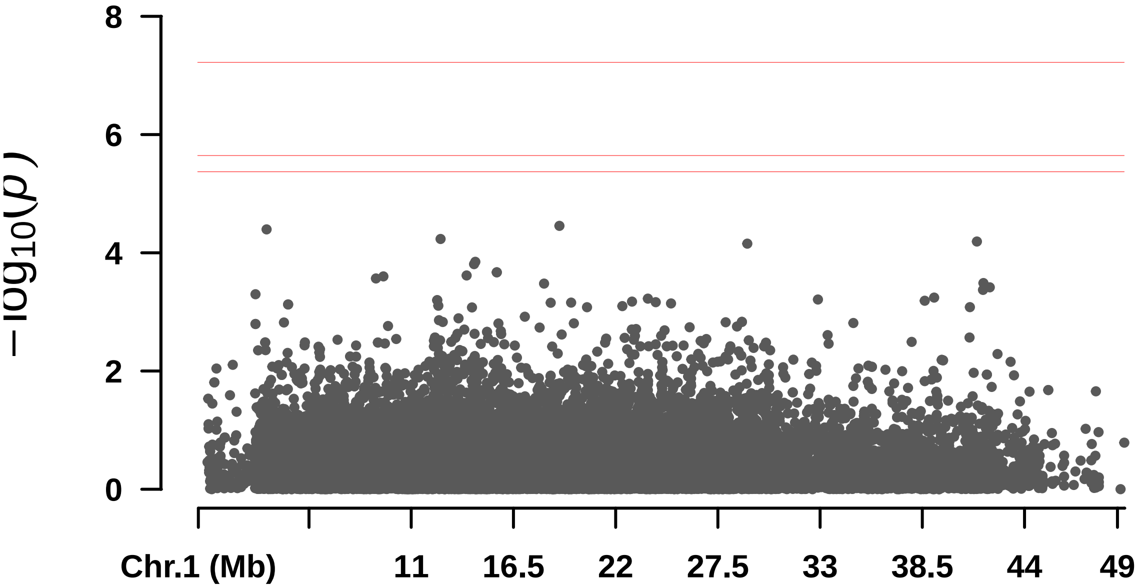


```
##  Rectangular-Manhattan Plotting MLMM.
```

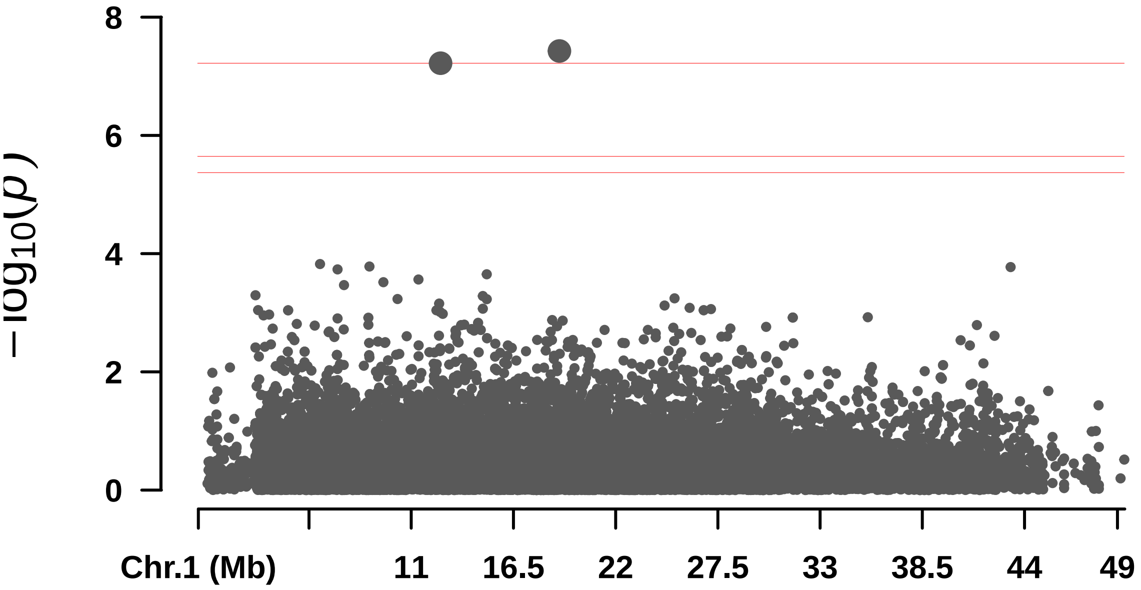
8
6
4
2
0
−log10(p)
Chr.1 (Mb)
11
16.5
22
27.5
33
38.5
44
49

—

#### Largura_pina

```
## Warning in max(x[m <= 0.05]): no non-missing arguments to max; returning -Inf

## Warning in max(x[m <= 0.05]): no non-missing arguments to max; returning -Inf

## Warning in max(x[m <= 0.05]): no non-missing arguments to max; returning -Inf

## Warning in max(x[m <= 0.05]): no non-missing arguments to max; returning -Inf

##         BLINK      FarmCPU          GLM          MLM         MLMM
## 1.950863e-07          -Inf         -Inf         -Inf         -Inf

##  Rectangular-Manhattan Plotting BLINK.
```

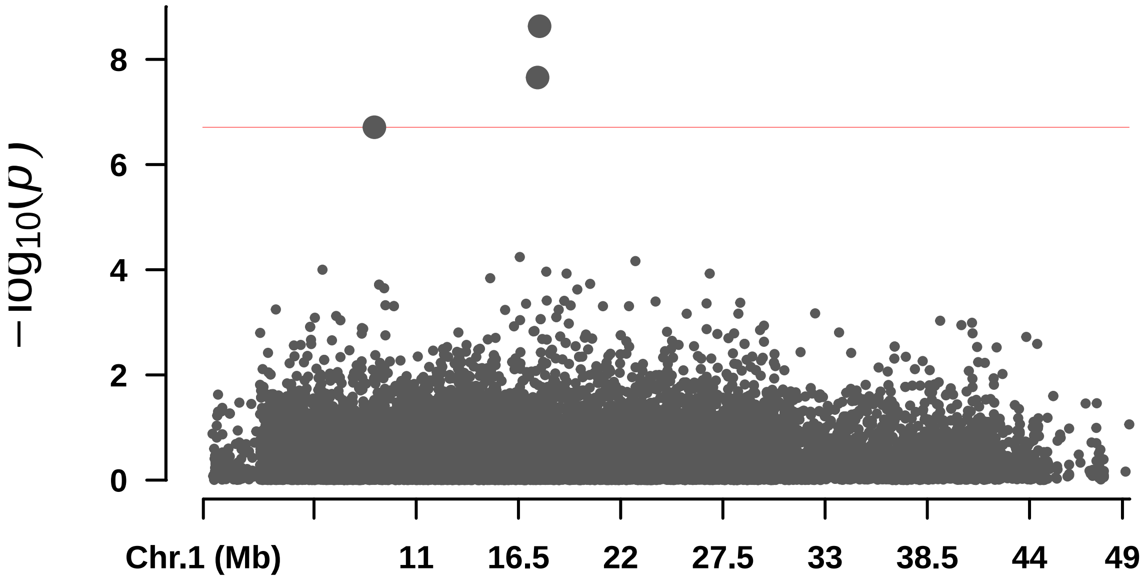


```
##  Rectangular-Manhattan Plotting FarmCPU.
```

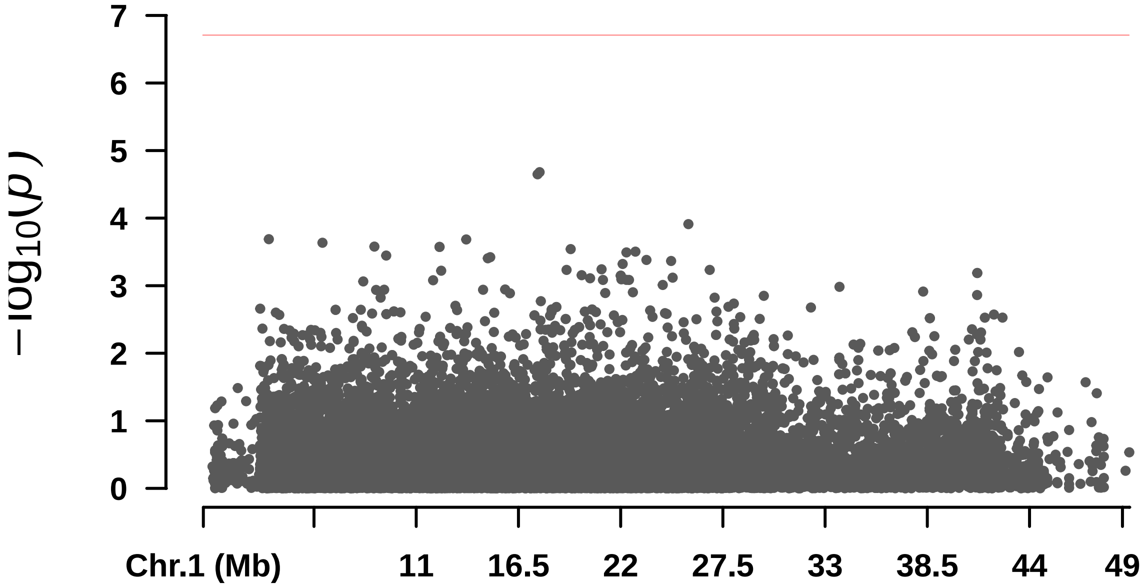

```
##  Rectangular-Manhattan Plotting GLM.
```

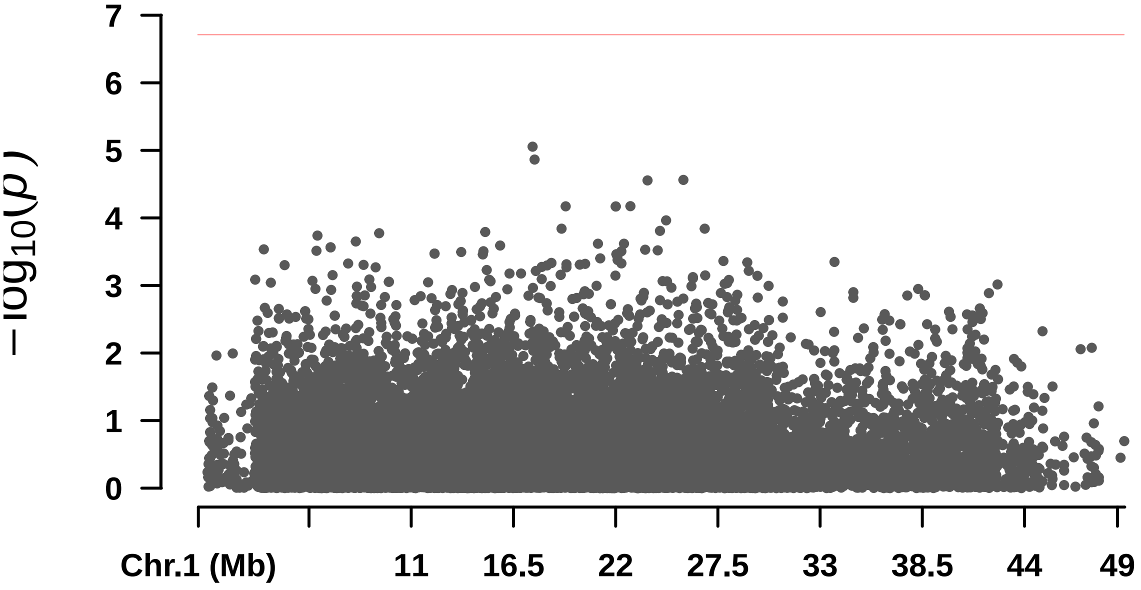


```
##  Rectangular-Manhattan Plotting MLM.
```

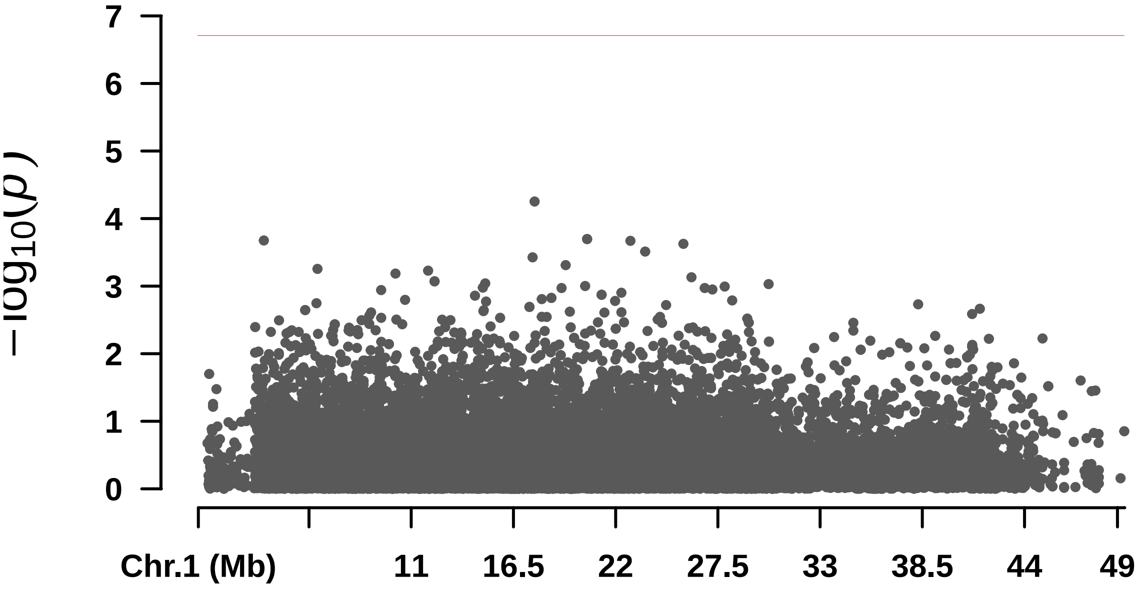


```
##  Rectangular-Manhattan Plotting MLMM.
```

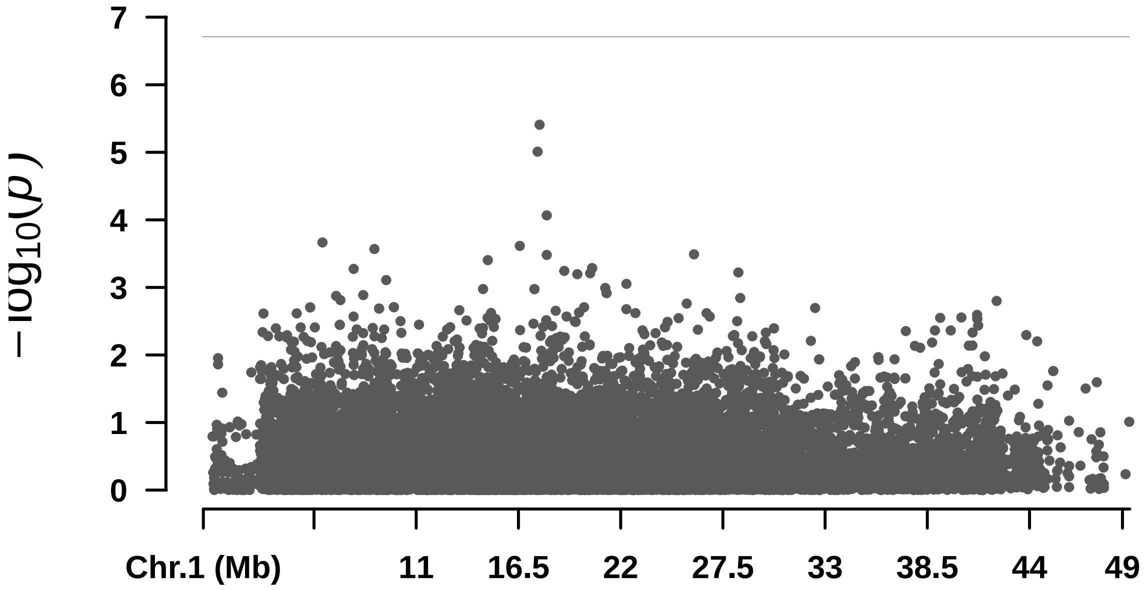
−log10(p)
7
6
5
4
3
2
1
0
Chr.1 (Mb)
11
16.5
22
27.5
33
38.5
44
49

#### Oil_content

```
## Warning in max(x[m <= 0.05]): no non-missing arguments to max; returning -Inf

## Warning in max(x[m <= 0.05]): no non-missing arguments to max; returning -Inf

##         BLINK      FarmCPU          GLM          MLM         MLMM
## 5.243642e-07 4.089495e-06 1.590756e-05         -Inf         -Inf

##  Rectangular-Manhattan Plotting BLINK.
```

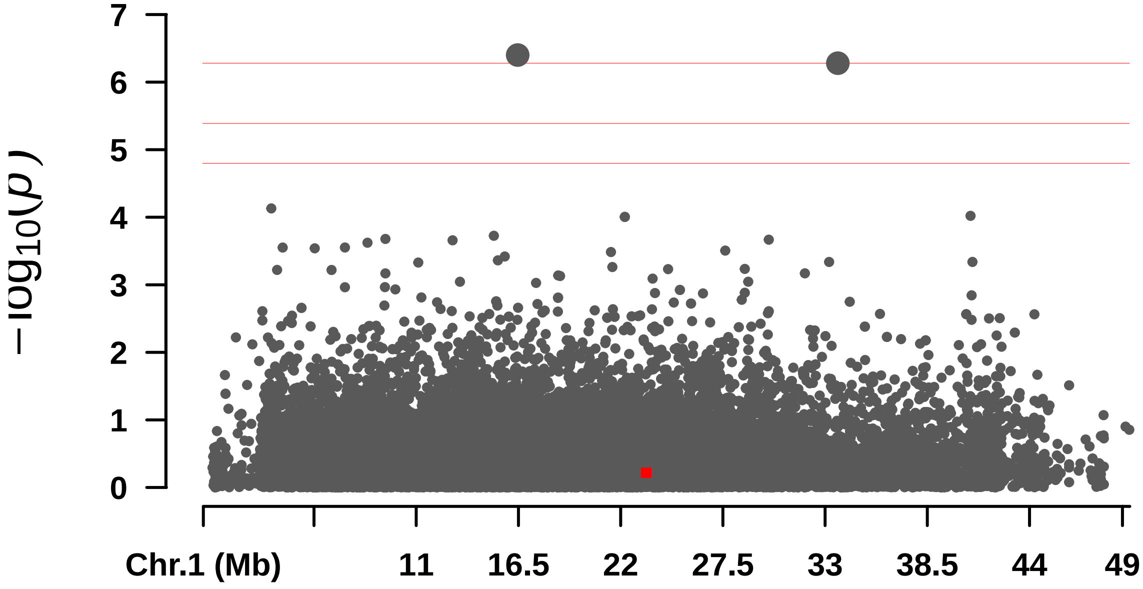


```
##  Rectangular-Manhattan Plotting FarmCPU.
```

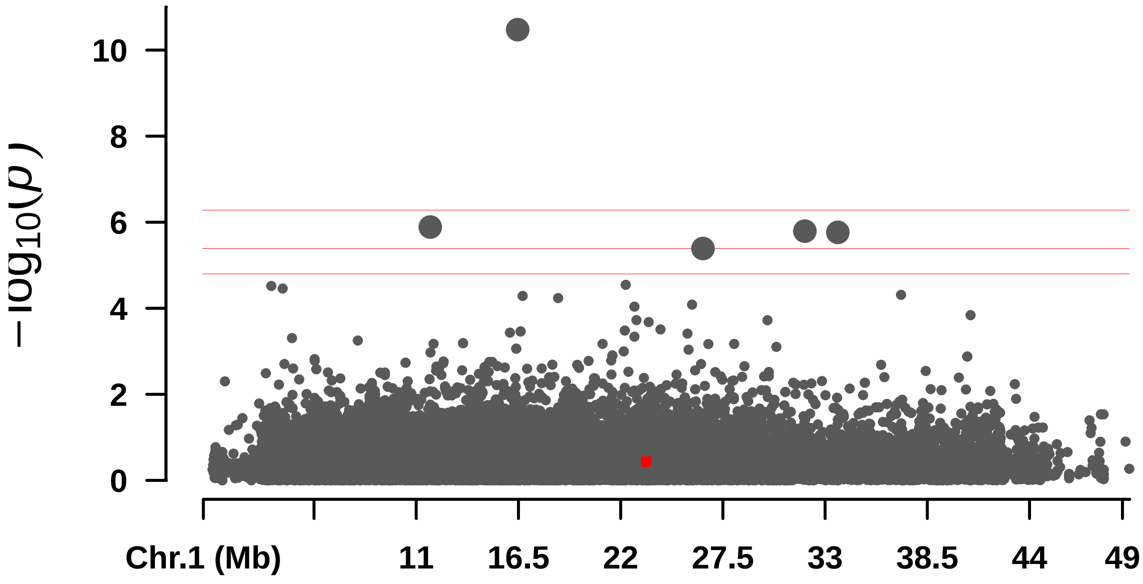


```
##  Rectangular-Manhattan Plotting GLM.
```

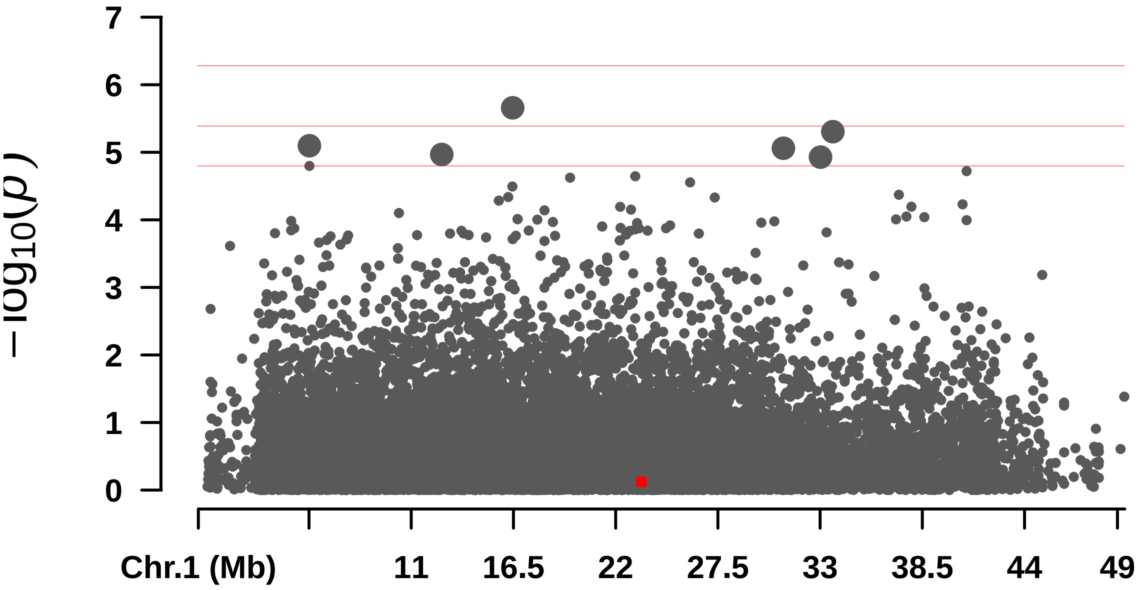


```
##  Rectangular-Manhattan Plotting MLM.
```

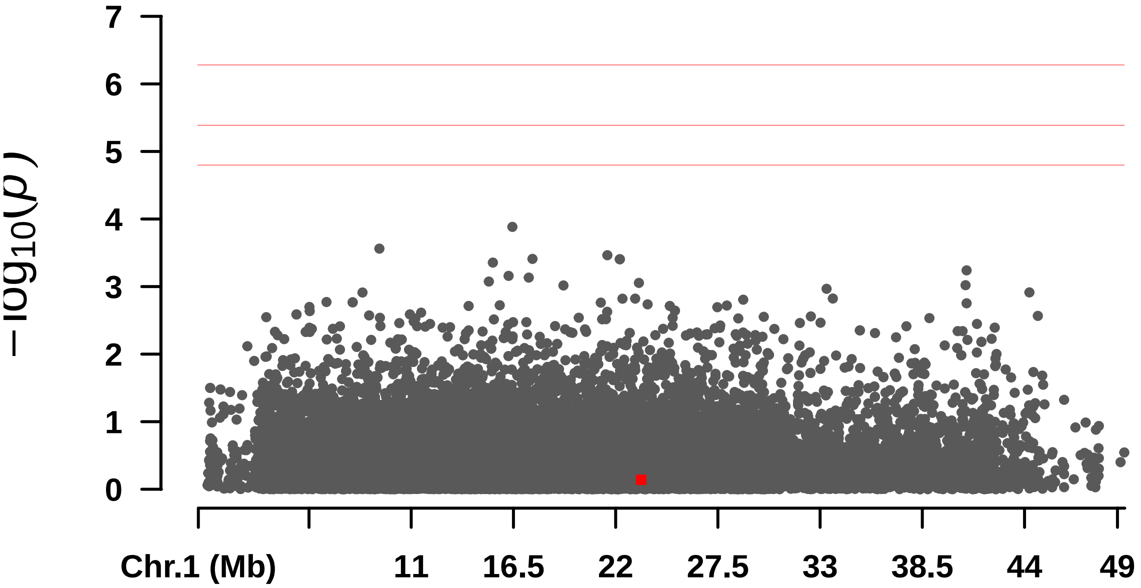


```
##  Rectangular-Manhattan Plotting MLMM.
```

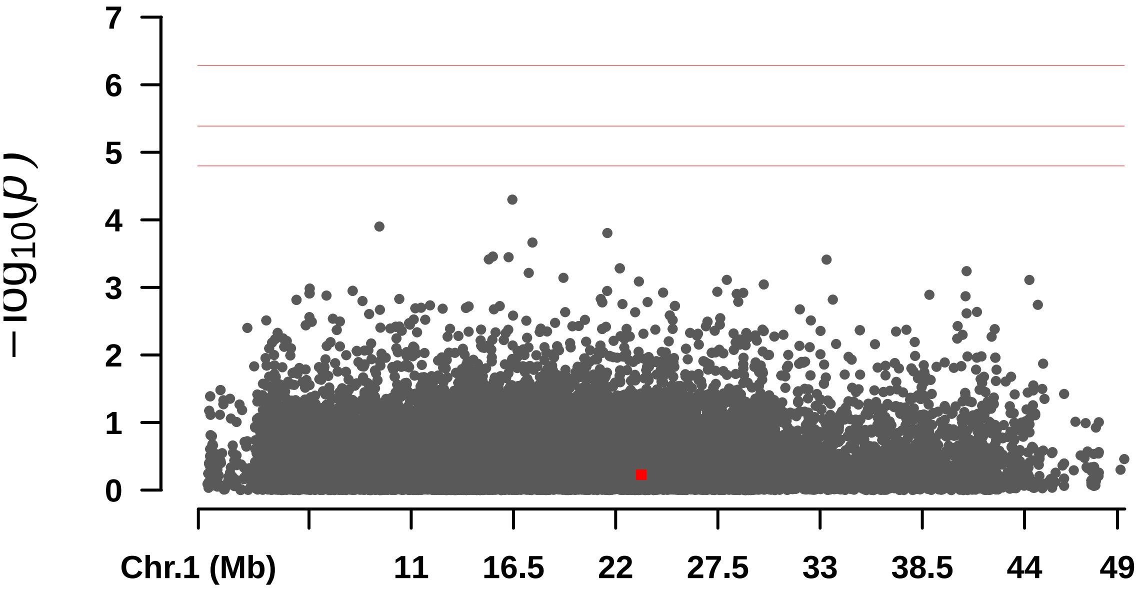
7
6
5
4
3
2
1
0
−log10(p)
Chr.1 (Mb)
11
16.5
22
27.5
33
38.5
44
49

—

#### S_Casca

```
## Warning in max(x[m <= 0.05]): no non-missing arguments to max; returning -Inf
```

```
##         BLINK      FarmCPU          GLM          MLM         MLMM
## 1.552968e-07 7.870854e-06 5.476808e-06         -Inf 4.569461e-06
```

```
##  Rectangular-Manhattan Plotting BLINK.
```

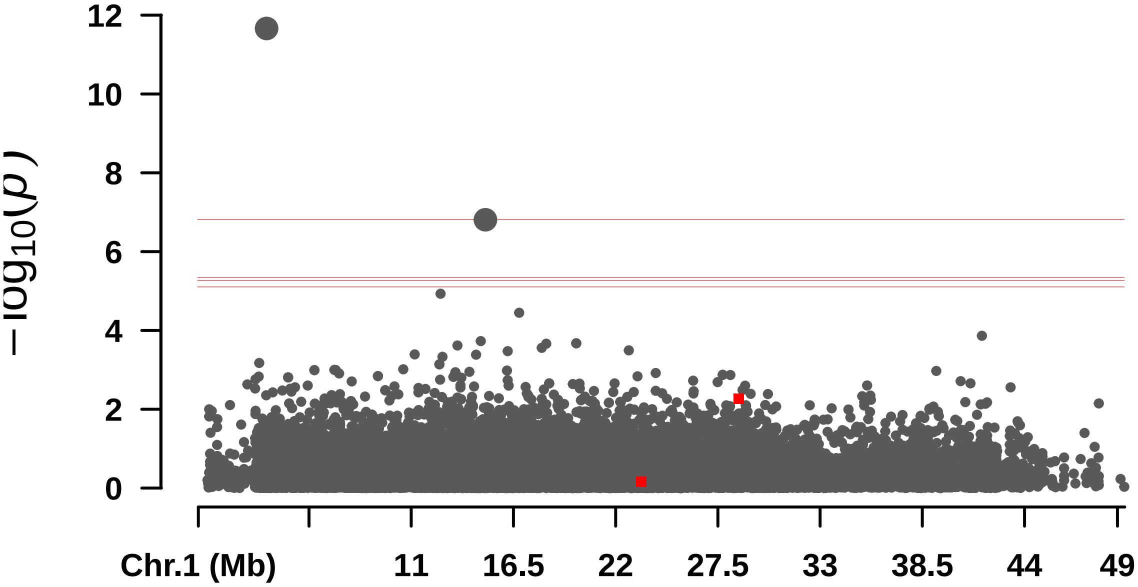


```
##  Rectangular-Manhattan Plotting FarmCPU.
```

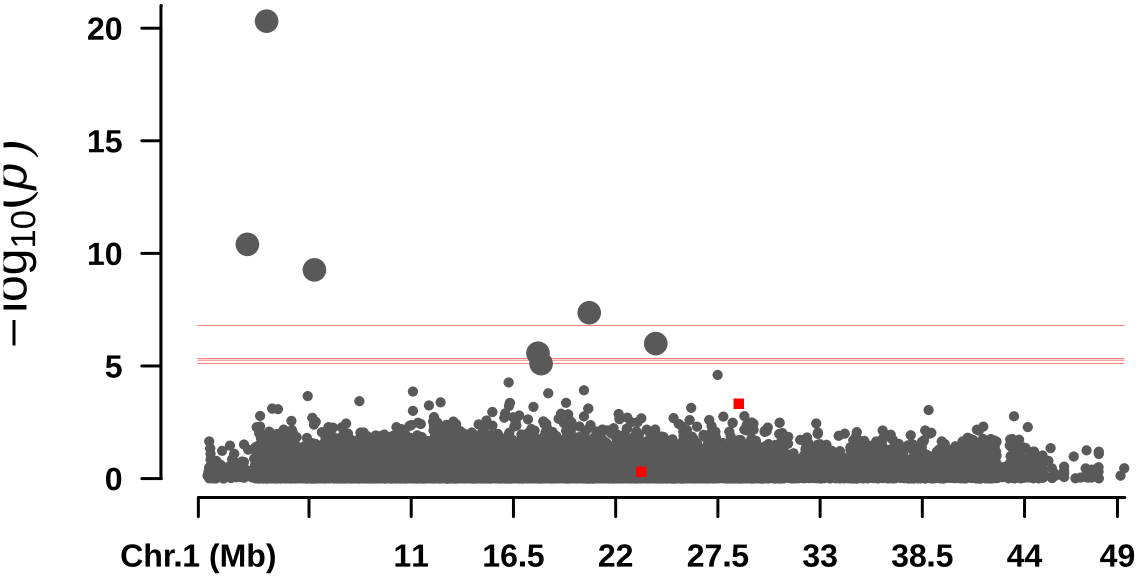


```
##  Rectangular-Manhattan Plotting GLM.
```

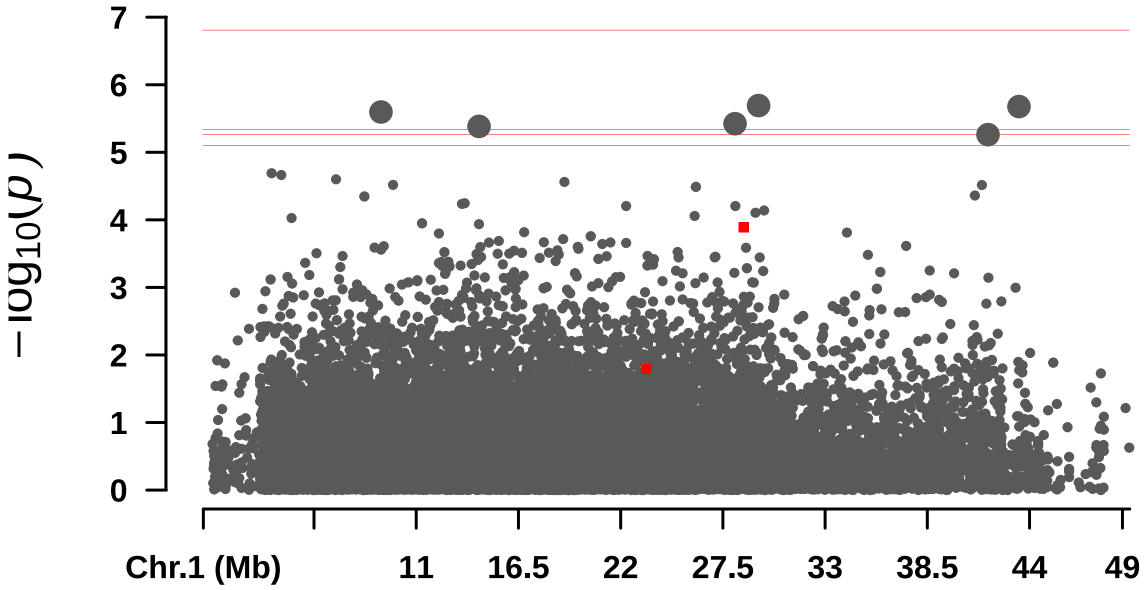


#### Rectangular-Manhattan Plotting MLM.

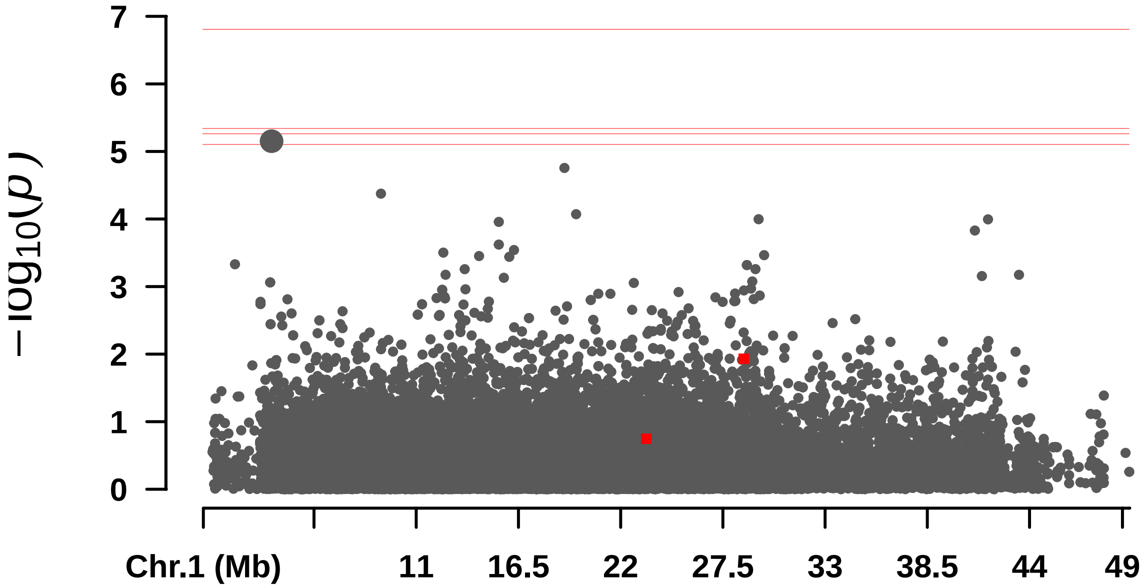


#### Rectangular-Manhattan Plotting MLMM.

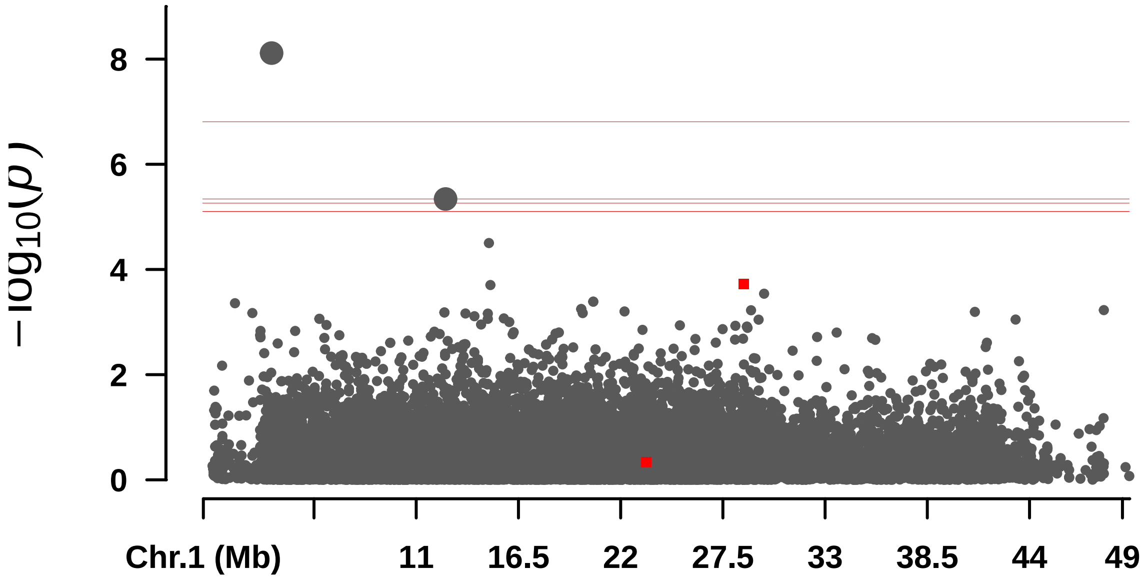

$-\log_{10}(p)$
8
6
4
2
0
Chr.1 (Mb)
11
16.5
22
27.5
33
38.5
44
49

#### S__Endo

```
## Warning in max(x[m <= 0.05]): no non-missing arguments to max; returning -Inf

## Warning in max(x[m <= 0.05]): no non-missing arguments to max; returning -Inf

##         BLINK      FarmCPU          GLM          MLM         MLMM
## 5.648924e-07         -Inf 5.436219e-07         -Inf 1.324425e-06

##  Rectangular-Manhattan Plotting BLINK.
```

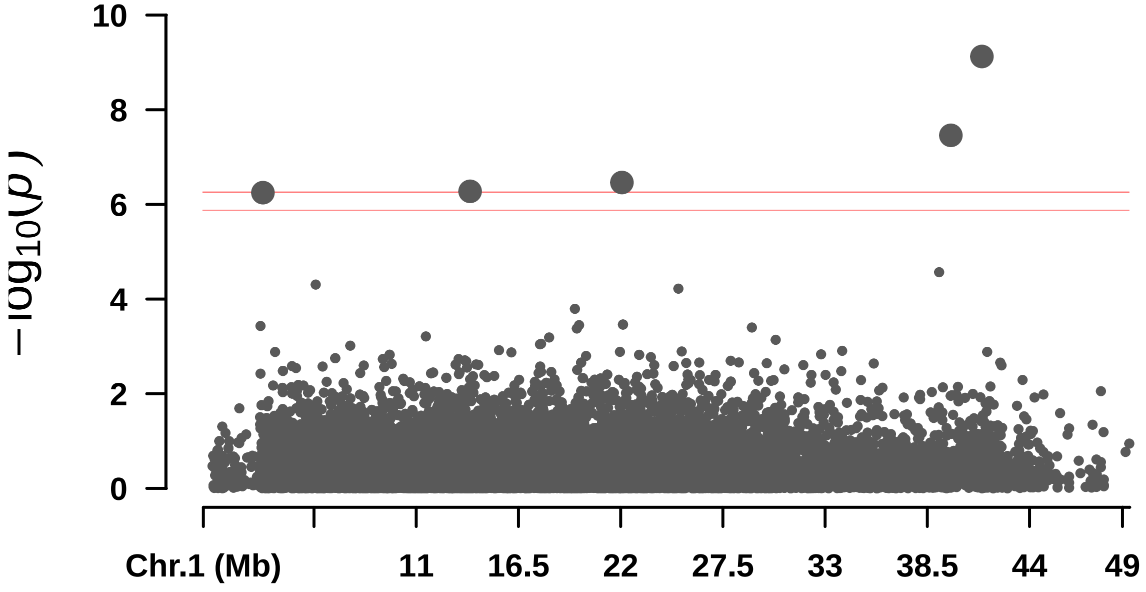


```
##  Rectangular-Manhattan Plotting FarmCPU.
```

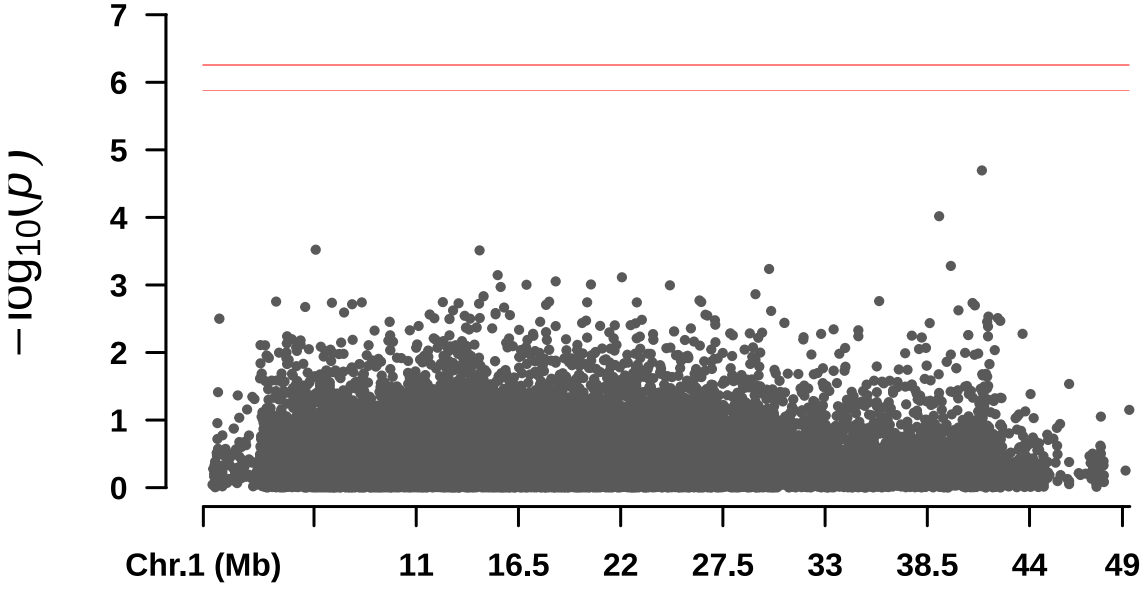


```
##  Rectangular-Manhattan Plotting GLM.
```

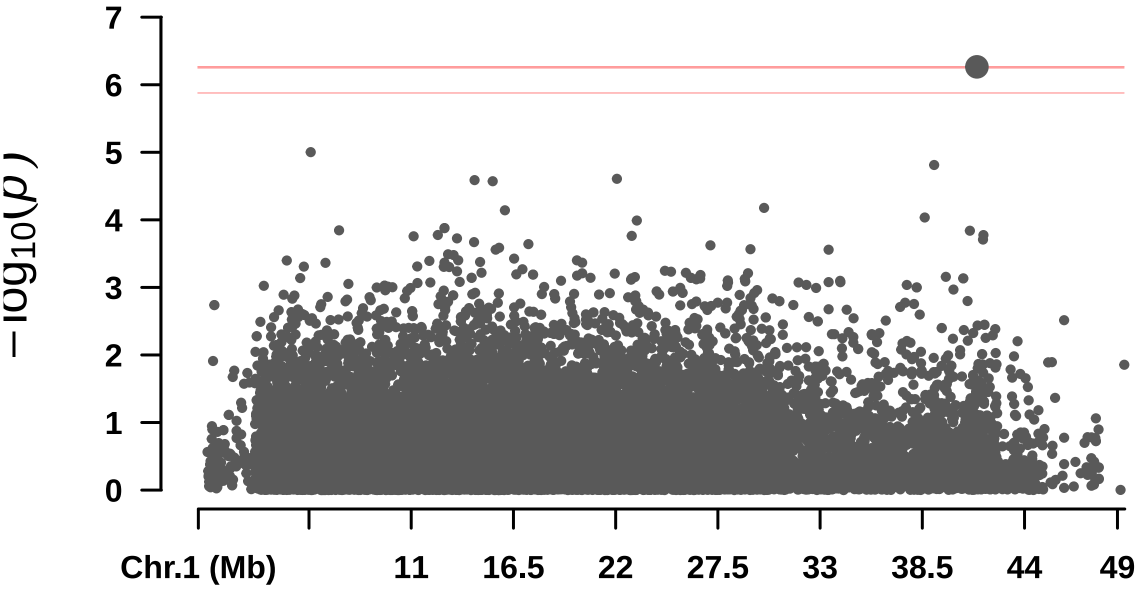


```
##  Rectangular-Manhattan Plotting MLM.
```

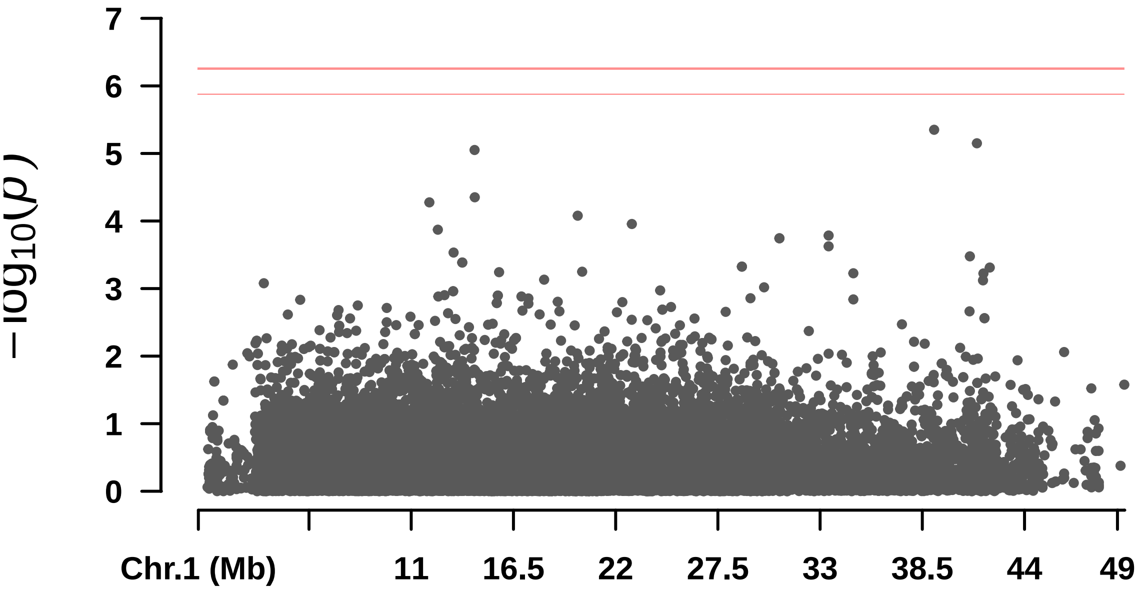


```
##  Rectangular-Manhattan Plotting MLMM.
```

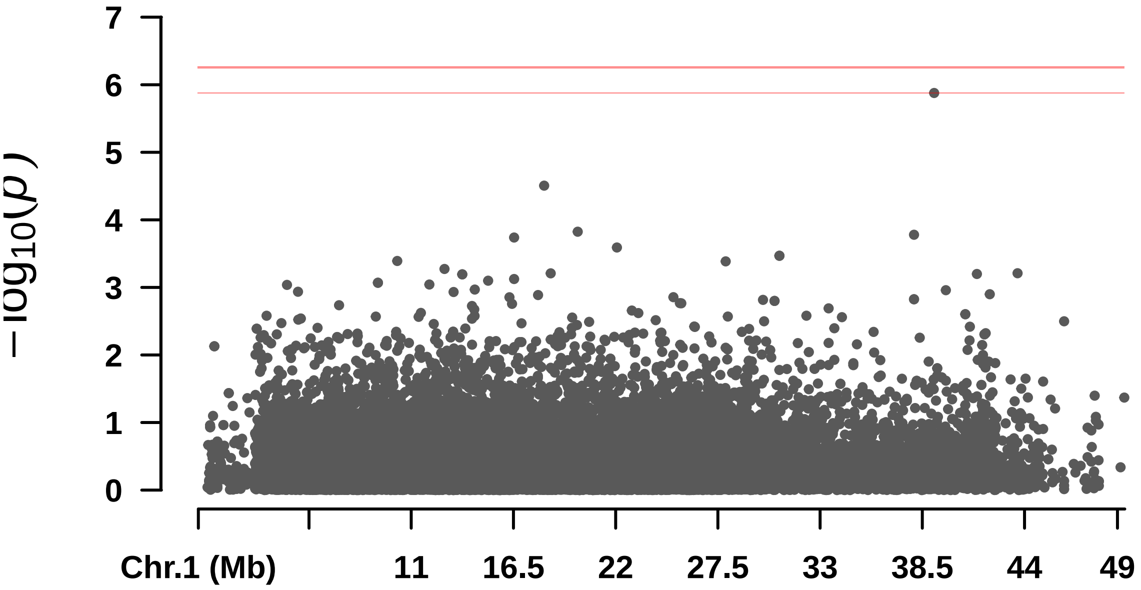

$-\log_{10}(p)$
7
6
5
4
3
2
1
0
Chr.1 (Mb)
11
16.5
22
27.5
33
38.5
44
49

#### S_Polpa

```
##        BLINK      FarmCPU          GLM          MLM         MLMM
## 1.042425e-07 3.767452e-06 1.085381e-04 7.675832e-06 1.377769e-06
```

```
##  Rectangular-Manhattan Plotting BLINK.
```

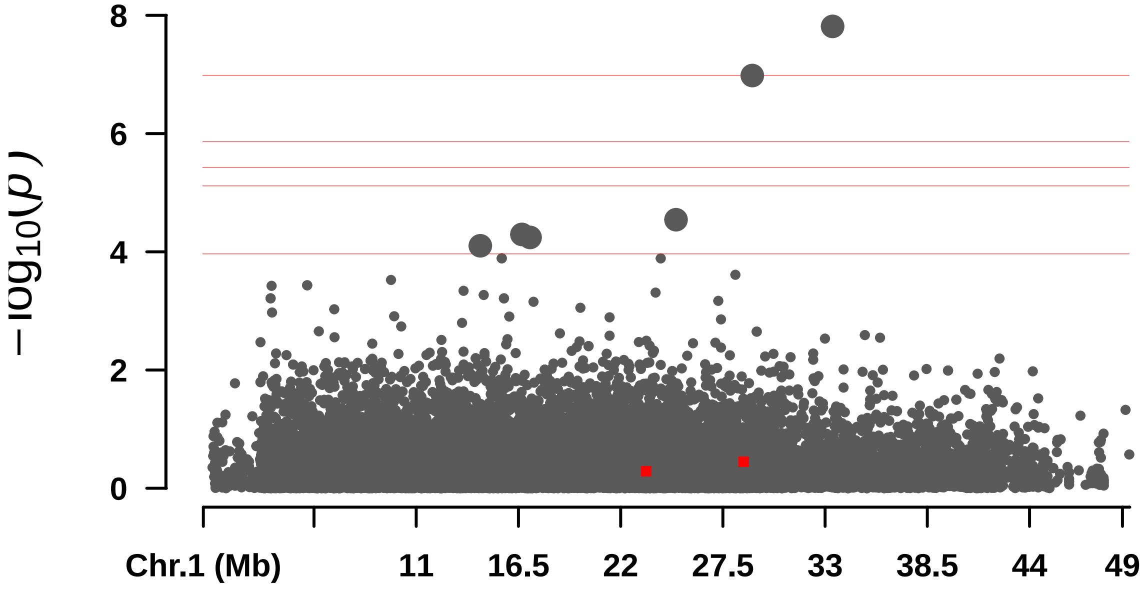


```
##  Rectangular-Manhattan Plotting FarmCPU.
```

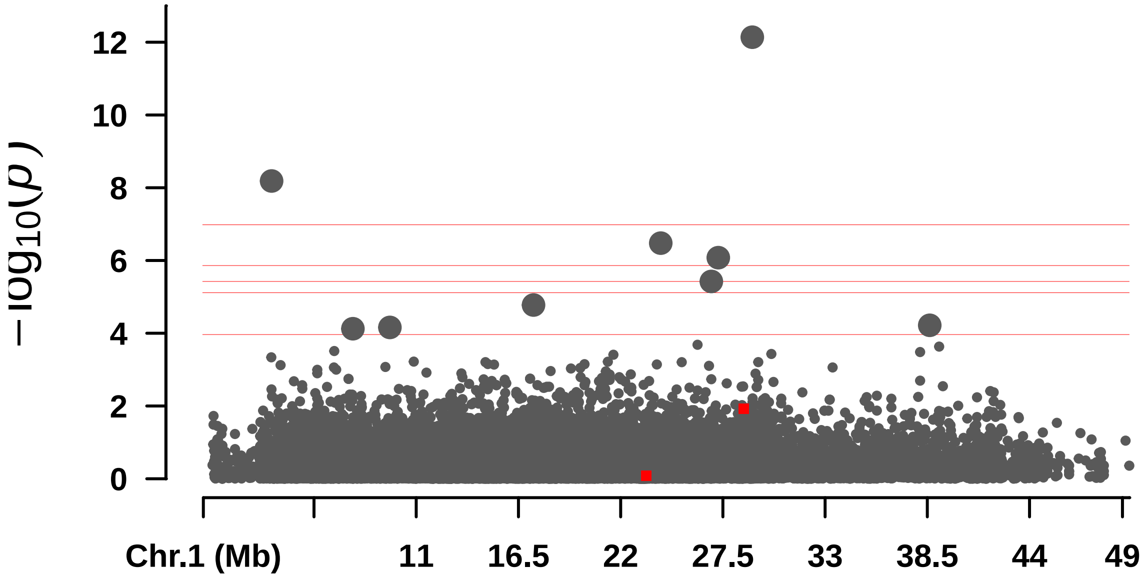


```
##  Rectangular-Manhattan Plotting GLM.
```

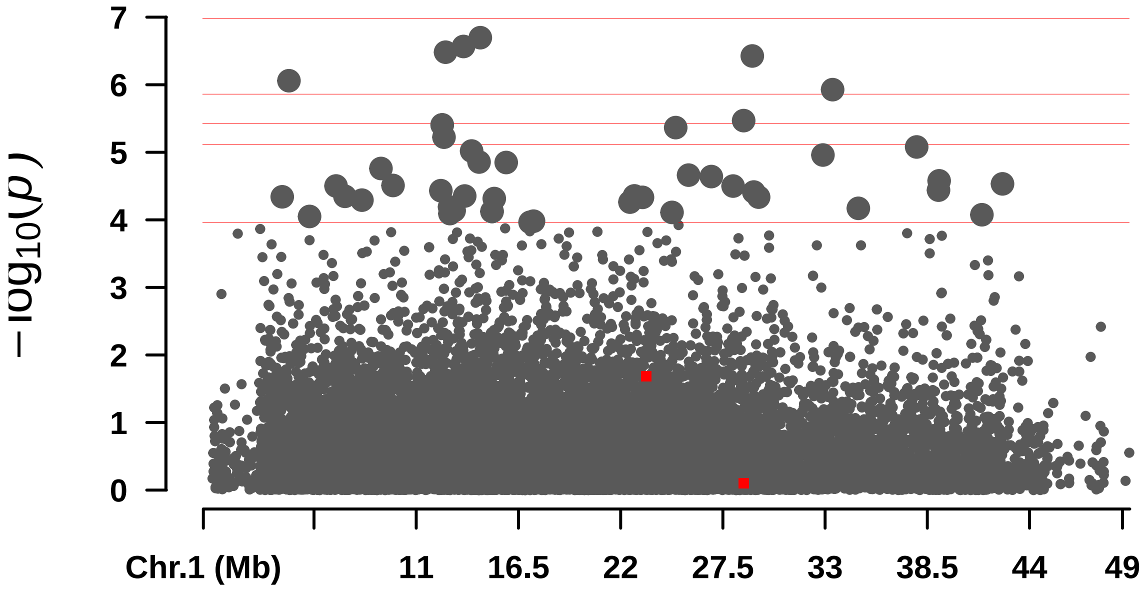


####  Rectangular-Manhattan Plotting MLM.

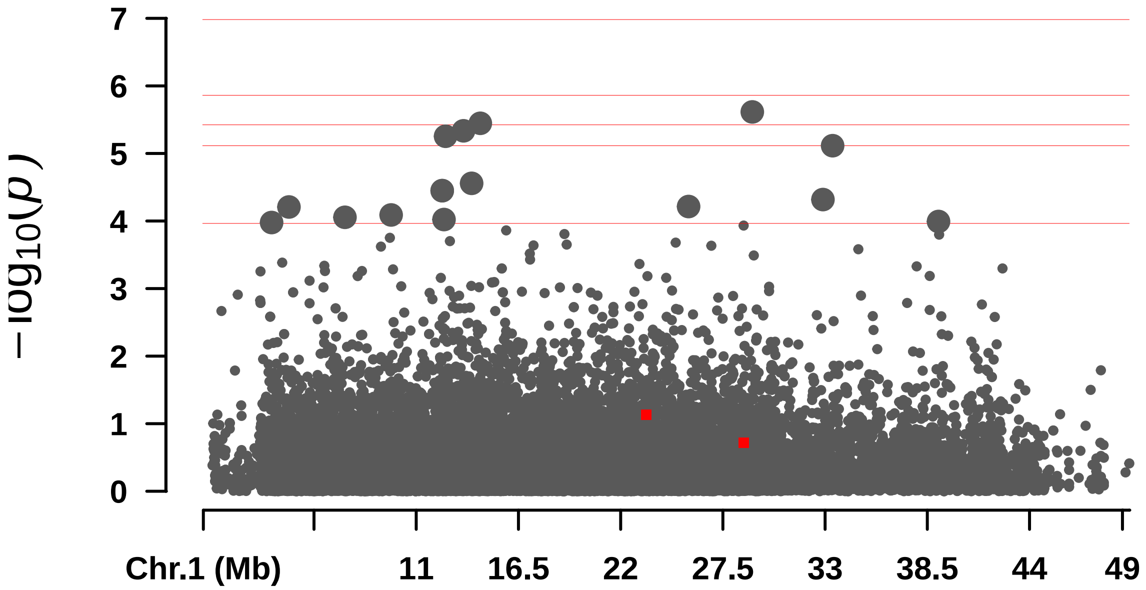


####  Rectangular-Manhattan Plotting MLMM.

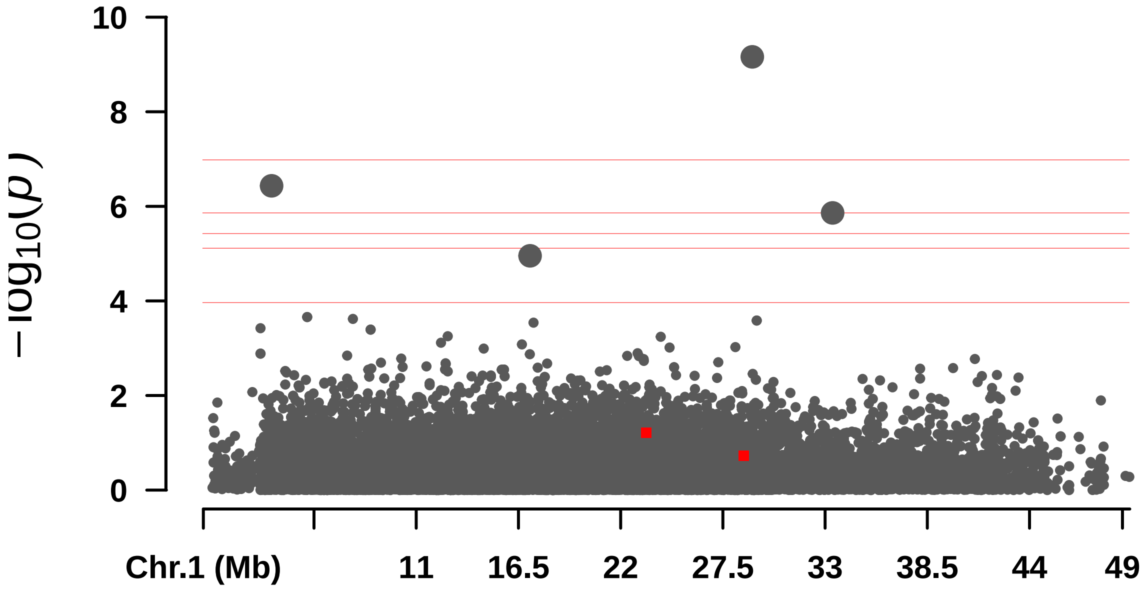

10
8
6
4
2
0
$-\log_{10}(p)$
Chr.1 (Mb)
11
16.5
22
27.5
33
38.5
44
49

—

#### S_Sem

```
## numeric(0)
```

```
##  Rectangular-Manhattan Plotting BLINK.
```

```
##  Rectangular-Manhattan Plotting FarmCPU.
```

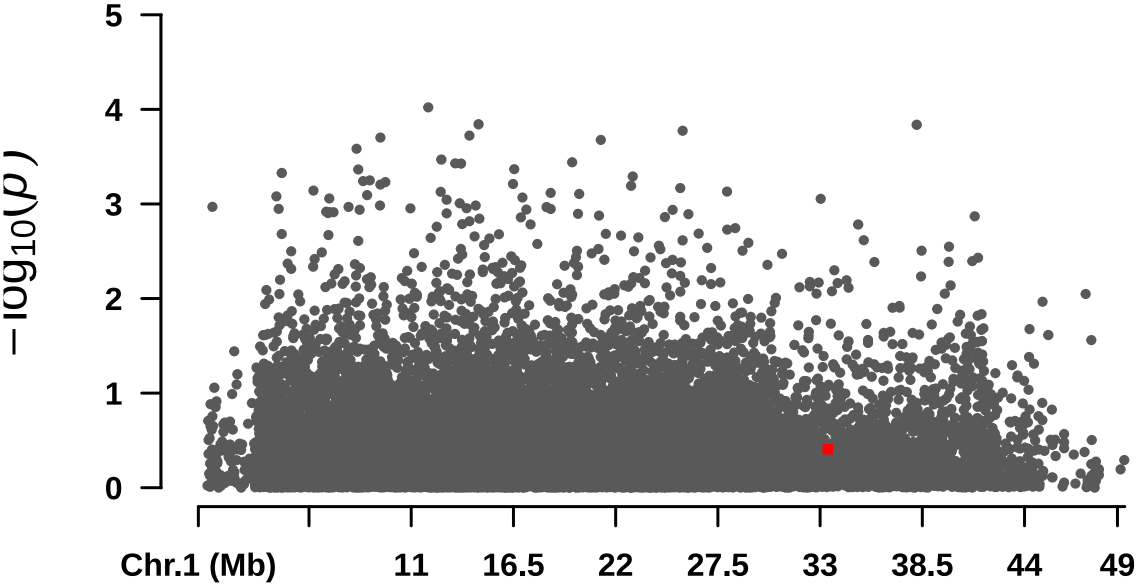


```
##  Rectangular-Manhattan Plotting GLM.
```

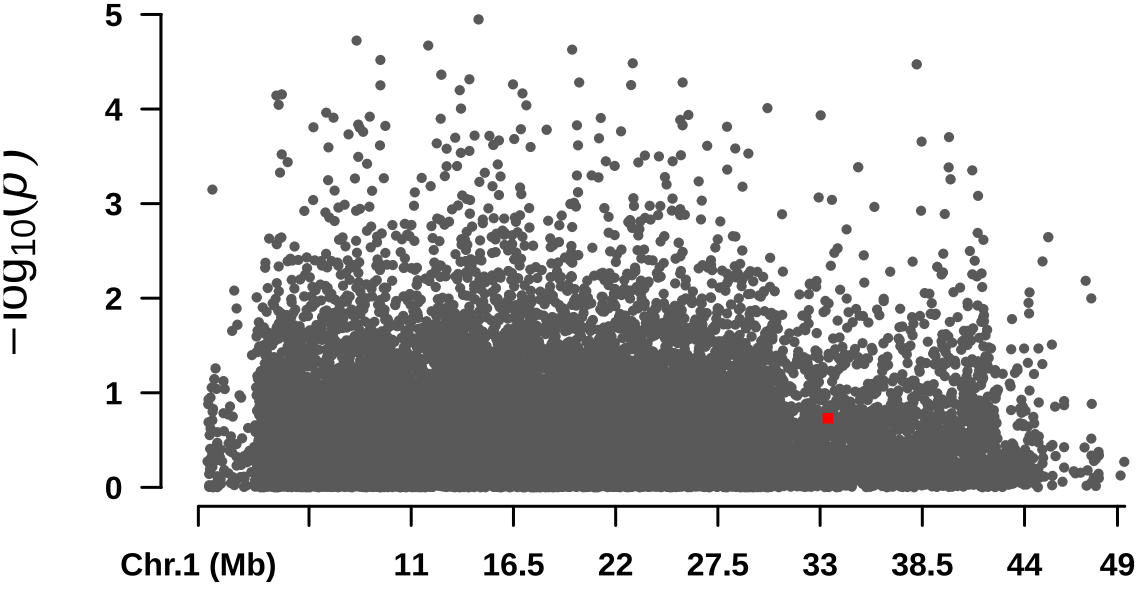


```
##  Rectangular-Manhattan Plotting MLM.
```

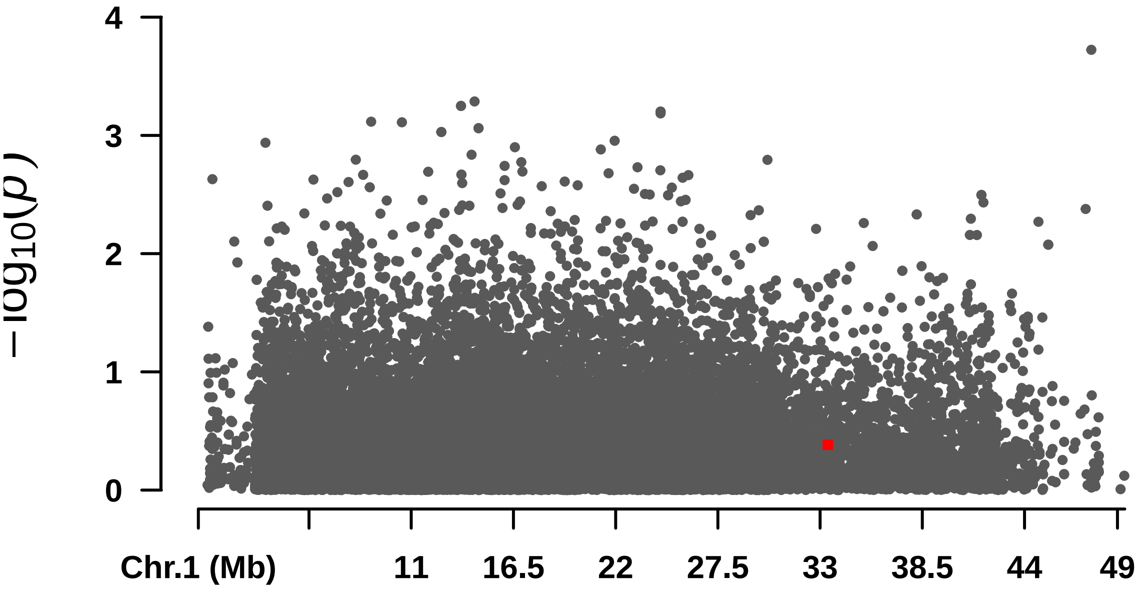


```
##  Rectangular-Manhattan Plotting MLMM.
```

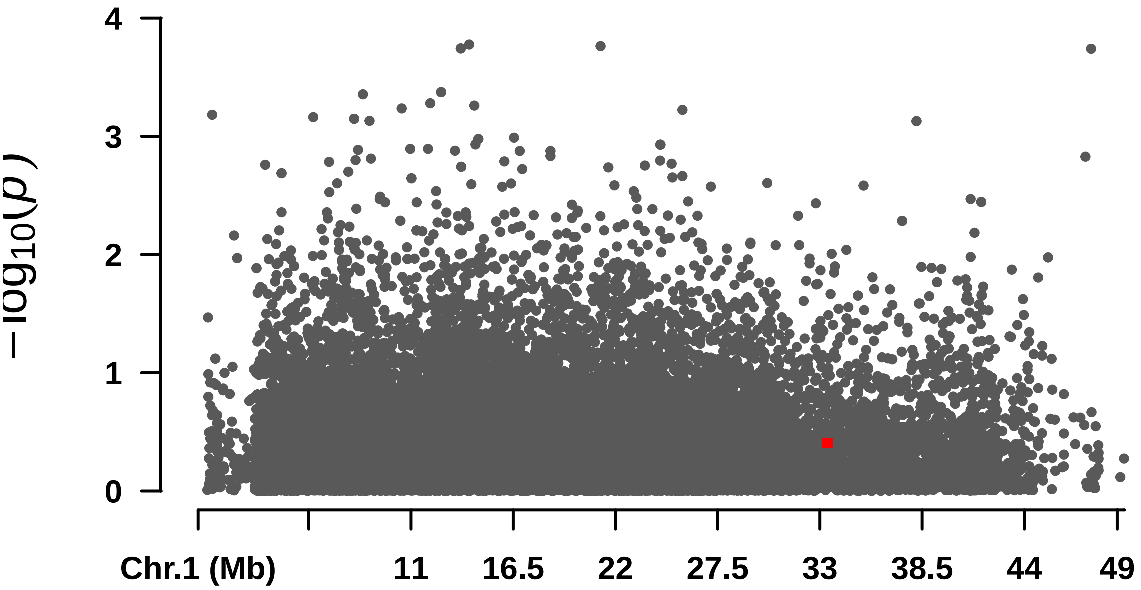

—

## S_T

```
## Warning in max(x[m <= 0.05]): no non-missing arguments to max; returning -Inf

##        BLINK      FarmCPU          GLM          MLM         MLMM
## 2.321189e-06 1.168518e-05 2.102759e-05         -Inf 1.348660e-08

##  Rectangular-Manhattan Plotting BLINK.
```

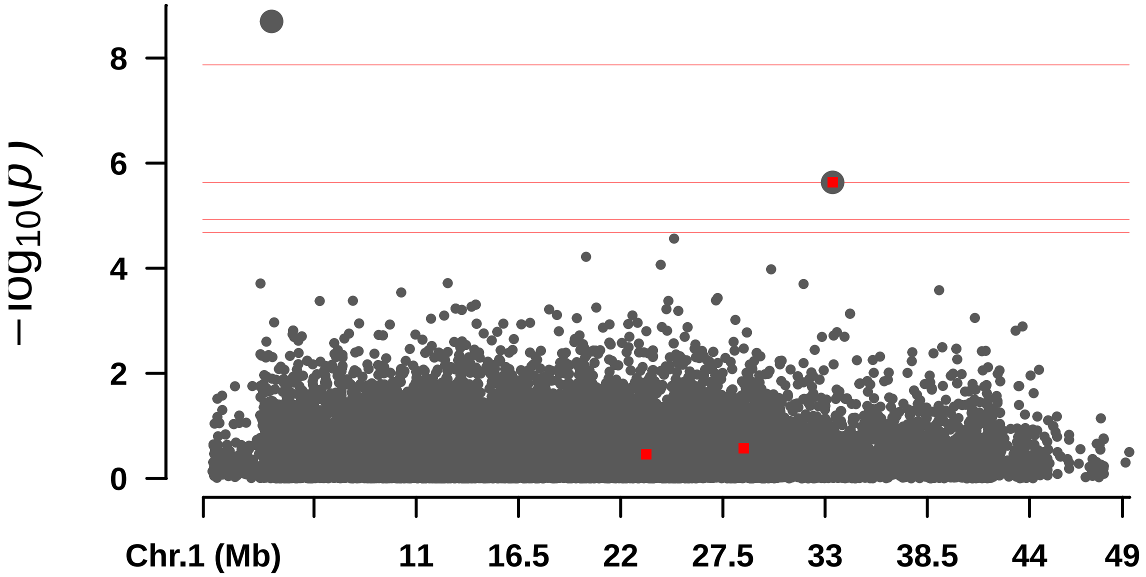


```
##  Rectangular-Manhattan Plotting FarmCPU.
```

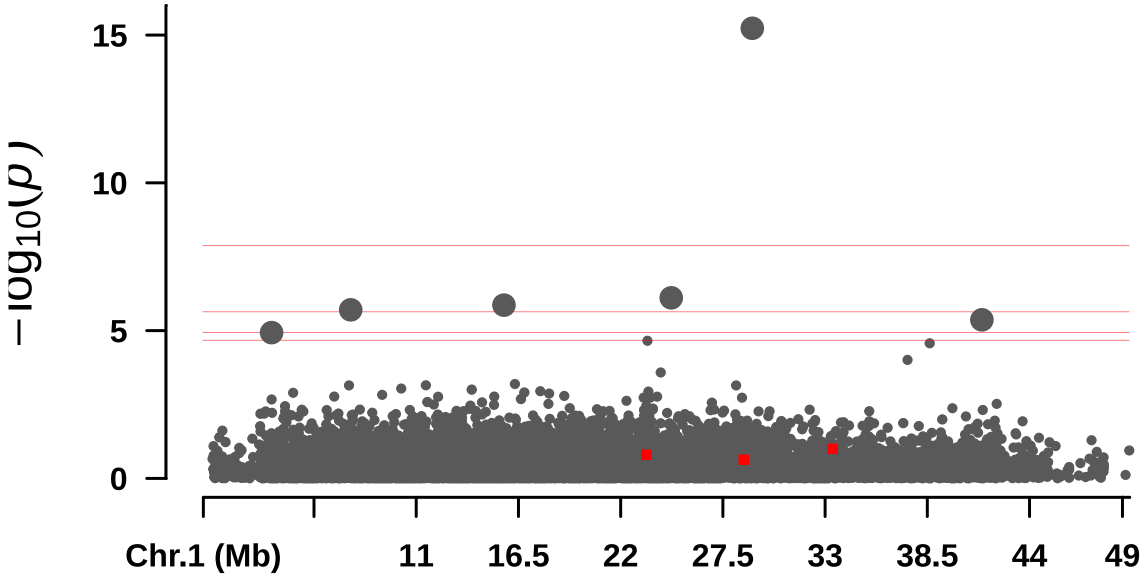


```
##  Rectangular-Manhattan Plotting GLM.
```

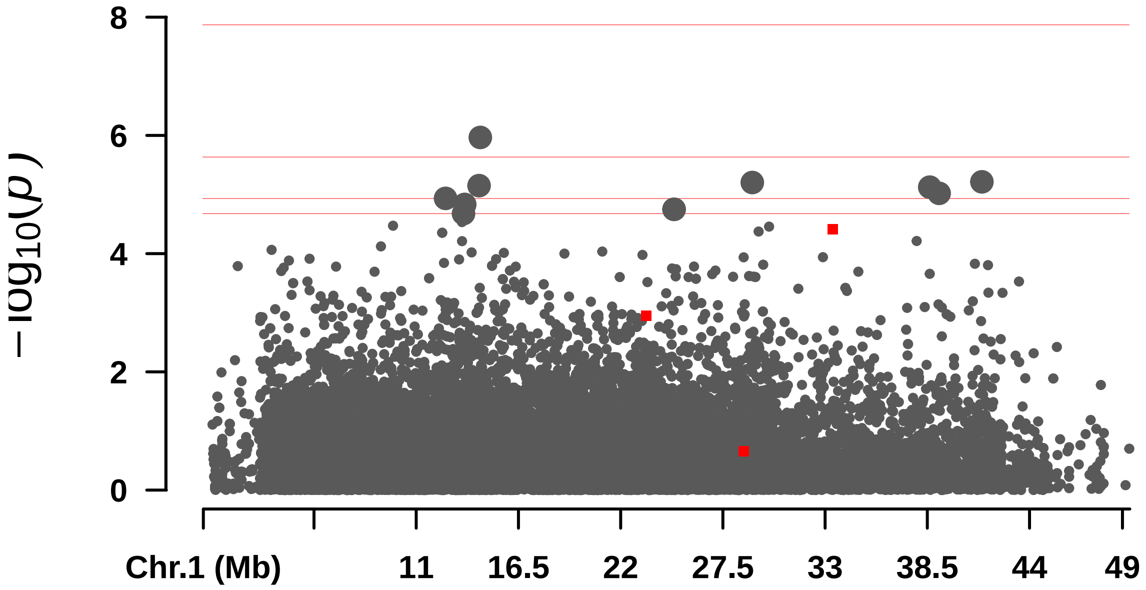


```
##  Rectangular-Manhattan Plotting MLM.
```

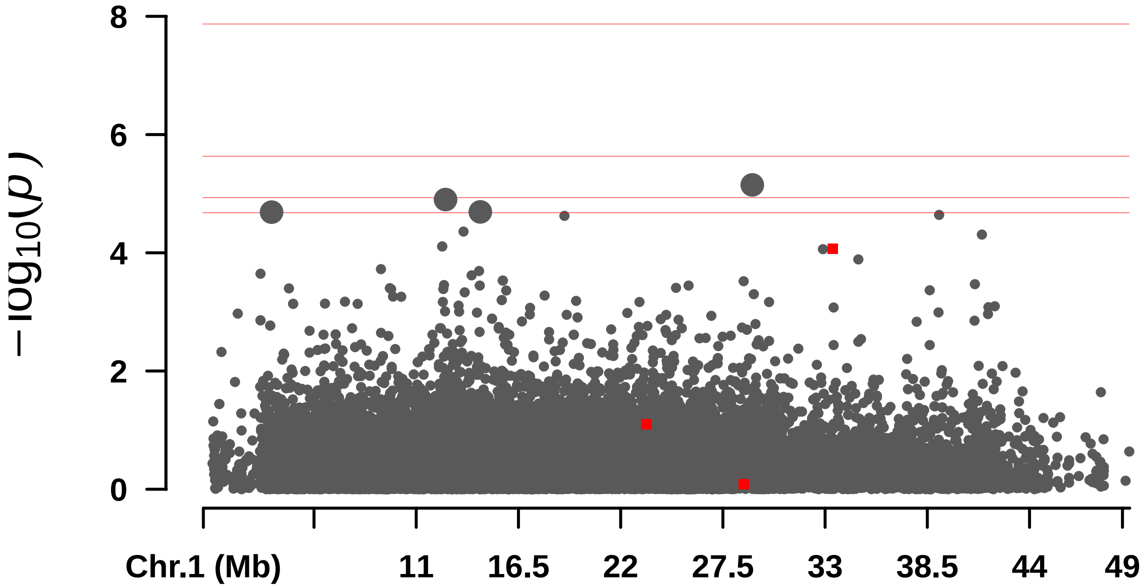


```
##  Rectangular-Manhattan Plotting MLMM.
```

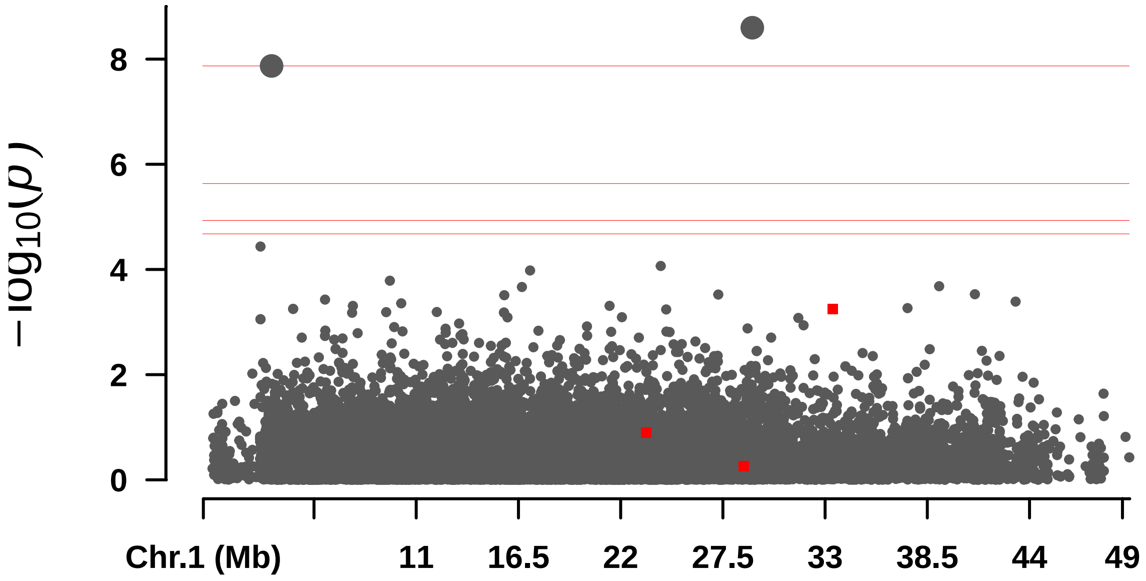

8
6
4
2
0
$-\log_{10}(p)$
Chr.1 (Mb)
11
16.5
22
27.5
33
38.5
44
49
